# Sex allocation conflict and sexual selection throughout the lifespan of eusocial colonies

**DOI:** 10.1101/454512

**Authors:** Piret Avila, Lutz Fromhage, Laurent Lehmann

## Abstract

Models of sex allocation conflict are central to evolutionary biology but have mostly assumed static decisions, where resource allocation strategies are constant over colony lifespan. Here, we develop a model to study how the evolution of dynamic resource allocation strategies is affected by the queen-worker conflict in annual eusocial insects. We demonstrate that the time of dispersal of sexuals affects the sex allocation ratio through sexual selection on males. Furthermore, our model provides three predictions that depart from established results of classic static allocation models. First, we find that the queen wins the sex allocation conflict, while the workers determine the maximum colony size and colony productivity. Second, male-biased sex allocation and protandry evolve if sexuals disperse directly after eclosion. Third, when workers are more related to new queens, then the proportional investment into queens is expected to be lower, which results from the interacting effect of sexual selection (selecting for protandry) and sex allocation conflict (selecting for earlier switch to producing sexuals). Overall, we find that colony ontogeny crucially affects the outcome of sex-allocation conflict because of the evolution of distinct colony growth phases, which decouples how queens and workers affect allocation decisions and can result in asymmetric control.

## Introduction

Eusocial Hymenopteran colonies may superficially appear to function as single organisms, where queens and workers could be viewed as the germinal and somatic tissues of multicellular organisms (Macevicz and Oster, 1976). However, such individuals are usually not clonal, whereby some genes, for instance those influencing sex allocation or reproductive ability of workers, can experience diverging selection pressures in different individuals (e.g. Bourke and Franks, 1995; Haig, 2003; Hamilton, 1967; Ratnieks et al., 2006).

One of the most intensively studied genetic conflicts is the queen-worker conflict over sex allocation. In an outbred haplodiploid population where each colony is headed by a singly-mated queen, natural selection on resource allocation strategies favors alleles in queens that code for equal resource allocation to males and (sexual) females and alleles in workers that code for a 3:1 (sexual females to males) allocation ratio (e.g. Frank, 1998; Trivers and Hare, 1976; West, 2009). Factors such as multiple related queens per colony and multiple matings by the queen, reduce the extent of the genetic conflict over sex allocation because they reduce relatedness asymmetries between individuals within colonies (e.g. Frank, 1998; Ratnieks et al., 2006; West, 2009).

The long-term evolutionary “outcome” of the sex allocation conflict – the uninvadable resource allocation schedule, is determined by the mechanisms through which the opposing “parties” can influence how colony resources are allocated into producing individuals of different types. In a colony founded by a single queen there are two opposing parties: the genes in the workers and the genes in the colony-founding queen. The genetic control over resource allocation decisions can be achieved through different genetic, behavioural, and physiological processes (Beekman and Ratnieks, 2003; Helanterä and Ratnieks, 2009; Mehdiabadi et al., 2003). Hereinafter, if one party fully determines a given resource allocation trait, then this party is said to be “in control” of that trait (here, “in control” has a related but more restricted meaning than “having power” as in e.g. Beekman and Ratnieks, 2003). In general, there are reasons to expect that the genes in the queen and workers simultaneously control different resource allocation decisions, because both parties are known to have means to control different resource allocation decisions and selection for a party to seize control over a resource allocation decision can be strong if there are means to do so (Bourke and Franks, 1995; Helanterä and Ratnieks, 2009; Trivers and Hare, 1976). Furthermore, it is often considered most likely that the genes in the queen determine the primary sex allocation ratio (allocation of resources to females versus males) and the workers control the developmental fate of the female eggs (Bourke and Franks, 1995; Helanterä and Ratnieks, 2009; Trivers and Hare, 1976). Hereinafter, we refer to this scenario as “mixed control”.

Theoretical models of sex allocation conflict provide three important insights into fundamental questions in evolutionary biology (e.g. Bourke and Chan, 1999; Bourke and Ratnieks, 1999; Pamilo, 1991a; Pen and Taylor, 2005; Reuter et al., 2004; Reuter and Keller, 2001). Firstly, they provide clear predictions that allow to test how relatedness affects selection on social traits (Crozier and Pamilo, 1996). Secondly, they allow to predict which party is in control of the underlying resource allocation decisions, given that one has sex-allocation data. Thirdly, they enable to predict to what extent the conflicts can be “resolved” (sensu Ratnieks et al., 2006, i.e. conflict outcome with modest colony-level costs) under various assumptions about the mechanisms of genetic control over the resource allocation decisions. However, all of the aforementioned models consider static allocation decisions without explicitly taking colony ontogeny into account. Nevertheless, it is known that many annual eusocial insect species (e.g. vespid wasps, bumble bees, and sweat bees) grow in two distinct phases (see references in Mitesser et al. (2007a), Crone and Williams (2016)), such that, in the beginning of the season only workers are produced, i.e. the ergonomic phase followed by a drastic shift into exclusive production of sexuals (males and future queens), i.e. the reproductive phase. This life-history schedule was shown to be an evolutionary outcome in annual eusocial colonies assuming clonal reproduction by Macevicz and Oster (1976). However, only a few theoretical studies (Bulmer, 1981; Ohtsuki and Tsuji, 2009) have considered sexually reproducing species (thereby including the possibility of genetic conflicts) and time-dependent resource allocation decisions in the context of colony life-history. The importance of colony ontogeny in studying within-colony conflict was demonstrated by Ohtsuki and Tsuji (2009) who showed (in the context of worker policing) that the expression of conflict depends on the phase of colony ontogeny.

In his seminal work, Bulmer (1981) showed using a dynamic allocation model (i.e., time-dependent decisions) that the sex allocation conflict can have a detrimental effect on colony productivity (sexual biomass) under mixed control since relatively few resources are allocated into producing workers. Indeed, he predicted that the production of workers is expected to halt earlier under mixed control, but he did not consider the entire colony ontogeny and his predictions relied on some additional restrictive assumptions. For example, he assumed that the worker generations do not overlap within a season (i.e. a colony grows in separate generation of workers within a season) and that sexuals can only mate at the very end of the season. Hence, theoretical understanding of the life-history decisions of eusocial colonies has mostly relied on the assumption of clonal reproduction with no genetic conflicts (Macevicz and Oster, 1976; Mitesser et al., 2007a).

The importance of considering dynamic resource allocation decisions for studying within-colony conflict is demonstrated by the fact that the static and dynamic resource allocation models can make contradicting predictions about which party wins the sex allocation conflict under mixed control (Bulmer, 1981; Reuter and Keller, 2001). Indeed, the static resource allocation model by Reuter and Keller (2001) predicts a sex allocation ratio under mixed control that is intermediate between the evolutionary prediction corresponding to worker and queen control. In contrast, Bulmer’s (1981) dynamic model predicts that the queen wins the sex allocation conflict by laying only haploid eggs at the penultimate generation causing the colony to die one generation before the end of the season if the sex allocation ratio in the population is female-biased. However, the generality of Bulmer’s predictions are limited due to the aforementioned restrictive assumptions of his model.

Furthermore, in another study assuming queen control of resource allocation traits and the possibility of sexuals to mate before the end of the season, Bulmer (1983) showed that sexual selection on males will lead to protandry (males being produced before sexual females) if mating can occur over some period of time. Indeed, sexual selection may thus play an important role for colony ontogeny, since protandry is found among many annual eusocial insects, e.g. in paper wasps and bumble bees (Bourke, 1997; Strassmann and Hughes, 1986). Evolution of protandry however contradicts the earlier model by Bulmer (1981) for mixed control, since it predicted that males are produced in the very end of the season. Hence, there are no theoretical predictions for time-dependent colony resource allocation decisions and conflicts under mixed control, where individuals can mate over a finite period of time during the season with sexual selection occurring throughout.

In this paper, we address the limitations of previous studies by developing a dynamic resource allocation model where we consider three alternative scenarios of genetic control of resource allocation decisions: queen control, worker control, and mixed control; and two alternative scenarios of dispersal of sexuals: delayed dispersal (all sexuals simultaneously disperse at the end of the season to mate) and direct dispersal (sexuals disperse immediately after eclosion to mate). In light of previous work, the purpose of this paper is to address the following questions: (i) How does conflict affect colony growth? (ii) How does sexual selection affect the order at which sexuals are produced? (iii) Which party wins the sex allocation conflict for different scenarios of dispersal of sexuals?

## Model

### Life-cycle

We consider a seasonal population of haplodiploid eusocial insects consisting of a large (ideally infinite) number of colonies or breeding sites each occupied by a mated queen. The life cycle over a season is assumed to consist of the following four events. (1) *Reproduction*: at the start of the season of total length *T*, each queen occupying one of the *n* breeding sites initiates a colony that can grow throughout the season, and where workers, males, and future queens can be produced. (2) *Dispersal*: sexuals disperse out of their natal colony, such that no inbreeding, local mate competition, or local resource competition takes place; we consider two alternative scenarios for the timing of dispersal (to be detailed below). (3) *Mating*: random mating occurs and all queens mate exactly with *M* ≥ 1 males. (4) *Regulation*: all individuals die at the end of the season, except (juvenile) queens who randomly compete for the *n* breeding slots to initiate colonies of the next generation.

### Dispersal and mating

The two dispersal scenarios are as follows: (i) delayed dispersal, where sexuals all disperse at the same time at the end of the season, and (ii) direct dispersal, where sexuals disperse immediately after being produced. Females mate immediately with *M* males in the mating pool after which they will exit the mating pool. In contrast, males continue on mating until they die. Hence, the mating success of a male depends on his mortality rate and the availability of mating females. In order to gain fitness, females have to survive until the end of the season, while males have to inseminate females who survive until the end of the season.

### Colony growth and production of sexuals

We model explicitly colony population dynamics during stage (1) of the life cycle. To describe our model, we start by considering that the population is monomorphic for all phenotypes, and will later introduce variation and selection. The size of a focal colony at time *t* ∈ [0, *T*] in the (monomorphic) population is *y*_w_(*t*), which gives the number of sterile workers (including the colony founding queen, who has been counted as a worker) in the colony at time *t*. In addition, by time *t*, the colony has produced *y*_q_(*t*) surviving (juvenile) queens and *y*_m_(*t*) surviving (juvenile) males. By the term “juvenile” we only want to emphasize that these sexual individuals are regarded as offspring in the current generation and that they will reproduce in the next generation. For simplicity, we assume that all individuals are equally costly to produce, which allows to equate the investment allocation ratio to the numerical sex ratio. However, the assumption of equal production cost has no fundamental effect on the evolutionary process, since selection acts only on total investment in the sexes and not on their numbers and hence is independent of the production costs of different individuals (West, 2009).

Workers acquire resources from the environment to produce offspring. Let *b* denote the individual productivity rate of a worker (i.e. the net rate at which a worker acquires resources for the colony, measured in individuals produced per unit time). For simplicity, we assume that the availability of resources in the environment is constant over time and the rate at which resources are acquired scales linearly with the colony size (i.e. *b* is constant). The latter assumption implies that there are enough resources in the environment to sustain constant per-worker rate of resource acquisition and the egg-laying rate of the queen is constrained only by the resources available to the colony.

The number *y*_*k*_(*t*) of type *k* ∈ {w, q, m} individuals alive at time *t* that were produced in the focal colony is assumed to change according to

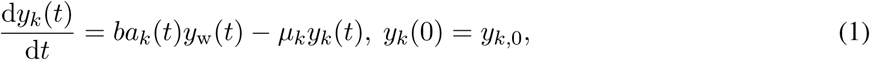

where *a*_*k*_(*t*) is the fraction of resources allocated into producing type *k* individuals at time *t, µ*_*k*_ is the mortality rate of individuals of type *k*, and *y*_*k*,0_ is the number of type *k* individuals in the colony in the beginning of the season. The initial condition (number of individuals at the beginning of the season) for the colony is *y*_w,0_ = 1 (the colony founding queen is counted as a worker, since she can for example recover some resources from her body fat), *y*_q,0_ = 0 (no juvenile queens), and *y*_m,0_ = 0 (no juvenile males). Note that the number of juvenile queens *y*_q_(*t*) and males *y*_m_(*t*) are counted regardless if they have dispersed from the colony or not.

It will turn out to be useful to keep track of the number of queens that the males from a focal colony have inseminated. Let *y*_iq_(*t*) be the expected number of females alive at time *t*, who have been inseminated by males from a focal colony, given that females mate only once (i.e. under a monandrous mating system, *M* = 1) and it changes according to

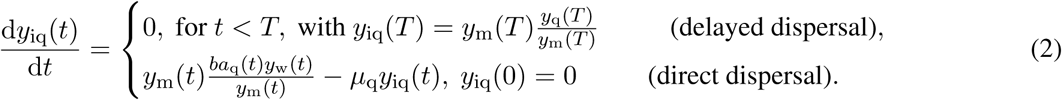

Under delayed dispersal, all females are inseminated at time *t* = *T*, where a total number of *ny*_m_(*T*) males compete for *ny*_q_(*T*) females. Hence the mating success of a male produced in a focal colony is *y*_q_(*T*)/*y*_m_(*T*), and the number of males in that colony at the end of the season is *y*_m_(*T*). Under direct dispersal, females mate immediately after being produced, whereby at time *t* a total number of *nba*_q_(*t*)*y*_w_(*t*) females are available to mate (after which they will leave the mating pool). In contrast, males stay in the mating pool, hence at time *t*, an average number of *ny*_m_(*t*) males compete for the access to females. Therefore, the mating rate of a male produced in a focal colony is *ba*_q_(*t*)*y*_q_(*t*)/*y*_m_(*t*) at time *t* and the last term in the second line of eq. (2) takes into account the mortality of the inseminated females. If females mate *M* times, then there are on average *M* times more matings available to males at any given time. Hence, the number of (surviving) females at time *t*, who have been inseminated by males from a focal colony is *My*_iq_(*t*) in a population where females mate *M* times.

### Resource allocation traits

We assume that the allocation schedule, *a*_*k*_(*t*) (*k* ∈ {w, q, m}), that governs the dynamics of individuals produced in the focal colony (recall eq. 1), is controlled by two traits

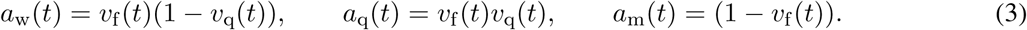

The first trait 0 ≤ *v*_f_ (*t*) ≤ 1 is the proportion of resources allocated to producing females (individuals destined to become workers or queens) at time *t*. The second trait 0 ≤ *v*_q_(*t*) ≤ 1, gives the proportion of resources allocated to producing queens from resources allocated to females at time *t* ∈ [0, *T*]. Thus (1 − *v*_f_ (*t*)) is the proportional allocation to males and (1 − *v*_q_(*t*)) is the proportional allocation of resources directed to producing workers from resources allocated to females.

Our aim is to investigate the evolution of the resource allocation schedule during the whole colony ontogeny, i.e., the evolution of **v** = {*v*_f_ (*t*), *v*_q_(*t*)}_*t*∈[0,*T*]_. In species where workers are sterile (as assumed here) the queen is often thought to control the allocation between females and males (trait *v*_f_) because she decides at which rate she lays female and male eggs. However, the genes in the workers can influence *v*_f_, if they are able to redirect resources from male brood to female brood (Chapuisat et al., 1997; Sundström et al., 1996), but for simplicity we do not consider this scenario in our paper. In many species, the genes in the workers control the developmental fate of the female larvae (trait *v*_q_) by differential feeding, as the diet provided to the larvae by workers determines the caste of the female offspring (Berens et al., 2015; Ratnieks et al., 2006; Schwander et al., 2010). However, in some species, queens can also alter the caste determination of females by producing different types of diploid eggs (Wheeler, 1986). It is believed that in many eusocial insects, the queen and the workers are in control of different resource allocation decisions simultaneously and it is often considered most likely that the queen determines the primary sex ratio (ratio of female to male eggs), while the workers control the developmental fate of the female eggs (Bourke and Franks, 1995; Helanterä and Ratnieks, 2009; Trivers and Hare, 1976). Hence, in light of the empirical evidence of genetic control of resource allocation decisions, we will examine three possible scenarios of genetic control over these traits: queen control (i.e. the genes in the queen determine resource allocation decisions), worker control (i.e. the genes in the queen determine resource allocation decisions) and mixed control, where the genes in the queen control *v*_f_ (the proportional investment into females versus males) and the genes in the workers control *v*_q_ (the proportional investment into new queens versus workers). Our assumptions of the genetic control are in accordance with the corresponding assumptions of the static resource allocation model by Reuter and Keller (2001), where they also considered these three scenarios with the corresponding static traits.

In order to analyse the long-term evolution of the resource allocation traits, we perform an evolutionary in-vasion analysis (see section 1 of S.I. for more information). That is, we consider the fate (invasion or extinction) of a single mutant allele (an allele determines the entire allocation schedule, i.e., a trajectory of the trait over *t* ∈ [0, *T*]) introduced into a population of resident individuals and ask what is the (candidate) uninvadable allo-cation schedule 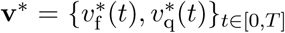 namely, the allocation schedule resistant to invasion by any mutant schedule that deviates from **v**\*. We determine the (candidate) uninvadable allocation schedule **v**\* analytically using Pontryagin’s maximum principle (see sections 3–6 of S.I.), which gives a necessary condition for optimality, and we confirm these results numerically using GPOPS–II (Patterson and Rao, 2014), which gives support to the attainability of the uninvadable schedules (see section 11 in S.I.).

## Results

### Marginal value, relatedness asymmetry, and potential for conflict

#### Dynamic marginal value result

Consider a mutant allocation schedule **u** = {*u*_f_ (*t*), *u*_q_(*t*)}_*t*∈[0,*T*]_ that deviates slightly from a candidate uninvadable schedule **v**\*, such that a trait *u*_*τ*_ (*t*) (*τ* ∈ {f, q}) can be expressed as

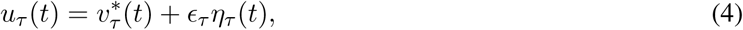

where *η*_*τ*_ (*t*) is a feasible phenotypic deviation from the resident trait 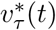 and *ϵ*_*τ*_ ≪ 1 scales the magnitude of this variation. By a feasible phenotypic deviation we mean any deviation *η*_*τ*_ (*t*) such that the mutant strategy *u*_*τ*_ (*t*) satisfies the constraints of the model (i.e., 0 ≤ *u*_*τ*_ (*t*) ≤ 1 for all *t* ∈ [0, *T*], e.g., see Sydsæter et al., 2008, p. 129 and 308).

Let us now denote by *y*_*k*_(**u**) *≡ y*_*k*_(*T*) the number of type *k* ∈ {q, iq} individuals at the end of the season where the resident allocation schedule **v** in eqs. (1) and (2) has been replaced by the mutant allocation schedule **u**. Then, the first-order condition for a schedule **v**\* to be uninvadable when party *c* ∈ {q, w} is in control of the trait of type *τ* ∈ {f, q} can be written as

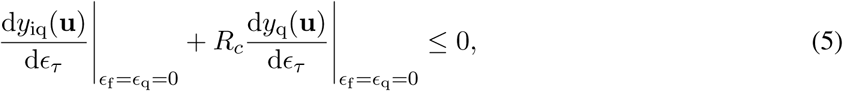

which has to hold for all feasible phenotypic deviations *η*_*τ*_. For mixed control inequality (5) must hold simultaneously for each trait being under the control of the respective party (see sections 2.1–2.2 of S.I. for a proof). Here, d*y*_*k*_(**u**)/ d*ϵ*_*τ*_ is a Gâteaux derivative (a type of functional derivative, e.g., Troutman, 2012, p. 45–50, Luenberger, 1997, p. 171–178, see also section 2.1 of S.I.) measuring the change in the number of individuals *y*_*k*_(**u**) of type *k* ∈ {q, iq} produced by the end of the season in a mutant colony (and we here emphasized that this number depends on the whole allocation schedule, recall eqs. 1–2), due to the infinitesimal deviation *ϵ*_*τ*_ *η*_*τ*_ (*t*) of the trait of type *τ* throughout the entire season *t* ∈ [0, *T*]. Eq. (5) is not a strict equality, since the (pointwise) selection gradient does not vanish when a population evolves towards the boundary of the set of possible allocation strategies (e.g., when only workers are produced over some time span, see 2–3 of S.I. for more details, especially sections 2.3–2.4 for pointwise selection gradient and first-order condition).

The first-order condition (5) says that at the uninvadable state, the marginal (gene) fitness return (“marginal return” for short) from allocating more resources to male production (measured in the currency of inseminated queens) cannot exceed the marginal loss from allocating less resources to queen production weighted by *R*_*c*_, which is the so-called *relatedness asymmetry* (Boomsma and Grafen, 1991, p. 386) defined as

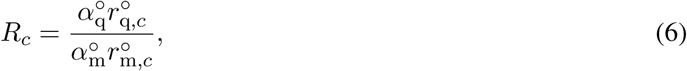

where 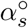 is the (neutral) reproductive value of all individuals of class *s* ∈ {q, m}, i.e., the probability that a gene taken in the distant future descends from an individual in class *s* ∈ {q, m} and 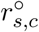 is the (neutral) coefficient of relatedness between an individual of type *s* ∈ {q, m} and an average individual whose genes are in control of the resource allocation trait. In section 2 of S.I. (eqs. S30–S32), we detail that the relatedness asymmetry can be interpreted as giving the ratio of sex-specific (queen/male) contributions, of genes in party *c*, to the gene pool in the distant future (under a neutral process). For haplodiploids the relatedness asymmetry is *R*_q_ = 1 (queen control) and *R*_w_ = (2 + *M*)/*M* (worker control).

Eq. (5) is a generalised formulation of Fisher’s (1930) theory of equal allocation (under queen control) and the standard static marginal value result of sex allocation theory (e.g., Taylor and Frank, 1996, eq. 22). The novelty of eq. (5) is that it results from a dynamic model, where the marginal return of producing an individual is time-dependent, and natural selection favours an allocation schedule that produces males and queens in such a way that the ratio of surviving inseminated queens and produced queens is equal to the relatedness asymmetry. Note that eq. (5) does not directly give the ratio of total amount of resources invested (“overall investment” ratio) in each sex, which depends on the characteristics of the life cycle. Furthermore, we show that the overall investment ratios can depart from classic static results of sex allocation theory under direct dispersal in our model.

#### Proportional relatedness asymmetry

It follows from the first-order condition that the marginal value result is given by the relatedness asymmetry, i.e. the ratio of sex-specific asymptotic contributions to the gene-pool (eq. 5). However, it will turn out to be useful to define the proportional contribution of genes of party *c* through queens to the future gene pool, i.e.

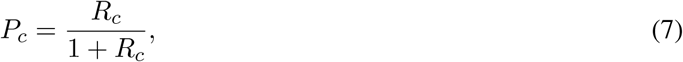

which can be thought of as a proportional relatedness asymmetry. This quantity evaluates to *P*_q_ = 1/2 (queen control) and *P*_w_ = (2+*M*)/(2(1+*M*)) (worker control), and it is equal to the (overall) uninvadable proportional allocation into females according to the classical static models of sex allocation theory under single-party control (Boomsma and Grafen, 1991; Reuter and Keller, 2001; Trivers and Hare, 1976).

The conflict between workers and the queen is absent when the proportional relatedness asymmetries for queens and males are equal, i.e. *P*_w_/*P*_q_ = 1. However, when *P*_w_/*P*_q_ *>* 1, then future queens are more valuable to workers than to the queen in contributing genes to the future gene pool. Hence, the ratio

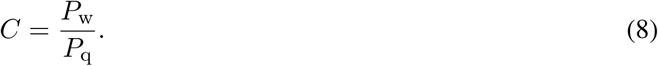

can be interpreted as the potential for conflict. In other words, whenever *C* ≠ 1, then there is potential for conflict between the queen and the workers over sex allocation. In haplodiploids, the potential for conflict *C* = *C*(*M*) = (2 + *M*)/(1 + *M*) decreases with the increase in polyandry *M* (Ratnieks and Boomsma, 1995), since *P*_w_ → *P*_q_ with the increase in queen mating frequency. Hence, the potential conflict *C*(1) = 1.5 is maximal when the queen mates once. It turns out that the proportional relatedness asymmetry *P*_*c*_ and the potential for conflict *C* are key quantities describing the properties of the uninvadable allocation schedule **u**\*, to which we next turn.

### The candidate uninvadable resource allocation schedule

In order to determine how selection shapes the colony growth schedule, we need to determine the uninvadable allocation schedule **v**\* that satisfies the first-order condition (recall eq. 5). We now present this schedule assuming equal mortality in (juvenile) queens and males (i.e. *µ*_q_ = *µ*_m_ = *µ*_r_) and later discuss the relaxation of this assumption.

#### The colony growth schedule

The uninvadable allocation schedule **v**\* consists of two phases: (i) **the ergonomic phase** 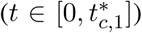 during which workers are produced and (ii) **the reproductive phase** 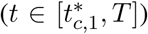 during which sexual offspring are produced (see sections 5 and 6 of SI for derivations). Here, 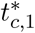 marks the switching time from the ergonomic phase to the reproductive phase and the subscript *c* ∈ {w, q, mx} emphasizes the scenario of genetic control. Resource allocation during the reproductive phase depends on the scenario of dispersal of sexuals: (i) under delayed dispersal, resources should be allocated such that the sex allocation ratio at the end of the season is given by the relatedness asymmetry *R*_*c*_ and (ii) under direct dispersal, males are produced before queens. The switching time 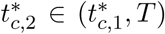 from male production to queen production depends on the scenario of genetic control *c* ∈ {w, q, mx} and the sex allocation ratio is more male-biased than under delayed dispersal

In Figs. 1–2 we have depicted the analytically and numerically determined uninvadable allocation schedules **u**\* in terms of proportional allocation to workers 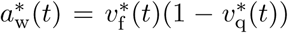, queens 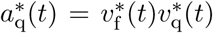, and males 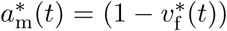 and in Figs. 3–4 we have depicted the respective number of (surviving) individuals (assuming queen monandry (*M* = 1)).

**Figure 1:**
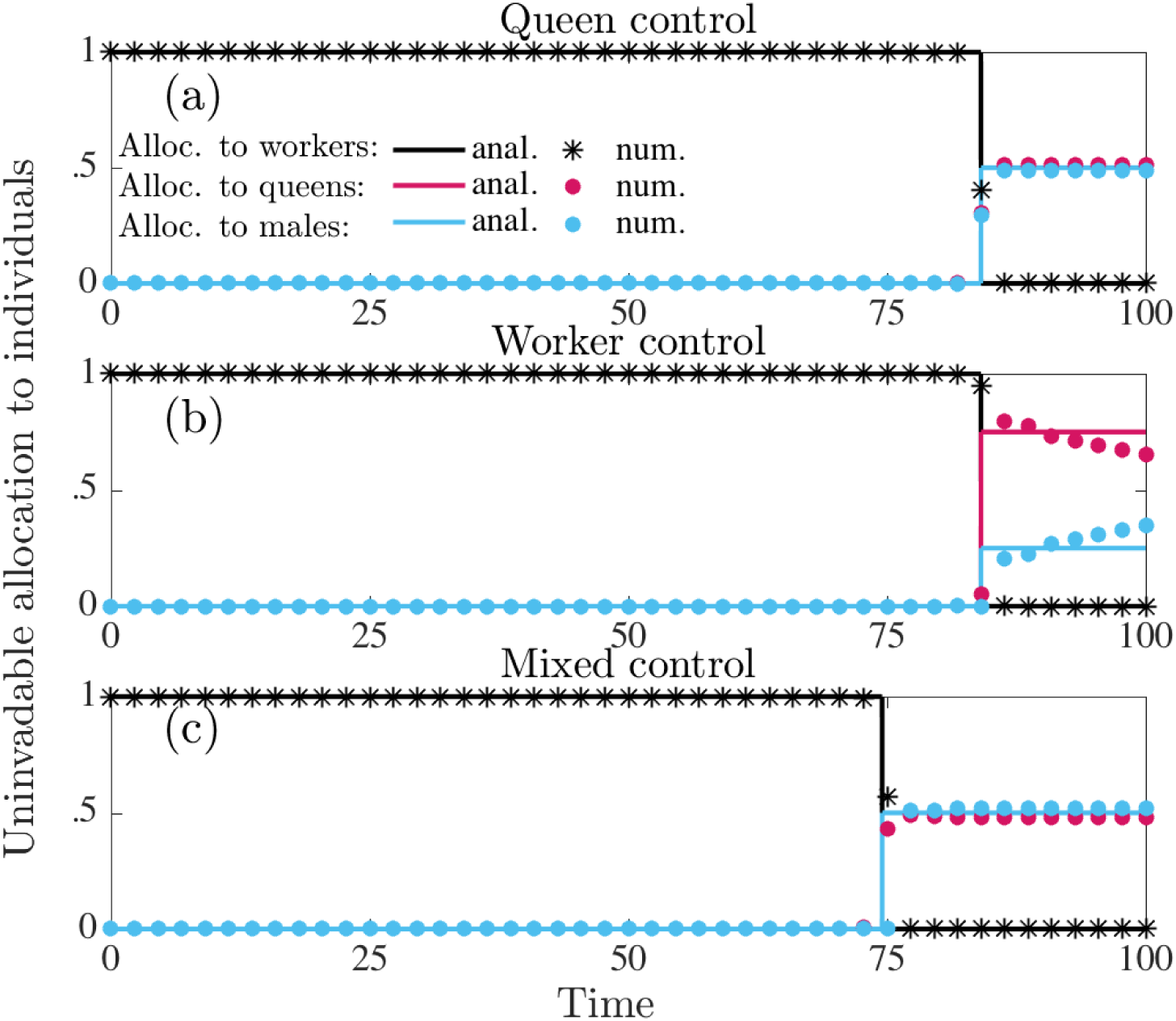
Uninvadable proportional allocation (under delayed dispersal) to workers 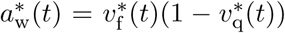 (black), queens 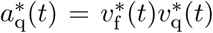 (red), and males 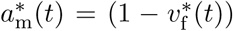 (blue). Solid lines are analytically predicted results and the correspondingly colored symbols represent the numerical results. Panel (a): queen control. Panel (b): worker control. Panel (c): mixed control. Proportional allocation to queens and males exactly match for queen and mixed control, which is why red lines do not appear in the corresponding panels. Notice that the numerical results slightly deviate from the analytical results, since any strategy that gives the sex ratio (queens to males) at the end of the season, equal to relatedness asymmetry *R*_*c*_ of the party in control of *v*_f_ (*t*) has equal invasion fitness (see Fig. 3). Parameter values: *M* = 1, i.e. *C* = 1.5 (queen monandry), *b* = 0.07, *µ*_w_ = 0.015, *µ*_q_ = *µ*_m_ = 0.001, *T* = 100.

**Figure 2:**
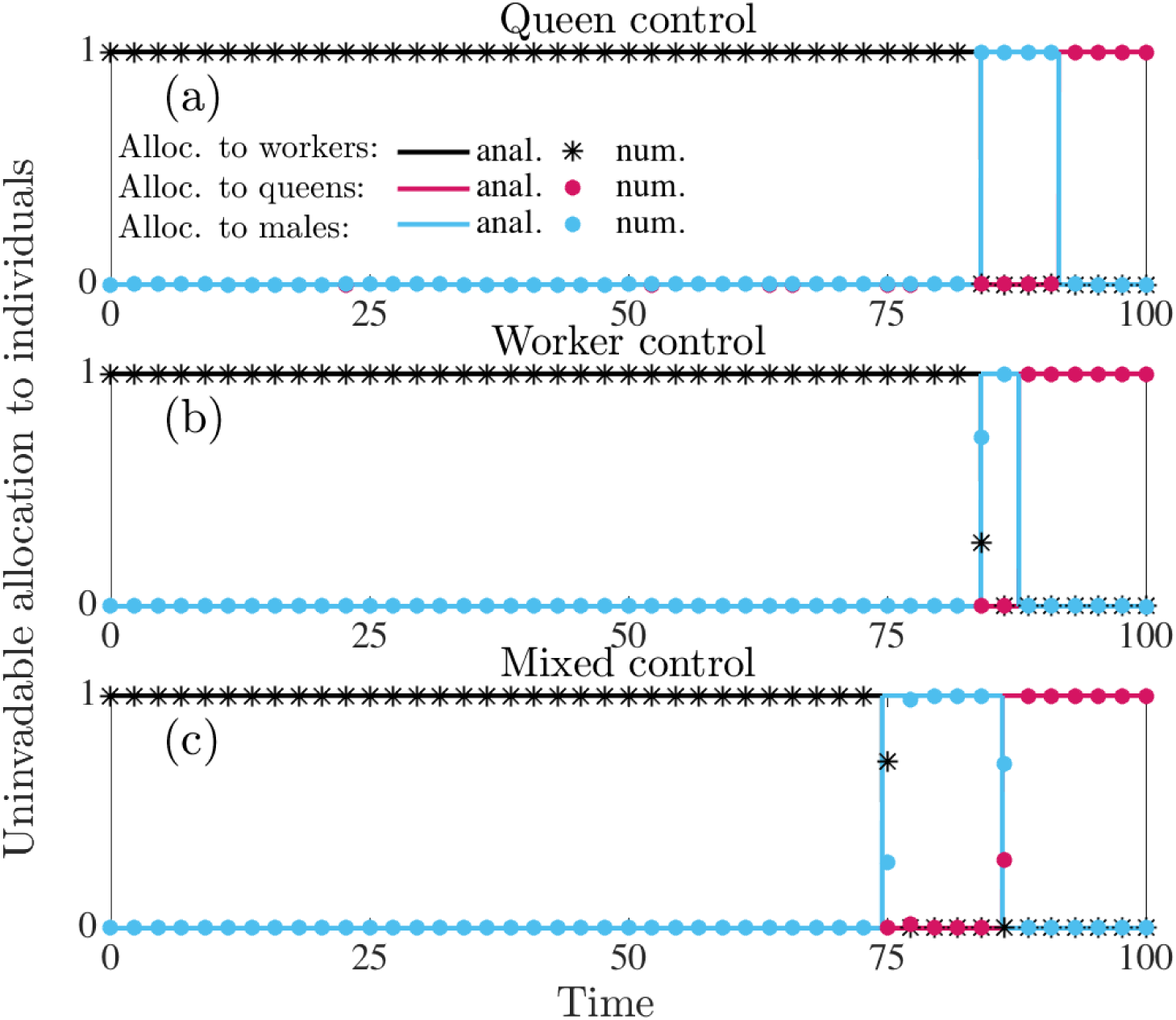
Uninvadable proportional allocation (under direct dispersal) to workers 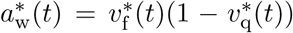 (black), queens 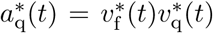 (red), and males 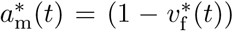 (blue). Solid lines are analytically predicted results and the correspondingly colored symbols represent the numerical results. Panel (a): queen control. Panel (b): worker control. Panel (c): mixed control. Parameter values: *M* = 1, i.e. *C* = 1.5 (queen monandry), *b* = 0.07, *µ*_w_ = 0.015, *µ*_q_ = *µ*_m_ = 0.001, *T* = 100.

**Figure 3:**
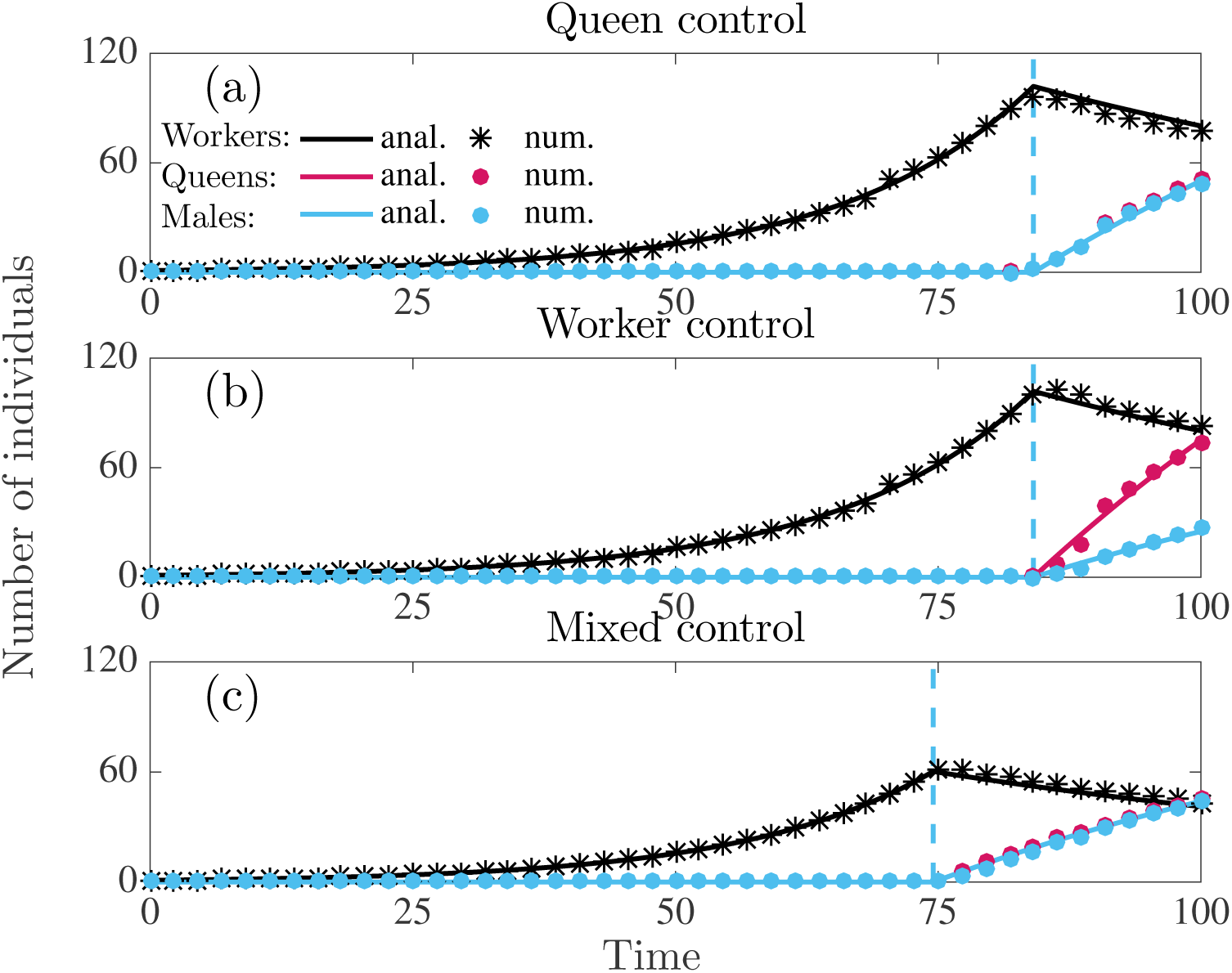
Number of individuals produced in a colony following the uninvadable resource allocation schedule **v**\* under delayed dispersal. Number of workers (black), queens (red), males (blue). Solid lines are analytically predicted results and the correspondingly colored symbols represent the numerical results. Panel (a): queen control. Panel (b): worker control. Panel (c): mixed control. Parameter values: *M* = 1, i.e. *C* = 1.5 (queen monandry), *b* = 0.07, *µ*_w_ = 0.015, *µ*_q_ = *µ*_m_ = 0.001, *T* = 100.

#### Production of workers in the ergonomic phase

The switching time 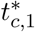 from the ergonomic to the reproductive phase determines the overall amount of resources allocated to workers versus sexuals and it depends on the scenario of genetic control over the resource allocation traits; namely,

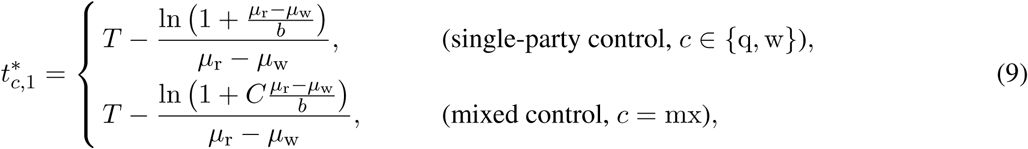

(see sections 5 and 6 of S.I. for derivation, especially see eqs. S109–S117, S119–S122, S133–S150, S140–S147). Under single–party control, this switching time is equal for queen and worker control (i.e. 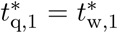). Furthermore, in this case it is identical to eq. (6) of the clonal model of Macevicz and Oster (1976), by setting *b* = *bR, µ*_w_ = *µ*, and *µ*_r_ = ***v*** (see section 13 of S.I. for an overview of how our model relates to previous work).

It follows from eq. (9) that the switch from the ergonomic to the reproductive phase under mixed control 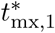 depends on the potential for conflict *C* ≥ 1. Furthermore, this switch happens earlier in the season under mixed control than under single-party control (i.e. 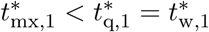, see also Fig. 1 for delayed dispersal and Fig. 2 for direct dispersal, assuming queen monandry, i.e. *C* = 1.5). The switching time 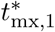 under mixed control happens earlier and, hence, the ergonomic phase is shorter if the potential for conflict *C* is larger. It turns out that the switching time 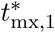 under mixed control is determined by the workers (see section 8 in S.I. for more detailed explanation). Eq. (9) also implies that the onset of early reproduction under mixed control is more pronounced in poor habitats where resource acquisition rate is low and thus reproduction is slow (*b* is small), but colony per-capita productivity still scales linearly as the colony grows (*b* is constant and does not depend on colony size). Increased mortality of workers (*µ*_w_) and decreased mortality of sexuals (*µ*_r_) also cause the time difference between optimal switching time and switching time under mixed control to be larger (see eq. 9).

#### Production of males and queens in the reproductive phase

Under delayed dispersal, selection favors any allocation schedule that produces an allocation ratio of females and males at the end of the season which is equal to the relatedness asymmetry. There are several uninvadable strategies that can satisfy this condition, the most simple one being the constant allocation, i.e. proportional allocation to queens (during the reproductive phase) is given by

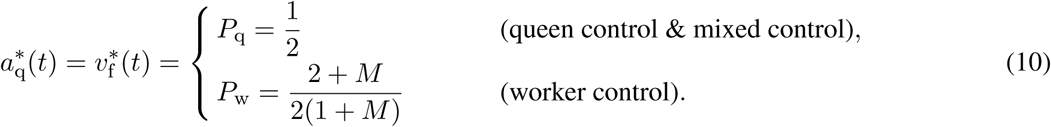

Under direct dispersal, selection favours the production of males before queens (protandry). This is because the reproductive success of males and queens depends asymmetrically on the time they are produced. The switching time 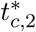 from male production to queen production happens for *M* = 1 when

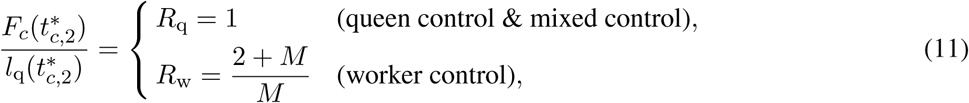

where the left hand side is the ratio of the cost to the benefit to (gene) fitness of producing a queen instead of a male at 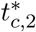 and the right hand side is the exchange rate between inseminated females and queens, which is given by the relatedness asymmetry (see section 9 of S.I. for proof). The cost of producing a queen instead of a male (at 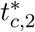) is equal to the potential mating success of a male (born at 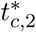), measured in the “currency” of expected number 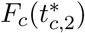 of inseminated queens who survive until the end of the season. The benefit of producing a queen (at 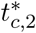) is equal to the probability 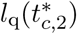 that she survives until the end of the season. Note that in a population where the queens mate *M* times, the expected number 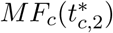 of surviving queens inseminated by males born at time 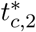, has to be divided by the the queen mating frequency *M* (since the focal male is expected to father only 1/*M* of the diploid offspring). Hence, eq. (11) holds under any queen mating frequency *M*.

The queen is in control of the switch from male production to queen production under mixed control, since under both queen and mixed control the switch happens at the time when producing a male instead of a surviving queen yields one surviving inseminated queen (recall eq. 11). However, this does not imply that the switching time under queen control 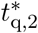 and mixed control 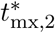 is equal and it follows from eq. (11) that the switching time is

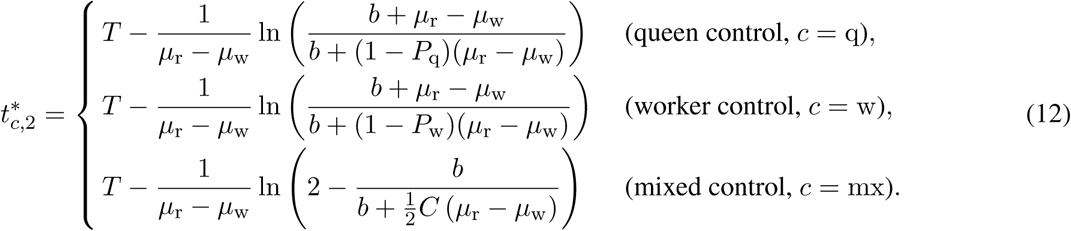

This shows that the switch to production of queens happens later under queen control than under worker control 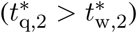, because *P*_q_ *< P*_w_ and it implies that more resources are expected to be allocated to queens under worker control than under queen control (since the length of the reproductive phase is the same under single-party control, i.e. 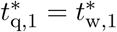). The switch to production of queens happens later under mixed control for higher values of the potential conflict *C*. Furthermore, the switch to queen production happens later when per-worker productivity *b* is small, worker mortality rate *µ*_w_ is large, and the mortality rate *µ*_r_ of sexuals is large.

#### Switching times when the mortality rate of workers and sexuals is equal

In our model (1/*b*) can be loosely interpreted as the time it takes for one worker to help produce one offspring. We show in S.I. (see sections 5.1.3, 5.1.4, and 6.2) that if the mortality rate of sexuals is roughly equal to the mortality rate of workers, then the switching time from the ergonomic to the reproductive phase 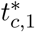 under single-party control (*c* = {q, w}) approaches to the time (1/*b*) it takes for a worker to help produce an offspring before the season end (i.e. 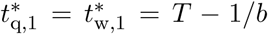); only the individuals produced at the end of the season are reproductive. However, under mixed control the switch happens *C* times earlier (i.e. 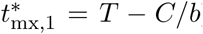). For example, when females mate only once (i.e. *M* = 1 and *C* = 1.5) then the switch to reproductive phase happens at time *T* − 3/(2*b*).

### Colony level traits

#### Colony size at maturity and colony productivity

During the ergonomic phase the number of workers in the colony grows exponentially until it reaches size 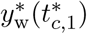 at maturity (i.e. maximum size, see Fig. 3 for delayed dispersal and Fig. 4 for direct dispersal). During the ensuing reproductive phase, sexuals are produced at rate 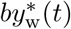. We define as colony productivity the total number 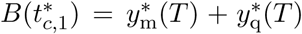 of (surviving) males and queens produced from 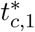 until the end of the season. This can also be interpreted as the total sexual biomass produced in a colony (since we have assumed that males and females are equally costly to produce) and is a quantity often used as a fitness proxy in social insects (Wills et al., 2018). We show that under single-party control the switching time 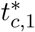 from the ergonomic to the reproductive phase happens exactly at the time that maximizes colony productivity (see section 7.2 in S.I. for proof). Under mixed control the switch from the ergonomic to the reproductive phase happens earlier, especially for higher values of potential conflict *C*. Therefore, we predict that colony size at maturity and colony productivity would decrease with the increase in potential conflict *C* (that can be caused by e.g. low queen mating frequency *M*). See also Fig. 5 for illustration, table 1 for the summary of parameter dependence, and sections 7.1–7.2 of S.I. for more technical details.

**Table 1:**
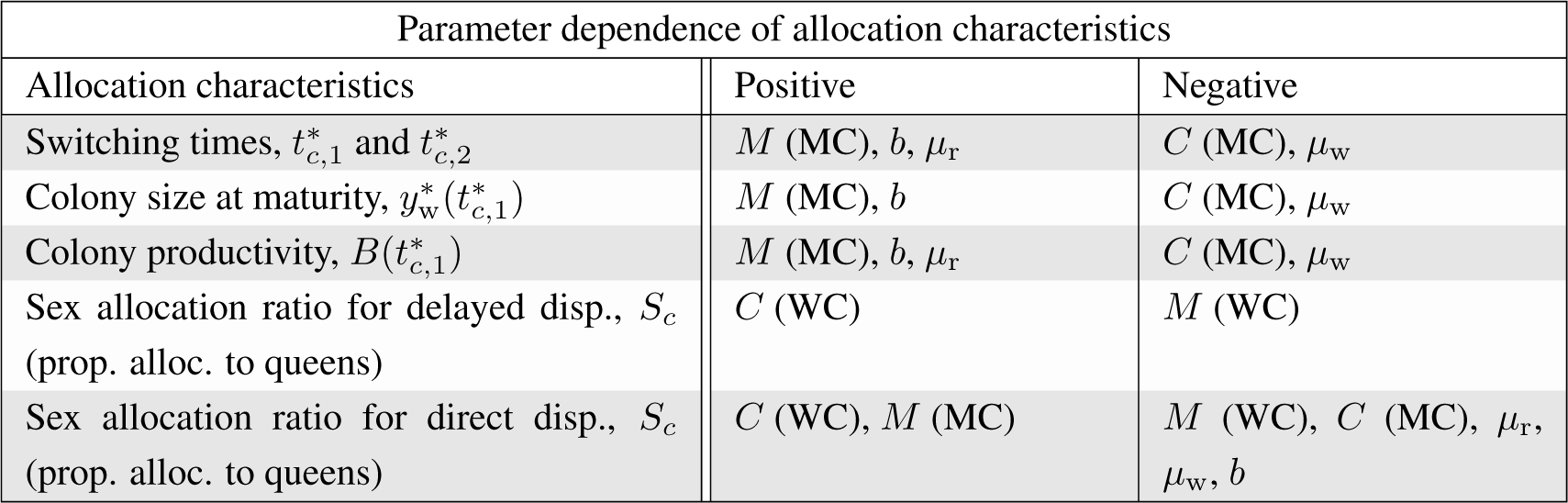
Parameter dependence of colony resource allocation characteristics for biologically meaningful parameter values (*µ*_w_ *>* 0, *µ*_r_ *>* 0, *b > µ*_w_, and *b > µ*_r_). We predict positive relationship between the allocation characteristics and the parameters listed under “Positive” column and negative dependence between the allocation characteristics and the parameters listed under “Negative” column. Here, “(MC)” and “(WC)” that follows after the parameter, emphasizes that this relations only holds for mixed or worker control, respectively.

| Parameter dependence of allocation characteristics |  |  |
| --- | --- | --- |
| Allocation characteristics | Positive | Negative |
| Switching times, $t_{c,1}^*$ and $t_{c,2}^*$ | $M$ (MC), $b$, $\mu_r$ | $C$ (MC), $\mu_w$ |
| Colony size at maturity, $y_w^*(t_{c,1}^*)$ | $M$ (MC), $b$ | $C$ (MC), $\mu_w$ |
| Colony productivity, $B(t_{c,1}^*)$ | $M$ (MC), $b$, $\mu_r$ | $C$ (MC), $\mu_w$ |
| Sex allocation ratio for delayed disp., $S_c$<br>(prop. alloc. to queens) | $C$ (WC) | $M$ (WC) |
| Sex allocation ratio for direct disp., $S_c$<br>(prop. alloc. to queens) | $C$ (WC), $M$ (MC) | $M$ (WC), $C$ (MC), $\mu_r$,<br>$\mu_w$, $b$ |

**Figure 4:**
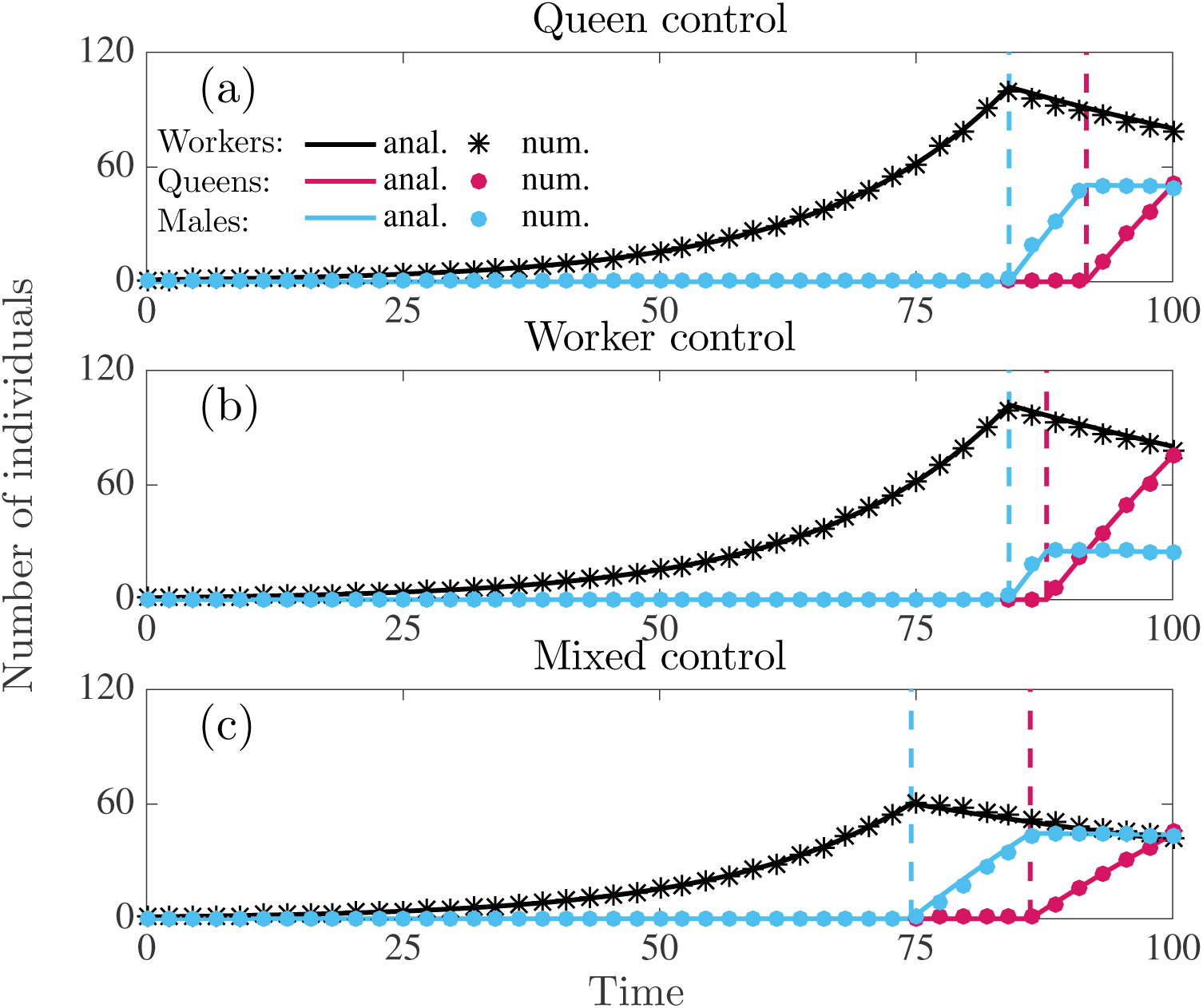
Number of individuals produced in a colony following the uninvadable resource allocation schedule **v**\* under direct dispersal. Number of workers (black), queens (red), males (blue). Solid lines are analytically predicted results and the correspondingly colored symbols represent the numerical results. Panel (a): queen control. Panel (b): worker control. Panel (c): mixed control. Parameter values: *M* = 1, i.e. *C* = 1.5 (queen monandry), *b* = 0.07, *µ*_w_ = 0.015, *µ*_q_ = *µ*_m_ = 0.001, *T* = 100.

**Figure 5:**
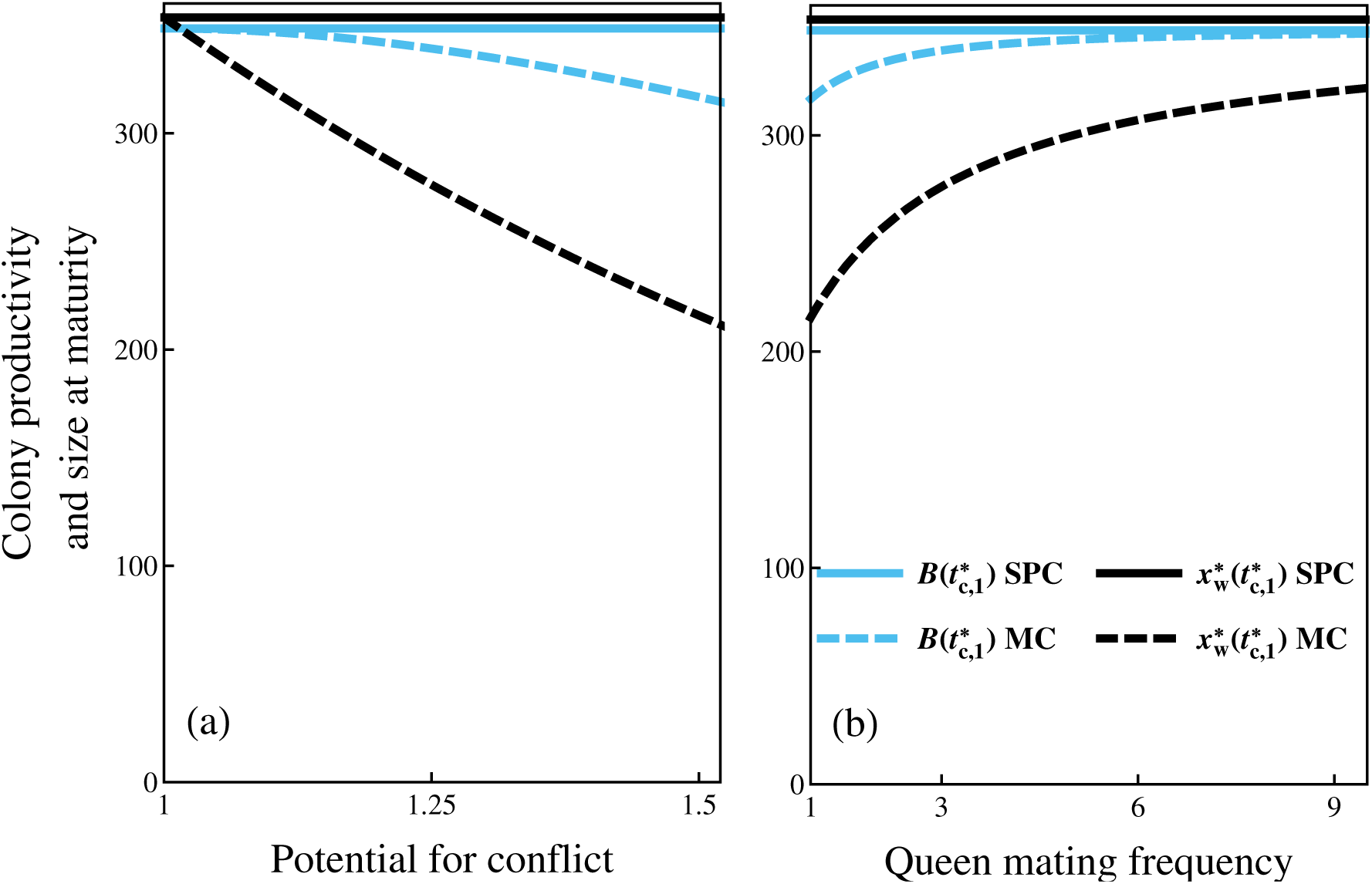
Colony productivity 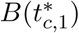 (blue lines) and size at maturity 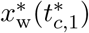 (black lines) under single-party (SPC, solid lines) and mixed control (MC, dashed lines) as a function of the potential for conflict *C* (panel a) and as a function of queen mating frequeny *M* (panel b) for the uninvadable resource allocation schedule **u**\*. Recall that *C* = (2 + *M*)/(1 + *M*). Parameter values: *b* = 0.07, *µ*_w_ = 0.0015, *µ*_q_ = *µ*_m_ = 0.001, *T* = 100.

#### Sex allocation ratio

We define the overall sex allocation ratio *S*_*c*_ at the evolutionary equilibrium as the proportion of the colony resources allocated to queens from the resources allocated to sexuals over the entire season (irrespective of whether they survive to reproduce), where the subscript *c* ∈ {q, w, mx} emphasizes the dependence on the scenario of genetic control (see section 7.3 of S.I. for a formal definition). *S*_*c*_ can be interpreted as the overall proportion of queens among sexuals produced in the colony, since we assume that males and queens are equally costly to produce.

Under delayed dispersal, the overall sex allocation ratio is given by (section 7.3 of S.I., eqs. S172–S175)

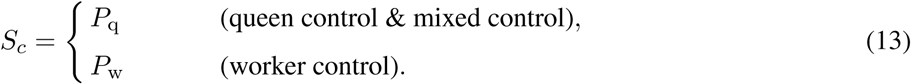

Hence, under delayed dispersal the overall sex allocation ratio is given by the proportional relatedness asymmetry (via eq. 10 and recall eq. 7). It follows from eq. (13) that the prediction for the uninvadable overall sex allocation ratio under single-party control is equal to the corresponding prediction from the standard static models of sex allocation theory (Boomsma and Grafen, 1991; Reuter and Keller, 2001; Trivers and Hare, 1976).

Under direct dispersal, the overall sex allocation ratio is given by (section 7.3 of S.I., eqs. S176–S178)

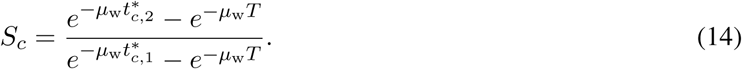

Note that the overall sex allocation ratio under direct dispersal, in contrast to delayed dispersal, depends also on other life-history characteristics of the species and not only on the proportional relatedness asymmetry in the colony (which enters into the equation via 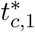 and 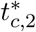).

The overall sex allocation ratio is more male-biased under direct dispersal than under delayed dispersal and compared to results from static models of sex allocation theory (e.g. Boomsma and Grafen, 1991; Trivers and Hare, 1976). Furthermore, the male-bias is more pronounced under mixed control than under single-party control. We illustrate in Figs. 6–7 that this male-bias can be substantial for higher values of mortality rates of sexuals and workers, e.g. *S*_mx_ *≈* 0.35 for mixed control under monandry, compared to *S*_mx_ = 0.5 under delayed dispersal and *S*_mx_ *≈* 0.56 according to the corresponding static allocation model (see Table S3 in section 12 of S.I., see also Table 1 for a summary of how *S*_*c*_ depends qualitatively on the parameters of the model). Mortality of sexuals increases male-biased allocation because it increases the mating success of males produced before the emergence of queens (see Discussion for more elaborate explanation).

**Figure 6:**
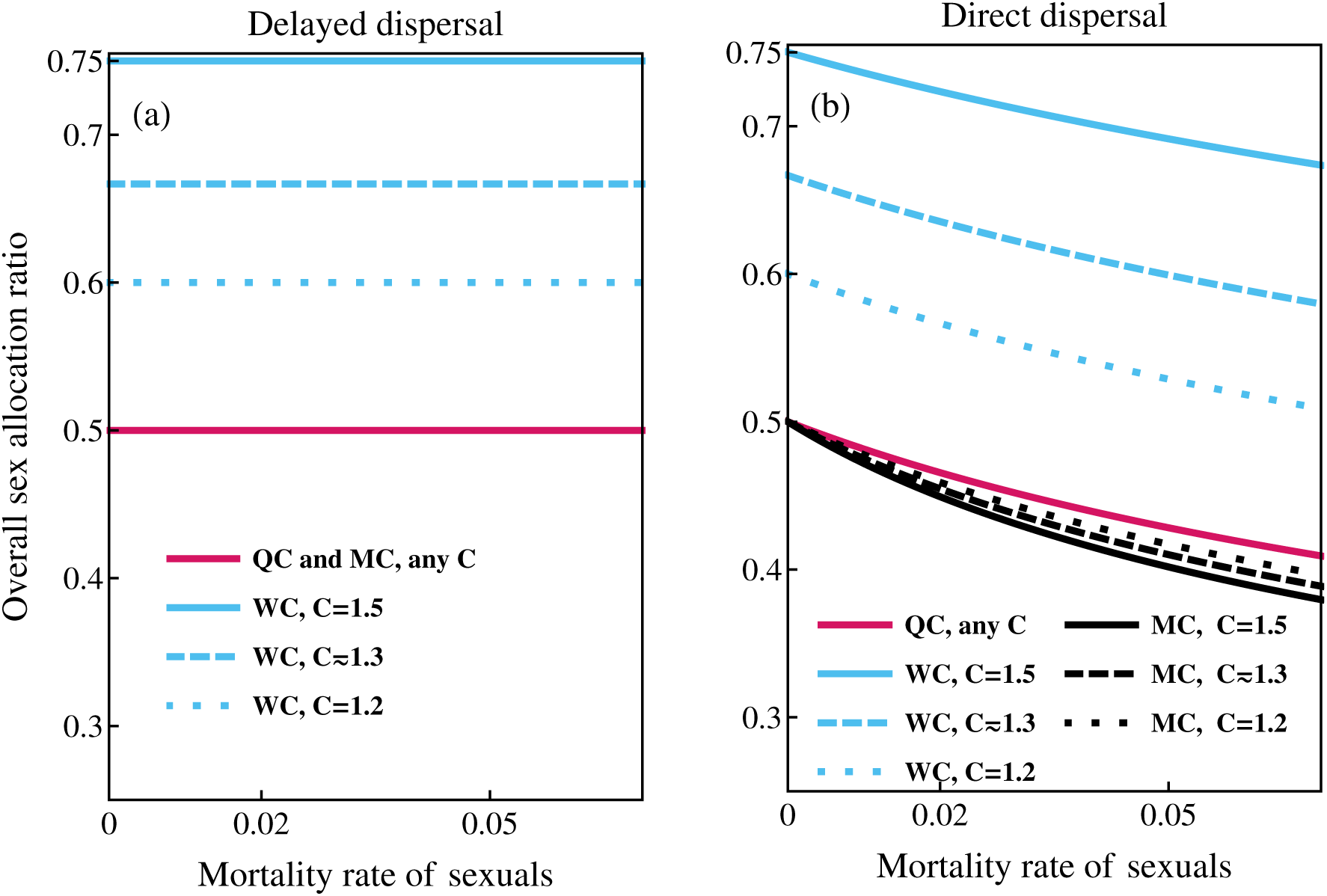
Overall proportional sex allocation ratio *S*_*c*_ (proportional investment into queens) as a function of mortality rate of the sexuals *µ*_r_ for different values of potential for conflict *C*. Panel (a): delayed dispersal; queen and mixed control (QC and MC, red lines), worker control (WC, blue lines). Panel (b): direct dispersal; queen control (QC, red lines), worker control (WC, blue lines), mixed control (MC, black lines). Other parameter values: *b* = 0.07, *µ*_w_ = 0.015, *T* = 100. Note that classical results from static models (e.g Reuter and Keller, 2001) only coincide with these results under delayed dispersal and single-party control.

This effect of mortality in inducing male-biased allocation is stronger under mixed control, especially for higher values of the potential for conflict *C*, since proportionally more sexuals die when the reproductive phase is longer (as it is under mixed control for high values of *C*). Hence, under mixed control and direct dispersal, the overall proportional allocation to queens is lower for higher values for the potential for conflict *C* (i.e. for lower values of queen mating frequency *M*, see Fig. 6–7).

Regardless of the order in which sexuals are produced, the primary sex allocation ratio 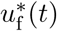 during the reproductive phase determines the overall sex allocation ratio. Hence, the queen is in control of the overall sex allocation ratio under mixed control (see also section 8 in S.I. for more detailed explanation).

#### Unequal mortality rates of sexuals

We now discuss how relaxing the assumption of equal mortality (*µ*_q_ = *µ*_m_ = *µ*_r_) used in the derivation of the above results qualitatively affects these results. From further analysis (section 5.2 of S.I) and our numerical solutions, we find that under delayed dispersal, if the mortality rate of queens and males is not equal, then the sex with the lower mortality rate should be produced earlier, such that by the end of the season the sex ratio of queens to males would be given by *R*_*c*_ under single party control and *R*_q_ under mixed control (assuming that males and queens are equally costly to produce).

We also find that the main conclusions of our results under direct dispersal hold qualitatively if *R*_*c*_*µ*_q_ *≥ µ*_m_under single-party control and *R*_q_*µ*_q_ *≥ µ*_m_ under mixed control. Under direct dispersal, if *R*_*c*_*µ*_q_ *< µ*_m_ then the overall sex allocation under single-party control can be more female-biased than the static models of sex allocation theory predict (e.g. Boomsma and Grafen, 1991; Trivers and Hare, 1976). Similarly, if *R*_q_*µ*_q_ *< µ*_m_ then the overall sex allocation under mixed control and direct dispersal can be female-biased. Furthermore, we find that under mixed control, if the mortality of queens is significantly lower than that of males, then males and queens are produced simultaneously after the switch to the reproductive phase, until there is a switch to producing only females (see section 6.3 of S.I.).

## Discussion

Ontogenetic development of social insect colonies causes behavioural trait expressions of individuals to be necessarily time-dependent (Oster and Wilson, 1979). In this paper, we formulated a mathematical model to analyse how sex allocation conflict affects the dynamic (time-dependent) allocation of resources to workers, queens, and males in annual eusocial monogynous species. We have considered three alternative scenarios of control of colony trait expression (full queen, full worker, and mixed control) and two alternative scenarios of dispersal of sexuals: direct dispersal after eclosion (common among bees and wasps) and delayed dispersal at the end of the season, which resembles the life-history of species, where nuptial flights are synchronized (more commonly found in ants, e.g. see Heinze, 2016 and references therein). Our model extends static allocation models with genetic conflict and dynamic allocation models without conflict and it allows to shed light on a number of questions about colony ontogeny, such as: how does sex allocation conflict affect colony growth? How does sexual selection affect the production of sexuals? Which party wins the sex allocation conflict?

Our results suggest that the marginal benefit of allocating a unit resource to a queen rather than to a male is weighed by the relatedness asymmetry, regardless of any details of colony life-cycle or growth dynamics, thereby generalizing the standard static first-order condition of sex allocation theory (e.g., Boomsma and Grafen, 1991; Taylor and Frank, 1996) to any pattern of colony ontogeny. Solving the first-order condition under our specific life-cycle assumptions using optimal control theory (a non-trivial task, see S.I. sections 5 and 6), we find that selection tends to favor a colony resource allocation schedule that consists of two qualitative phases. First, an ergonomic phase with production of only workers, which determines the colony size at maturity. Second, a reproductive phase with resource allocation to queens and males, which determines the colony productivity and overall sex-allocation ratio. Sexuals can be produced according to various schedules, possibly including switching between producing only males or females (or vice versa), depending on life-cycle assumptions. Colony traits, such as the switching times between different phases of colony growth, maximum colony size, colony productivity, and overall sex-allocation ratio are influenced by the assumptions about the genetic control of resource allocation traits and individual dispersal behaviour.

### How does sex allocation conflict affect colony growth?

Our results confirm earlier predictions derived from dynamic resource allocation models (Macevicz and Oster, 1976; Ohtsuki and Tsuji, 2009) that colony resource allocation should consist of an ergonomic phase and a reproductive phrase. During the ergonomic phase, the marginal return of workers is higher than the return of investment into sexuals. Workers have a higher early marginal return because colony productivity rate (*by*_w_) increases linearly with colony size (hence exponentially during the ergonomic phase), allowing for the production of more sexuals later in the season. Sexuals have a lower early marginal return because they need to survive (queens need to survive until the end of the season and males need to survive until they can reproduce with the surviving queens). The colony switches from the ergonomic to the reproductive phase when producing workers no longer yields the highest marginal return.

We find that under mixed control, colonies switch earlier to the reproductive phase than under single-party control. This early switch evolves because under mixed control the queen controls the sex allocation ratio (for why this is so, see section “Which party wins the sex allocation conflict?” below), meaning that workers can not increase allocation to queens during the reproductive phase, even though producing more queens would increase the fitness of genes residing in workers. Hence, workers start rearing female eggs (destined to become workers under single-party control) into queens earlier, in order to increase the allocation to queens. Hence, asymmetric control over the sex allocation ratio causes the switching time to the reproductive phase to be controlled by the workers (see also section 8 of S.I. for more technical explanation).

Colony size at maturity and colony productivity are expected to be smaller under mixed control than under single party control. Under single-party control the colony productivity is maximized, but not under mixed control (see section 7.2 in S.I. for proof and Fig. 5). This is so because in the latter case the switch to the reproductive phase occurs earlier, causing colony size at maturity to be smaller (there is less time for worker numbers to increase exponentially during the ergonomic phase). Therefore, there are fewer workers to produce sexuals in the reproductive phase, which results with a decline in colony productivity (colony-level cost of sex allocation conflict).

A loss in colony productivity due to sex allocation conflict was already predicted using a static (Reuter and Keller, 2001) and a dynamic allocation model assuming delayed dispersal (Bulmer, 1981). But for the latter model, the outcome of the resource allocation conflict is different from ours. Indeed, Bulmer (1981) concluded that colonies die one generation before the end of the season if the sex allocation at the population level is biased towards queens, since the queens are producing only males in the penultimate generation. His conclusion relied on the assumption that colony growth is divided into discrete generations, such that worker generations within a season do not overlap and in his model he only considered two generations before the end of the season. Our analysis not only extends the results of Bulmer (1981) to less restrictive life-cycle assumptions and to direct dispersal of sexuals, but it also provides quantitative predictions for the the switching time from the ergonomic to the reproductive phase. Indeed, we predict that the premature switch from the ergonomic to the reproductive phase is earlier in species where the resource acquisition rate is low, the mortality rate of workers is high and that of sexuals low. We also show that the switching time from the ergonomic to the reproductive phase under mixed control are equal for both delayed dispersal and direct dispersal. This implies that sexual selection and the evolution of protandry do not have an effect on the cost of sex allocation conflict that manifests itself through loss of colony productivity.

The switching time to the reproductive phase under mixed control depends on the potential for conflict *C*, which is the ratio of party-specific proportional contribution of genes through queens to the future of the gene pool (eq. 8), and a decreasing function of the mating number *M* of a queen. Our results imply that colonies with lower potential for conflict *C* are expected to grow larger and have higher colony productivity. Similar effects can be expected to hold for other factors that reduce the queen-worker conflict over sex allocation, for example polygyny of related queens or worker production of male eggs (Ratnieks et al., 2006; Reuter and Keller, 2001). We have assumed monogyny, but allowing for multiple queens per colony should be a relatively straightforward extension to our model. Our analysis implies that polyandry is expected to evolve under mixed control, given that the workers are able to assess the mating frequency of the queen (Pamilo, 1991b). However, empirical evidence suggests that polyandry is generally less common in annual eusocial insects but has been found, for example, in *Polistes* (Seppä et al., 2011) and *Vespula* (Johnson et al., 2009).

The so-called “bang-bang” schedule of colony growth, such that allocation to workers and sexuals never occurs simultaneously, represents a general life-history principle of growth and reproduction in annual organisms for which productivity rate scales linearly with size and environmental fluctuations that can cause variations in the length of the season or food availability are small (Cohen, 1971; King and Roughgarden, 1982). A key result of our analysis is that the sex allocation conflict does not affect the overall shape of the colony growth curve, but only the time of the switch between growth and reproduction. This is not an obvious result, since trade-offs between producing different types of individuals are not linear. It has been shown before (assuming clonal reproduction) that selection favours a singular control (sometimes called a graded control; i.e. workers and sexuals are produced simultaneously) if the productivity rate (i.e. *by*_w_) scales non-linearly with colony size, such that *b ≡ b*(*y*_w_) (Beekman et al., 1998; Poitrineau et al., 2009), but not for environmental fluctuations acting alone (Mitesser et al., 2007b). The properties of the relationship between productivity rate and colony size affects the way the marginal value of producing a worker changes over time, but not the marginal value of producing queens and males. In principle, this could affect the outcome of the sex-allocation conflict and it would be interesting to see if the results of our model change when the productivity rate would scale non-linearly with colony size.

Inherently, our model assumes that individuals in the colony possess some physiological mechanism that enables them to estimate the timing of the switch from the ergonomic phase to the reproductive phase. Currently, the underlying mechanism behind the timing of the switch from the ergonomic to the reproductive phase is not known (but it has been shown that *Bombus terrestris* queens are able to control the switching time endogenously, Holland et al., 2013). Nevertheless, the framework of our model can be used to also study the evolution of eusociality, when we allow for the brood to have control over their own developmental fate. Current models that study the emergence of eusociality that explicitly track colony growth usually fix the switch from ergonomic to reproductive phase to happen at arbitrary size of the colony (e.g. Avila and Fromhage, 2015). Hence, extending our model to study evolution of eusociality could explain how life-history interacts with other mechanisms that are known to drive the evolution of eusociality.

### How does sexual selection affect the production of sexuals?

Our model predicts simultaneous production of queens and males under delayed dispersal and protandry (males produced before females) under direct dispersal. Under delayed dispersal, both males and queens have to survive until the end of the season to mate and their reproductive success depends symmetrically on the time that they are produced. Under direct dispersal, males have to survive until there are females available to mate, while queens have to survive until the end of the season. This asymmetry leads to protandry.

Our prediction about the evolution of protandry relies on the assumption that the females mate immediately and indiscriminately after dispersal with the males currently in the mating pool. However, there is some evidence of female choice in some social insects (Baer, 2003 and references therein). Nevertheless, there is also evidence that earlier emergence of males can give them an advantage in mating success through precopulatory sexual behaviours or through the use of mating plugs (Baer, 2003, 2014; Baer et al., 2000; Foster, 1992).

### Which party wins the sex allocation conflict?

We show that the queen wins (more accurately, the genes in queens win) the sex allocation conflict, because the evolution of distinct phases of colony growth constrains the ability of workers to manipulate the overall sex allocation ratio. Indeed, during the reproductive phase, the ratio at which the queen lays the female versus male eggs determines the overall sex allocation ratio, since workers can only influence the developmental fate of the female eggs. Therefore, the only option for workers to increase the allocation to queens is to switch to the reproductive phase earlier at the expense of reduced colony productivity, while queens, regardless of the early switch, can always further affect the sex-ratio without disturbing colony productivity.

The evolution of different phases of colony growth is thus crucial as it decouples the trade-offs experienced by the queens. During the ergonomic phase, there is a latent trade-off between producing males versus workers (since workers rear all the female-eggs into workers), while during the reproductive phase there is a trade-off between producing queens versus males (since workers rear all the female-eggs into queens). The distinct phases of colony growth also decouples how queens and workers can affect the allocation decisions in the colony, impeding the ability of workers to influence the overall sex allocation during the reproductive phase and the ability of queens to influence the proportional allocation to workers versus sexuals (see also section 8 of S.I. for more detailed explanation). Our results thus suggest that the queen is always expected to win the sex allocation conflict, as long as workers and sexuals are produced during separate phases of colony growth and workers can only influence the developmental fate of the female eggs.

### The overall sex allocation ratio

In our model, the overall sex allocation ratio depends on the scenario of dispersal of sexuals. Under mixed control, the overall sex allocation ratio is expected to be even under delayed dispersal and male-biased under direct dispersal (given that the mortality rate of males and queens is equal). Under single-party control and delayed dispersal, the overall sex allocation ratios predicted by our model are in accordance with the classical static models (e.g. Boomsma and Grafen, 1991; Trivers and Hare, 1976) and do not depend on the life-history characteristics of the species (e.g. mortality rate of sexuals or workers). However, under direct dispersal, we observe more male-biased overall sex allocation ratios than occur in the static models of sex allocation theory (e.g. Boomsma and Grafen, 1991; Trivers and Hare, 1976), especially for higher mortality rates of sexuals (see Fig. 6) and lower mortality rates of workers (see Fig. 7).

**Figure 7:**
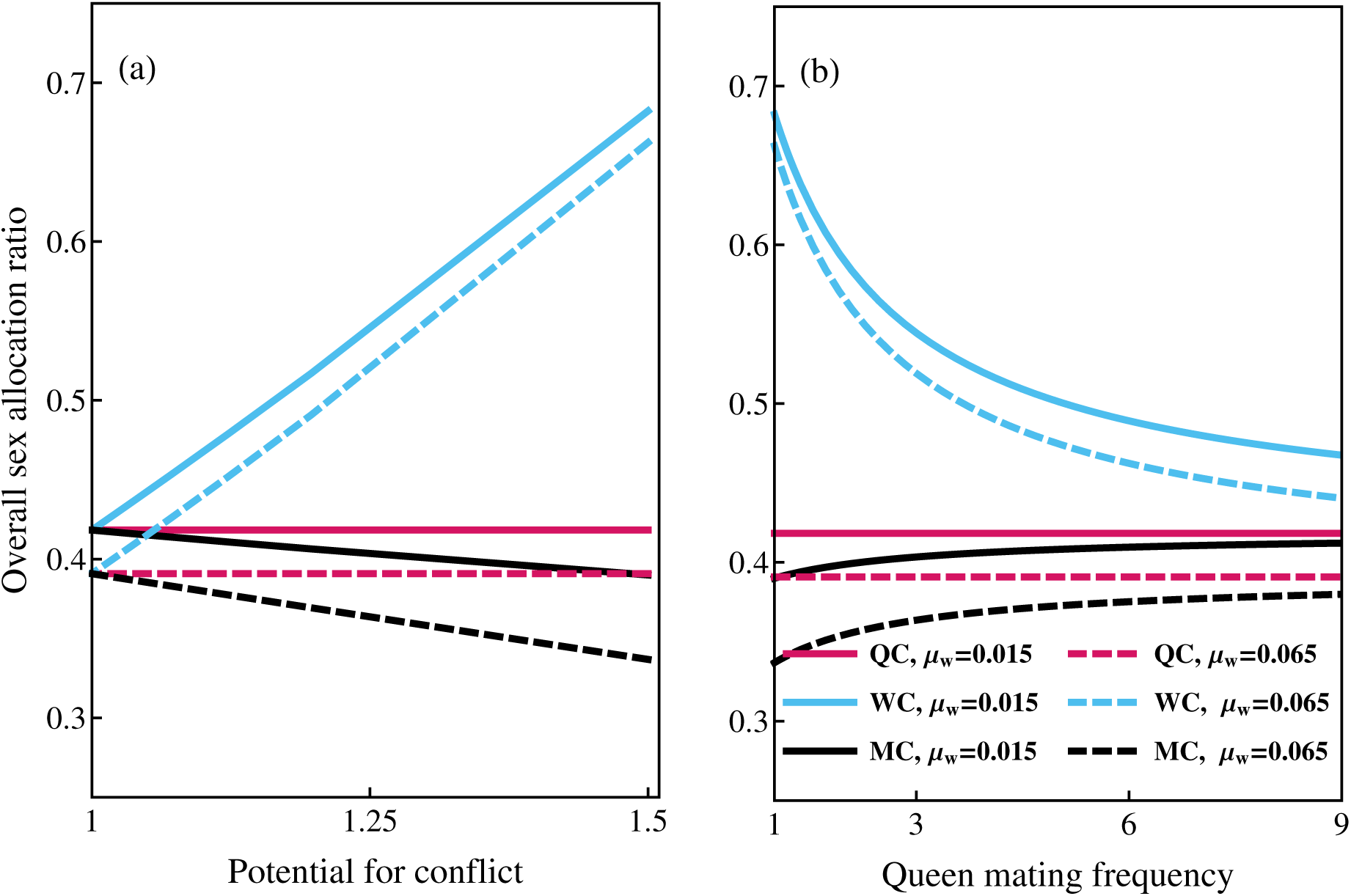
Overall proportional sex allocation ratio *S*_*c*_ (proportional investment into queens) under direct dispersal as a function of the potential for conflict *C* (panel a) and queen mating frequency *M* (panel b) for different values of mortality of workers *µ*_w_. Queen control (QC, red lines); worker control (WC, blue lines); mixed control (MC, black lines). Parameter values: *b* = 0.07, *µ*_r_ = 0.06, *T* = 100.

More male-biased sex allocation ratios evolve under direct dispersal because mortality affects the coevolution of protandry (that evolves due to sexual selection on males) and sex allocation ratio. The sex allocation ratio is determined by the switching time from male production to queen production. This happens when pro-ducing a male yields *R*_*c*_ (surviving) inseminated queens, instead of producing a (surviving) queen. Hence, the relative mating success of males compared to the survival probability of queens determines the switching time from male production to queen production. When mortality of sexuals is high, males produced later in the season (just before the emergence of queens) have a higher mating success for higher values of mortality of sexuals, since there are fewer surviving males to compete with. Hence, higher mortality of sexuals delays the switch to queen production because it increases the mating success of males (see section 9 of the S.I. for a more detailed analysis and explanation). Our result that mortality affects the sex allocation ratio appears to be at variance with Fisher’s (1930) result that mortality after parental investment (either differential between the sexes or not) should not affect the uninvadable sex allocation ratio (see e.g. West, 2009, p. 19–20). The reason for this apparent discrepancy is that, in our model, mortality causes resources that are invested into sexuals earlier to yield lower fitness returns (since early-produced sexuals have a lower chance to contribute to the next generation). So, mortality causes males to be produced more cheaply (at a time when allocating resources yield smaller returns). Hence over-production of males under higher mortality is in fact consistent with Fisher’s prediction that more offspring should be produced of the cheaper sex.

Under direct dispersal, the overall sex allocation ratio is more male-biased for mixed control than for queen control, even though for both queen and worker control, the switch from male production to queen production happens when producing a male instead of a surviving queen yields one surviving inseminated queen. This is because, for mixed control, the reproductive phase is longer during which proportionally more males die before they can mate, which increases the mating success of males produced later. This is why the overall allocation is more male-biased under mixed control for higher values of mortality of sexuals (see Fig. 6) and for other life history characteristics that cause the reproductive phase to be longer, such as higher values of the mortality rate of workers *µ*_w_ (see Fig. 7). Hence, we find that in protandrous species, proportionally more resources are expected to be allocated into producing males.

Surprisingly, under direct dispersal and mixed control the overall sex allocation ratio *S*_mx_ becomes more male-biased as the workers become more related to the female brood (their sisters) (i.e. if the potential for conflict *C* increases or the queen mating frequency *M* decreases, see Fig. 7). This prediction follows from the combined effect of protandry under direct dispersal and a longer duration of the reproductive phase for higher values of the potential for conflict under mixed control. If workers are more related to the female brood (e.g. for higher values of the potential conflict *C*), then the mating success of males produced later is higher, because proportionally more males have died due to early switch to the reproductive phase. For these reasons, worker relatedness to female brood is expected to correlate negatively with the proportional investment into queens when resource allocation is under mixed control. This prediction contradicts standard results from the static models of sex allocation theory (Boomsma and Grafen, 1991; Trivers and Hare, 1976) that predict the opposite correlation. We expect that other factors that reduce the queen-worker conflict over sex allocation have qualitatively similar effects on overall proportional allocation to queens.

Most comparative studies about population-wide sex allocation of eusocial Hymenoptera come from ants, where sex-allocation is mostly female-biased (Bourke and Franks, 1995; Ratnieks et al., 2006; Sundström et al., 1996), although it is not universal (Fjerdingstad et al., 2002; Helms, 1999; Helms et al., 2000; Passera et al., 2001). However, most ant species are perennial and their life-cycles diverge in many respects from the assumptions of our model. In bumble bees, who are annual and mostly monogynous species, the population-wide sex allocation tends to be overwhelmingly male-biased (Bourke, 1997). Indeed, Bourke (1997) found that the median proportional allocation to queens is only 0.32 (range 0.07–0.64) among 11 populations of seven bumble bee species. Interestingly, Johnson et al. (2009) found that in a social wasp (*V. maculifrons*) nestmate relatedness is negatively associated with overall investment into queens which would be in accordance with our model for mixed control under direct dispersal with male protandry (see Fig. 6). However, these results arise from a dataset where the queens have a relatively high mating frequency and the variation between mating frequencies is not very large (hence, the effect size is not very large) and male protandry in that species is not entirely clear (Johnson et al., 2009).

### Static and dynamic approaches to resource allocation conflicts

Corresponding static and dynamic models can make different predictions for the outcome of the conflict. This can be seen when comparing the predictions of our model under delayed dispersal with the predictions of a corresponding static model by Reuter and Keller (2001). See section 12 of S.I., for a proof that our model is indeed comparable to that of Reuter and Keller (2001), even though there is a slight deviation in the assumption about how productivity scales with colony size (since this assumption does not affect qualitatively their results). We followed their approach on modeling conflict by way of using mixed control of colony allocation traits, but our result that queen wins the sex allocation conflict contradicts with theirs. Indeed, they predicted that the sex allocation ratio under mixed control is intermediate between sex allocation ratios predicted for queen and worker control (the exact values depending on the assumption about how productivity scales with colony size). This contradiction arises, because in our dynamic model the sex allocation ratio is determined during the reproductive phase by the queen, while in the model of Reuter and Keller (2001) behavioural decisions can not vary over time, meaning that the two parties make their decisions simultaneously for the whole season *T*. Hence, this way of modelling links all the allocation decisions together to happen simultaneously, which leads to the result that workers can influence the sex allocation ratio by rearing some worker–destined female brood into queens.

It has been shown by Pen and Taylor (2005) that if the two parties make their allocation decisions sequentially (the so-called Stackelberg equilibrium, such that the queen acts first and workers respond), then the queen is expected to win the sex allocation conflict even assuming static resource allocation decisions. Pen and Taylor (2005) studied a static resource allocation model similar to the model of Reuter and Keller (2001)), but they also looked at the effect of information exchange between the two parties. While they arrived at a conclusion similar to ours about the overall sex allocation ratio, our result implies that the workers do not have to have the information about the ratio at which the queen lays the male to female eggs.

Reuter and Keller (2001) also generally argue that complete control by a single party is not evolutionarily stable, since the conflict over sex allocation strongly selects for the other party to manipulate the sex allocation leading to a stable evolutionary equilibrium where the sex allocation is intermediate between the predicted evolutionary outcomes for full control of the two parties. However, under the dynamic model, we show that under the assumptions of mixed control, an intermediate sex allocation will not evolve.

## Conclusion

We showed that when dynamic properties of resource allocation are considered, sex allocation conflict can substantially affect colony ontogeny, and thus overall patterns of growth and productivity. Helanterä (2016) has argued that life-history trade-offs may be easier traits to conceptualize as organismal traits (i.e. traits evolving like group-selected adaptations), as opposed to traits more heavily contingent on conflicts among genes in different individuals, such as traits involving sex allocation and dispersal behaviour. In contrast, our model suggests that colony life-history traits can generally not be viewed in isolation from traits that are influenced by genetic conflicts, and hence both, the “morphology” and “physiology” of a colony are likely to be affected by them, leading to a general breakdown of the “organismic” perspective of eusocial insect colonies.

## Supporting information for

## 1 Evolutionary analysis

### 1.1 Mutant-resident system

We analyze the evolution of allocation schedules by performing an evolutionary invasion analysis (e.g., Caswell, 2001; Charlesworth, 1994; Eshel and Feldman, 1984; Ferrière and Gatto, 1995; Fisher, 1930; Lehmann et al., 2016; McNamara et al., 2001; Metz, 2011; Otto and Day, 2007) whereby we consider the fate (invasion or extinction) of a single mutant allele introduced into a population of resident individuals, where the mutant allele determines an allocation schedule that is different from the resident allocation schedule throughout the entire season. We assume that the traits are determined at separate single loci with two segregating alleles (resident and mutant).

Let *v*_*τ*_ (*t*) and *u*_*τ*_ (*t*) denote the resident and mutant resource allocation phenotypes for a trait of type *τ* ∈ {f, q}, respectively. It will turn out to be useful to define the mutant phenotype *u*_*τ*_ (*t*) as a trait expressed by a (hypothetical) colony where all the genes in control of the trait are mutant alleles (i.e. individuals whose genes are in control of the trait are homozygous for the mutant allele).

Since the fate (invasion or extinction) of a mutant allele is determined when it is rare in the population, then only one of the colony “founding” individuals is carrying a single copy of the mutant allele (heterozygous diploid female or a hemizygous haploid male). By writing that a mutant male “founds” a colony, we mean that a mutant male has mated with a resident female that gives rise to a focal colony, where the mutant allele is present in the genes of the workers. Furthermore, in haplodiploid systems, where females are diploid and males are haploid, the phenotypes of colonies founded by mutant individuals of opposite sexes will be different. The distinction between the phenotypes expressed in these two types of colonies will also turn out to be useful when describing these phenotypes under the various assumptions of genetic control that different parties have over the traits.

Thus, we will denote by 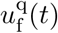 (and respectively, by 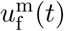 the proportion of resources allocated at time *t* to producing females in a colony founded by a focal mutant heterozygous female (hemizygous male) and by 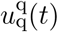 (and respectively, by 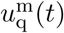) the proportion of resources allocated to producing queens from resources allocated to females at time *t* in a colony founded by a focal mutant heterozygous female (hemizygous male). Let 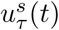 denote the resource allocation phenotype for a trait of type *τ* ∈ {f, q} of a colony founded by a heterozygous (hemizygous) individual of type *s* ∈ {q, m} and it can be expressed as (assuming additive genetic effects)

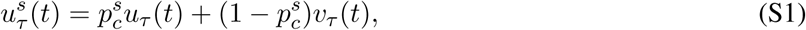

where 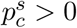 is the expected frequency of the mutant allele in party *c* ∈ {q, w} in full control of the trait of type *τ* in a colony founded by a mutant individual of type *s* (*s* ∈ {q, m}). Hereinafter, the subscript *c* = q denotes a scenario of full queen control and *c* = w denotes a scenario of full worker control, and *c* = mx denotes a scenario of mixed control.

Under queen control of the trait, the expected colony phenotype 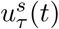 is determined from the frequency of the mutant allele in the colony-founding queen. If the colony is founded by a heterozygous mutant female then the frequency of the mutant allele in the colony-founding queen is 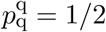. Under queen control, mutant males who have mated with a colony-founding queen have no genetic influence on the resource allocation traits and thus 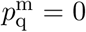. Hence, female mating frequency will also have no affect on the trait under queen control. Under worker control of the trait, the expected colony phenotype 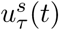 is determined from the expected frequency of the mutant allele in workers. The expected frequency of the mutant allele in workers in a colony founded by a heterozygous mutant female is 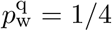 and it is not affected by the mating frequency of the queen because a mutant female will only encounter resident males since the mutant allele is considered to be rare. The expected frequency of the mutant allele in workers in a colony founded by a mutant male is 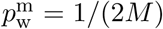, where *M* is the number of times the female has mated (when the mutant allele is rare, only one of the males is carrying the mutant allele).

Let **v** = {*v*_f_ (*t*), *v*_q_(*t*)}_*t*∈[0,*T*]_ denote the full allocation schedule of a colony founded by resident individuals, i.e. it describes how colony resources are allocated throughout the entire season *t* ∈ [0, *T*]. Similarly, let **u** = {*u*_f_ (*t*), *u*_q_(*t*)}_*t*∈[0,*T*]_ denote the full allocation schedule of a colony founded by individuals who carry only mutant alleles for both of the evolving traits. Similarly, let 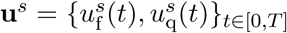 denote the full allocation schedule of a colony founded by a heterozygous (hemizygous) individual of type *s* for each of the evolving traits, hence **u**^*s*^ depends on **u**. This notation turns out to be useful for performing the invasion fitness analysis, but it does not imply that we are assuming pleiotropic effects.

Let 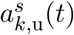 be the proportion of resources allocated to producing type *k* ∈ {w, q, m} individuals in a colony founded by a heterozygous individual of type *s* ∈ {q, m}, where the subscript “u” in 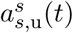 emphasizes that it is the mutant allocation schedule, which, according to eq. (3) is

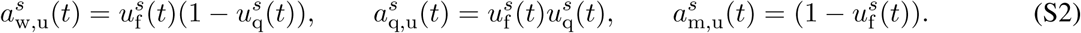

The rate of change in the number of type *k* ∈ {w, q, m} individuals alive at time *t*, that have been produced in a colony founded by a mutant individual of type *s*, is given by the equation

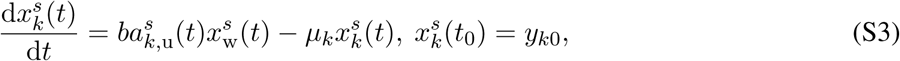

where 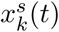 denotes the number of individuals of type *k* alive at time *t* that have been produced in a colony founded by a mutant individual of type *s*. The rate of change of females 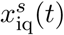 alive at time *t*, who have been inseminated by the males produced in the focal colony (under a monandrous mating system) founded by a mutant individual of type *s*, is given by the equation

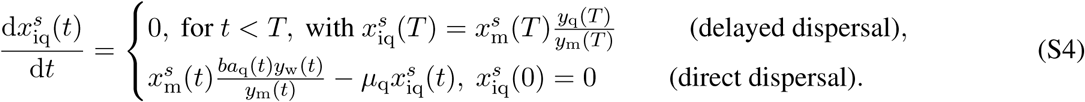

Note that the number of females alive at time *t*, who have been inseminated by the males produced in the focal colony founded by a mutant individual of type *s* in a mating system where females mate *M* times is *M* 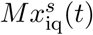.

The rate of change of individuals of type *k* ∈ {w, q, m} produced in a resident colony and females insemi-nated by males from the resident colony are given by eq. (1) and eq. (2), respectively.

### 1.2 Fitness functions of mutant individuals

We will express the invasion fitness of the mutant allele in terms of gene transmission frequencies and fitness functions of mutant individual of type *s* (*s* ∈ {q, m}), who have founded colonies in the current season. The fitness is measured as the expected number of daughters and sons who become colony founders in the next generation. Note that, since females mate with *M* males, a son can become a colony founder for multiple colonies. Hence, the number of sons who become colony founders in the next generation can be measured in the number of females inseminated by sons, who will become colony-founding queens in the next generation (Bulmer, 1994, p. 213). Hence, one fitness measurement cycle lasts from the beginning of the current season to the beginning of the following season and we keep track of the genes in mutant colony-founding individuals of type *s*.

Let *w*_*s′ s*_(**u**^*s*^, **v**) denote the expected number of mutant colony-founding individuals of type *s′* ∈ {q, m} in the following season that descend from a current colony-founding mutant individual of type *s* ∈ {q, m} in a resident population. The fitness function *w*_*s′ s*_(**u**^*s*^, **v**) is a function of the allocation schedule **u**^*s*^ of a colony founded by an individual of type *s* (by way of eqs. S2–S4). Note that the fitness function *w*_*s′ s*_(**u**^*s*^, **v**) is ultimately a function of the mutant schedule **u**, the frequency 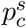 of the mutant allele in the average individual in control of a resource allocation trait, and of the resident allocation schedule **v** (by way of eq. S1). However, since the mutant allele is considered to be rare for the invasion analysis and the population size is large, then the fitness function *w*_*s′ s*_(**u**^*s*^, **v**) is independent of the number (or frequency) of mutants in the population.

For calculating the fitness functions, we only need to specify the number of individuals alive at the end of the season *t* = *T*. To that end it is useful to set

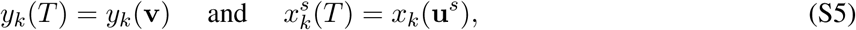

which gives the number of individuals of type *k* ∈ {w, q, m, iq} at the end of the season *t* = *T* associated with a resident colony and a colony founded by a mutant individual of type *s*, respectively (by way of eqs. 1–2 and S3–S4). Note that type *k* ∈ {w, q, m} individuals are individuals produced in a focal colony and type *k* = iq individuals are females inseminated by sons produced in the focal colony. In eq. (S5) we have emphasized the functional dependence of the number of individuals at the end of the season on the allocation schedules, **u**^*s*^ and **v** (recall eqs. S2 and 3).

Next, we derive the fitness functions *w*_*s′ s*_(**u**^*s*^, **v**). A colony-founding female is expected to have *x*_q_(**u**^q^) surviving daughters (juvenile queens) at the end of the breeding season and her sons are expected to have inseminated *Mx*_q_(**u**^q^) surviving females at the end of the breeding season. The probability that a daughter or a female inseminated by a son will gain any one of the *n* breeding spots is *n/ny*_q_(**v**), since there are total number of *ny*_q_(**v**) juvenile queens competing for these spots. Hence, the number of mutant colony-founding individuals of type *s′* in the next generation that descend from a mutant colony-founding female can be written as

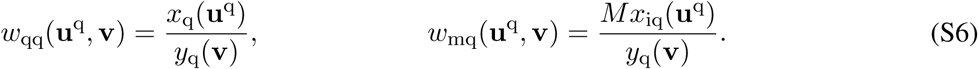

Since females mate with *M* males, each colony-founding male is expected to only father 1/*M* of the offspring (here we have not yet taken into account the transmission frequencies specific to different genetic systems). Hence, a colony-founding male is expected to father *x*_q_(**u**^m^)/*M* surviving daughters at the end of the breeding season and his sons are expected to have inseminated *Mx*_iq_(**u**^m^) surviving females at the end of the breeding season. Here we formulate the fitness functions for any genetic system and by “sons” we mean males produced by a queen that the male has mated with. Note that, in haplodiploids, males do not pass their genes to male offspring. The probability that the daughters and the females inseminated by sons will gain any one of the *n* breeding spots is *n/ny*_q_(**v**). Hence, the number of colony-founding individuals of type *s′* in the next generation that descend from a colony-founding male can be written as

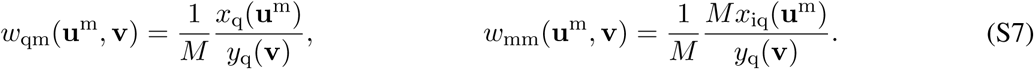

The number of individuals at the end of the season *y*_q_(**v**), *x*_q_(**u**^*s*^) and *x*_iq_(**u**^*s*^) (*s* ∈ {q, m}) in eqs. (S6)–(S7) are determined from eq. (1) (with eq. 3) and eqs. (S3)–(S4) (with eqs. S1 and S2), respectively.

### 1.3 The invasion fitness

We now have all the elements to obtain an expression for the invasion fitness, which allows to ascertain the fate of the mutant allele. Let us denote by *n*_q,u_ (and respectively, by *n*_m,u_) the number of mutant allele copies in females (males with whom the females have mated with), measured at time *t* = *T* in the population. The change in the vector **n**_u_ = (*n*_q,u_, *n*_m,u_)^T^ of number of gene copies from one generation to the next 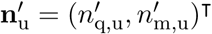, when the mutant allele for a trait that is under genetic control of party *c* ∈ {q, w} is still rare in the population, is given by the matrix

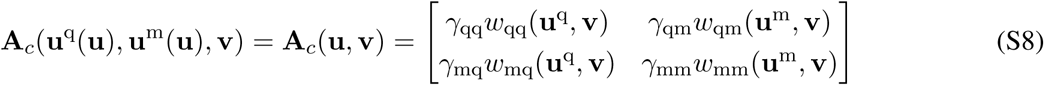

where *γ*_*s′ s*_ is the probability that a gene sampled in an individual of type *s′* ∈ {q, m} was contributed by an individual of type *s* ∈ {q, m}, i.e. a transmission frequency of type *s* to type *s′* (for haplodiploids *γ*_qq_ = 1/2, *γ*_qm_ = 1/2, *γ*_mq_ = 1, *γ*_mm_ = 0). Hence, elements *a*_*s*_*Is* of matrix **A**_*c*_(**u**, **v**) give the expected number of mutant gene copies in a type *s′* ∈ {q, m} individual that descends from an individual of type *s* ∈ {q, m} carrying the mutant allele. Note that in eq. (S8), the dependence on the party *c* ∈ {q, w} who has the genetic control enters into the right-hand-side implicitly via the mutant schedules **u**^q^ and **u**^m^ (recall eq. S1).

The invasion fitness *W*_*c*_(**u**, **v**) of the mutant allele is then given by the leading eigenvalue of the matrix **A**_*c*_(**u**, **v**) (eq. S8), where the subscript *c* ∈ {q, w} emphasizes the party in control of the focal trait. Hence, it satisfies

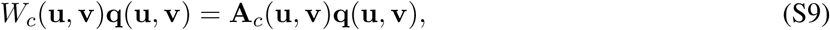

where **q**(**u**, **v**) = (*q*_q_(**u**, **v**), *q*_m_(**u**, **v**))^T^ is the normalized right leading eigenvector of **A**_*c*_(**u**, **v**). Here, normalization means that *q*_q_(**u**, **v**) + *q*_m_(**u**, **v**) = 1. Pre-multiplying eq. (S9) by the vector (1, 1) yields

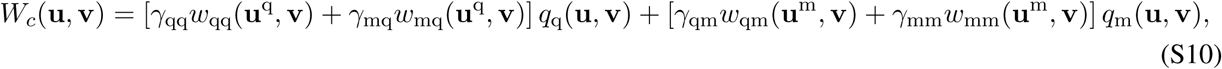

since *q*_q_(**u**, **v**) = (1 − *q*_m_(**u**, **v**)) (see Lehmann et al., 2016, Appendices A-C for more details of how to express invasion fitness in terms of leading left and right eigenvectors of the transition matrix). Note that in eq. (S10), the dependence on the party *c* ∈ {q, w} who has the genetic control, enters into the right-hand-side implicitly via the mutant schedules **u**^q^ and **u**^m^ (recall eq. S1).

The invasion fitness can be interpreted here as the geometric growth rate (generational growth rate) of the mutant allele. This is the asymptotic per capita number of mutant copies produced by the mutant lineage descending from the initial mutation, when overall still rare in the population (see Lehmann et al., 2016 for more details and connections to different fitness measures used in evolutionary biology).

Direct calculation of the normalized right eigenvectors yields

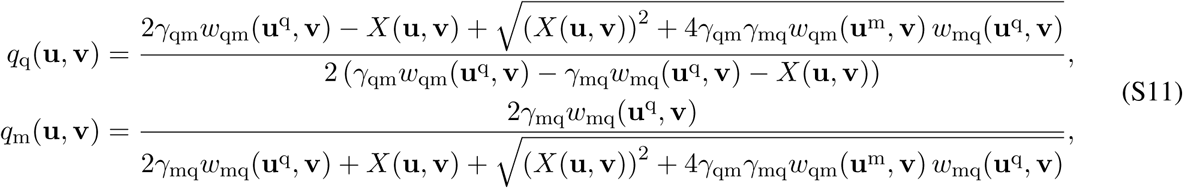

where *X*(**u**, **v**) = *γ*_qq_*w*_qq_(**u**^q^, **v**) − *γ*_mm_*w*_mm_(**u**^m^, **v**).

The quantity *q*_*s*_(**u**, **v**) can be interpreted as the asymptotic probability that a mutant allele is sampled in a class *s* individual. It follows that the maximization of the invasion fitness (S10) depends on both the fitnesses of carriers of the mutant allele (the *w*_*s′ s*_(**u**^*s*^, **v**) functions) and how the mutant allele is distributed across classes (the *q*_*s*_(**u**, **v**) functions which also depend on the evolving traits themselves).

### 1.4 Uninvadable allocation schedule

An uninvadable schedule 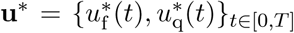 is a resident schedule that is resistant to invasion by any mutant **u** ∈ 𝕌 = 𝕌_f_ *×* 𝕌_q_ schedule. Here, 𝕌 is a set of all possible allocation schedules, while 𝕌_f_ and 𝕌_q_ are sets of full trajectories of the traits **u**_f_ = {*u*_f_ (*t*)}_*t*∈[0,*T*]_ and **u**_q_ = {*u*_q_(*t*)}_*t*∈[0,*T*]_ under consideration. Notice that in order to simplify notations in the main text we used 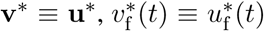, and 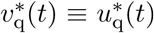 for the uninvadable schedule, but in this S.I. it is more convenient to use the letter *u* for that, basically throughout the S.I. we always distinguish mutant and resident, both at the level of state variables (*x* vs. *y*) and at the level of evolving traits (*u* vs. *v*).

If party *c* ∈ {q, w} is in full control of the two traits (i.e. single-party control), then the uninvadable schedule **u**\* satisfies the condition

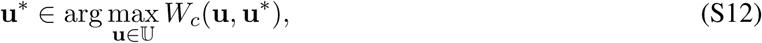

that is, a mutant that adopts the resident schedule **u**\* has the highest invasion fitness from all possible strategies in a population, for a population expressing schedule **u**\*. Hence, an uninvadable schedule **u**\* is a candidate endpoint of the evolutionary process.

Under mixed control, where the queen is in control of the trait *u*_f_ and the workers are in control of the trait *u*_q_, the uninvadable schedule **u**\* satisfies condition

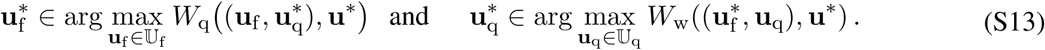

Hence, the uninvadable allocation schedules to individuals of type *k* (*k* ∈ {w, q, m}) can be written as follows

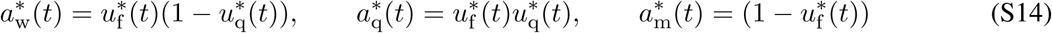

and we denote by 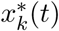 the resulting number of individuals at time *t*.

## 2. First-order condition

In this section, we derive the first-order necessary condition for uninvadability (eq. 5 of the main text) and show that it has a similar structure that first-order conditions in static allocation models (e.g. eq. (1)–(2) in Reuter and Keller, 2001) and applies regardless of colony growth dynnamic. In the next section, we the show how to formulate the necessary first-order condition in terms of pointwise marginal change using optimal control theory and then solve explicitly for the (candidate) uninvadable schedule for our model.

### 2.1 Eigenvalue perturbation

#### 2.1.1 Perturbations in terms of Gâteaux derivatives and relatedness assymetry

We consider a small variation 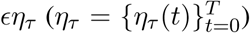 in the trait 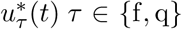 of the uninvadable schedule, such that the mutant trait can be written as

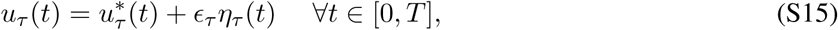

for any feasible deviation *η*_*τ*_ (*t*) (such that 0 ≤ *u*_*τ*_ (*t*) ≤ 1) from the resident schedule 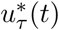, where *ϵ*_*τ*_ ≪ 1 is a small parameter measuring the intensity of the mutant deviation. Hence, we consider a change in the candidate uninvadable allocation trait 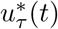 that remains very close to it for all *t* ∈ [0, *T*]. The direction of selection for trait 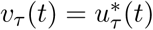 is indicated by the sign of perturbation in invasion fitness

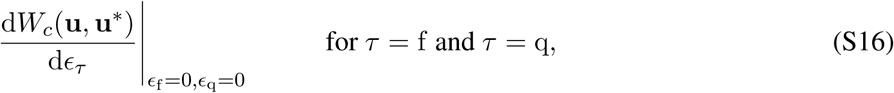

for any feasible deviation *η*_*τ*_ (*t*) from the uninvadable schedule **u**\*. The derivative in eq. (S16) is a Gâteaux derivative (a type of functional or variational derivative) of invasion fitness (e.g., Weber and Arfken, 2003, p. 827–830, Troutman, 2012, p. 45–50, Luenberger, 1997, p. 171–178, Gelfand and Fomin, 1963, p. 54–63). In other words, it gives the infinitesimal change in invasion fitness resulting from a change in the whole mutant schedule into the direction of *η*(*t*) (Gâteaux derivative can be thought of as a generalization of directional derivative from differential calculus). Gâteaux derivatives are useful to generalize evolutionary stability conditions (e.g., Eshel, 1983, eq. 3, Taylor, 1989, eq. 2.1) to function-valued traits.

Because the functional derivative, d*W*_*c*_(**u**, **u**\*)/ d*ϵ*_*τ*_ is an ordinary function in *ϵ*_*τ*_, it follows from standard results of eigenvalue perturbation (Caswell, 2001, p. 209, eq. 9.10) that

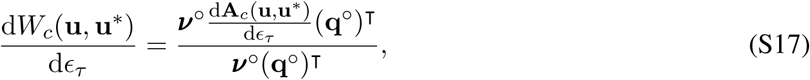

where superscript ⊤ denotes transpose, 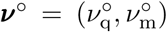 is a vector of neutral reproductive values of colony-founding individuals of type *s* ∈ {q, m} and 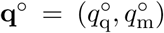 is a vector of the neutral frequencies of class *s* ∈ {q, m} individuals. Throughout, the superscript ° will denote a quantity that is evaluated in the absence of natural selection, i.e., by a process determined by the monomorphic resident population. Substituting eq. (S8) into (S17) and given that ***v***°(**q**°)^T^ = 1 (total class reproductive values of all individuals add up to one) yields

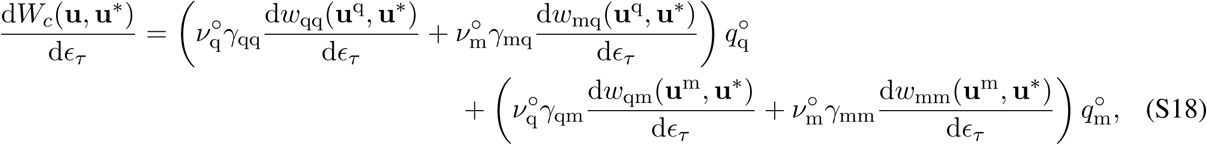

where all derivatives are evaluated at *ϵ*_*τ*_ = 0.

In the absence of natural selection, the number of gene copies from one generation to the next can be described by a matrix

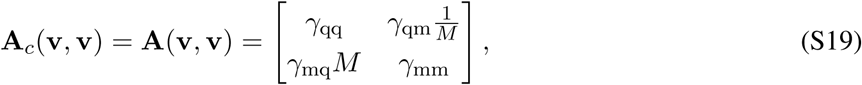

which does not depend on the mode of control and whose dominant eigenvalue is one (given that *γ*_qq_ = 1/2, *γ*_qm_ = 1/2, *γ*_mq_ = 1, *γ*_mm_ = 0). The reproductive values ***v****°* and class frequencies **q***°* are, respectively, given by the left and right unit eigenvectors of **A**(**v**, **v**), and we normalize these vectors such that the total class reproductive values defined by

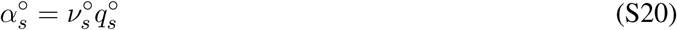

(e.g., Rousset, 2004; Taylor, 1990; Taylor and Frank, 1996) of all individuals add up to one: 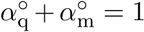. This normalization entails the use of the perturbation formula eq. (S18) (e.g., Caswell, 2001), with which we obtain

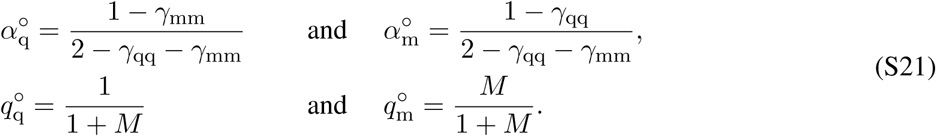

It follows from the class frequencies 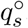, that under neutrality there are *M* times as much colony-founding males than females, which is in accordance with the fact that females mate *M* times.

Substituting the transmission frequencies *γ*_*s′ s*_ for haplodiploids [*γ*_qq_ = 1/2, *γ*_mq_ = 1, *γ*_qm_ = 1/2, *γ*_mm_ = 0] then we have the class reproductive values for haplodiploids

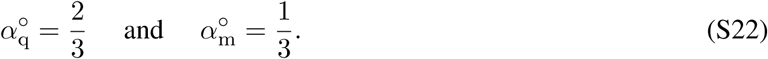

In eq. (S18), the derivative d*w*_*s′ s*_(**u**^*s*^, **v**)/ d*ϵ*_*τ*_ is the total variation of individual fitness with respect to mutant values, which acts on **u**^*s*^ (by way of eq. S1). By substituting eq. (S15) into eq. (S1) (where we take 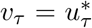), we have for *τ* ∈ {f, q} that

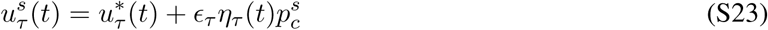

and owing to eq. (S15) and the constant factor rule in differentiation, we can write

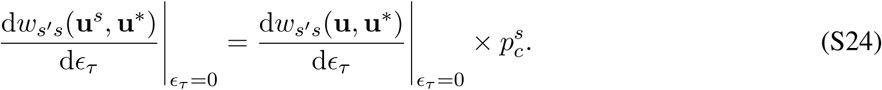

Substituting eq. (S24) into eq. (S18), we have for control mode *c* ∈ {q, w} that

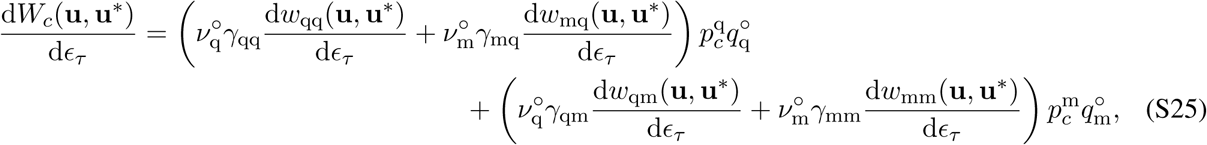

where all derivatives are evaluated at *ϵ*_*τ*_ = 0 and thus all trait values (allocation schedules) are set to the resident schedule **v**. Substituting eq. (S20) into eq. (S25) yields

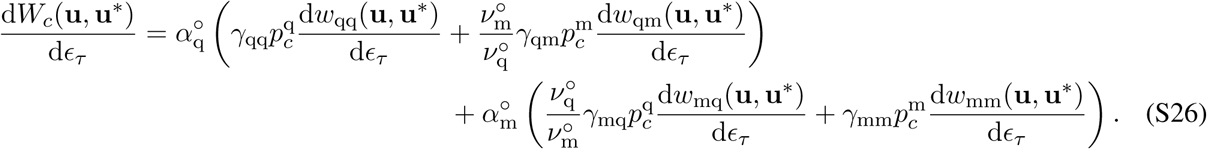

Substituting the fitness functions (S6)–(S7) into eq. (S26) yields

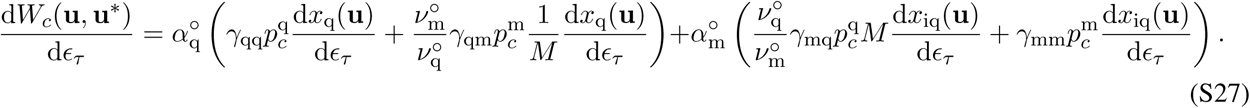

By considering that eqs. (S20) and (S21) yield that 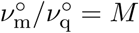 and 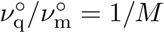, then eq. (S27) as

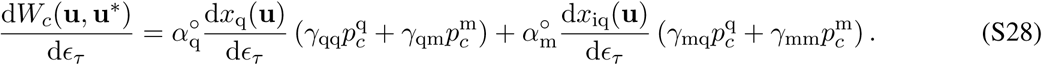

Rearranging, we can write eq. (S28) as

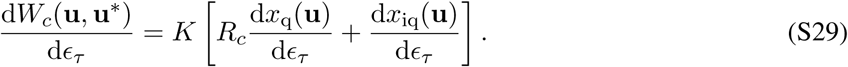

where 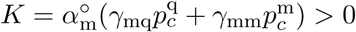 is a positive constant and

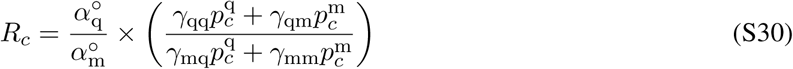

is the so-called relatedness asymmetry (see Boomsma and Grafen, 1991, p. 386 and section 2.1.2 for the biological interpretation).

#### 2.1.2 Interpretation of relatedness asymmetry

The *relatedness asymmetry R*_*c*_ gives the ratio of sex-specific potentials for the party *c* in control to contribute genes into the distant future (Boomsma and Grafen, 1991, p. 386). In order to see this, we note that owing to eq. (S21), the first ratio in eq. (S30) is the ratio 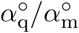 of class reproductive values. Furthermore, notice that 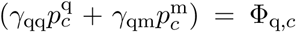 and 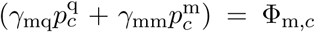 are the probabilities that a gene randomly sampled in a recipient female and male, respectively, is identical-by-descent to a gene randomly sampled from party *p* in control of resource allocation; that is, the *coancestry* (or *consanguinity*) between a female (male) and the (average) individual whose genes are in control of the resource allocation trait. Hence, the relatedness asymmetry is

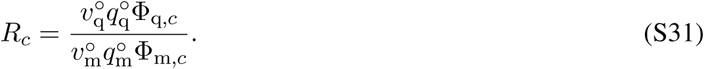

Here, 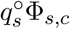 is the asymptotic probability that a randomly sampled gene from a colony-founding individual finds itself in an individual of type *s* and is a replica copy of a gene sampled from a party *c*. Then, since 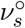 is the long-term contribution of genes in individual of type *s* to the gene pool, we can interpret the relatedness asymmetry as giving the ratio of sex-specific potentials for party *p* in control to contribute (in a neutral process) to the gene pool in the distant future.

Since the ratio of consanguinity is equivalent to the ratio of relatedness, we can write the second ratio in eq. (S30) as 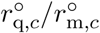, where 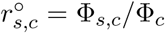 is the relatedness between an individual of type *s* and the average individual whose genes are in control of the resource allocation trait, and this depends on the coefficient of coancestry Φ_*c*_ of the average individual in control of the resource allocation trait with itself (i.e. the probability that two homologous genes, drawn randomly with replacement from party *c*, are identical by descent). With this eq. (S31) is also

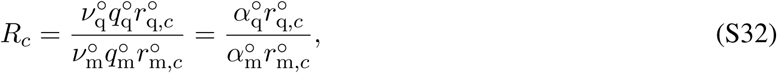

where the second equality displays the classical form of the relatedness asymmetry (Boomsma and Grafen, 1991, p. 386). For haplodiploids eq. (S32) simplifies to

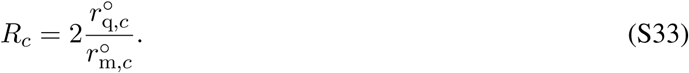

Since relatedness if given by the ratio of the coefficient of coancestry of party *c* with an individual of type *s* (Φ_*s,c*_, which is given by the transmission frequencies *γ*_*s′ s*_ and the expected frequency 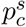 of mutant allele residing in party *c*) to the coefficient of coancestry Φ_*c*_ of party *c* with itself (Φ_q_=Φ_w_=1/2). Substituting the frequencies for haplodiploids [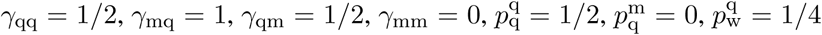, and 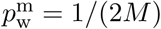] gives the relatedness coefficients for haplodiploids

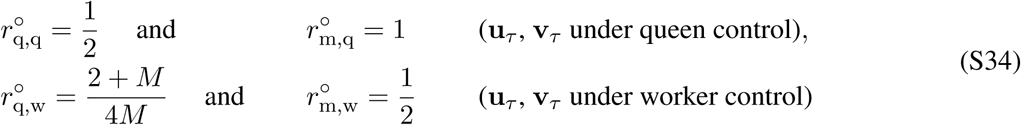

which are classical expressions (e.g., Frank, 1998, Fig. 10.4, p. 209). Substituting the relatedness coefficients into eq. (S33) yields the relatedness asymmetry for haplodiploids

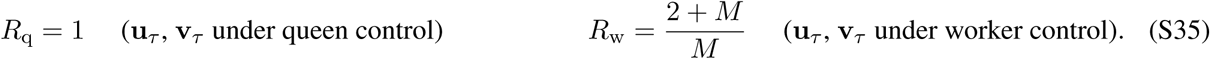

### 2.2 First-order condition for uninvadability

The necessary first-order condition for the candidate uninvdable schedule 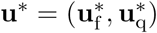 is given by

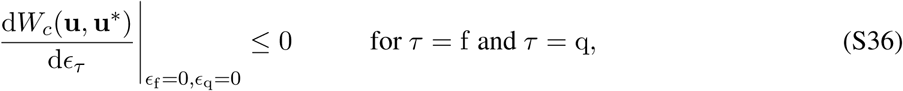

for any feasible deviation *η*(*t*). Substituting eq. (S29) into (S36) yields that we can express the necessary first-order condition for uninvadability under queen (*c* = q) or worker (*c* = w) control as

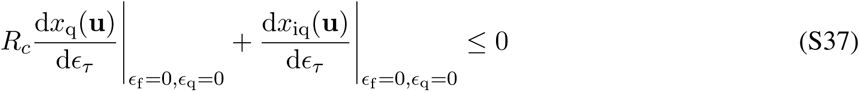

and under mixed control as

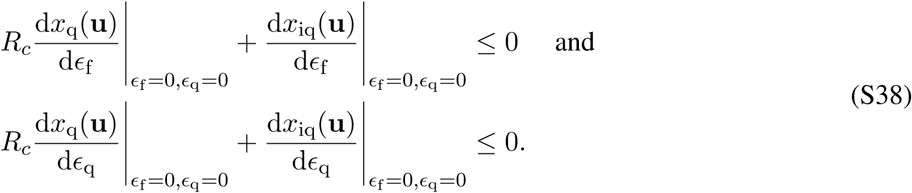

Hence, the first-order condition given by eqs. (S37) and (S38) can be expressed in terms of variational derivatives d*x*_*k*_(**u**)/ d*ϵ*_*τ*_ and relatedness asymmetry *R*_*c*_. The variational derivative d*x*_*k*_(**u**)/ d*ϵ*_*τ*_ measures the change in the number of individuals of type *k* ∈ {q, iq} associated with a focal colony where phenotype **u** is expressed. In the next section we give the interpretation for relatedness asymmetry. Note that the first-order condition given by eqs. (S37) and (S38) is a dynamic version of first-order condition in a comparable static allocation model (e.g. eq. (1)–(2) in Reuter and Keller, 2001). Note that we the first-order condition (given by eqs. S37 and S38) in the main text (recall 5) using a different notation (to simplify the readability for the general audience), where *y*_q_(**u**) *≡ x*_q_(**u**) and *y*_iq_(**u**) *≡ x*_iq_(**u**).

### 2.3 Pointwise eigenvalue perturbation

It is useful to also consider pointwise perturbations in invasion fitness, which would allow to describe the direction of selection on trait 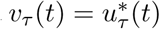 for each *t*. That is, we consider for every *t* ∈ [0, *T*]

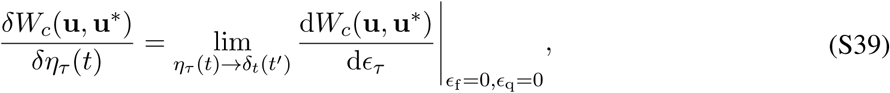

where the derivative on the left-hand-side is a pointwise functional derivative (the so-called Volterra derivative) of invasion fitness at time *t* ∈ [0, *T*] (see e.g. Parr and Yang, 1989, p. 246-247 and eq. (3a) in Dieckmann et al., 2006) and *δ*_*t*_(*t′*) = *δ*(*t′ - t*) is the Dirac delta function, which is 0, except at *t′* = *t*, when it is 1 (here, *t′* is just a dummy variable for time *t′* ∈ [0, *T*]) Note that using the *δ*-notation (not to be confused with the Dirac delta function) to refer to the pointwise functional derivative is standard notation in the physical literature (see e.g. Giaquinta and Hildebrandt, 1996, p. 18).

It follows from eq. (S29) that we can express the pointwise perturbations as follows

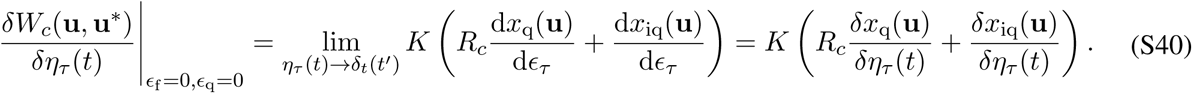

### 2.4 Pointwise first-order condition for a singular arc and the marginal substitution rate

We call the uninvadable allocation trait 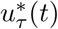 a singular arcs, when it does not reside on the bounds of the feasible set (i.e., when 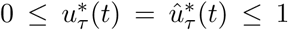) over a finite period of time. Now we will show that the pointwise first-order conditions for singular arcs can be expressed in terms of marginal substitution rates. We will show in section 3 (see eqs. S66 and S73) that at the singular arc

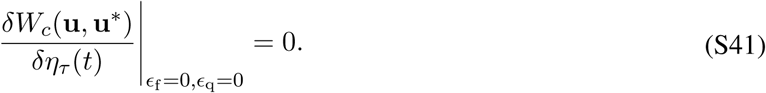

Hence, we can express the necessary first-order condition for the singular arc 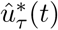 to be uninvadable under queen (*c* = q) or worker (*c* = w) as

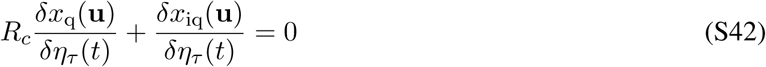

and under mixed control as

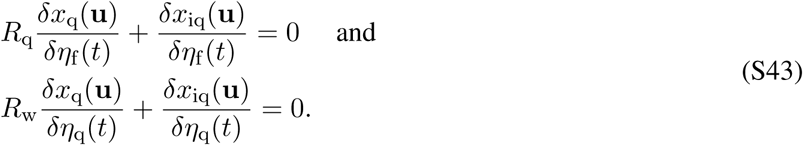

Rearranging eqs. (S42) and (S43) yields for queen (*c* = q) and worker (*c* = w) control

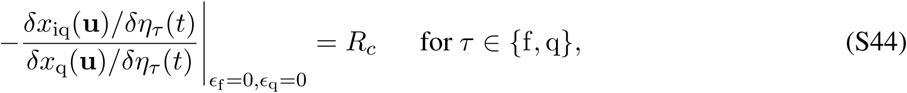

and for mixed control

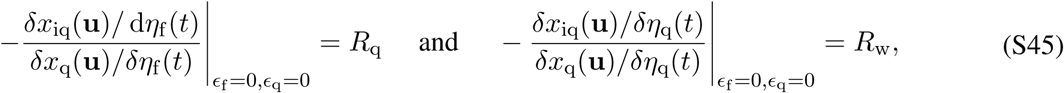

The left-hand side in eqs. (S44) and (S45) gives the ratio of the marginal change in the number of inseminated queens to the marginal change in the number of queens produced when the allocation schedule is varied. This ratio is expressed in terms of a variational derivatives *δx*_*k*_(**u**)/*δη*_*τ*_ (*t*) measuring the change in the number of individuals of type *k* ∈ {q, iq} associated with a focal colony where phenotype **u** is expressed. Hence, we have showed that when 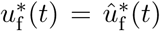 is a singular arc, the marginal substitution rate of inseminated queens with produced queens is given be the relatedness asymmetry *R*_*c*_.

#### 2.4.1 The critical sex ratio under delayed dispersal

In this section, we derive the condition for the (critical) sex ratio at the end of the season (*t* = *T*) under delayed dispersal. Equation (S4) for delayed dispersal together with eq. (S5) yields that

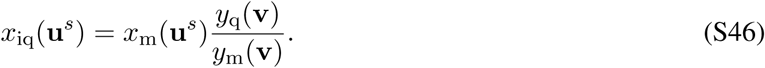

Hence, it follows from eq. (S23) that under delayed dispersal

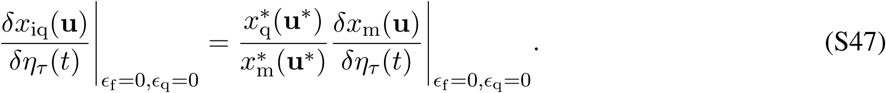

We will show later in section 5 that under delayed dispersal 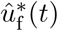 is a singular arc during 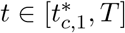, where 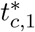 is the time, when 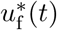 becomes a singular arc under the control mode *c* ∈ {q, w, mx}. Substituting eq. (S47) into eqs. (S44)–(S45) yields for queen (*c* = q) and worker (*c* = w) control

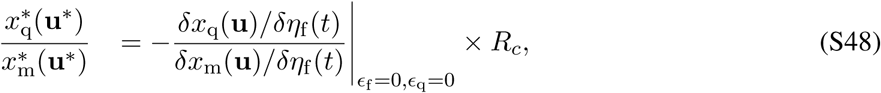

and mixed control

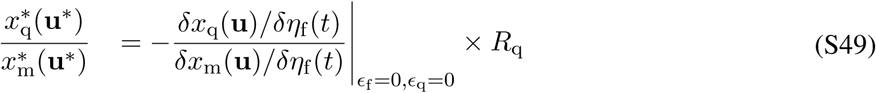

where the right-hand side depends on the ratio of the marginal values of males relative to queens (i.e., the marginal rate of substitution of producing males instead of new queens). If males and juvenile queens are equally costly to produce and they have the same mortality rate (*µ*_q_ = *µ*_m_ = *µ*_r_) and the same growth schedule, then the marginal product is negative one. Hence, the (critical) sex ratio at the end of the season (*t* = *T*) under delayed dispersal for equal mortality rates of males and queens is equal to the relatedness asymmetry (recall eq. S35), i.e.

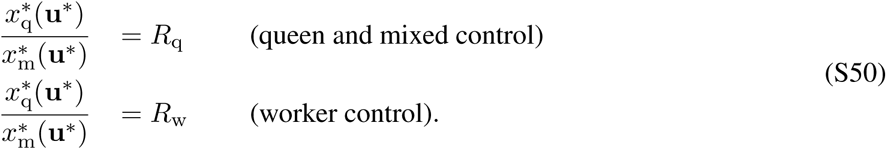

## 3 First-order condition expressed in terms of optimal control problem

In this section, we use optimal control theory in order to solve the marginal value equation (S44) for different scenarios of our model by way of applying Pontryagin’s weak maximum principle (e.g., Bryson and Ho, 1975; Sydsæter et al., 2008 for broad introductions and Day and Taylor, 2000; González-Forero et al., 2017; Iwasa and Roughgarden, 1984; Macevicz and Oster, 1976; Perrin, 1992 for previous application to evolutionary biology).

### 3.1 Formulation of the optimal control problem

#### 3.1.1 The basic problem

We start by formulating the dynamic resource allocation problem as a classical optimal control problem. Finding the candidate uninvadable schedule entails establishing an optimal pair

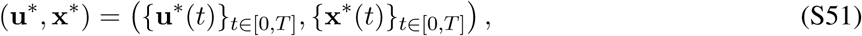

where 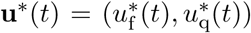 and 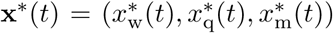 are vectors of uninvadable (optimal) control and state variables, respectively. The optimal pair (**u**\*, **x**\*) is a solution to eqs. (S12) and (S13) under single-party control and mixed control, respectively. That is, it maximizes the invasion fitness *W*_*c*_(**u**, **v**) (as given by eq. S10).

The so-called control variables for the maximization problems are the (resource allocation) phenotypes expressed in colonies founded by individuals who are homozygous for the mutant allele (recall eq. S1)

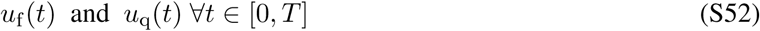

Where

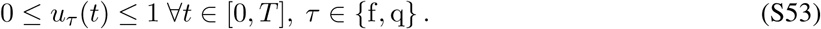

For delayed dispersal, the vector of the so-called state variables for the optimal control problems can be expressed as

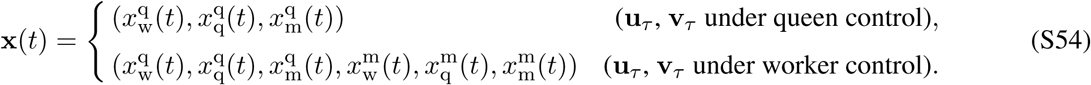

For direct dispersal, the vector of state variables for the optimal control problems can be expressed as

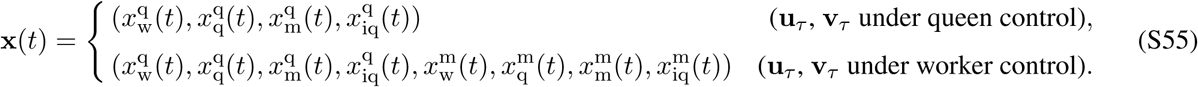

In addition, the vector of dynamical variables involved in the invasion fitness (eqs. S62–S62) for the optimal control problems can be expressed as

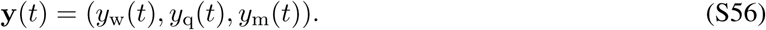

The rate of change in state variables appearing in eqs. (S54)–(S55) is described by the differential equations

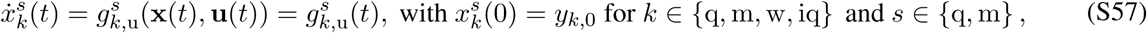

where upper “·” denotes the time derivative, (*y*_w,0_, *y*_q,0_, *y*_m,0_, *y*_iq,0_) = (1, 0, 0, 0) (fixed) and 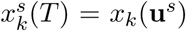 is free and the differential equations can be expressed as

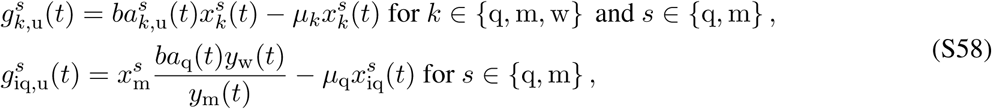

where the mutant allocation schedules 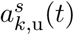 and resident allocation schedules *a*_*k*_(*t*), are given by eqs. (S2) and (3) of the main text, respectively.

The rate of change in dynamic variables appearing in eq. (S56) is described by the differential equations

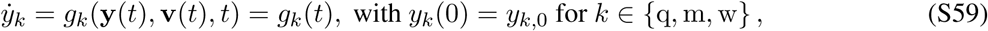

where (*y*_w,0_, *y*_q,0_, *y*_m,0_) = (1, 0, 0) (fixed) and *y*_*k*_(*T*) = *y*_*k*_(**v**) is free and the differential equations can be expressed as

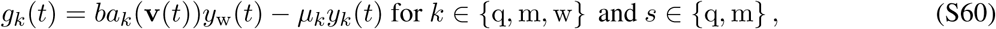

where the resident allocation schedules, *a*_*k*_(**v**(*t*)), are given by and eq. (3) of the main text.

#### 3.1.2 Explicit expression for invasion fitness function

Henceforth, we write the invasion fitness of a mutant allele as *W*_*c*_(**u**, **v**) *≡ W*_*c,d*_(**u**, **v**), where the additional subscript *d* ∈ {del, dir} emphasizes the scenario of dispersal of sexuals, delayed and direct dispersal, respectively. Substituting the transmission frequencies for haplodiploids (*γ*_qq_ = 1/2, *γ*_mq_ = 1, *γ*_qm_ = 1/2, *γ*_mm_ = 0) into eq. (S10) and using eq. (S11) we can simplify the expression for the invasion fitness (eq. S10) under delayed dispersal to

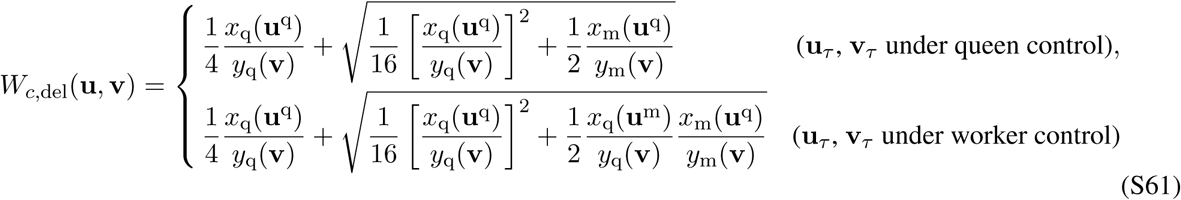

and under direct dispersal to

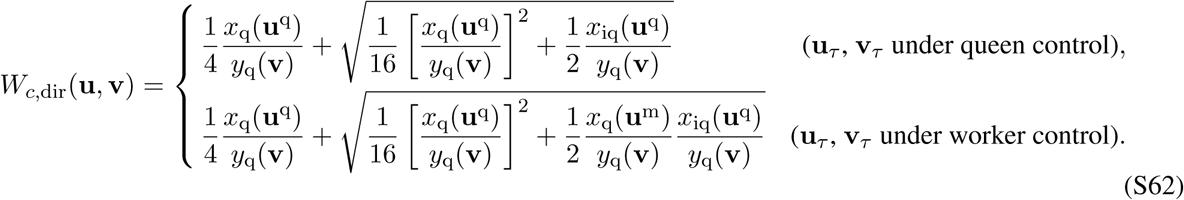

Note that for mixed control we have the invasion fitness function under queen control *W*_q,*d*_(**u**, **v**) to determine 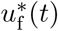 and the invasion fitness function under worker control *W*_w,*d*_(**u**, **v**) to determine 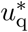 These simplified expressions of invasion fitness will turn out useful for solving numerically the optimal control problem (see section 11) and also conceptually, because it makes it explicit how the invasion fitness depends on the state **x**(*t*) and dynamic variables **y**(*t*).

### 3.2 Pontryagin’s weak maximum principle and the Hamiltonian

The necessary first-order condition for uninvadability (given by eq. S36) can be expressed in terms of pointwise functional derivatives (given by eq. S39, see e.g. Parr and Yang, 1989, p. 246); that is

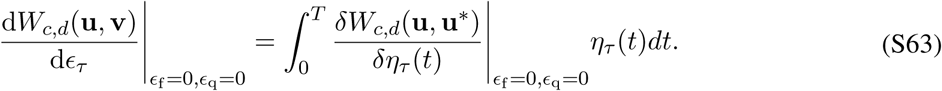

This expression can be thought of as a functional analogue of the formula for the total derivative of a function *W* (*η*_1_(*t*), *η*_2_(*t*), *…*): d*W/* d*t* = Σ_*i*_(*∂W/∂η*_*i*_)(*∂η*_*i*_/*∂t*) (see e.g. Parr and Yang, 1989, p. 246).

Hence, the first-order condition for uninvadability (eq. S36) can be expressed in terms of point-wise marginal change, which can be expressed under single-party control as

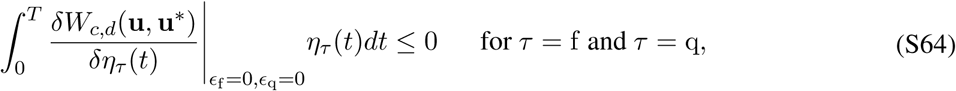

and for mixed party control as

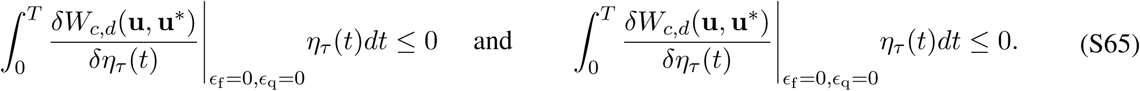

for any feasible *η*_*τ*_ (*t*).

Pontryagin’s weak maximum principle yields that the pointwise marginal change (as in eq. S63) can be restated for both single-party (see e.g. Speyer and Jacobson, 2010, p. 61) and mixed control (see e.g. Mazalov, 2014, p. 372, Theorem 10.8) as follows

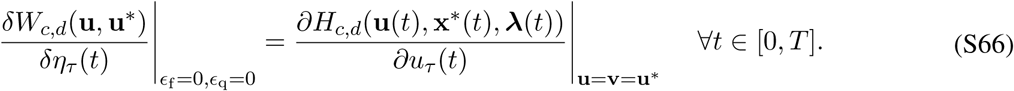

Here, *H*_*c,d*_(**u**(*t*), **x**\*(*t*), ***λ***(*t*)) is the Hamiltonian function, which allows to transform a dynamic optimization problem into a sequence of static optimization problems by providing a marginal condition that must be satisfied over the whole time schedule (see section 3.3 for the full interpretation of the Hamiltonian), and which for our problem can be expressed as

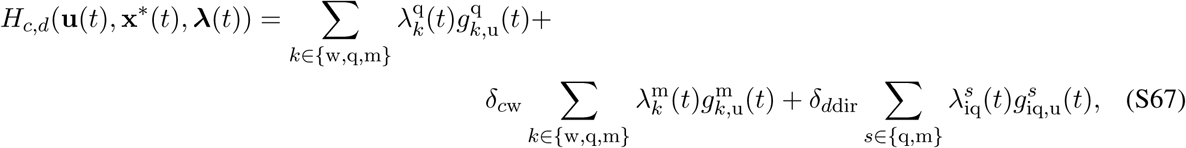

where index *c* ∈ {q, w, mx} emphasizes the mode of control and *d* ∈ {del, dir} emphasizes the time of dispersal of sexuals. In eq. (S67), *δ*_*ij*_ is the Kronecker delta function, i.e.

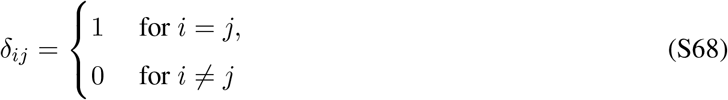

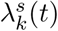 is a costate variable associated with the state variable 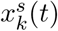 and ***λ***(*t*) is a vector of costate variables and for delayed dispersal it can be expressed as

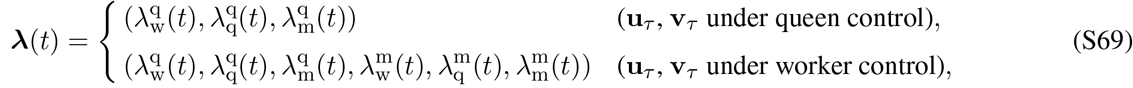

and for direct dispersal it can be expressed as

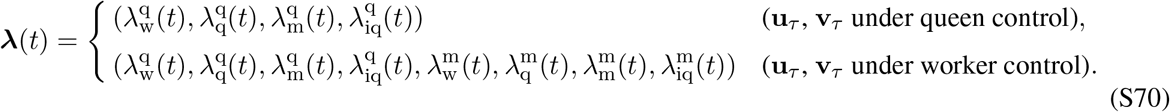

The differential equations for the costate variables appearing eqs. (S69)–(S70) are given by the derivatives of the Hamiltonian with respect to the corresponding state variables, i.e.

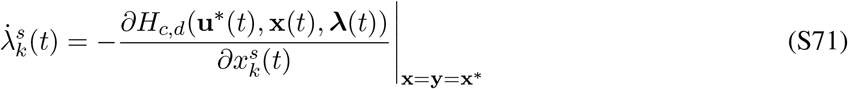

Since **x**(*T*) is free, the transversality conditions for the co-state variables are given by

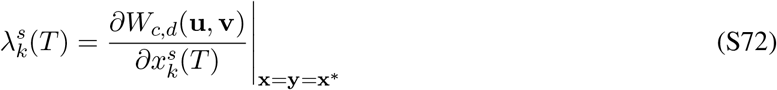

(e.g., Bryson and Ho, 1975; Sydsæter et al., 2008).

### 3.3 Interpretation of the Hamiltonian and costate variable

The quantity *H*_*c,d*_(**u**(*t*), **x**\*(*t*), ***λ***(*t*)) d*t* = *H*_*c,d*_(*t*) d*t* can be interpreted as the total contribution to the invasion fitness *W*_*c,d*_(**u**, **v**) by an increase in the production of individuals of different types for a certain (constant) allocation schedule 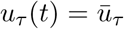 during the interval [*t, t* + d*t*] (e.g. Dorfman, 1969, Sethi and Thompson, 2006, p. 34). As a consequence, the control variables *u*_*τ*_ (*t*) for a given interval should be chosen such that to maximize *H*_*c,d*_(*t*). This implies that the dynamic optimization problem of maximizing the invasion fitness *W*_*c,d*_(**u**, **v**) can be transformed into a sequence of static problems of maximizing the corresponding Hamiltonian *H*_*c,d*_(*t*) at instants *t* ∈ [0, *T*]. Hence, the Hamiltonian can be interpreted as a rate at which the invasion fitness (which is defined at final time *T*) increases at time *t* and *∂H*_*c,d*_(*t*)/*∂u*_*τ*_ (*t*) represents a variation in invasion fitness due to a unit impulse (Dirac function) in *u*_*τ*_ (*t*) at time *t*, while satisfying the state equations (Bryson and Ho, 1975, p. 49). More precisely, *∂H*_*c,d*_(*t*)/*∂u*_*τ*_ (*t*) measures the net effect on invasion fitness that the marginal change in the trait value *u*_*τ*_ (*t*) has through immediate change in the trait value *u*_*τ*_ (*t*) at time *t* and through the cascading effects that this change has on the invasion fitness by changing the state variables (the numbers of individuals of different types) from time *t* onward until time *T*.

A costate variable 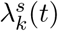 can be interpreted as the effect on invasion fitness for a marginal change in the corresponding state variable 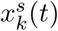 when colony resources are allocated to production of type *k* individuals at time *t* in a colony founded by a mutant individual of type *s*. Therefore, informally, a costate variable 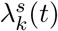 gives the value (measured as the increase in invasion fitness) of each unit resource invested at time *t* into the production of type *k* individuals at time *t* in a colony founded by a mutant individual of type *s*. Hence, the costate variables are of extreme importance since only the individuals that yield the highest investment value in invasion fitness should be produced at any given time.

### 3.4 Derivatives of the Hamiltonian

It follows from eqs. (S64), (S65), (S66) and (S53) (see e.g. Kamien and Schwartz, 2012, p. 185-186 for full explanation) that for all *t* ∈ [0, *T*]

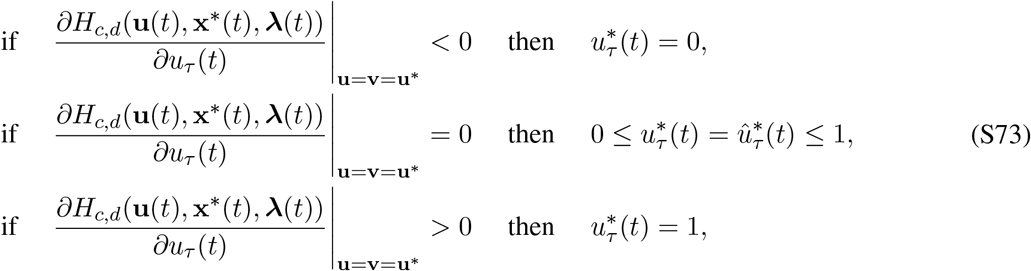

where 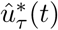 denotes that the control 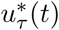 is a singular arc (Sethi and Thompson, 2006, p. 407). An allocation trait is a singular arc 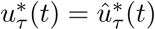 when the Hamiltonian is linear (or more strictly, affine) in the control and the derivative *∂H*_*c,d*_(**u**(*t*), **x**\*(*t*), ***λ***(*t*))/*∂u*_*τ*_ (*t*)|_**u**=**u**_* = 0. Hence, the first-order condition (given by eq. S64 or eq. S65) is satisfied, but the control variable *u*_*τ*_ (*t*) does not directly appear in the first-order condition. More generally, an optimal control is a singular arc, if the value of variational Hamiltonian is unchanged to the second order from a weak first-order variation of the control at each point of the arc (Robbins, 1967).

In order to ascertain the uninvadable allocation schedule **u**\* from eq. (S73), we need to determine the derivatives of the Hamiltonian with respect to *u*_*τ*_ (*t*). Substituting eq. (S58) into eq. (S67) and taking the derivative with respect to *u*_*τ*_ produces

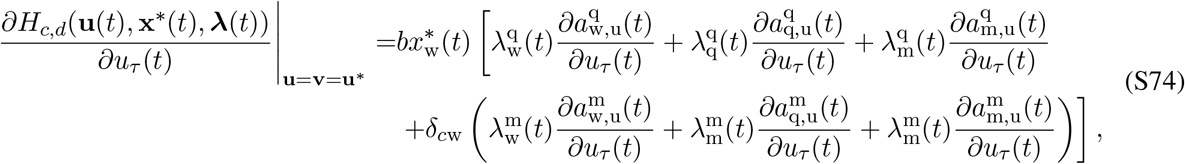

with partial derivatives

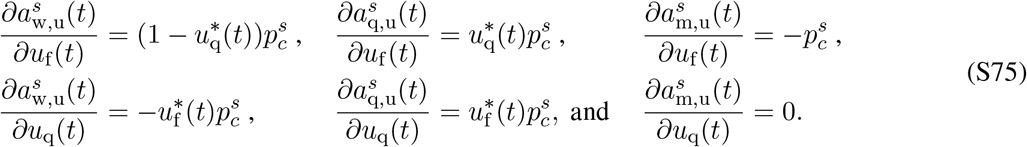

Hence, the derivatives of the Hamiltonian with respect to controls *u*_f_ and *u*_q_ can be written as

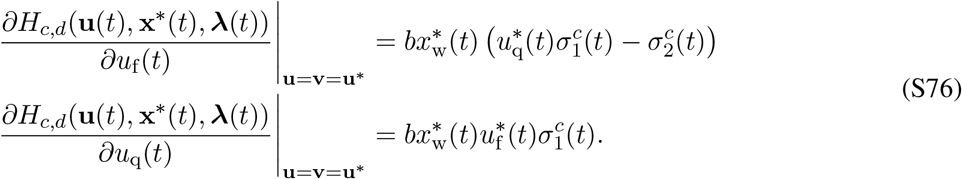

Expressions 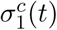 and 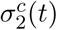 and 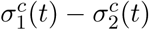 in eq. (S76) are functions of the costate variables 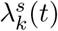 and the expected frequencies 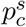 of the mutant allele in the party *c*

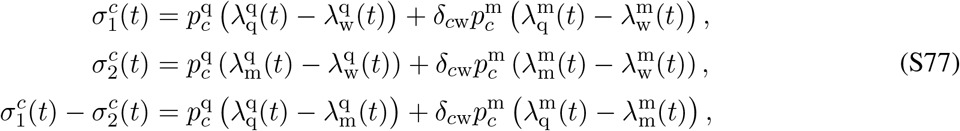

where 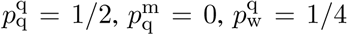, and 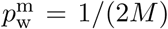. It follows from eq. (S76) and (S73) that 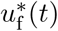 is determined from the sign of expression 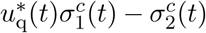 and 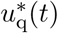 is determined from the sign of expression 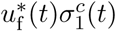, since *b >* 0 and 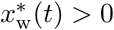 for biological reasons. Since, the signs of functions 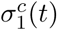 and 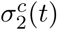 will be instrumental in determining the signs of these expressions, we call them the switching functions, which are analogous to the switching functions in linear optimal control problems (e.g., Bryson and Ho, 1975, p. 111).

Substituting the expected frequencies 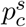 of the mutant allele in the party *c* and eq. (S91) into eq. (S77) yields that for queen control

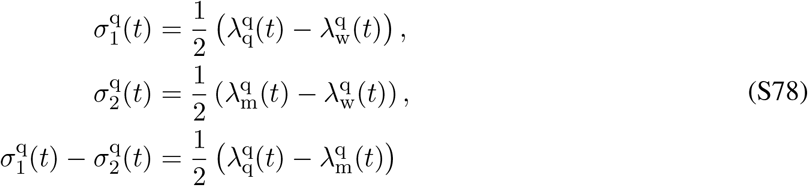

and for worker control

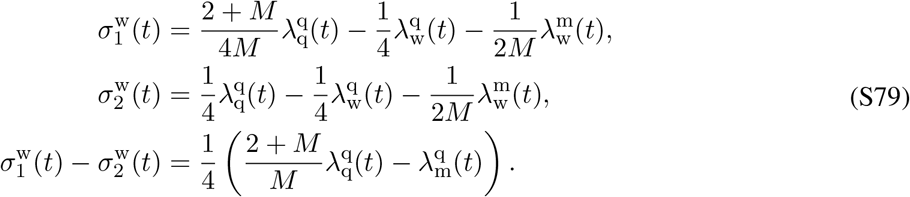

Under delayed dispersal if the mortality rates of males and queens are equal (i.e. *µ*_q_ = *µ*_m_ = *µ*_r_), it follows from the costate equations (S82) and from the transversality conditions (S86) that the switching functions for worker control further simplify to

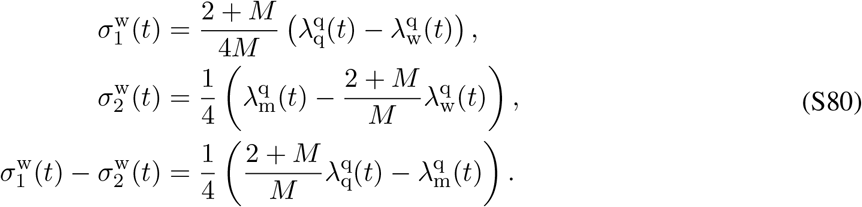

## 4 Global qualitative properties of the uninvadable allocation schedule

Here we describe the scheme of deriving the uninvadable allocation schedule **u**\* under different assumptions of the model. First we present the conditions that the candidate allocation schedule has to satisfy to be consistent with the first-order condition for uninvadability, which gives rise to different phases of colony growth. Then we describe the scheme of determining the uninvadable allocation schedule that consists of these possible phases.

### 4.1 Conditions for candidate uninvadable allocation schedules

We now have all the elements to characterize the first-order conditions given by eqs. (S64) and (S65). We have from eqs. (S73) and (S76) that the conditions for the candidate optimal controls can be expressed as

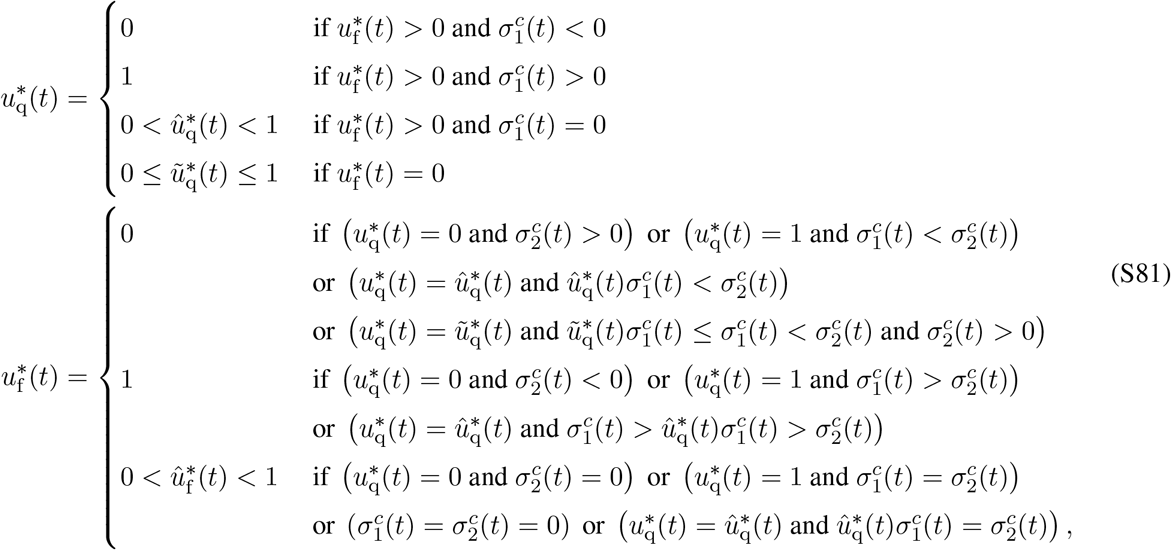

where 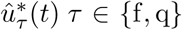 denotes that the uninvadable control variable 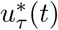 is a singular arc and 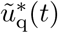 denotes that 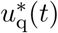 can not be determined and hence can take any value in the range [0, 1] (reflecting the fact that during the phase when only males are produced the trait that affects how resources are allocated between different types of females does not have any effect on the allocation to individuals of different types). It follows from the conditions for the candidate optimal controls, given by eq. (S81), that there are seven possible phases of colony growth (see table S1). We will call phase W (exclusive worker production) the ergonomic phase of colony growth, while phases F (production of only queens), M (production of only males) and FM (production of queens and males simultaneously) are called the reproductive phases, since all the colony resources are directed towards producing sexuals. During phases WF, WM, FM, WFM there is simultaneous production of individuals of different types and in the context of our optimal control problem, it means that one or both of the control variables are singular arcs. In the following sections, we determine from the candidate optimal controls given by eq. (S81), the uninvadable allocation schedules for different assumptions of the time of dispersal of sexuals and different scenarios of genetic control.

**Table S1:**
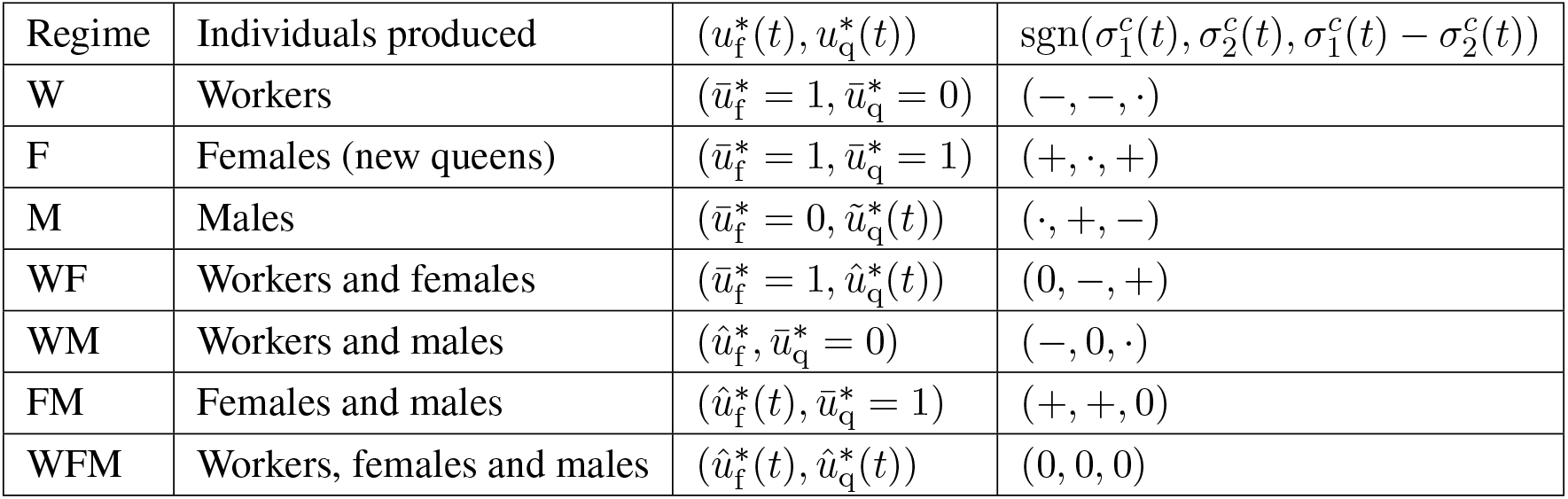
Candidate optimal controls and conditions for the signs of switching functions for all possible regimes of colony growth. Note that “*·*” means any sign, 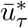 denotes that 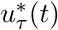 is constant, 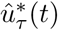 denotes that 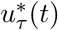 is a singular arc (if the singular arc is constant we write it as 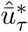) and 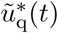 denotes that 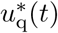 is undetermined because no females are produced, since it does not appear in the Hamiltonian when 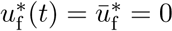.

### 4.2 Short description of the derivations of the analytical results

We describe the scheme of obtaining the uninvadable allocation schedule **u**\* that consists of possible phases outlined in the table S1. The uninvadable allocation schedule **u**\* can explicitly be determined from the table S1, given that we know the switching functions 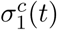 and 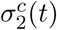 throughout the period *t* ∈ [0, *T*]. According to eq. (S77) the switching functions depend on the costate variables 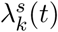 and the expected frequencies 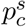 of the mutant allele in party *c*. Hence, establishing the uninvadable allocation schedule **u**\* reduces to determining the costate variables throughout the period *t* ∈ [0, *T*]. The dynamical behaviour of the costate variables is given backwards in time by the differential equations in eq. (S71) and the terminal conditions (transversality conditions) in eq. (S72). During candidate phases W, F, and M of colony growth, the allocation variables 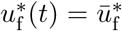 and 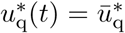 are constant. Hence, it will turn out to be useful to derive the equations for costate variables from eq. (S71), assuming that 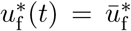 and 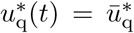 are constant (see eqs. S90–S91). The costate variables at time *t* = *T* are given by the transversality conditions (S83), (S86), and (S87), which deduce from eq. (S72). In addition, we derive eqs. (S88) and (S89) that describe the dynamical behaviour of the state and dynamic variables, respectively, which are solutions to eqs. (S57), (S58), (S57), and (S58) assuming that 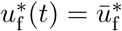 and 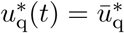 are constant. These expressions together with table S1 give the necessary elements to obtain the uninvadable allocation schedule **u**\*.

We then proceed by determining the switching functions 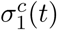 and 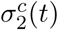 backwards in time. At time *t* = *T*, the switching functions can be directly determined by substituting the transversality conditions (S83), (S86), and (S87) into eq. (S77). Considering that the costate variables are continuous, then it follows that we can determine **u**\*(*t*) for some time interval 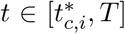 before the end of the season, where 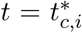 is the time for which one or more of the switching functions (S77) change their signs. Here, the subscript “*c*” emphasizes the scenario of genetic control and “*i*” denotes the number of switches between different phases that make up the uninvadable allocation schedule **u**\*. Hereby, we have established the uninvadable allocation strategy **u**\*(*t*) for the last phase 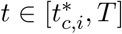. We proceed by using information about **u**\*(*t*) during the last phase and determining the switching functions 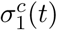 and 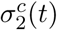 for the penultimate phase 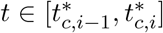. We iterate this process, until the switching functions will not change their signs any more. This scheme allows us to determined all the *i* + 1 phases of the uninvadable allocation schedule **u**\* and all the switching times 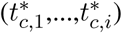 between the different phases.

Since the properties of the uninvadable resource allocation strategies under delayed dispersal and direct dispersal turn out to be substantially different, then we will present the derivations for these two cases separately. For delayed dispersal, we were able to derive the uninvadable resource allocation schedule analytically only assuming equal mortality of sexuals *µ*_q_ = *µ*_m_ = *µ*_r_. For direct dispersal, we were able to derive the full uninvadable resource allocation schedule analytically under single party control only for when *R*_*c*_*µ*_q_ *≥ µ*_m_ and under mixed control only for when *µ*_q_ = *µ*_m_.

Our analytical results are in accordance with the numerical results derived with GPOPS that employs the direct approach of finding the uninvadable allocation schedule instead of the indirect approach given by Pontryagin’s maximum principle. Note that the derivation of some of our analytical results (for direct dispersal) entails using the intuition from our numerical results about the general properties of the optimal allocation schedule and then verifying that this solution satisfies the first-order conditions for uninvadability (eqs. S64–S65).

### 4.3 State and costate variables

We outlined in section 4.2 that in order to determine the univadable allocation schedule **u**\* from the first-order condition, we need to derive the costate equations, transversality conditions and the equations that describe the dynamics of the costate and the state variables for a period of time [*t*_0_, *t*_1_] during which the resource allocation is constant. We now present these expressions.

#### 4.3.1 Costate equations and transversality conditions

Substitution of eqs. (S58) and (S67) into eq. (S71) yields the following differential equations for the costate variables

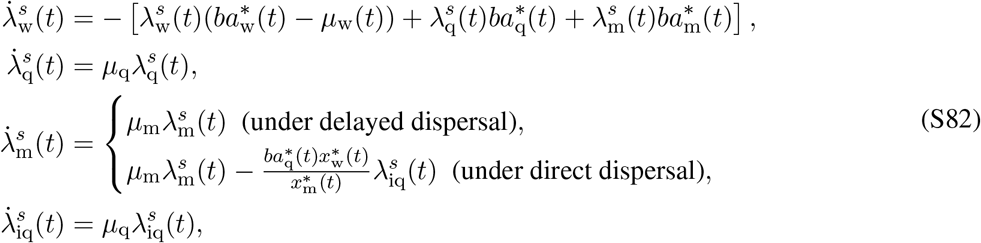

where 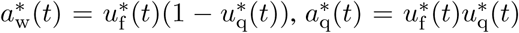, and 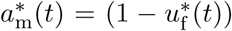. The terminal conditions for these differential equations (S82) are given by the transversality conditions (S72).

Because the number of workers does not appear in the expression of invasion fitness, we have, regardless of the mode of control of traits, that

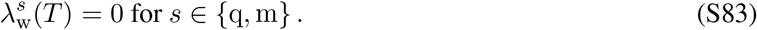

Otherwise, we have from the perturbation formula for eigenvalues (eq. S18 and S72) that

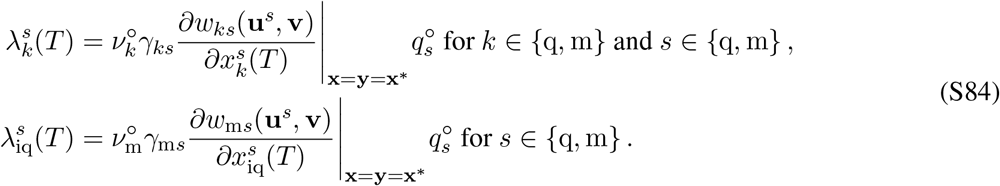

Furthermore 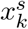 affects only the component *w*_*ks*_(**u**^*s*^, **v**) of invasion fitness for *k* ∈ {q, m} and 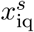 only affects the component *w*_m*s*_(**u**^*s*^, **v**). Hence, owing to eq. (S20) and the fitness functions (eqs. S6–S7), we have

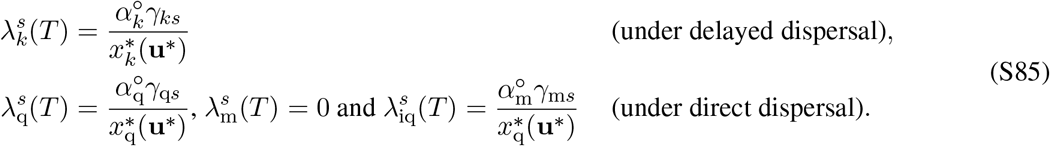

For haplodiploids, *γ*_qq_ = 1/2, *γ*_mq_ = 1, *γ*_qm_ = 1/2 and *γ*_mm_ = 0 and consequently eq. (S21) yields that *α*_q_ = 2/3 and *α*_m_ = 1/3. Substituting these parameters into eq. (S85) yields the following transversality conditions for sexuals under delayed dispersal

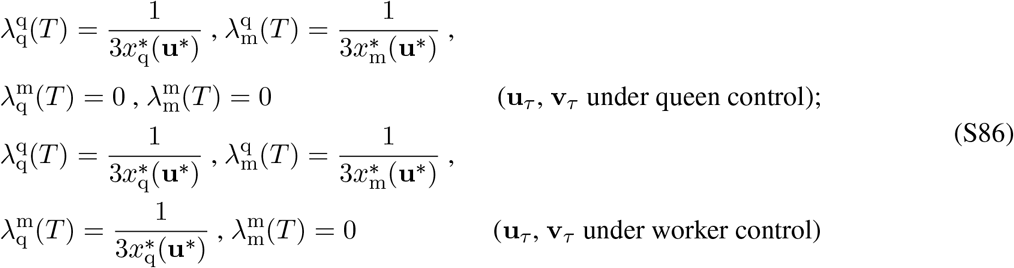

and under direct dispersal the transversality conditions are given by

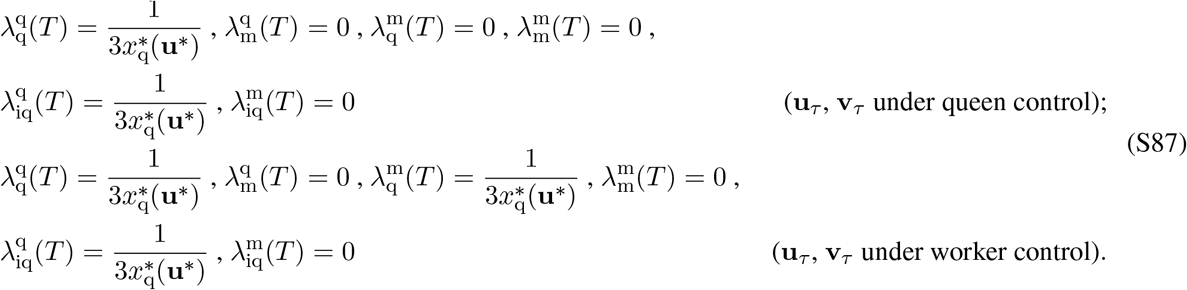

#### 4.3.2 Solutions to the state and costate equations

Let the upper bar denote that the variable is constant. Hence, let 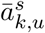 denote constant proportional allocation to individuals of type *k* in a colony founded by a mutant individual of type *s* during a given time interval [*t*_0_, *t*_1_]. The solutions to the differential equations (S57)–(S58) that give the expressions for the state variables at time *t* ∈ [*t*_0_, *t*_1_] are

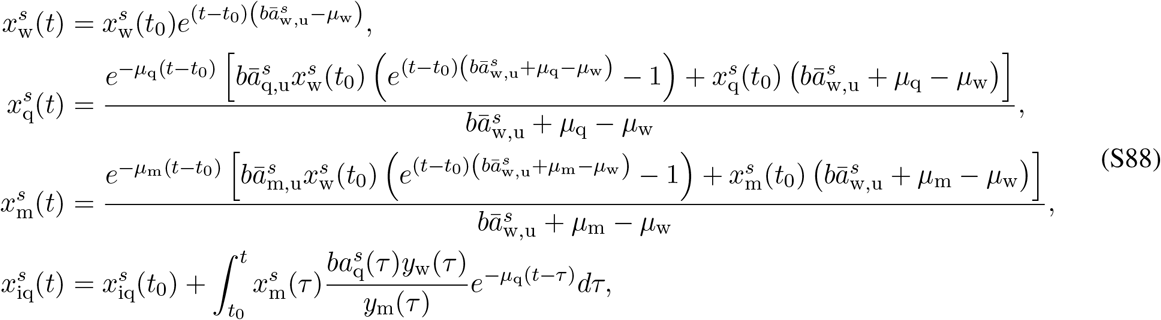

Let *ā*_*k*_ denote the constant proportional allocation to individuals of type *k* in a resident colony during a given time interval [*t*_0_, *t*_1_]. The solutions to differential equations (S59)–(S60) that give the expressions for the dy-namic variables at time *t* ∈ [*t*_0_, *t*_1_] are

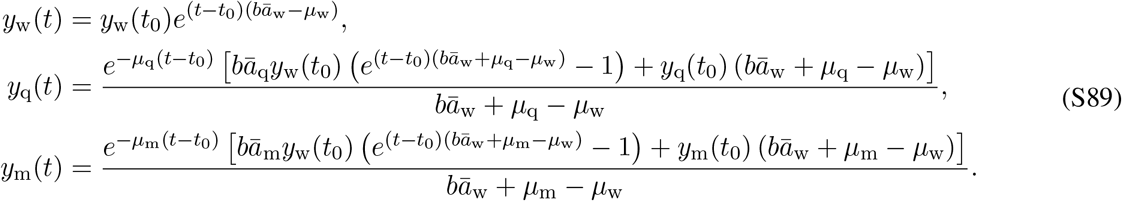

Let 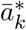 denote a constant uninvadable proportional allocation to individuals of type *k* during a given time interval [*t*_0_, *t*]. The solutions to differential equations eq. (S82) that give the expressions for the costate variables at time *t* ∈ [*t*_0_, *t*_1_] are

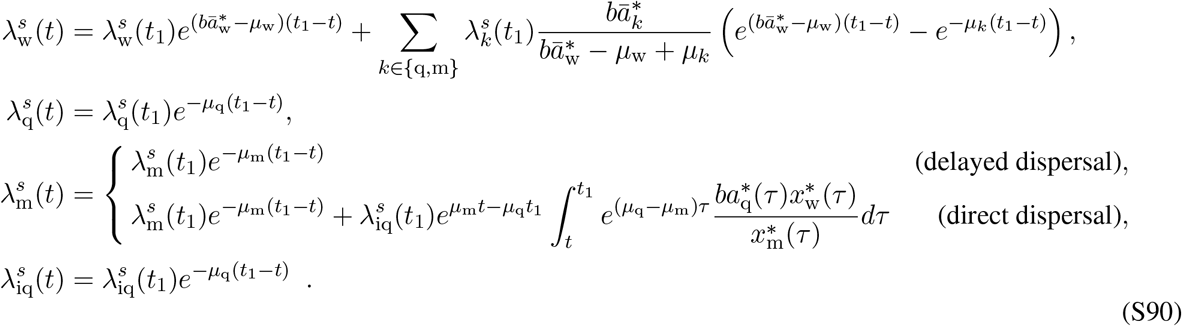

Note that the solutions given by eq. (S90) to costate equations hold for 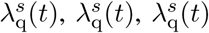 even if the unin-vadable allocation schedule **u**\*(*t*) is not constant during [*t*_0_, *t*]. As opposed to state variables, the dynamics of costate variables is described backwards in time, where 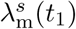 is the terminal condition. The transversality conditions (S86)-(S87) together with eq. (S90) imply that

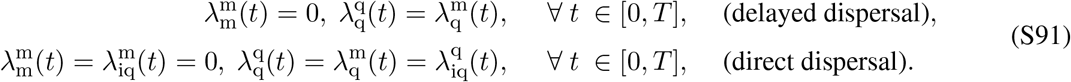

## 5 The candidate uninvadable allocation schedule under delayed dispersal

### 5.1 Equal male and female mortality (*µ*_q_ = *µ*_m_)

Here we show analytically that the candidate uninvadable allocation schedule under delayed dispersal (assuming equal mortality rates of sexuals, i.e. *µ*_q_ = *µ*_m_ = *µ*_r_) consists of only two growth regimes: (i) the ergonomic phase (regime W) and (ii) the reproductive phase, where males and new queens are produced simultaneously (regime FM). The uninvadable allocation schedule can be described by phases W and FM of table S1:

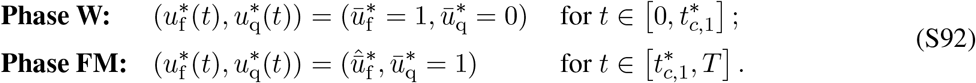

The uninvadable allocation schedule has this form regardless of the assumptions about the genetic control of the resource allocation traits. However, the switching time 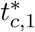 from the ergonomic phase to the reproductive phase depends on the scenario *c* of the genetic control of the resource allocation traits. We will proceed according to the scheme for deriving the uninvadable allocation schedule **u**\* outlined in section 4.2.

#### 5.1.1 Regime FM: 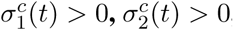, and 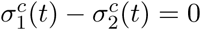

We know from biological considerations that 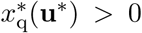 and 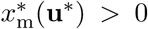 Taking this into account and substituting the transversality conditions (S83) and (S86) into the switching functions (S78) (for queen control) and (S80) (for worker control), implies that 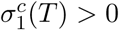 and 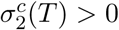. According to table S1, we need to also determine the sign of 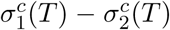. Assuming equal mortality rates of males and queens (*µ*_q_ = *µ*_m_ = *µ*_r_) and substituting eq. (S86) into eqs. (S78) (for queen control) and (S80) (for worker control) yields

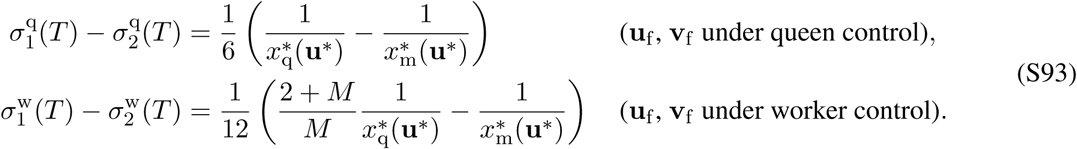

We know from eq. (S35) that the first-order condition implies that 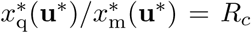, where *R*_*c*_ = 1 for queen control and *R*_*c*_ = (2 + *M*)/*M* for worker control (recall S35), which, on substitution, implies that

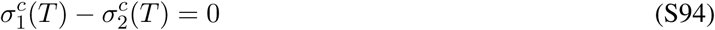

for both queen and worker control of the focal trait of type *τ*.

Hence, we have shown that 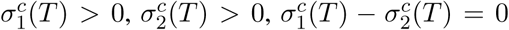. This implies via table S1 that 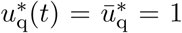 during the final growth regime. It also follows from eq. (S81) and 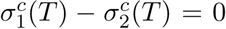, that 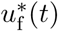 might be a singular arc during the final growth regime. More precisely, for 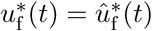 to be a singular arc,

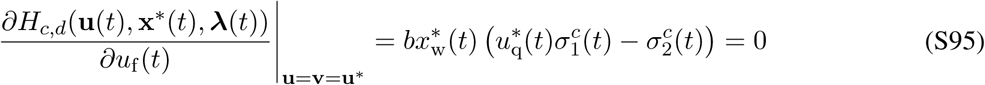

must hold for a finite interval of time (e.g., Bryson and Ho, 1975, p. 248). It follows, that also higher order time derivatives have to vanish along the singular arc and this condition can be used to determine 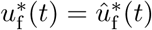, i.e. the constraints for the singular arc are given by the sequence

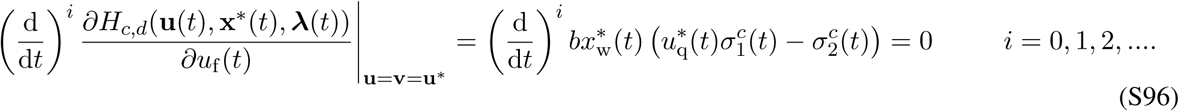

Note that *b >* 0 for biological reasons and 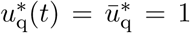 during the last growth regime. It follows from eq. (S88) (by taking 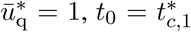, and *t*_1_ = *T*) that 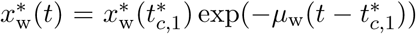) during the last growth regime. Since, 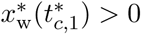, then eq. (S96) simplifies to

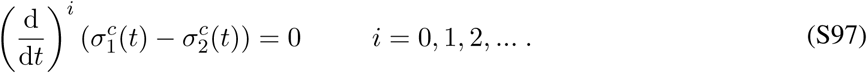

Substituting the switching functions eq. (S78) (for queen control) and (S80) (for worker control) into eq. (S97) implies for *i* = 1 that on a singular arc

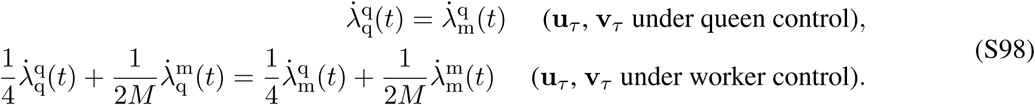

must hold. Simplifying eq. (S98) by using eq. (S91); namely, 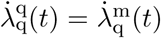 and 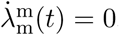, and further using the expression for relatedness asymmetry *R*_*c*_ given by eq. (S35), we obtain

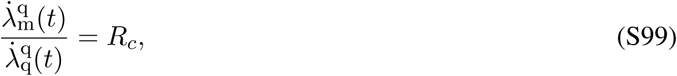

where *R*_*c*_ is the relatedness asymmetry associated with the party *c* ∈ {q, w} in control of the trait of type f. Substituting the differential equations for costate variables (S82) into eq. (S99) gives

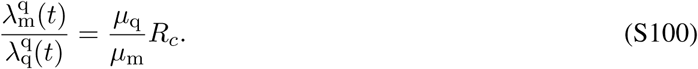

Note that the control variable *u*_f_ (*t*) itself does not appear in eq. (S100). Furthermore, it follows from the simple form of the costate equations (S82) that the *i*-th order time derivative of the coefficient 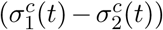 produces

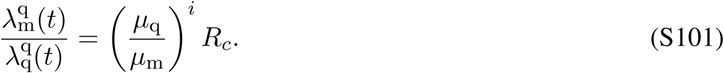

Since the control variable does 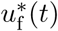 not appear in any order time derivative of the coefficient 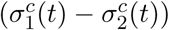, we can use the constraints (S101) that the singular arc has to satisfy to indirectly obtain the expression for the singular arc 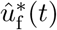. Substituting the costate equations (S90) (and setting *t*_1_ = *T*) into eq. (S101), using the transversality conditions given by eq. (S86), and assuming that the mortality rates of queens and males are equal (*µ*_q_ = *µ*_m_) yields

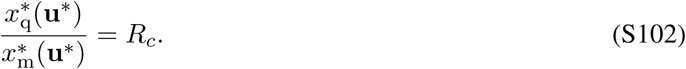

Here, we have recovered the critical sex ratio given by eq. (S50), which essentially implies that if males and queens have equal mortality rates, there are no additional dynamic constraints besides the critical sex ratio at the end of the season. Since, there are no additional dynamic constraints for the control variable, we assume (for simplicity) that it is constant, i.e. 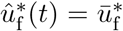 This allows us to substitute the equations for state variables for constant allocation strategy 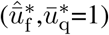, given by eq. (S88) with eq. (3) into eq. (S102) for the time interval 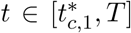 and assuming that *µ*_q_ = *µ*_m_ = *µ*_r_ and 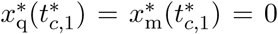 (at the start of the reproductive phase, there are no males or juvenile queens), we obtain

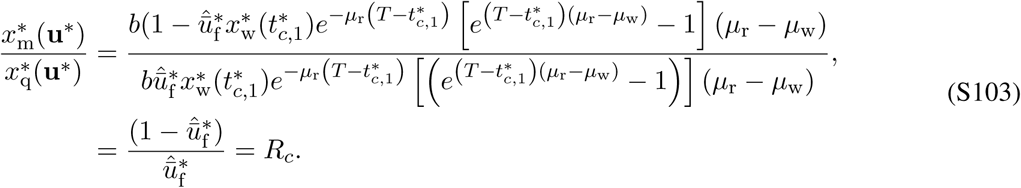

Solving for 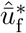 yields

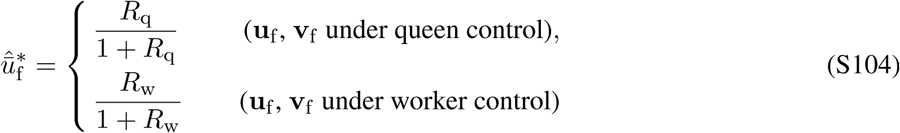

and for haplodiploids it simplifies to

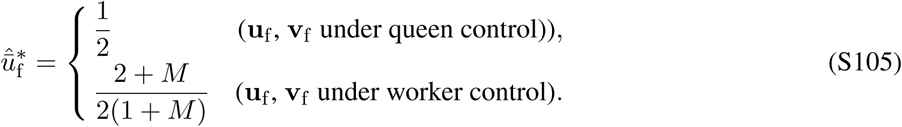

Hence, we have determined the uninvadable allocation schedule **u**\*(*t*) for the final growth regime FM 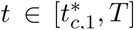, assuming equal mortality rates of males and queens.

#### 5.1.2 Regime W: 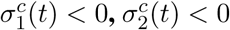, and 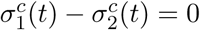

In order to determine the preceding phase, we need to determine which switching function expression 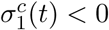, 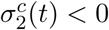, and 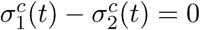 changes their sign.

Lets first examine expression 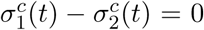. Substituting the costate variables given by eq. (S90) (for the time interval *t* ∈ [0, *T*]) into the expression 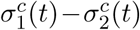 given by eq. (S78) (for queen control) and eq. (S80) (for worker control) yields for any *t* ∈ [0, *T*]

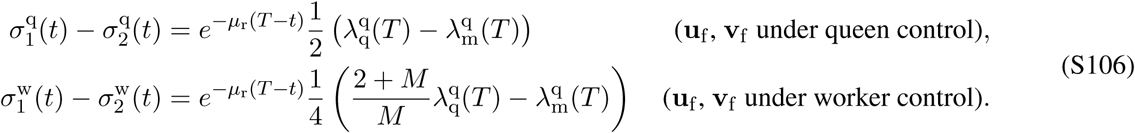

Notice that from the expression 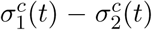 given by eqs. (S78) (for queen control) and (S80) (for worker control) that eq. (S106) can also be expressed in terms of

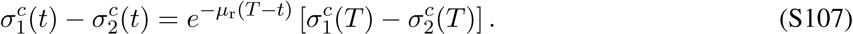

It follows from eq. (S94) that 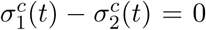 throughout the entire time interval *t* ∈ [0, *T*] for both queen and worker control.

Next, we will show that there is a switch from regime FM to regime W. If there exists at least one root of *t* in equation 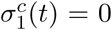, given that the costate variables in 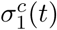 are obtained through integrating eq. (S82) over the final growth regime 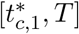, assuming that 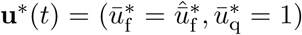, then there is a switch in *u*_q_(*t*) and *u*_f_ (*t*) (since 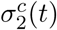 also changes its sign because 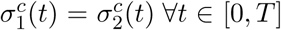. The switching time 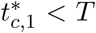 from phase FM to phase W is given by the largest root of 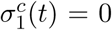 (assuming that 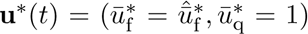 in the last phase) and its existence is shown in sections 5.1.3 for single-party control and 5.1.4 for mixed control, respectively. We can infer from our numerical solutions that *u*_q_ is not a singular arc. Hence, 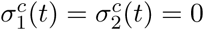 only at time 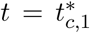 and 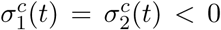 for 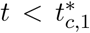. Hence, we have determined that phase W (with allocation schedule 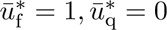 is the penultimate phase.

In order to determine if there are additional switches during time 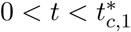, one can further look for roots of the switching function 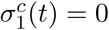 that satisfy 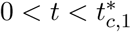. It follows from 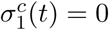 and eq. (S78) for queen control and eq. (S80) for worker control that

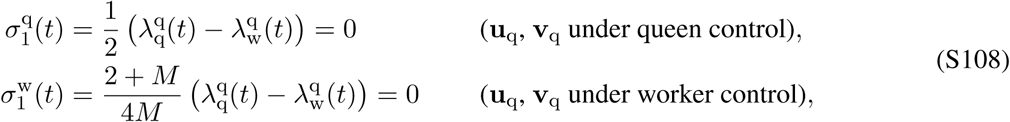

where costate variables at time *t* are evaluated by using eq. (S82) 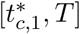, assuming that 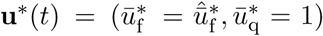 during 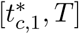 and 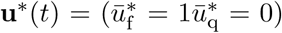 during [*t, T*]. Our numerical solutions indicate that there are no additional growth regimes (see Figs. 1–2). This can be also shown analytically, by showing that there are no additional switches in the switching functions for 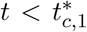 However, for conciseness, we do not provide the proofs here.

In conclusion, we have determined that the optimal allocation schedule consists of two growth regimes: starting with the ergonomic regime W followed by reproductive regime FM, where the switch from regime W to regime FM happens at time 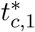 We also verified that our analytical results are in line with our numerical solutions (see Fig. 1, where *µ*_q_ = *µ*_m_ = *µ*_r_).

#### 5.1.3 Switching time for single-party control

We have determined the candidate optimal controls for the ergonomic 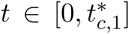 and the reproductive 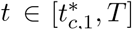 phase (eq. S92 and S104), and we are now going to determine the switching time 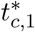 for single-party control (i.e. *c* ∈ {q, w}) that marks the time when the growth schedule switches from one regime to another.

We showed in the previous section that the control variable 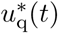 switches its value from 1 to 0, when 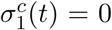. Hence, solving the equation 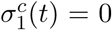 for *t* gives the switching time 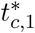 For queen control of **u**_q_ and **v**_q_, eq. (S78) and 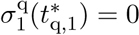 this yields

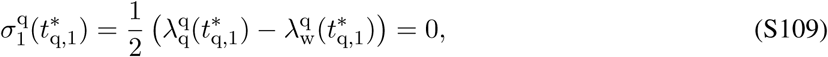

and using the solutions for costate equations (S90), transversality conditions (S83) and (S86), and assuming *µ*_q_ = *µ*_m_ = *µ*_r_ it leads to the following transcendental equation for finding 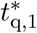

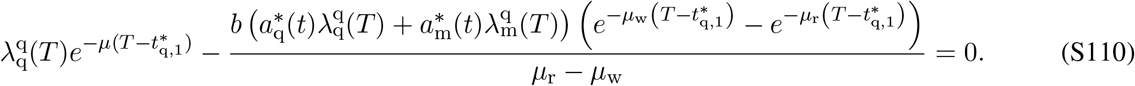

Similarly, for worker control of **u**_q_ and **v**_q_, eq. (S80) and 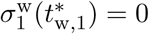 yields

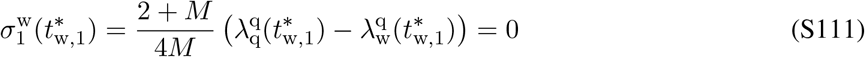

and using the solutions for costate equations (S90), transversality conditions (S83) and (S86), and assuming *µ*_q_ = *µ*_m_ = *µ*_r_, it leads to the following transcendental equation for finding 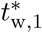

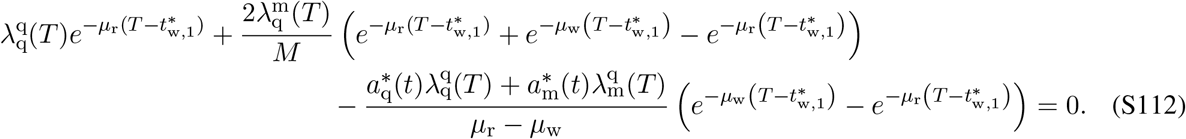

By solving eq. (S110) for 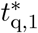 and (S112) for 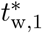 and taking 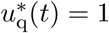, we obtain

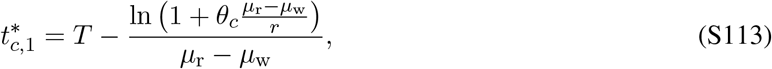

where

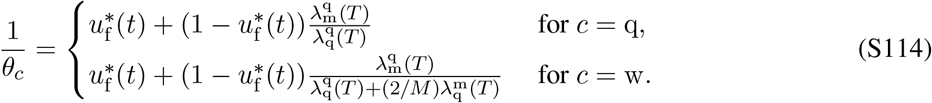

Using eq. (S86) we obtain

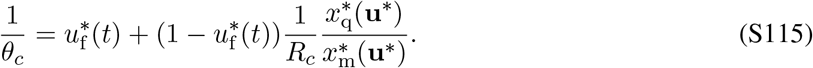

After substituting the control variable 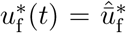 from equation (S104) for the respective case of control and using eq. (S102), we finally have

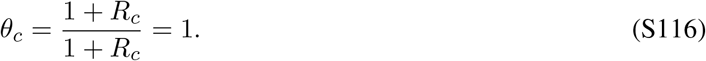

We have obtained that the switching time for single-party control is

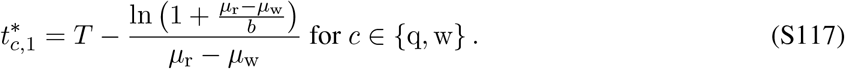

In the limit where the mortality of sexuals becomes equal to the mortality of workers (*µ*_r_ → *µ*_w_) the switching time simplifies to

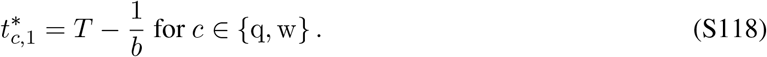

Note that in our model (1/*b*) can be loosely interpreted as the time it takes for one worker to help produce one offspring, i.e. a generation time. Hence, when the mortality rate of sexuals is roughly equal to the mortality rate of workers, then the switching time from the ergonomic to the reproductive phase 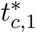 under single-party control (*c* = {q, w}) approaches to one generation time (1/*b*) before the end of the season, i.e. only the last generation of brood is reproductive.

#### 5.1.4 Switching time for mixed control

It follows from eqs. (S65) and (S76) that under mixed control the trait 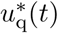 is determined from the sign of 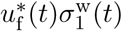 and the trait 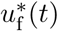 is determined from from the sign of 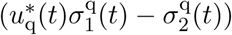. Hence, the switching time 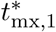 (the switch in the trait 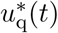 from ergonomic phase to reproductive phase under mixed control can be found by solving 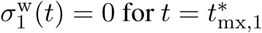, which by way of eq. (S80) yields

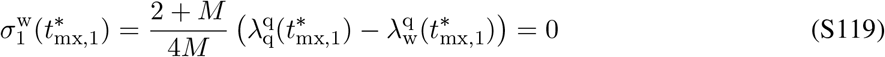

and using the solutions for costate equations (S90), transversality conditions (S83) and (S86), and assuming *µ*_q_ = *µ*_m_ = *µ*_r_, it leads to the following transcendental equation for finding 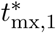

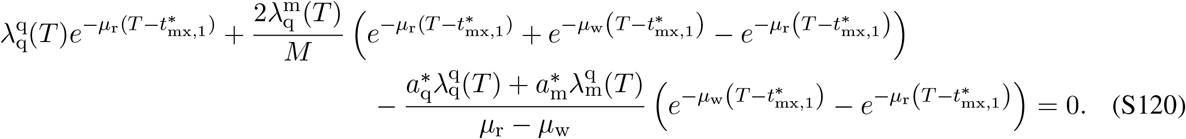

Solving eq. (S120) for 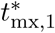 yields

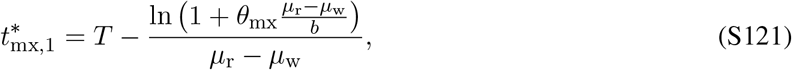

where

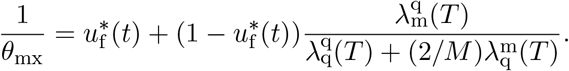

Since according to eq. (S76) for mixed control, the trait 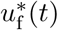 is determined from equation 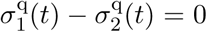. We have shown earlier (see eqs. S98–S104 for queen control) that this leads to an equation for 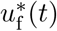 given by eq. (S104) (for queen control). Substituting 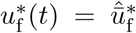 from eq. (S104) (for queen control) and with transversality conditions (S86) and eq. (S102) (for queen control) and using eq. (8) of the main text, we obtain

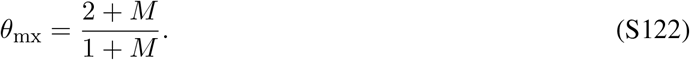

It follows from eq. (S121) that 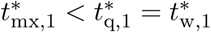 and as *M* → *∞*, 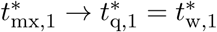 (see eq. S117). We should also mention that equations (S117) and (S121) hold if *µ*_w_ *< b/θ*_*c*_ + *µ*_r_. This not biologically restrictive, since the reproduction rate has to be significantly higher than worker mortality otherwise the population will go extinct. Finally, in the limit where the mortality of sexuals approaches the mortality of workers (*µ*_r_ → *µ*_w_) the switching time simplifies to

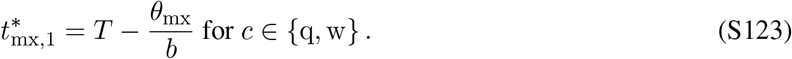

Hence, when the mortality rate of sexuals is roughly equal to the mortality rate of workers, then under mixed control the switch happens *θ*_mx_ generations earlier. For example, when females mate only once (*M* = 1) the switch to reproductive phase happens one and a half generations before the end of the season.

### 5.2 Unequal male and female mortality (*µ*_q_≠ *µ*_m_)

The above analytical results hold for *µ*_q_ = *µ*_m_. Our numerical solutions indicate that if the mortality rates of queens and males are not equal (*µ*_q_≠ *µ*_m_), then the sex that has the lower mortality rate is produced first. Furthermore, the numerical solutions confirm that males and queens are produced such that by the end of the season the ratio of queens to males is given by the relatedness asymmetry, i.e. eq. (S50) holds, regardless of the mortality rates of males and queens. In Figs. S1–S2, we have depicted our numerical results for the uninvadable proportional allocation 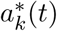 to individuals of different types and the corresponding number of individuals 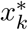, respectively, assuming that queen mortality is lower than male mortality.

**Figure S1:**
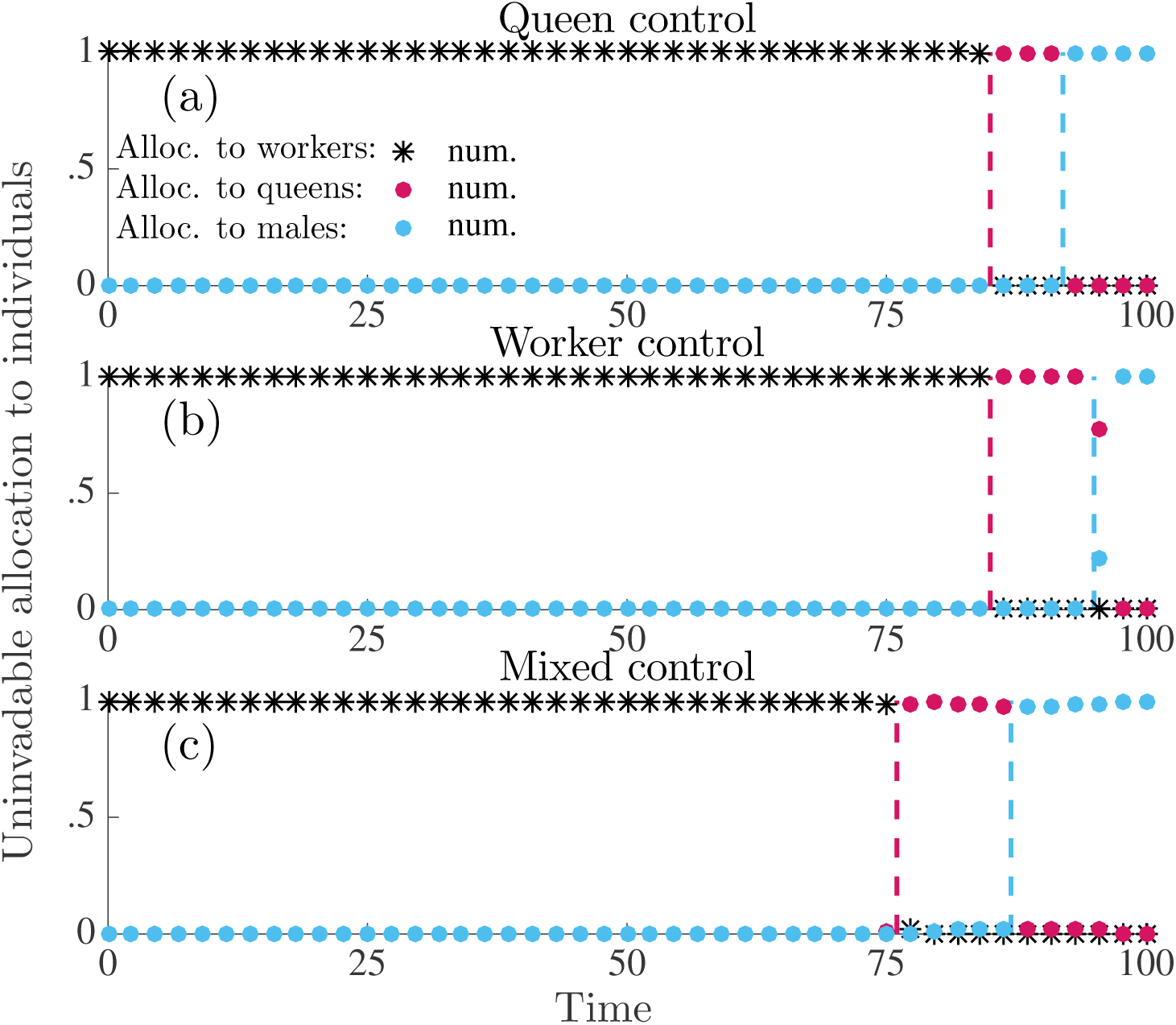
Uninvadable proportional allocation (under delayed dispersal) to workers 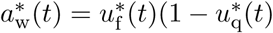 (black asterisks), queens 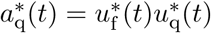 (red circles), and males 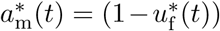 (blue circles). Results here are only numerically derived. Panel (a): queen control. Panel (b): worker control. Panel (c): mixed control. Parameter values: *M* = 1 (queen monandry), *b* = 0.07, *µ*_w_ = 0.015, *µ*_q_ = 0.001, *µ*_m_ = 0.02, *T* = 100.

**Figure S2:**
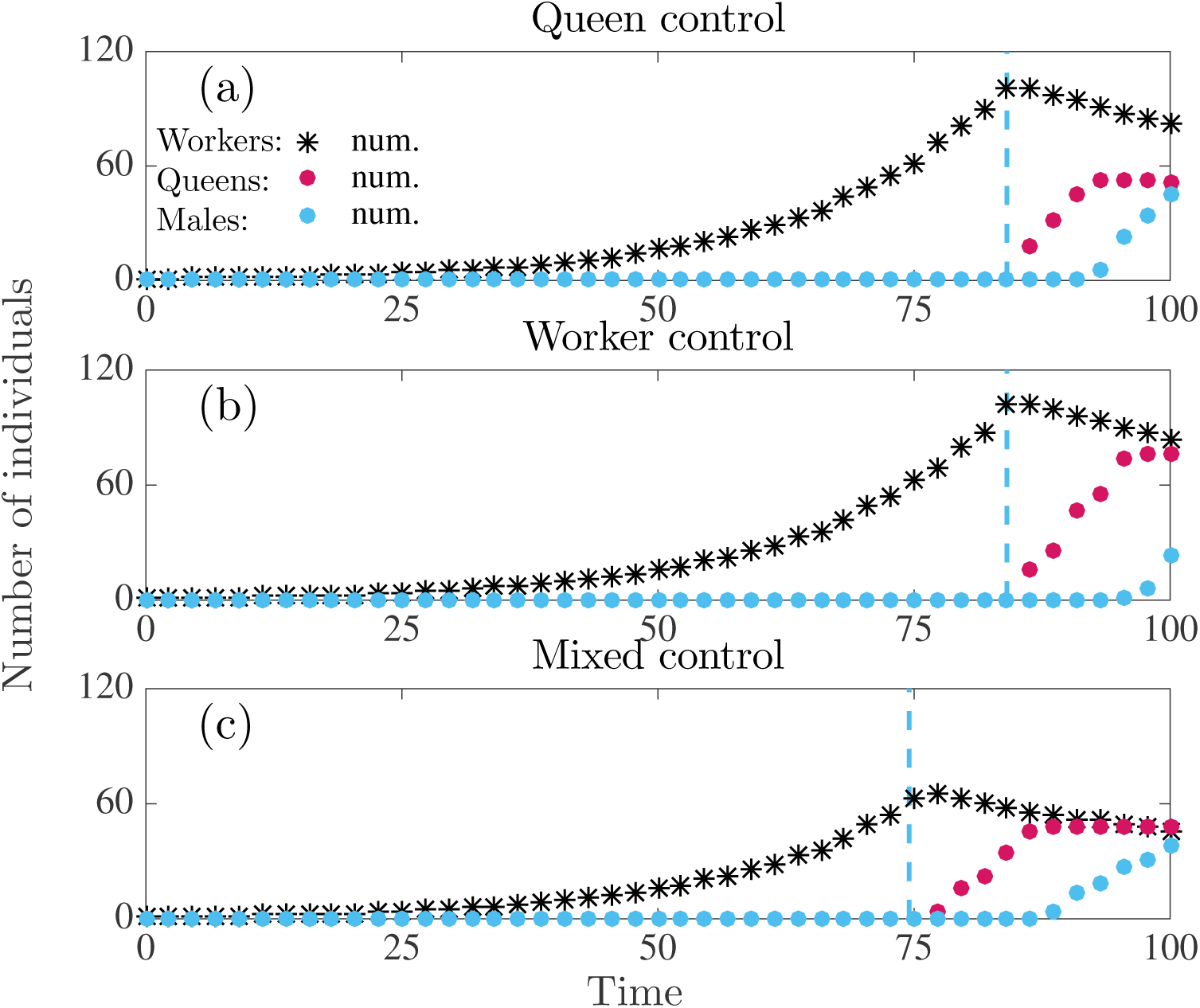
Number of individuals produced in a colony following the uninvadable resource allocation schedule **u**\* under delayed dispersal. Number of workers (black asterisks), number of juvenile queens (red circles), number of males (blue circles). Results here are only numerically derived. Panel (a): full queen control. Panel (b): full worker control. Panel (c): mixed control. Parameter values: *M* = 1 (queen monandry), *b* = 0.07, *µ*_w_ = 0.015, *µ*_q_ = 0.001, *µ*_m_ = 0.02, *T* = 100.

## 6 The candidate uninvadable allocation schedule under direct dispersal

### 6.1 The cases *R*_*c*_*µ*_q_ *≥ µ*_m_ (single-party control) and *R*_q_*µ*_q_ ≥ *µ*_m_ (mixed control)

Here, we determine analytically the candidate uninvadable allocation schedule under direct dispersal assuming *R*_*c*_*µ*_q_ *≥ µ*_m_ (single-party control) and *R*_q_*µ*_q_ *≥ µ*_m_ (mixed control). We find that the optimal allocation schedule consists of the following three growth regimes: (i) ergonomic phase (production of workers), (ii) reproductive phase where only males are produced, (iii) reproductive phase where only new queens are produced. Under these conditions, the uninvadable allocation schedule has the following properties

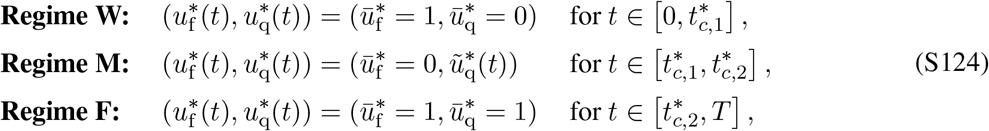

where 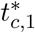 and 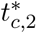 in denote the switching times from ergonomic to reproductive phase and from male production to queen production, respectively, and they depend on the mode of control *c* ∈ {q, w, mx}. We now derive this schedule by working backwards in time.

#### 6.1.1 Regime F: 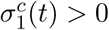 and 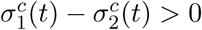

The transversality conditions (S83) and (S87) yield that 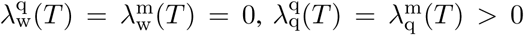, and 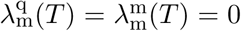. Hence, it follows from eqs. (S78) and (S79) that 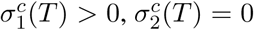, and 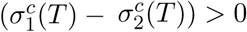. Therefore, from table S1 it follows that 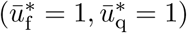 during 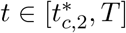, where 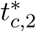 marks the beginning of the last growth regime.

#### 6.1.2 Regime M: 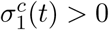 and 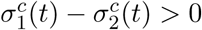

If at least one of the switching functions (S77) changes its sign at time 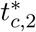, then one of the three alternative conditions must hold: (i) 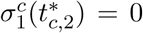, which implies a change in the control variable 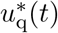, (ii) 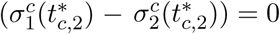, which means that the control variable 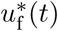 changes, or (iii) 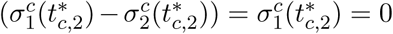, which means that both control variables change. The switching time 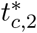 of an uninvadable allocation schedule (S124) is given by the largest root 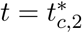 that satisfies one of the conditions given by these scenarios.

Next, we will solve eqs. 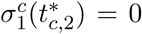 and 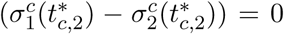 for 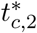, taking into account that that 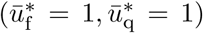 during 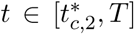 and eq. (S91) that implies that 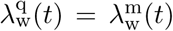. After which we will compare the the roots 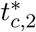 for these two equations in order to determine which of the three alternative above-mentioned scenarios holds.

Firstly, we determine the root 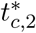 of eq. 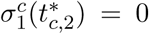. Substituting the transversality conditions (S83) and (S87) into eq. (S90), where we take *t*_1_ = *T* and assume that 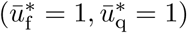 we obtain for 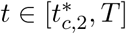

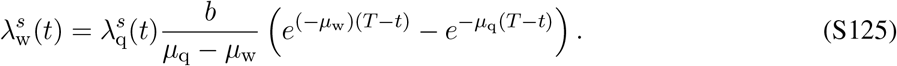

Substituting eq. (S91) into eq. (S125) yields that 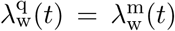. Taking this into account and substituting eqs. (S78) and (S79) into 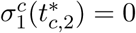 yields

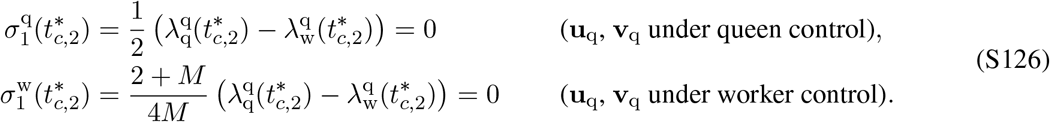

Substituting the solutions to the costate equations (S90) (assuming that *t*_1_ = *T* and 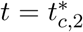), the transversality conditions (S83), (S87) into eq. (S126) assuming that 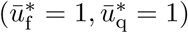 during 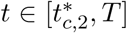 and solving for 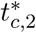 yields

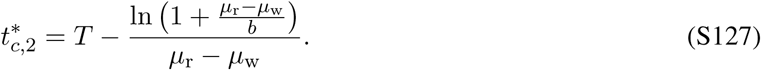

Note that the derivation of eq. (S127) from eq. (S126) is not shown here, since it is very similar to derivation of eq. (S117) from eqs. (S109) and (S111).

Secondly, we need to determine the root 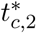 of eq. 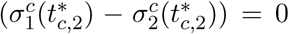. Taking into account that 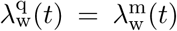 for 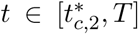 (recall the implications of eqs. S125 and S91) and substituting eqs. (S78) and (S79) (assuming that 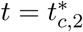) into 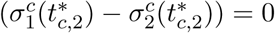 yields for 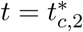 that

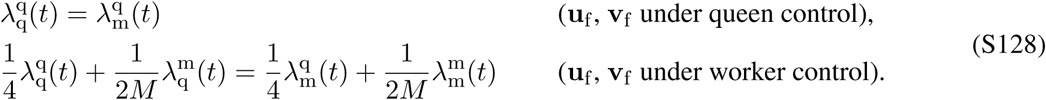

We will show in sections 6.1.4 and 6.1.5 that substituting the solutions to the costate equations (S90) (assuming that *t*_1_ = *T* and 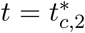) and the transversality conditions (S83), (S87) into eq. (S128) yields a switching time 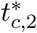 given by eq. (S139) for single-party control (assuming *R*_*c*_*µ*_q_ *≥ µ*_m_) and eqs. (S147) and (S146) for mixed control (assuming *µ*_q_ = *µ*_m_).

Finally, comparing the root 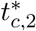 of eq. 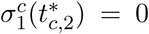 given by eq. (S127) with roots of eq. 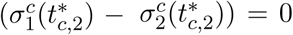, given by eq. (S139) (for single-party control) and eqs. (S147) and (S146) (for mixed control), yields that for biologically realistic parameter values, the root 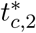 of eq. 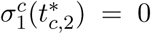 is smaller than the roots of eq. 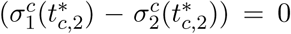. Hence, we have verified that the switch 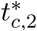 is given by the root of 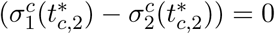 and it follows that the control variable 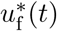 changes its sign at this time.

We have established that 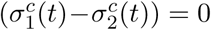 at time 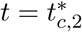 Next we have to determine if 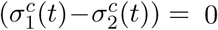 only at time 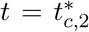 or if 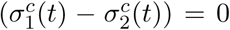 during a finite period of time that ends at time 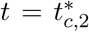. It follows from the definition of the singular arc that if 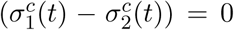 holds for a finite interval of time then 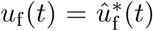 is a singular arc during that time (e.g., Bryson and Ho, 1975, p. 246–249). Furthermore, if 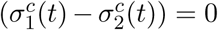 during a finite period of time then it also follows that 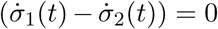 holds during that time. Furthermore, it follows from the time derivative of eq. (S128) that a condition for a singular arc to exist is given by

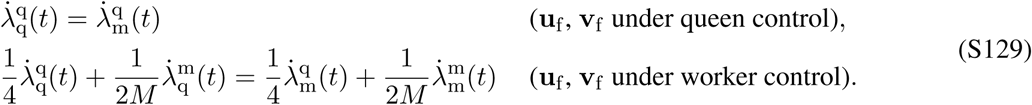

Simplifying eq. (S129) by considering that 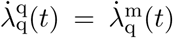 and 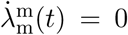 (by way of eq. S91) and using eq. (S35) we obtain

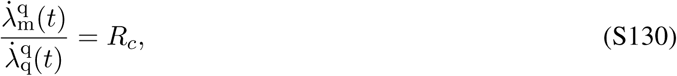

where *c* denotes the party in control of **u**_f_ and **v**_f_. Substituting the costate equations (S82) into eq. (S130) yields

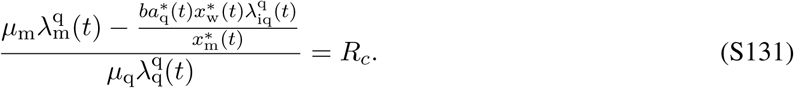

Taking this account together with eq. (S91) and 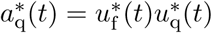 implies that

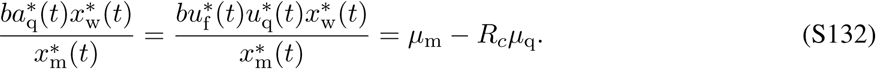

Given that 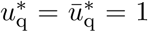 for 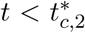 (since 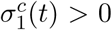 for 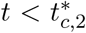) then eq. (S131) implies that 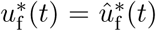 can only be positive if (*µ*_m_ − *R*_*c*_*µ*_q_) *>* 0. Note that here *c* denotes the party in control of **u**_f_ and **v**_f_. Hence, 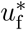 can not be a singular arc before 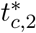 if juvenile male mortality is lower or equal than that of *R*_*c*_ times juvenile queen mortality (i.e. *µ*_m_ ≤ *R*_*c*_*µ*_q_).

Hence we have determined that in the penultimate phase, which ends at time, 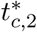 that 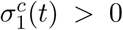 and 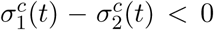 if *R*_*c*_*µ*_q_ *≥ µ*_m_ (under single-party control) or *R*_q_*µ*_q_ *≥ µ*_m_ (under mixed control). This means that if *R*_*c*_*µ*_q_ *≥ µ*_m_ (under single-party control) or *R*_q_*µ*_q_ *≥ µ*_m_ (under mixed control) then regime M (exclusive production of juvenile males) precedes the final regime F, where control variables are given by 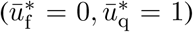. Later in this section we will revisit the case when *R*_*c*_*µ*_q_ *< µ*_m_ (under single-party control) and *R*_q_*µ*_q_ *< µ*_m_ (under mixed control).

#### 6.1.3 Regime W: 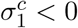 and 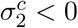

We use the intuition from our numerical solutions (see Fig. 2) that there exists only one additional switching time 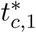, when 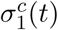 and 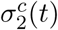 become negative, which represent the first growth regime of the uninvadable allocation schedule. Hence, regime W (worker production) is the first growth regime of the uninvadable alloca-tion schedule, where 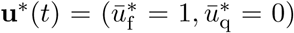. We can determine the switching time 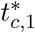 from the condition 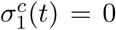, where the costate variables are obtained by integrating them over the last two growth regimes. It turns out that we can explicitly calculate the switching times 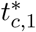 and 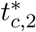 only if *R*_*c*_*µ*_q_ *≥ µ*_m_ under single-party control and only if *µ*_q_ = *µ*_q_ = *µ*_r_ under mixed control.

#### 6.1.4 Switching times under single-party control *R*_*c*_*µ*_q_ *≥ µ*_m_

If *µ*_m_ ≤ *R*_*c*_*µ*_q_ then the condition for the switching time 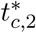 which marks the transition from production of exclusively males to the production of exclusively sexual females is given by (resulting from 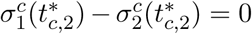)

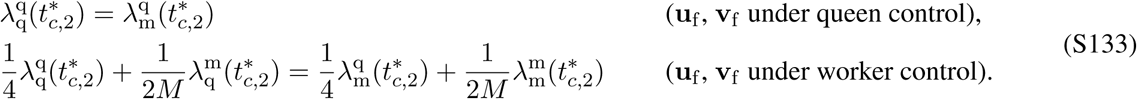

Using eq. (S90) for phase 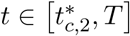, where 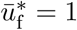 and 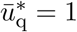 and simplifying, we get

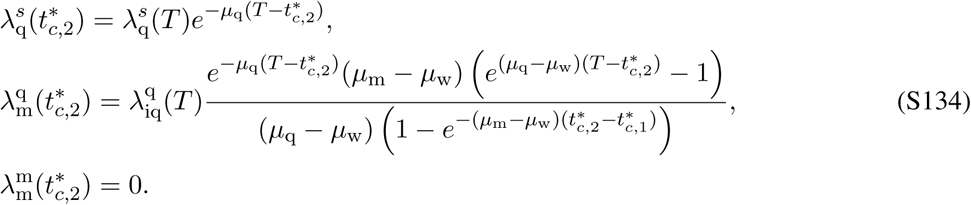

Substituting of eq. (S134) into (S133) and using eq. (S87) implies

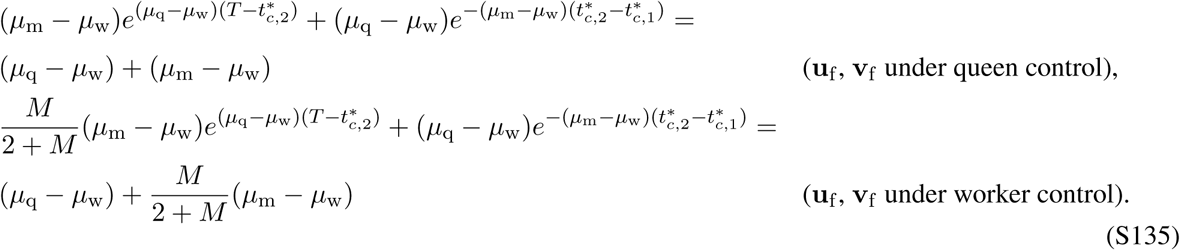

The switching time 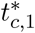 which marks the transition from production of exclusively workers to the production of exclusively males can be found by solving the eq. 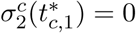 for 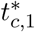 (see table S1)

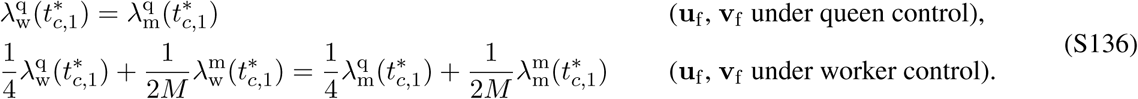

Using eq. (S90) for phase 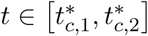, for which 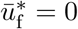 and simplifying, we get

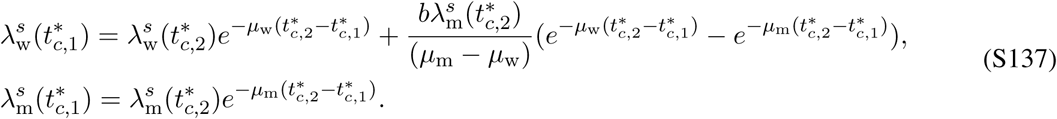

Substituting of eq. (S137) into (S136) and solving for 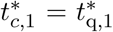 (under queen control) and 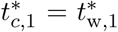 (under worker control) implies

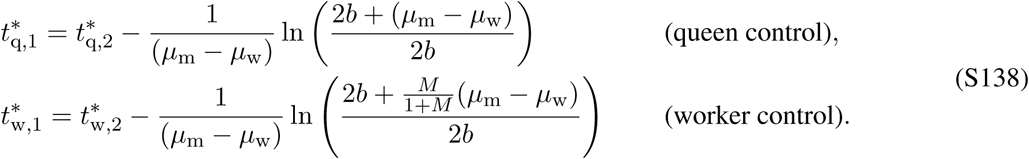

Substituting eq. (S138) into eq. (S135) and solving for 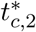 we obtain

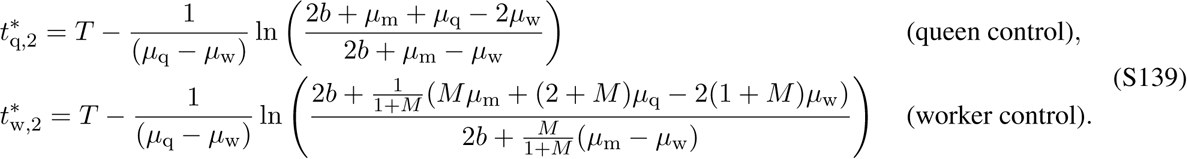

#### 6.1.5 Switching time under mixed control when *µ*_q_ = *µ*_m_ = *µ*_r_

For mixed control, we will derive the switching times 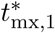 and 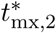 assuming that juvenile queen and male mortality is equal, i.e. *µ*_m_ = *µ*_q_ = *µ*_r_, since this represents the only case where we were able to derive analytical expressions. Under mixed control, the workers control the trait **u**_q_ and the queen controls the trait **u**_f_. Hence, it follows from eqs. (S76) and (S124) that the switching time 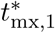 is determined from equation 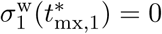 and 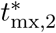 is determined from equation 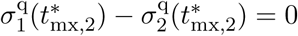.

It follows from eq. (S77) that condition 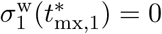 yields

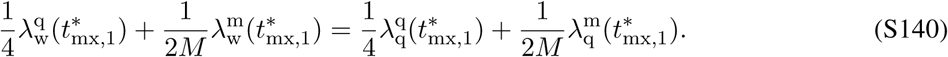

Using eq. (S90) for phase 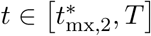, where 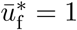 and 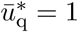 and simplifying, we get

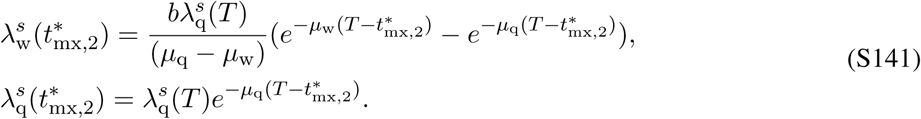

Using eq. (S90) for phase 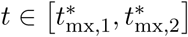, where 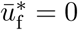 and simplifying, we get

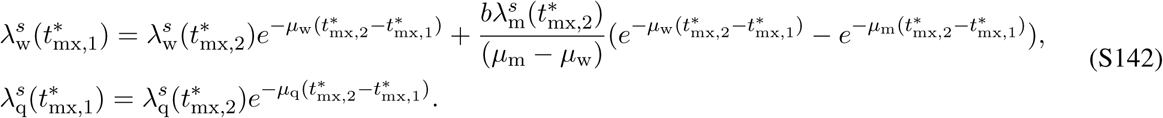

Substituting the the costate variables from eq. (S142) together with (S141) into eq. (S140) and solving for 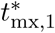 yields

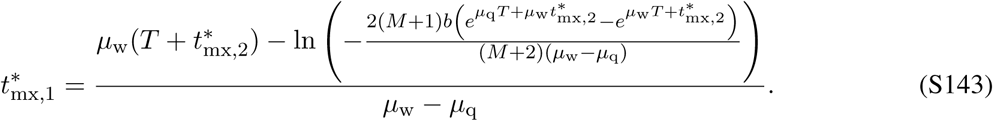

It follows from eq. (S77) that condition 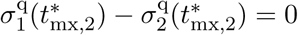 yields

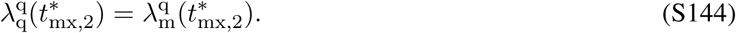

Substituting the costate variables from eq. (S134) and the switching time 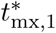 given by eq. (S143) into eq. (S144) and solving for 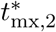 (assuming that *µ*_q_ = *µ*_q_ = *µ*_r_) yields

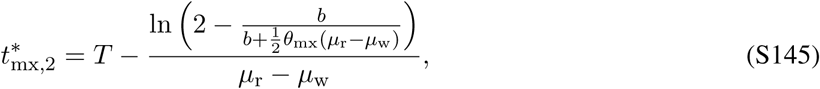

where

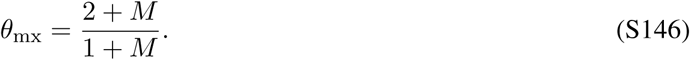

Substituting 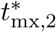 given by eq. (S145) back into eq. (S143) and simplifying yields

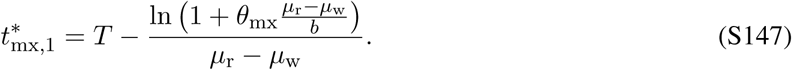

Hence, we have retrieved the same switching time from the ergonomic to the reproductive phase for mixed control under direct dispersal and delayed dispersal (given by eq. S121).

### 6.2 Equal mortality rates of males and queens (*µ*_q_ = *µ*_m_)

In this section, we present the results for the candidate uninvadable allocation schedule under direct dispersal assuming that the mortality rates of queens and males are equal (*µ*_q_ = *µ*_m_ = *µ*_r_). It turns out these results can be directly obtained section 6.1 by equating the mortality rates of queens and males are equal (*µ*_q_ = *µ*_m_ = *µ*_r_). This is because the results in section 6.1 were derived assuming that *R*_*c*_*µ*_q_ *≥ µ*_m_ (under single-party control) and *R*_q_*µ*_q_ *≥ µ*_m_ (under mixed control), where relatedness asymmetry *R*_*c*_ ≥ 1 (recall eq. (S35)).

The optimal allocation schedule consists of the following three growth regimes: (i) ergonomic phase (production of workers), (ii) reproductive phase where only males are produced, (iii) reproductive phase where only new queens are produced. The uninvadable allocation schedule has the following properties

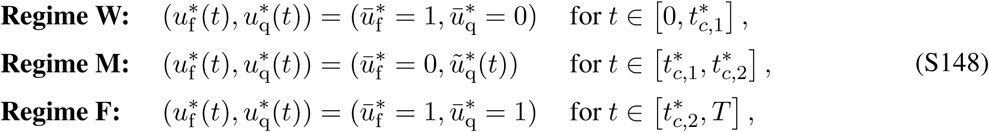

where 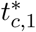 and 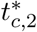 in denote the switching times from ergonomic to reproductive phase and from male production to queen production, respectively, and they depend on the mode of control *c* ∈ {q, w, mx}.

#### Single-party control

If the mortality of juvenile queens and males is equal (i.e. *µ*_q_ = *µ*_m_ = *µ*_r_) then the switching time 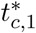 simplifies to

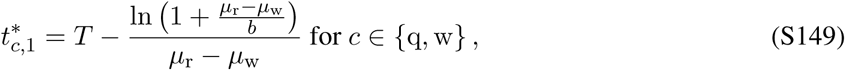

which is equal to the switching time obtained for single-party control under delayed dispersal (eq. S117). For equal juvenile queen and male mortality (i.e. *µ*_q_ = *µ*_m_ = *µ*_r_) the switching time 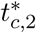 simplifies to

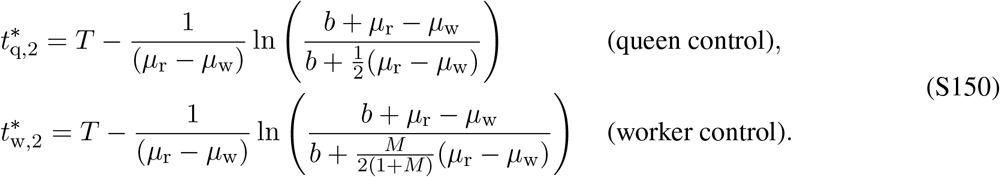

In the limit where the mortality of sexuals becomes equal to the mortality of workers (*µ*_r_ → *µ*_w_) the switching times 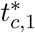 and 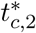 simplify to

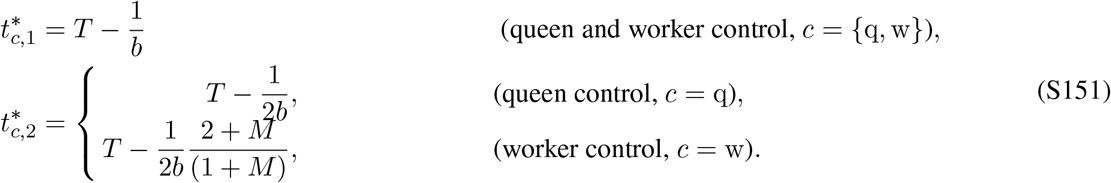

Hence, when the mortality rate of sexuals is roughly equal to the mortality rate of workers, then the switching time from the ergonomic to the reproductive phase 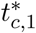 under single-party control (*c* = {q, w}) approaches to one generation time (1/*b*) before the end of the season, i.e. only the last generation of brood is reproductive.

#### Mixed control

We showed previously that if the mortality rates of queens and males are equal (i.e. *µ*_q_ = *µ*_m_ = *µ*_r_) then the switching time 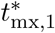 from the ergonomic to the reproductive phase can be expressed as

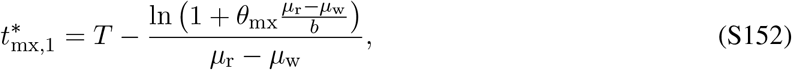

where

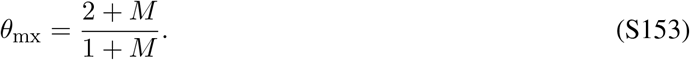

and the he switching time 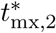 from the male production to the queen production can be expressed as

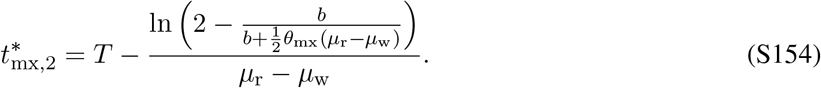

In the limit where the mortality of sexuals becomes equal to the mortality of workers (*µ*_r_ → *µ*_w_) the switching times 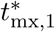 and 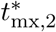 simplify to

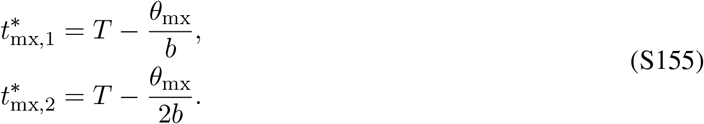

Hence, when the mortality rate of sexuals is roughly equal to the mortality rate of workers, under mixed control the switch happens *θ*_mx_ generations earlier. For example, when females mate only once (*M* = 1) the switch to reproductive phase happens one and a half generations before the end of the season.

### 6.3 The cases *R*_*c*_*µ*_q_ *< µ*_m_ (single-party control) and *R*_q_*µ*_q_ *< µ*_m_ (mixed control)

It follows from eq. (S132) that if *R*_*c*_*µ*_q_ *< µ*_m_ (single-party control) or *R*_q_*µ*_q_ *< µ*_m_ (mixed control), then 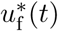 can possibly be a singular arc during some period before 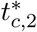, where *R*_*c*_ is the relatedness asymmetry associated with party *c* in control of the trait of type f. Lets denote this singular arc by 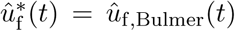, since it was originally derived under full queen control by Bulmer (1983). Furthermore, if the singular arc 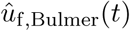 exists, it has to satisfy

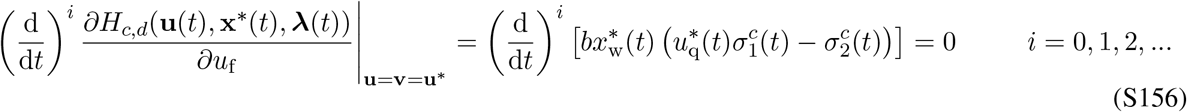

(e.g., Bryson and Ho, 1975, p. 248). And since we have shown previously that 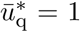 during the penultimate phase and hence 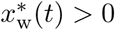 it follows that 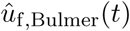 has to satisfy

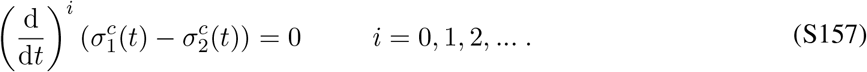

We have already shown that 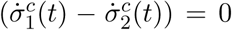 leads to to eq. (S132). Furthermore, 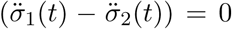 together with eqs. (S78), (S79), (S91) and (S35) implies that

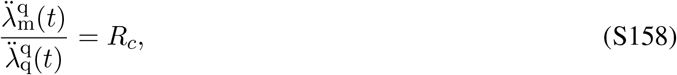

Considering that 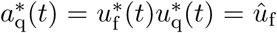, Bulmer 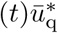 and 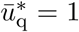 during the penultimate phase and substituting the costate equations (S82) into (S158) yields

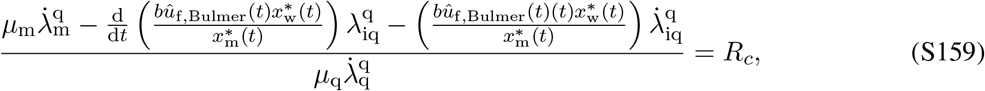

Substituting eq. (S132) and considering eqs. (S90) and (S91) yields

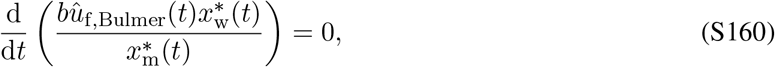

It follows from eq. (S160) that 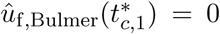, since 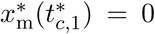. Using the quotient rule of taking derivatives yields

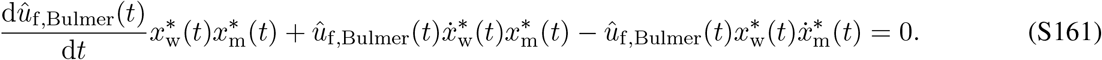

Substituting eqs. (S57) and (S58) into (S161) implies the following differential equation

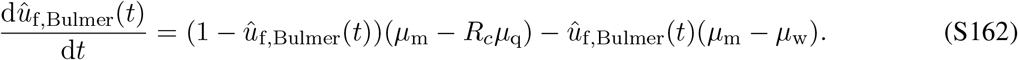

Solving the differential equation for 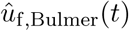 with initial condition 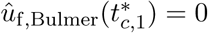 gives

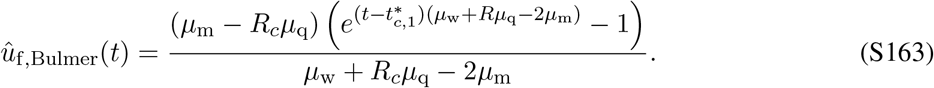

Thus far we have derived the singular arc from the first and second time derivative of the coefficient 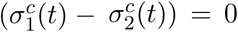. However, it follows from eq. (S131) that the control variable *û*_f,Bulmer_ first appears in the odd member (*i* is odd) in the sequence given by eq. (S157) (i.e. the degree of singularity of the singular arc is odd). It has been proven that if the degree of singularity of the singular arc is odd then it is necessarily non-optimal (Robbins, 1967). This means that if the control variable first appears in the time derivative of the coefficient 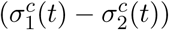 to an odd order, then this singular arc is non-optimal.

Hence, we will only rely on numerical solutions in order to approximate the uninvadable allocation schedule 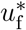 if *R*_*c*_*µ*_q_ *< µ*_m_. Our numerical solutions indicate that under single-party control 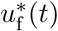 is close to 0 during the penultimate phase 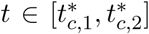 if *R*_*c*_*µ*_q_ *< µ*_m_. In Fig. S3 we demonstrate for single-party control that even if the mortality of queens is 20 times lower than that of males, approximately only males are produced in the penultimate phase 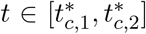. Hence, we find that under single-party control, for a large set of biologically realistic parameter values, approximately only males are produced in the penultimate phase.

We also observe from Fig. S3 that under mixed control 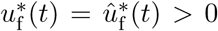 during the penultimate phase 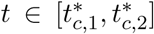 Hence under mixed control, we predict that males and queens are produced simultaneously under mixed control during the penultimate phase 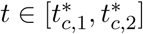 if *R*_*c*_*µ*_q_ *< µ*_m_.

We find that even if the mortality rate of queens is significantly lower than that of males, the overall sex allocation ratio *S*_*c*,dir_ under single-party control is only slightly more female-biased than the uninvadable sex allocation ratio predicted from the standard static models of sex allocation theory (Boomsma and Grafen, 1991; Reuter and Keller, 2001; Trivers and Hare, 1976). We find that the overall sex allocation ratio *S*_mx,dir_ under mixed control is close to the overall sex allocation ratio *S*_q,dir_ under full queen control (see Figs. S3–S4).

**Figure S3:**
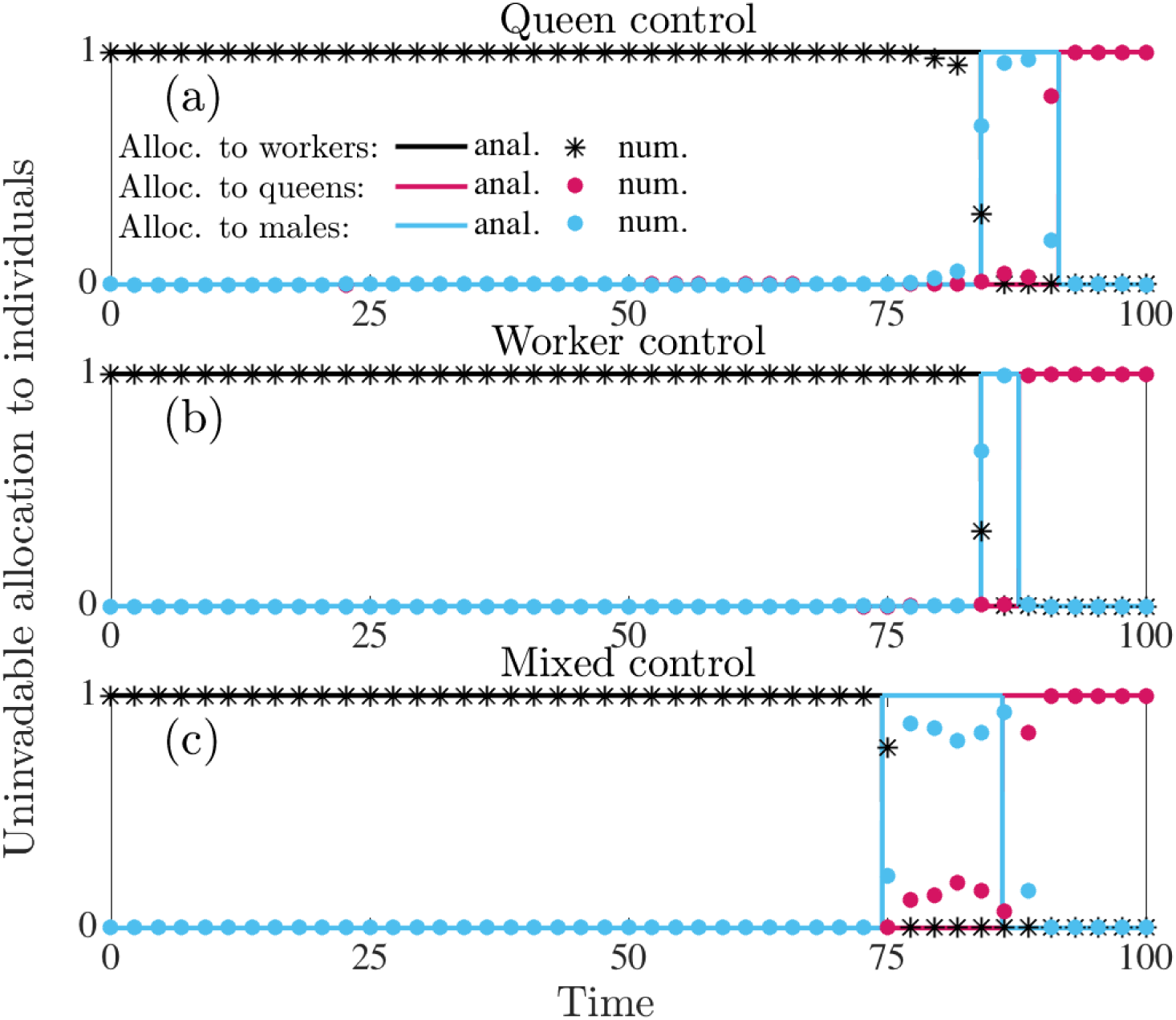
Uninvadable proportional allocation (under direct dispersal) to workers 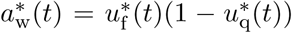 (black), queens 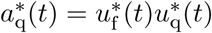 (red), and males 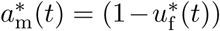 (blue). Panel (a): queen control. Panel (b): worker control. Panel (c): mixed control. Parameter values: *M* = 1 (queen monandry), *b* = 0.07, *µ*_w_ =0.015, *µ*_q_ = 0.001, *µ*_m_ = 0.02, *T* = 100. Results here are only numerically derived and the correspondingly colored lines are analytically predicted results assuming that *µ*_q_ = *µ*_m_ = 0.001. Notice that these analytical predictions approximate the numerically derived predictions quite well under single party control.

**Figure S4:**
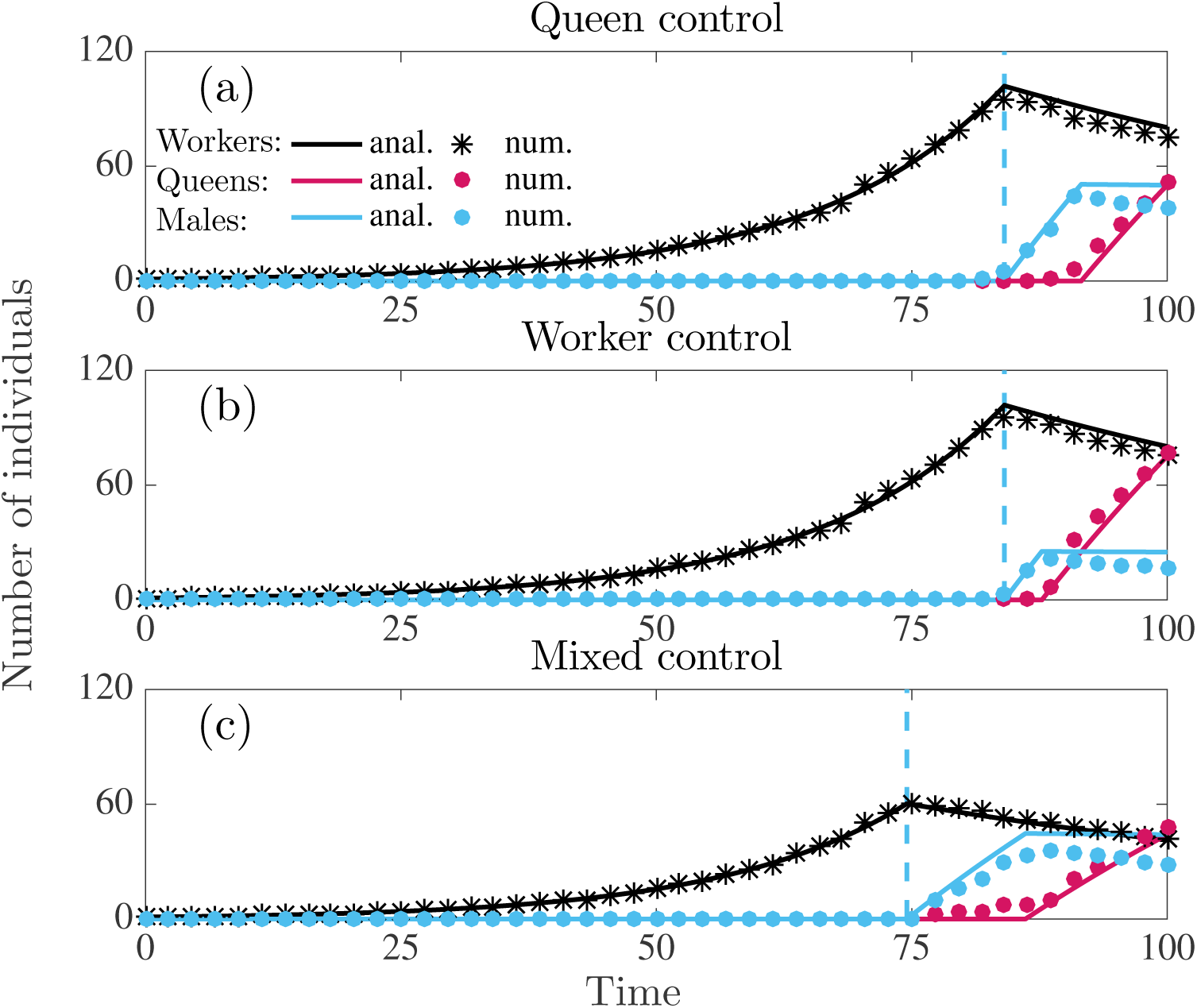
Number of individuals produced in a colony following the uninvadable resource allocation schedule **u**\* under direct dispersal. Panel (a): queen control. Panel (b): worker control. Panel (c): mixed control. Parameter values: *M* = 1 (queen monandry), *b* = 0.07, *µ*_w_ = 0.015, *µ*_q_ = 0.001, *µ*_m_ = 0.02, *T* = 100. The (numerical) overall sex allocation ratio *S*_q_ *≈* 0.51, *S*_w_ *≈* 0.78, *S*_mx_ *≈* 0.53. The correspondingly colored lines are analytically predicted results assuming that *µ*_q_ = *µ*_m_ = 0.001. Notice that these analytical predictions approximate the numerically derived predictions quite well under single party control.

## 7 Macroscopic quantities describing resource allocation in colonies

### 7.1 Colony size at maturity

It follows from eq. (S58) assuming that the allocation schedule to individuals corresponds to the uninvadable allocation schedule **u**\*, given by eq. (S92) (for delayed dispersal) and eq. (S124) (for delayed dispersal) that during the ergonomic phase the number of workers grows exponentially at rate *b-µ*_w_. Furthermore, the number of workers 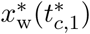 at the switching time 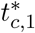 from the ergonomic phase to the reproductive phase determines the colony size at maturity, which is given by

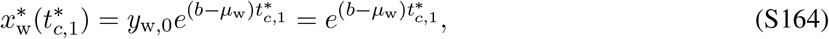

and owing to mortality of workers it is also the maximal colony size.

### 7.2 Colony productivity

The switching time 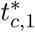 also determines the colony productivity, which we define as the total number of males and females produced that have survived until the end of the season

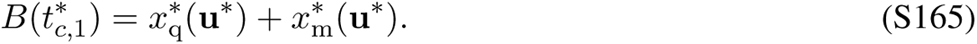

Substituting eq. (S58) for state variables into eq. (S165) assuming that the allocation schedule to individuals corresponds to the uninvadable allocation schedule **u**\* under delayed dispersal (given by eq. S92) and the mortality rate of queens and males is equal (*µ*_q_ = *µ*_m_ = *µ*_r_)

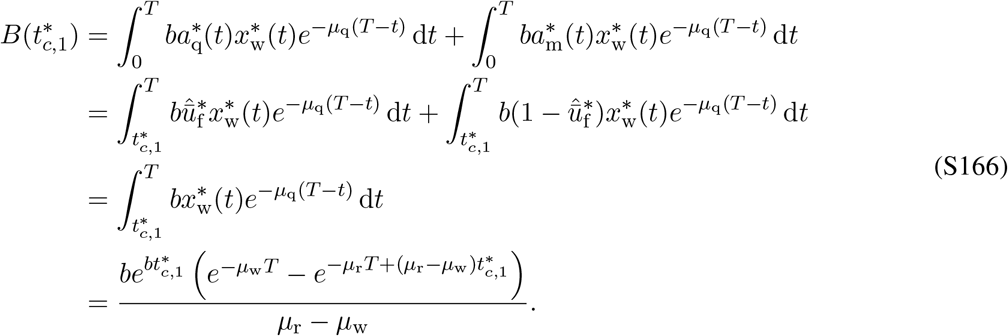

Substituting eq. (S58) for state variables into eq. (S165) assuming that the allocation schedule to individuals corresponds to the uninvadable allocation schedule **u**\* under direct dispersal (given by eq. S124) and the mortality rate of queens and males is equal (*µ*_q_ = *µ*_m_ = *µ*_r_)

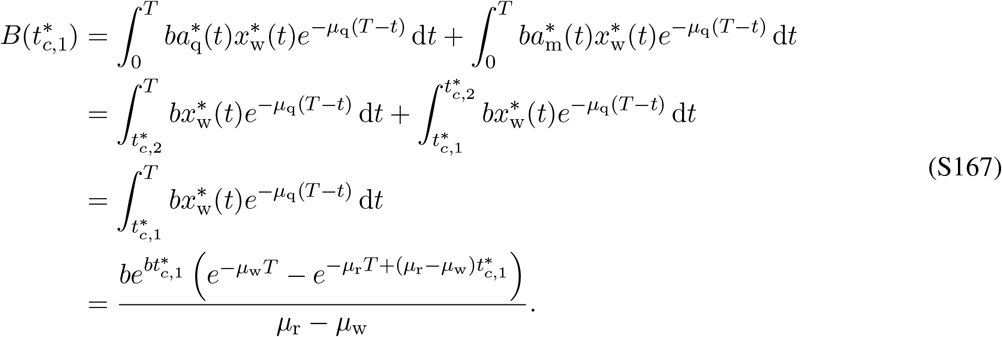

Hence, it follows from eqs. (S166) and (S167) that colony productivity (for delayed and direct dispersal) can be expressed as

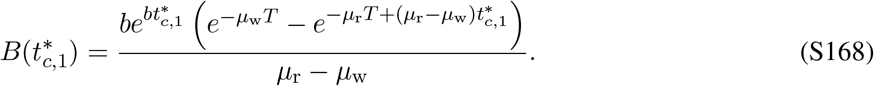

We can determine the switching time 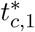 that maximizes colony productivity from

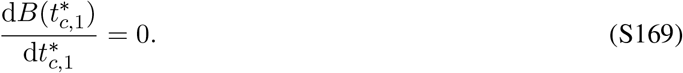

Substituting eq. (S168) into eq. (S169) implies

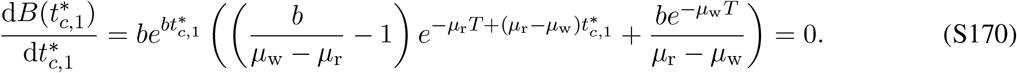

Solving eq. (S170) for 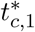 yields

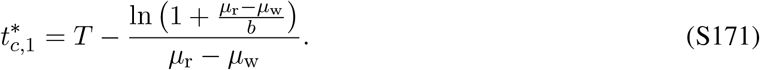

The switching time given by eq. (S171) that maximizes the colony productivity is equal to the switching time 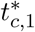 under single-party control (*c* ∈ {q, w}) for both delayed (given by eq. S117) and direct dispersal (given by eq. S149) assuming that the mortality rates of queens and males are equal (*µ*_q_ = *µ*_m_ = *µ*_r_).

### 7.3 Overall sex allocation ratio

We define the overall sex allocation ratio as the proportion of the colony resources allocated to queens from the resources allocated to sexuals over the entire season (irrespective of whether they survive to reproduce), and it is thus given by

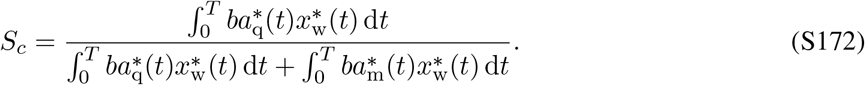

#### Sex allocation ratio under delayed dispersal

Substituting the uninvadable allocation schedule **u**\* for delayed dispersal given by eq. (S92) with the solutions to the state equations given by (S89) into eq. (S172) yields

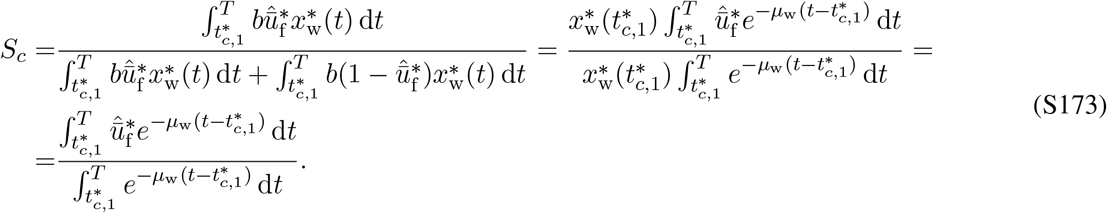

If males and queens are equally costly to produce, then 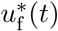 is constant in the reproductive phase and is given by eq. (S104). Hence, eq. (S173) simplifies to

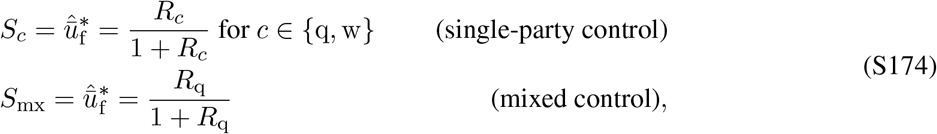

where *R*_*c*_ is the relatedness asymmetry given by eq. (S35) and for haplodiploids (S174) simplifies to

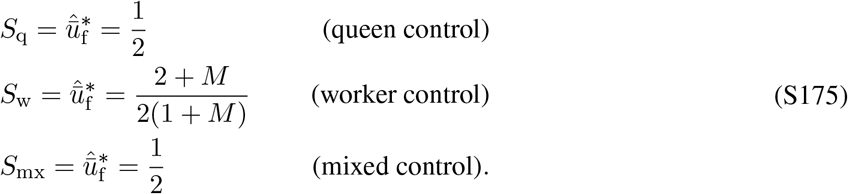

#### Sex allocation ratio under direct dispersal

Recall that the overall sex allocation ratio (the proportion of the colony resources allocated to queens from the resources allocated to sexuals over the entire season) is

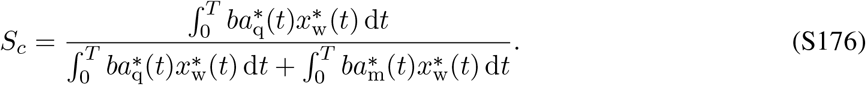

Substituting the uninvadable resource allocation schedule **u**\* for *µ*_m_ ≤ *µ*_q_ under direct dispersal given by eq. (S124) with the solutions to state equations given by (S89) into eq. (S176) yields

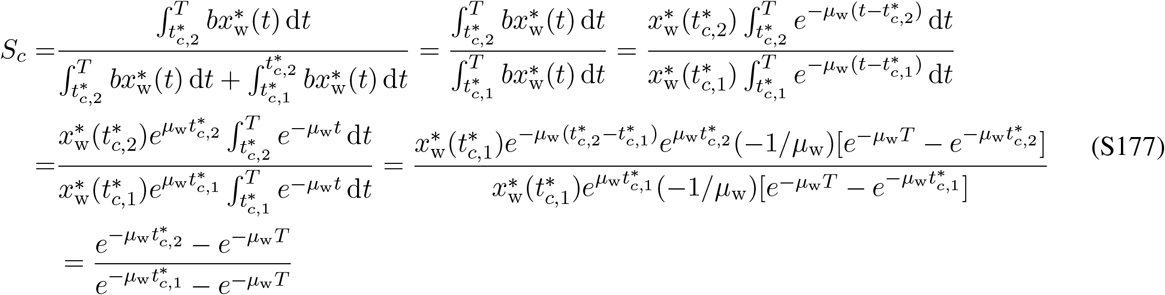

Hence, if *R*_*c*_*µ*_m_ > *µ*_q_ under single-party control and *µ*_m_ > *µ*_q_ under mixed control, then the overall sex allocation ratio under direct dispersal is

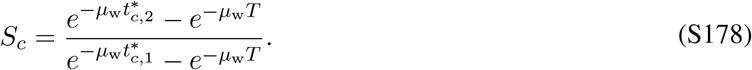

## 8 Marginal return of changing the allocation trait for the ergonomic and reproductive phase under mixed control

The aim of this section is to show that under mixed control the queen determines the overall sex allocation ratio and workers determine the switching time 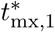 from the ergonomic to the reproductive phase (assuming equal mortality of males and queens, i.e *µ*_q_ = *µ*_m_ = *µ*_r_). We do this by analyzing the marginal return *∂H*_*c,d*_(**u**(*t*), **x**\*(*t*), ***λ***(*t*))/*∂u*_*τ*_ (*t*) = *∂H*_*c,d*_(*t*)/*∂u*_*τ*_ (*t*) of changing the allocation trait during the ergonomic 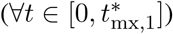 and the reproductive phase 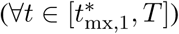 under mixed control.

It follows from the first-order condition for uninvadability under mixed control (recall eq. S65 and eq. S73) for both delayed and direct dispersal (*d* ∈ {del, dir}) that

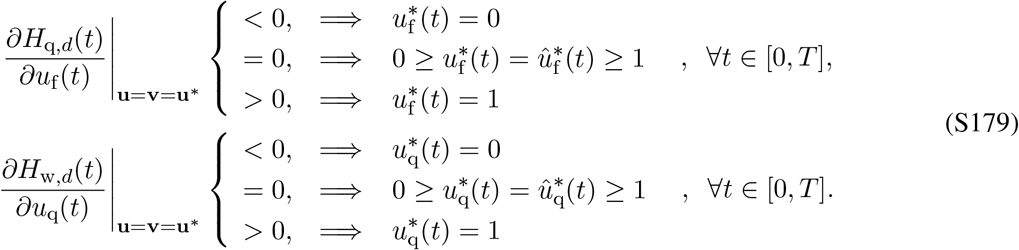

Hence, under mixed control the sign of *∂H*_q,*d*_(*t*)/*∂u*_f_ (*t*), which is under queen control, determines 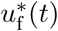, while the sign of *∂H*_w,*d*_(*t*)/*∂u*_q_(*t*), which is under worker control, determines 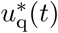.

Let sgn(·) denote a sign function, i.e.

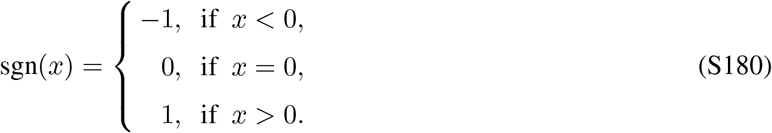

The signs of *∂H*_*c,d*_(*t*)/*∂u*_*τ*_ (*t*) (assuming equal mortality of males and queens, i.e *µ*_q_ = *µ*_m_ = *µ*_r_) can be inferred by way of eq. (S179) from the uninvadable allocation schedule **u**\* given by eq. (S92) for delayed dispersal and eq. (S148) for direct dispersal. Further, using eq. (S76), the fact that *b* > 0 and 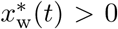, and the switching functions eq. (S77) (where 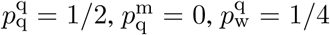, and 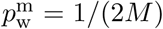), then we have for both delayed and direct dispersal (*d* ∈ {del, dir}) that during the ergonomic phase 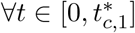:

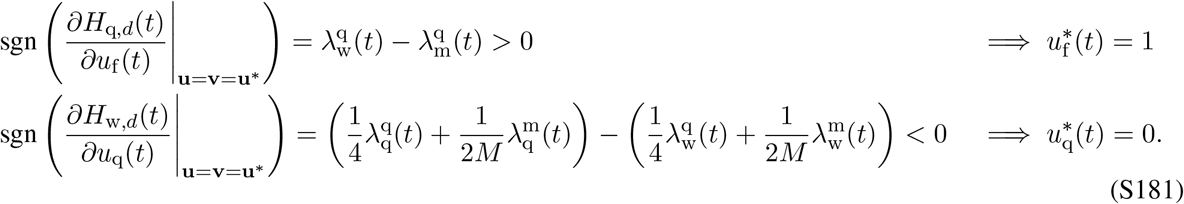

eq. (S181) shows that during the ergonomic phase the marginal return of workers is higher than that of queens and males. More precisely, the sign of *∂H*_q,*d*_(*t*)/*∂u*_f_ (*t*) implies that during the ergonomic phase only females are produced 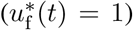, since the marginal return of producing workers 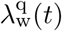 is higher than that of males 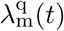 in colonies founded by queens carrying the mutant allele. In other words, during the ergonomic phase workers are more valuable than males to the genes residing in queens. Similarly, it follows from the the sign of *∂H*_w,*d*_(*t*)/*∂u*_q_(*t*) that all females produced during the ergonomic phase become workers 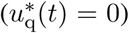, because the marginal return of queens 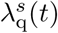 is lower that that of workers 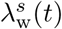, where the marginal returns have been weighed by the expected frequency of mutant alleles in workers in colonies founded by type *s* mutant individuals. In other words, workers are more valuable than queens to the genes residing in the workers. Hence, during the ergonomic phase, there is a latent trade-off between producing workers versus males from the perspective of the genes in the queens and a latent trade-off between producing workers versus queens from the perspective of the genes in the workers. Only workers are produced during the ergonomic phase, since workers have a higher marginal return for both parties.

During the reproductive phase 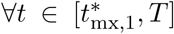, the signs of the marginal returns *∂H*_*c*,del_(*t*)/*∂u*_*τ*_ (*t*) are different for delayed and direct dispersal. Under delayed dispersal, we have

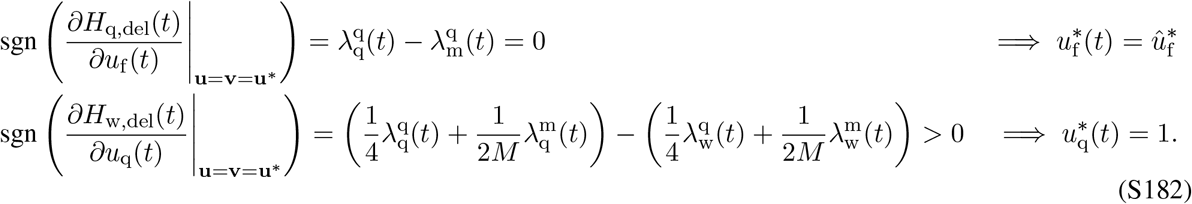

Under direct dispersal, we have during the time of male production 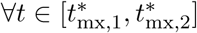 that

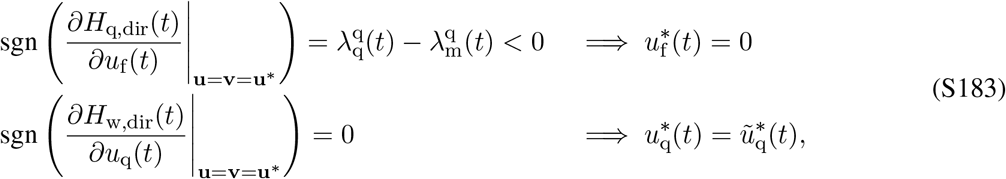

while during the time of queen production 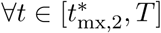:

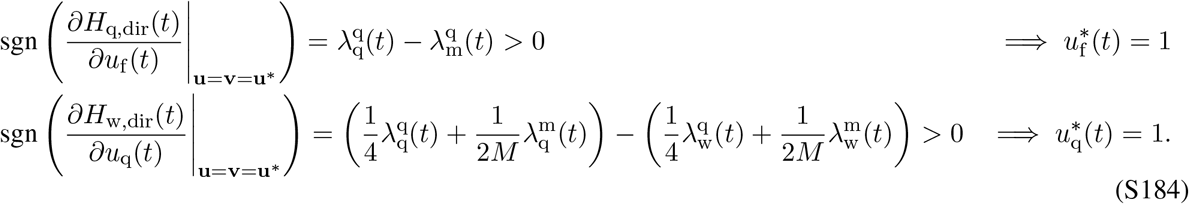

The sign of *∂H*_q,*d*_(*t*)/*∂u*_f_ (*t*) during the reproductive phase implies that females and males are produced simultaneously 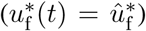 under delayed dispersal (eq. S182) since the marginal return of queens 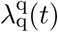 and males 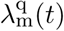 is equal in colonies founded by queens carrying the mutant allele. However, under direct dispersal (eqs. S183 and S184) males are produced first, since the marginal return of males 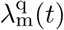 is initially higher and then becomes lower than that of queens 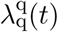 in colonies founded by queens carrying the mutant allele. The sign of *∂H*_w,*d*_(*t*)/*∂u*_q_(*t*) during the reproductive phase (eqs. S182, S183 and S184) implies that if any females are produced (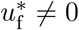 like in eq. S183) then all females become queens during the reproductive phase 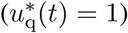, because the marginal return of queens 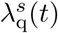 is higher that that of workers 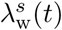, where the marginal returns have been weighed by the expected frequency of mutant alleles in workers in colonies founded by type *s* mutant individuals. In other words, queens are more valuable than workers to the genes residing in the workers. Under direct dispersal, during the production of males (eq. S183) the sign of *∂H*_w,dir_(*t*)/*∂u*_q_(*t*) = 0 because 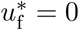. Hence, there is no directional selection on *u*_q_(*t*) for this time period, since it has no effect on invasion fitness (i.e. during the time only males are produced, the proportion at which workers rear female eggs into queens does not affect invasion fitness). Hence, during the reproductive phase, there is a latent trade-off between producing queens versus males from the perspective of the genes in the queens and a latent trade-off between producing workers versus queens from the perspective of the genes in the workers.

In summary, the signs of marginal returns *∂H*_*c,d*_(*t*)/*∂u*_*τ*_ (*t*) (eqs. S182, S183 and S184) defined over the entire season elucidate the main difference between the ergonomic and reproductive phase in terms of trade-offs between producing different individuals experienced by the two parties. From the perspective of the genes in the workers there is always a trade-off between producing queens versus workers, such that the balance is tipped in favour of workers during the ergonomic phase and in favour of queens during the reproductive phase. From the perspective of the genes in the queens, the trade-offs for the ergonomic and the reproductive phase are different. During the ergonomic phase, there is a trade-off between producing workers versus males and during the reproductive phase there is a trade-off between producing queens versus males. Intuitively, this “decoupling” of the trade-offs from the perspective of the queens happens because queens and workers are produced in separate phases. Hence, during the ergonomic phase the trade-off is only between producing workers and males, since workers rear all female eggs into workers and during the reproductive phase the tradeoff is only between producing queens and males, since workers rear all female eggs into queens. We can now turn to explaining how the trade-offs experienced by the two parties during different phases of colony growth have shaped the evolutionary outcome of the sex allocation conflict.

The queen determines the overall sex allocation ratio *S*_mx_ under mixed control. To see this, first note that the primary sex allocation ratio 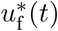 during the reproductive determines the overall sex allocation ratio *S*_mx_, since all females become queens (i.e. 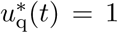) at the reproductive phase and sexuals are only produced during the reproductive phase (recall eqs. S92 and S148). It follows from eqs. (S182), (S183), (S184) that 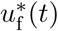 is determined by the difference between the marginal return of producing queens and males in a colony founded by queens carrying the mutant allele, i.e. the genes in the queens determines the overall sex allocation ratio *S*_mx_.

The workers determine the switching time 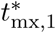 between the ergonomic and the reproductive phase under mixed control. Because the switching time 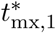 determines the colony size at its maturity 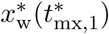 and colony productivity 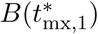 (recall eqs. S164 and S168), then it follows that the workers control also these quantities under mixed control. To see this, first recall that during the ergonomic phase there is a latent trade-off from the perspective of the queens between producing workers versus males and a latent trade-off from the perspective of the workers between producing workers versus queens (eq. S181). The ergonomic phase ends when for (at least) one of the parties, producing workers does not yield the highest marginal return anymore. For queens it means that the marginal value of producing males becomes higher than that of workers and for workers it means that the marginal value of producing queens becomes higher than that of workers. Hence, the switching time 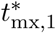 is determined by the party who prefers to end the ergonomic phase earlier. The switching times from the ergonomic to the reproductive phase are equal under full queen control and full worker control 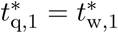 (see eq. 9). Hence, the marginal value of workers compared to sexuals is equal from the perspective of queens and workers. Note that the marginal value of queens and males depends on the overall sex allocation ratio *S*_mx_ (see eq. S86, S87, as boundary conditions for differential equations for the marginal returns S82). Since the queens control the overall sex allocation ratio *S*_mx_ under mixed control, then it means that it is more male-biased than preferred by the workers (*S*_w_ > *S*_mx_, see Fig. 6). Hence, from the perspective of workers, the marginal return of queens should be larger under mixed control than under full worker control, where the sex allocation ratio is also controlled by workers. Therefore, it follows from our intuitive explanation that the switch is controlled by the workers since the value of workers becomes equal to queens (from the perspective of workers) sooner than it becomes equal to males (from the perspective of queens). Hence, we can conclude that the genes in workers determine the switching time 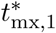 and therefore also the the colony size at its maturity 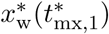 and colony productivity 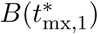.

## 9 Marginal return of producing a queen versus a male

The overall sex allocation ratio *S*_*c*_ is determined by the allocation trait 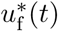 during the reproductive phase (see sections 7.3 and 8). This means that allocation to queens versus males is determined by the sign of the marginal return *∂H*_*c,d*_(*t*)/*∂u*_f_ (*t*) during the reproductive phase, which is given by eq. (S76) assuming that 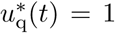, which yields

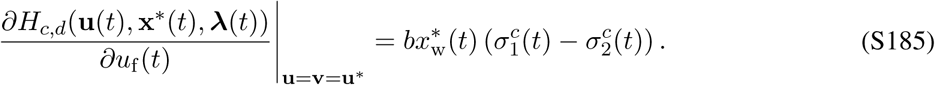

Substituting 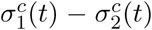 from eq. (S78) for queen control and eqs. (S79) and (S80) for worker control and using the expression for relatedness asymmetry *R*_*c*_ (eq. S35), we obtain

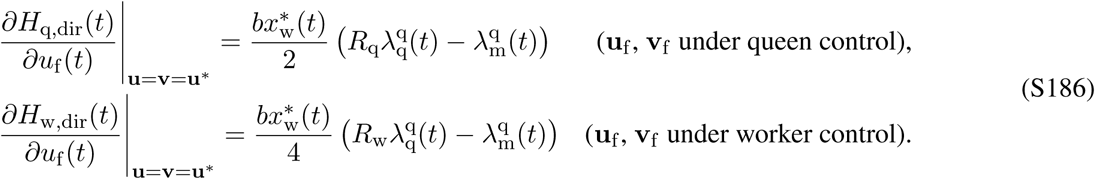

It follows from eq. (S90) that the marginal return of producing a male depends on the scenario of dispersal of reproductive individuals.

### 9.1 Delayed dispersal

Assuming that *µ*_q_ = *µ*_m_ = *µ*_r_, it follows from eq. (S90) (by taking *t*_1_ = *T*) for delayed dispersal

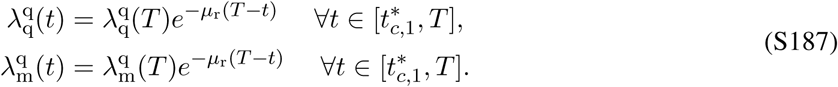

Substituting the transversality conditions (S86) and using eq. (S50), we obtain

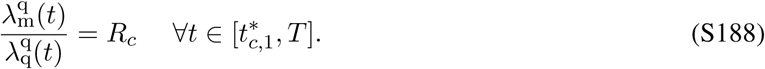

Hence the ratio of the marginal return of a male to the marginal return of a queen is equal to the relatedness asymmetry *R*_*c*_ at any time during the reproductive phase.

For consistency, we can substitute eq. (S188) into eq. (S186), which yields that *∂H*_*c,d*_(*t*)/*∂u*_f_ (*t*) = 0 throughout the reproductive phase 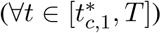.

### 9.2 Direct dispersal

It follows from eq. (S90), where *t*_1_ = *T* and 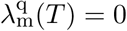 (from eq. S87) for direct dispersal and re-arranging

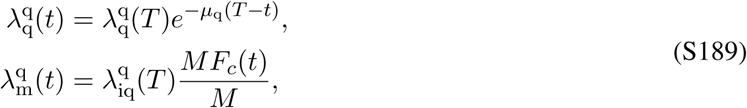

where

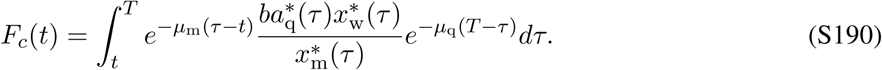

is the expected number of queens surviving until *T* that are inseminated by a male born at *t* (in a population, where the queens mates only once, i.e. *M* = 1). Note that, in a population where females mate *M* times, the expected number of surviving queens inseminated by a male born at *t* is equal to *MF*_*c*_(*t*). However, since the males are only expected to father 1/*M* of the offspring of the queen, that he would have fathered if the the queen would be singly mated, then the effect of multiple matings *M* by the queen cancels out. Lets denote by *l*(*t*) the probability that a queen produced at time *t* survives until the end of the season, which is given by

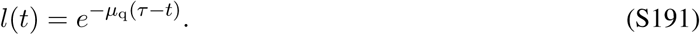

Under direct dispersal, *∂H*_*c*,dir_(*t*)/*∂u*_f_ (*t*) < 0 during 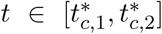 while only males are produced and *∂H*_*c*,dir_(*t*)/*∂u*_f_ (*t*) > 0 during 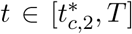 while only queens are produced. Hence, at time 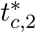 the marginal value 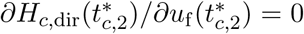. Assuming that the mortality of queens and males is equal (*µ*_q_ = *µ*_m_ = *µ*_r_), it follows then from eqs. (S186) and (S191) (S189)

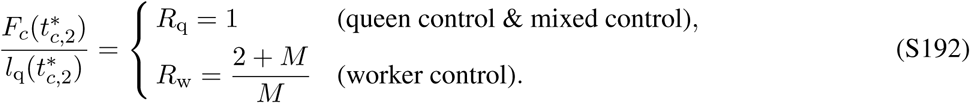

It follows from eq. (S192) that the switch from male production to queen production happens when producing a male instead of a surviving queen yields *R*_*c*_ (*R*_q_ = 1 and *R*_w_ = (2 + *M*)/*M*) surviving inseminated queens.

### 9.3 Verifying the consistency of eq. (S192)

Here, we show that the 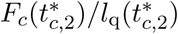 indeed satisfies eq. (S192) given the the explicit solutions for u^*^ x^*^ 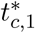, and 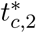 that we have already established. For this we need to evaluate *F* (*t*) at time 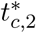 (assuming that the mortality of queens and males is equal, i.e. *µ*_q_ = *µ*_m_ = *µ*_r_) and the expression of 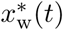 and 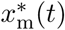 for the last phase 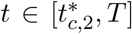.

In order to establish the initial conditions 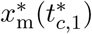 and 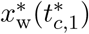 for the last phase, we use eq. (S88) for the penultimate phase 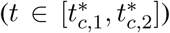 of the uninvadable state 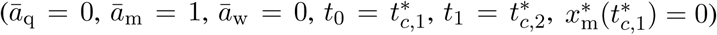 and assume that *µ*_q_ = *µ*_m_ = *µ*_r_. This allows us to express 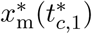 and 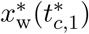 as

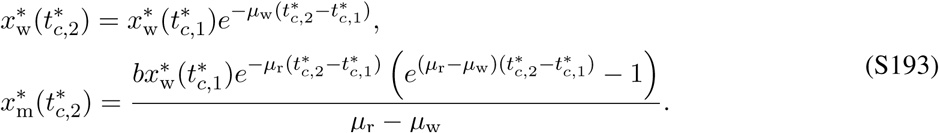

Next using eq. (S88) for the last phase 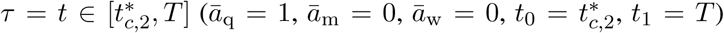, and taking eq. (S193) as an initial condition we obtain

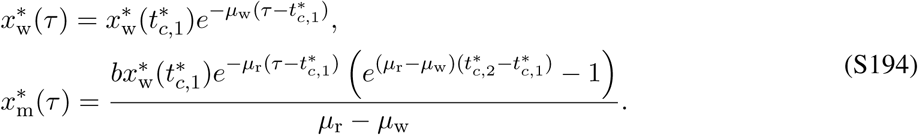

Now we can evaluate *F* (*t*) at time 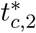 by substituting eq. (S194) into eq. (S190) (assuming *µ*_q_ = *µ*_m_ = *µ*_r_) and taking into account that 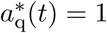 during the last phase 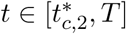 yields

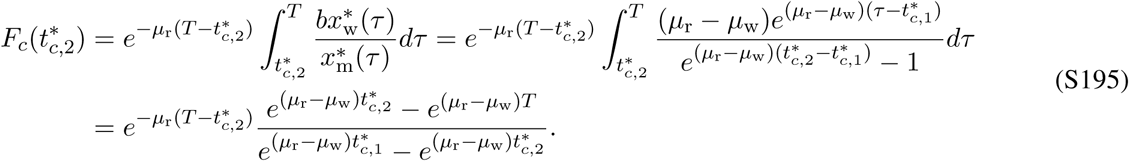

Substituting the expressions for the switching times under direct dispersal (eqs. S149, S150, S152, and S154), and simplifying, we get

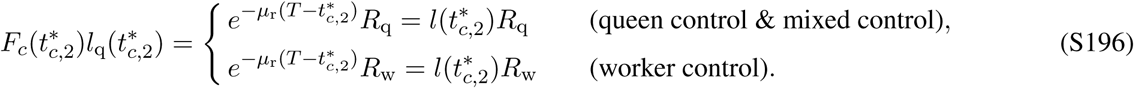

### 9.4 The overall sex allocation ratio and the ratio of expected surviving inseminated queens to surviving queens

It is significant that the necessary condition for **u**\* to be uninvadable under direct dispersal, is given by the condition that the ratio *F*_*c*_(*t*)/*l*(*t*) at time 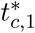 would be equal to the relatedness asymmetry and not on the condition on the overall sex allocation ratio *S*_*c*_. In this section we would like to make a connection between these two terms.

First, lets define the overall sex allocation ratio *Z*_*c*_ = *S*_*c*_/(1 − *S*_*c*_) (recall S172 for the formal definition of *S*_*c*_) as the ratio of the proportion of colony resources allocated queens versus males throughout the entire season (which would make it easier to compare overall allocation to queens versus males to the ratio 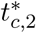. Hence, we can write *Z*_*c*_ as

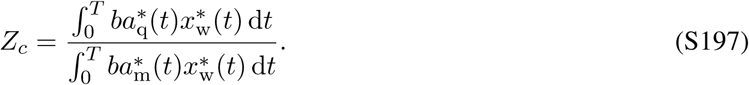

Substituting the uninvadable resource allocation schedule **u**\* for *µ*_m_ = *µ*_q_ = *µ*_r_ under direct dispersal given by eq. (S148) with the solutions to state equations given by (S89) into eq. (S197) yields

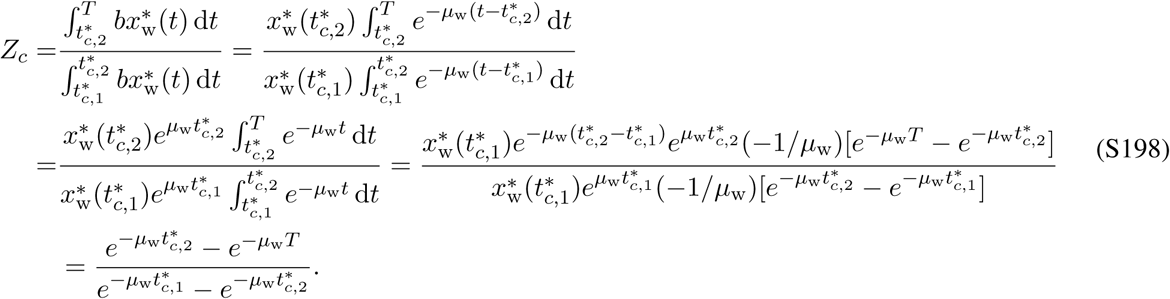

The overall allocation to queens versus males under direct dispersal is therefore given by

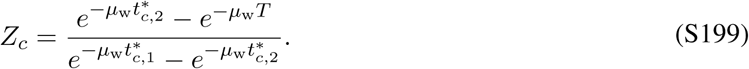

It follows from eq. (S195) that the ratio 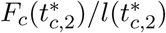 can be expressed as

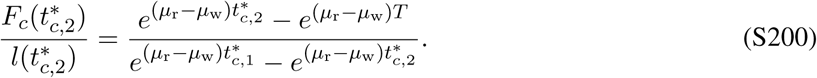

Hence, one can see from eqs. (S199) and (S200) that

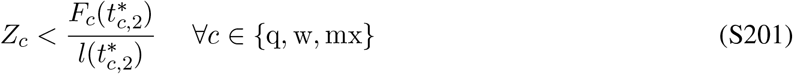

whenever *µ*_r_ > 0 and the difference 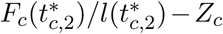 is larger for higher values of *µ*_r_. Hence, the overall sex allocation ratio is more male-biased under direct dispersal than expected from the classical results (e.g. Reuter and Keller, 2001) for higher values of mortality of reproductive individuals.

In order to get a better intuition why the overall sex allocation ratio is more male-biased for higher values of mortality *µ*_r_ of reproductive individuals, lets examine how *µ*_r_ influences *F*_*c*_(*t*)/*l*(*t*) during the reproductive phase which we can express as (using eq. S190 and S191)

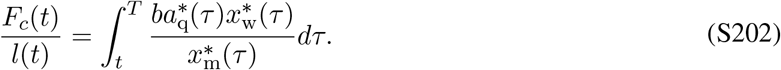

Here, 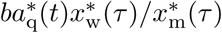 gives the mating success of a male (the number of queens available to mate per male at time *τ*). Here, 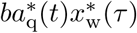 is independent of the mortality of sexuals *µ*_r_, since 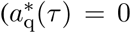, during male production and 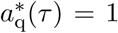 during queen production) and number of workers 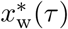 does not depend on the mortality of sexuals. In contrast, the number of males 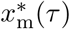 alive at time *τ* during the last phase, when only females are produced is smaller for higher values of *µ*_r_. Hence, it follows that *F*_*c*_(*t*)/*l*(*t*) is higher for higher mortality rate of sexuals (for a given *t* in the reproductive phase), because higher mortality increases the mating success of a male alive at a given time *t*. Since, the ratio *F*_*c*_(*t*)/*l*(*t*) gives the expected number of surviving queens inseminated by a male produced instead of a queen, then the surviving probability of a focal male together with the surviving probability of the queen(s) he inseminates cancels out with the surviving probability of a queen that would have been otherwise produced (since we assumed that the mortality of queens and males is equal). Because of this, the ratio *F*_*c*_(*t*)/*l*(*t*) increases with the increase in the mortality of sexuals via the mating success of a focal male 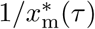.

## 10 Continuous stability of the candidate uninvadable allocation schedule

In this section we address the issue of (continuous) stability of the candidate uninvadable allocation schedule **u**\* given by eq. (S92) for delayed dispersal and eq. (S148) for direct dispersal. We only discuss the continuous stability of the candidate uninvadable allocation schedule **u**\* for equal mortality rate of queens and males (*µ*_q_ = *µ*_m_ = *µ*_r_) because we have fully derived the analytical results only under this assumption. Continuous stability is given by two separate properties of the of the candidate uninvadable allocation schedule **u**\*: (i) the local uninvadablity and (ii) convergence stability (e.g. see Christiansen, 1991; Eshel, 1983; Taylor, 1989 and for functioned-valued traits see Dieckmann et al., 2006). The candidate uninvadable allocation schedule **u**\* is locally uninvadable if a monomorphic population following the strategy **u**\* can resist invasion by any mutant whose strategy is close to the the strategy **u**\*. The candidate uninvadable allocation schedule **u**\* is convergence stable if a population will converge to this schedule **u**\* through recurrent substitutions, meaning that a mutant whose schedule is closer to **u**\* will invade a monomorphic population that follows a schedule further away from **u**\*.

Firstly, we would like to point out that the continuous stability of the candidate uninvadable allocation schedule **u**\* (assuming that *µ*_q_ = *µ*_m_ = *µ*_r_) is not directly given from the first-order condition only for 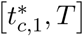 under delayed dispersal. Indeed, if *∂H*_*c,d*_(**u**(*t*), **x**\*(*t*), ***λ***(*t*))/*∂u*_*τ*_ (*t*)|_**u**=**u\***_ > 0 then a mutant allele with *u*_*τ*_ (*t*) > *v*_*τ*_ (*t*) can always spread (recall eq. S73). Similarly, if *∂H*_*c,d*_(**u**(*t*), **x**\*(*t*), ***λ***(*t*))/*∂u*_*τ*_ (*t*)|_**u**=**u\***_ < 0 then a mutant allele with *u*_*τ*_ (*t*) < *v*_*τ*_ (*t*) can always spread. However, for a finite period of time, for which *∂H*_*c,d*_(**u**(*t*), **x**\*(*t*), ***λ***(*t*))/*∂u*_*τ*_ (*t*)|_**u**=**u\***_ = 0 holds and the focal trait is a singular arc 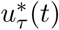, then there is no directional selection. Hence, the two properties of continuous stability of **u**\* has to be only addressed for 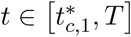 under delayed dispersal, where 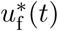 is a singular arc.

The conditions for local uninvadability and convergence stability are given in terms of second-order functional derivatives of the invasion fitness (S10) (Dieckmann et al., 2006). Since the formal continuous stability analysis for function-valued traits is out of the scope of this paper, we will not address the questions of continuous stability of **u**\* for 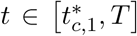 under delayed dispersal any further in this paper. However, since we recover the uninvadable allocation schedule numerically through iteration, which is analogous to the evolutionary dynamics of the population, then we might regard this as giving support that the proper evolutionary dynamics would also converge to this schedule. Under this heuristic approach, the iterative scheme of the best response map also implies continuous stability of the candidate uninvadable schedule found by it (Houston and McNamara, 1999).

## 11 Iterative scheme of the best response map

Here, we describe the computational technique for finding the locally uninvadable strategies for our optimal control problems. This method is known as the iterative scheme of the best response map (see p. 187 in Houston and McNamara, 1999).

A mutant schedule 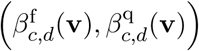 that yields the highest invasion fitness in a population, where resident schedule is **v**, i.e.

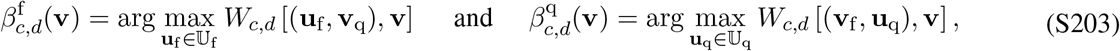

is said be the best response to the resident schedule **v**. Here, 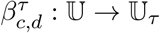, where *τ* ∈ f, q is the best-response correspondence which maps a resident schedule **v** ∈ 𝕌 to a (unique) trajectory 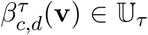 for a trait type *τ*, such that no other trajectory for a focal trait gives a higher invasion fitness to a mutant in a population, where resident individuals follow the schedule **v** ∈ 𝕌. Here, 𝕌 = 𝕌_f_ *×* 𝕌_q_ is a set of all possible allocation strategies, 𝕌_f_ and 𝕌_q_ are sets of all possible trajectories for the traits **u**_f_ (**v**_f_) and **u**_q_ (**v**_q_), respectively. In the notation of the best-response correspondence 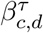, the subscripts *c* ∈ {q, w} and *d* ∈ {del, dir} emphasize the party in control and the time of dispersal of sexuals, respectively, and superscript *τ* ∈ {f, f} emphasizes the trait type.

Hence, under single-party control, where party *c* ∈ {q, w} is in full control, the best response schedule 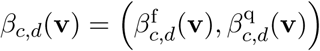 can be written as

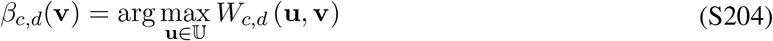

and under mixed control the best response 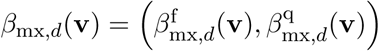 can be written as

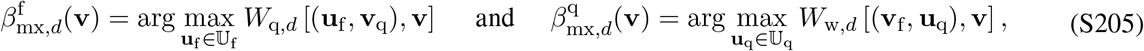

where *β*_*c*_: 𝕌 → 𝕌 under control mode *c* ∈ {q, w, mx} is the best-response correspondence which maps to a (resident) schedule **v** ∈ 𝕌 a schedule *β*_*c*_(**v**) ∈ 𝕌, such that no other schedule gives a higher invasion fitness to a mutant in a population, where resident individuals follow the schedule **v** ∈ 𝕌.

It follows from the definition of the uninvadable schedule given by eq. (S12) for single-party control and by eq. (S13) for mixed control that the uninvadable schedule is a best response to itself, i.e.

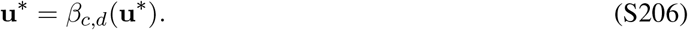

Note that, here we have assumed that the best response is always unique.

In order to approximate the uninvadable schedule numerically, we start out from some initial resource allocation schedule for the resident population **u**^0^ and using GPOPS (Patterson and Rao, 2014) we find the mutant schedule that has the highest fitness **u**^1^ = *β*_*c,d*_(**u**^0^). The software GPOPS uses a direct approach to find the best response *β*_*c,d*_(**v**) for a given environment **v** in contrast to the indirect approach of Pontryagin’s maximum principle (see section 3.2), which gives a necessary condition for optimality. We then update the resident schedule for the next iteration

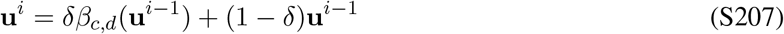

and repeat the process. Here, 0 > *d* > 1 is called the replacement factor. We can interpret this new resident schedule as a polymorphism - each individual adopting a schedule *β*_*c,d*_(**u**^*i*−1^) with probability *α* and schedule **u**^*i*−1^ with probability (1 − *δ*) (Houston and McNamara, 1999). To improve convergence after iterating from some while we can decrease *δ* with further iterations (Houston and McNamara, 1999; Krawczyk and Uryasev, 2000).

This iterative scheme forms a sequence of strategies (**u**^0^; **u**^1^; **u**^2^; *…*) where each schedule is derived from the best response to the previous schedule according to equation (S207). If the difference between the best response and resident schedule approaches zero as the number of iterations increases, i.e.

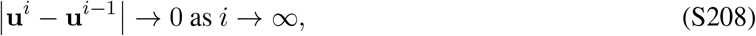

then we have arrived at the uninvadable schedule (Nash equilibrium).

For single-party control we use GPOPS to find the best response *β*_*c,d*_(**v**) that maximizes the objective *W*_*c,d*_(**u**, **v**) given by eq. (S10) of the party *c* in control. For mixed control, the best response 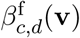 maximizes the the objective of the queen *W*_q,*d*_(**u**, **v**) and 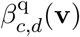 maximizes the objective of the workers *W*_w,*d*_(**u**, **v**).

## 12 Static resource allocation model with a linear relationship between colony productivity and colony size

In this section we re-derive the main results of Reuter and Keller (2001) with a slight modification in the assumption about how the overall colony productivity scales with colony size. Here we assume that the overall colony productivity scales linearly with colony size and show that the main predictions of their model about how sex allocation conflict affect allocation strategy in the colony are not affected by the assumption about how overall colony productivity scales with colony size. In order to make a direct comparison easier for the reader, we slightly modified the notation in Reuter and Keller (2001) to make it easier to compare our model with theirs, however the modelling approach in this section is exactly identical to Reuter and Keller (2001). Here we present the main features of the model assuming monogamy, for the full model and further details, see the original paper by Reuter and Keller (2001).

Let 0 ≤ *u*_f_ ≤ 1 (*u*_f_ = *f* in their notation) be the proportion of colony resources allocated into producing females in a focal colony and let 0 ≤ *u*_q_ ≤ 1 (*u*_q_ = 1 − *w* in their notation) be the proportion of resources allocated into queens from the resources allocated into females. Let the the corresponding population average traits be 0 ≤ *v*_f_ ≤ 1 and 0 ≤ *v*_q_ ≤ 1 (*F* and 1 − *W* in their notation), respectively. Here the allocation strategies *u*_f_, *u*_q_ (*v*_f_, *v*_q_) give the allocation of all colony resources over the entire season.

Since this model is a static resource allocation model, colony size is given by the proportional investment *u*_f_ (1*-u*_q_) into workers. Reuter and Keller (2001) assume that per-worker productivity declines with the number of workers in the colony, such that overall colony productivity *b*(*u*_f_, *u*_q_) (interpreted in their paper as the total biomass of all individuals produced) in the focal colony follows a diminishing return function

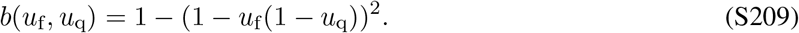

Next, they formulate an expression for the fitness function (*V*_*X*_ in their notation)

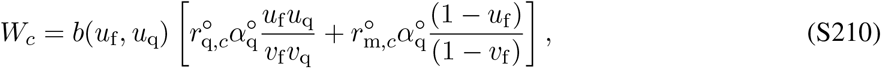

where the subscript *c* ∈ {q, w} emphasizes the party in control. See eq. 2 in Reuter and Keller, 2001 for further details about the interpretation of this fitness function. Essentially, *W*_*c*_ in (S210) is comparable to the invasion fitness *W*_*c*_ in (S9). However, we note that this way of defining an invasion fitness function is heuristic and implicitly makes the assumption that only a first-order analysis will be carried out (e.g. otherwise relatedness coefficients cannot be hold at neutrality), and so the eq. (S210) can not be used to check second order conditions (further it should be normalized so that in a resident population fitness is one). Candidate uninvadable allocation strategy 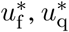 (called evolutionary stable strategy and noted as *f* *, *w*\* in Reuter and Keller, 2001) under single-party *c* control is given by

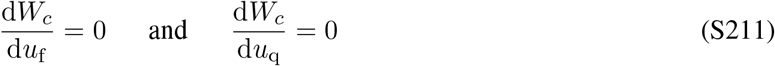

and under mixed control is given by

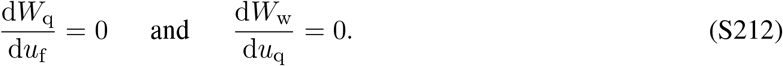

The results of Reuter and Keller (2001) assuming monogamy are outlined in table S2. The main results of Reuter and Keller (2001) can be summarized as follows: less resources are allocated into worker production under single-party control than under mixed control, the uninvadable sex allocation ratio is equal to the relatedness asymmetry under single-party control (*R*_q_ = 1 for queen control and *R*_w_ = 3 under worker control, assuming monogamy). Under mixed control the uninvadable sex allocation ratio has a value intermediate ≈ 1.26 between the relatedness asymmetries for queen and worker control.

**Table S2:**
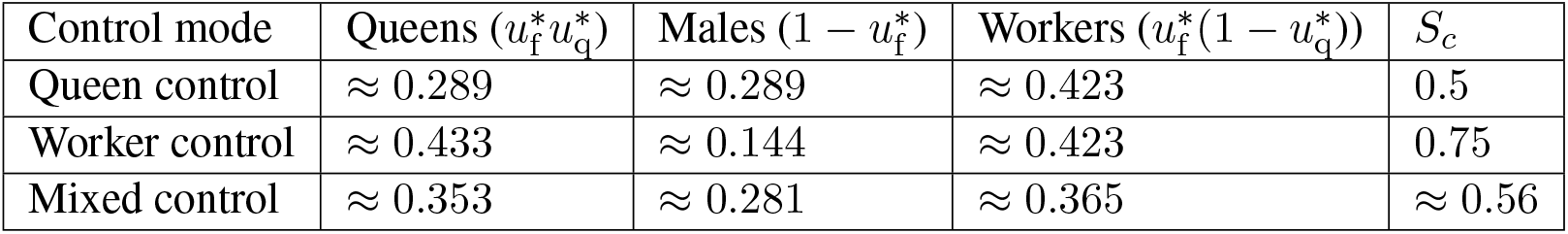
Uninvadable allocation into queen, males, and workers and the overall sex allocation ratio *S*_*c*_ (proportional allocation to queens from resources allocated to sexuals) predicted by Reuter and Keller (2001).

Lets now modify the overall colony productivity *b*(*u*_f_, *u*_q_), such that it increases linearly with the amount of resources invested into workers *u*_f_ (1 − *u*_q_).

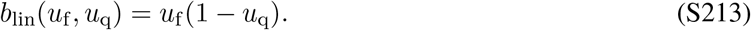

Substituting *b*_lin_(*u*_f_, *u*_q_) into the fitness function (S210) instead of *b*(*u*_f_, *u*_q_) and solving for the uninvadable allocation strategy according to eqs. (S211)–(S212), we obtain the uninvadable allocation strategy 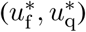 assuming that colony productivity scales linearly with colony size. The results are outlined in table S3. The main results of the static model, where colony productivity scaled linearly with colony size can be summarized as follows: less resources are allocated into worker production under single-party control than under mixed control, the uninvadable sex allocation ratio is equal to the relatedness asymmetry under single-party control (*R*_q_ = 1 for queen control and *R*_w_ = 3 under worker control, assuming monogamy). Under mixed control the uninvadable sex allocation ratio has a value intermediate ≈ 1.28 between the relatedness asymmetries for queen and worker control.

Hence, we can conclude, that the main results predicted by a static allocation model of Reuter and Keller (2001) about how sex allocation conflict affects allocation strategies in the colony are not qualitatively affected by the assumption about how colony productivity scales with colony size.

**Table S3:**
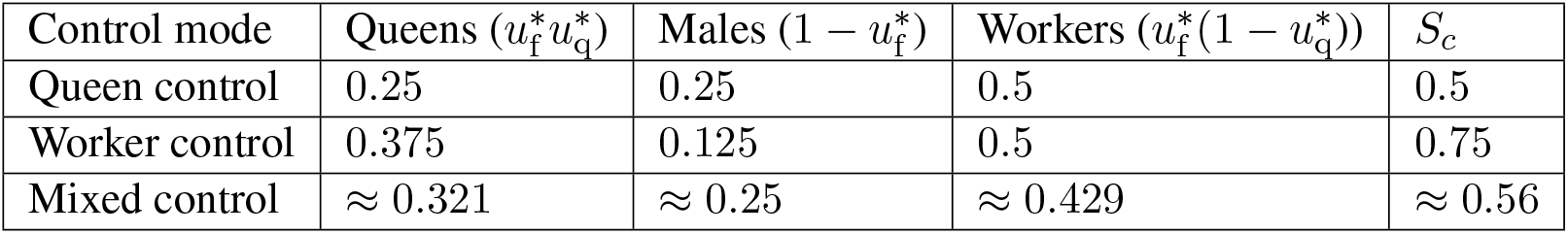
Uninvadable allocation into queen, males, and workers and the overall sex allocation ratio *S*_*c*_ (proportional allocation to queens from resources allocated to sexuals) predicted by a static model similar to Reuter and Keller (2001), assuming that colony productivity scales linearly with colony size.

## 13 How our model connects to previous tightly related literature

In this section we provide brief overview of how our model (under different assumptions about the time of dispersal and genetic control of the resource allocation traits) relates to a selection of previous models studying resource allocation in social insects.

### 13.1 Delayed dispersal

Macevicz and Oster (1976) were the first to develop a dynamic resource allocation model for eusocial insects, they assumed clonal reproduction, two classes of individuals (sexuals and workers) and delayed dispersal (sexuals have to survive until the end of the season to have reproduce). Our model first grew out of the idea to combine the works of Macevicz and Oster (1976) and Reuter and Keller (2001). Our model under delayed dispersal (for all three scenarios of genetic control) can be viewed as a dynamic extention of the static model by Reuter and Keller (2001). The only fundamental difference between our model under delayed dispersal and the model of Reuter and Keller (2001), besides that their model is static, is that they assumed that per-worker productivity decreases when colony size increases. We assume that per-worker productivity is constant and does not depend on the size of the colony. We show in section 12 that this assumption does qualitatively alter the predictions their model. However, this does not mean that we do not think that the assumption about how per-worker colony productivity scales with colony size can affect dynamic colony allocation strategies. Poitrineau et al. (2009) developed a dynamic resource allocation model similar to Macevicz and Oster (1976), to study how the non-linearity of colony productivity rate (i.e. *by*_w_) can affect the allocation strategy of the colony. Poitrineau et al. (2009) show that if per-worker productivity decreases with colony size then, the sexuals are workers are expected to be produced simultaneously through a significant period of time in the season, after which the colony switched to producing only sexuals. However, it is not known how sex allocation conflict interacts with non-linearity of colony productivity rate, since Poitrineau et al. (2009) assumed clonal reproduction. Bulmer (1981) was the first to study how sex allocation conflict can affect dynamic allocation strategies within a colony. However, he assumed discrete non-overlapping generations of workers within a season and he only studied how conflicts affects allocation strategies during the two generations at the end of the season. Furthermore, his model was not able to make any analytical predictions (e.g. when should colony switch from producing workers to producing sexuals, etc). Our prediction that the queen wins the sex allocation conflict contradicts the predictions of a static model by Reuter and Keller (2001), but is in accordance to the prediction by Bulmer (1981). However, the conflict outcome of our model is different from Bulmer (1981). He predicted that, due to the sex allocation conflict, the colony dies before the end of the season, which is a prediction linked with the assumption about discrete non-overlapping generation of workers within a season (if the sex allocation ratio at the population level is not even because workers rear worker-destined eggs into queens, then the queens only produce males during the penultimate generation). Ohtsuki and Tsuji (2009) also considered a dynamic resource allocation model assuming delayed dispersal under mixed control, but their model assumed a slightly different biological scenario, with reproductive workers and worker policing, which is not considered here. In addition, the results of Ohtsuki and Tsuji (2009) are only numerical.

### 13.2 Direct dispersal

Direct dispersal of sexuals is fundamentally a dynamic aspect, and hence can captured with a dynamic resource allocation model. Only paper that we are aware that has studied colony resource allocation assuming direct dispersal is that of Bulmer (1983). He studied the effect direct dispersal on resource allocation strategies assuming queen control. The result of his paper have never been previously extended to worker control nor to mixed control.

## 14 Summary of notation

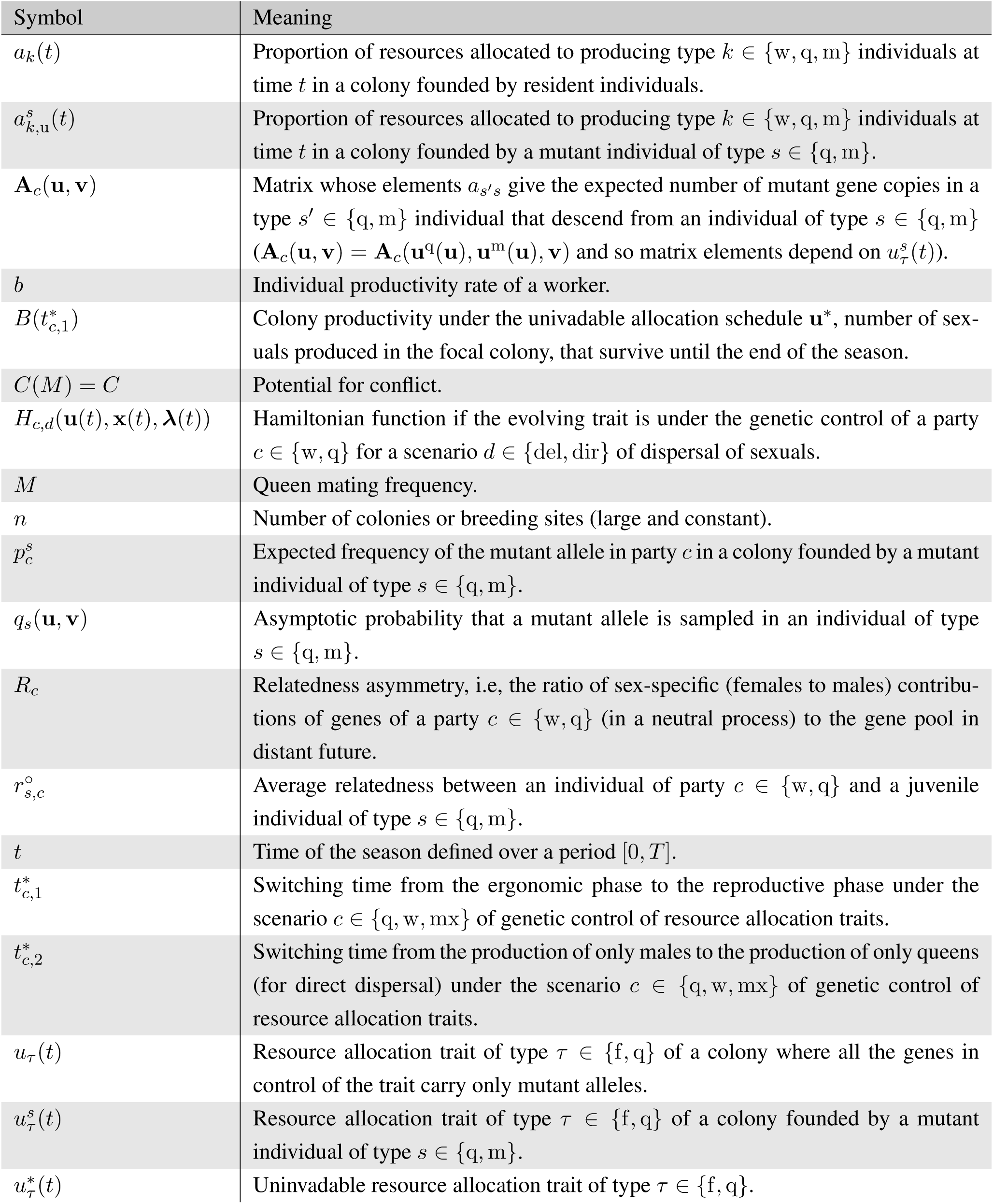
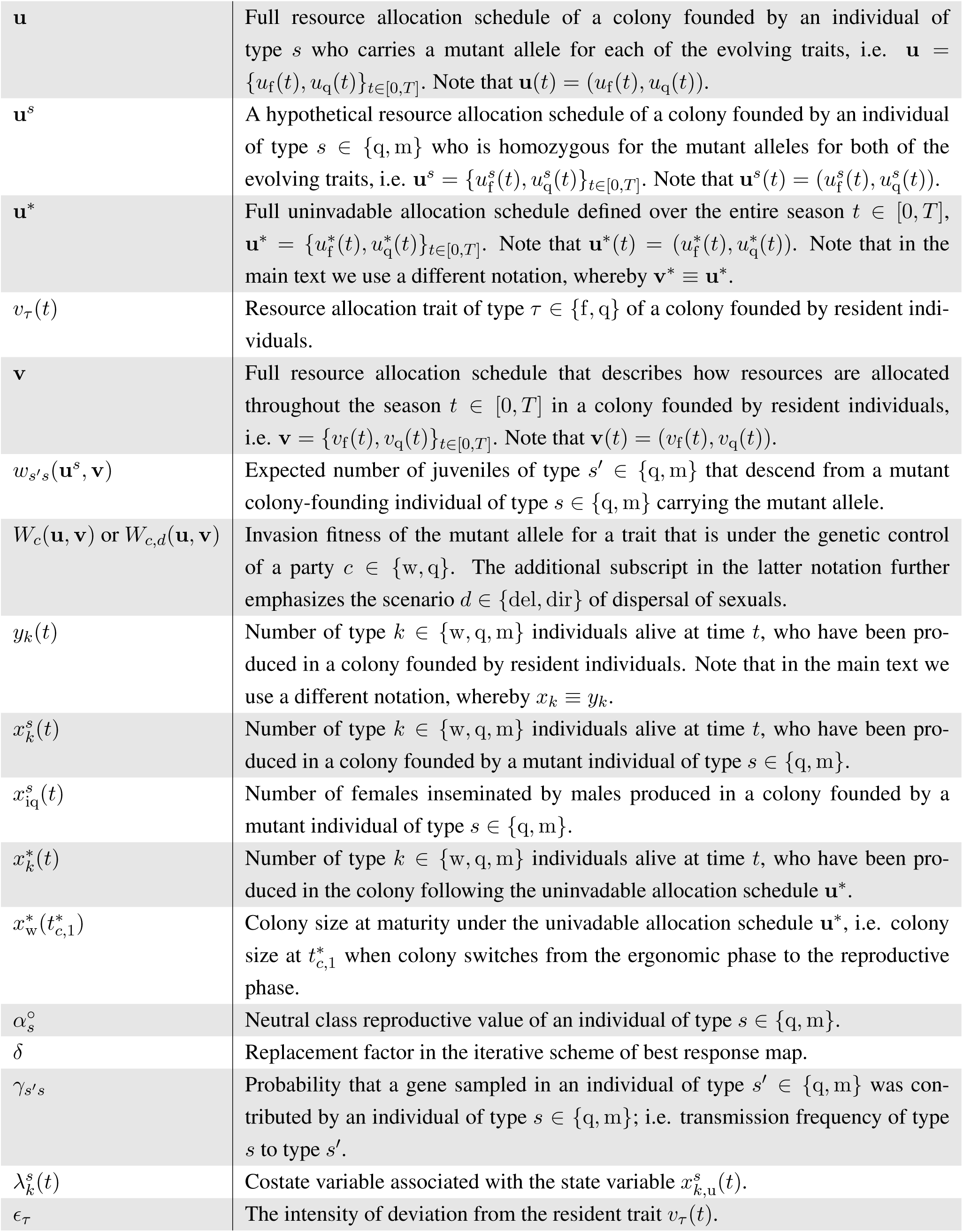
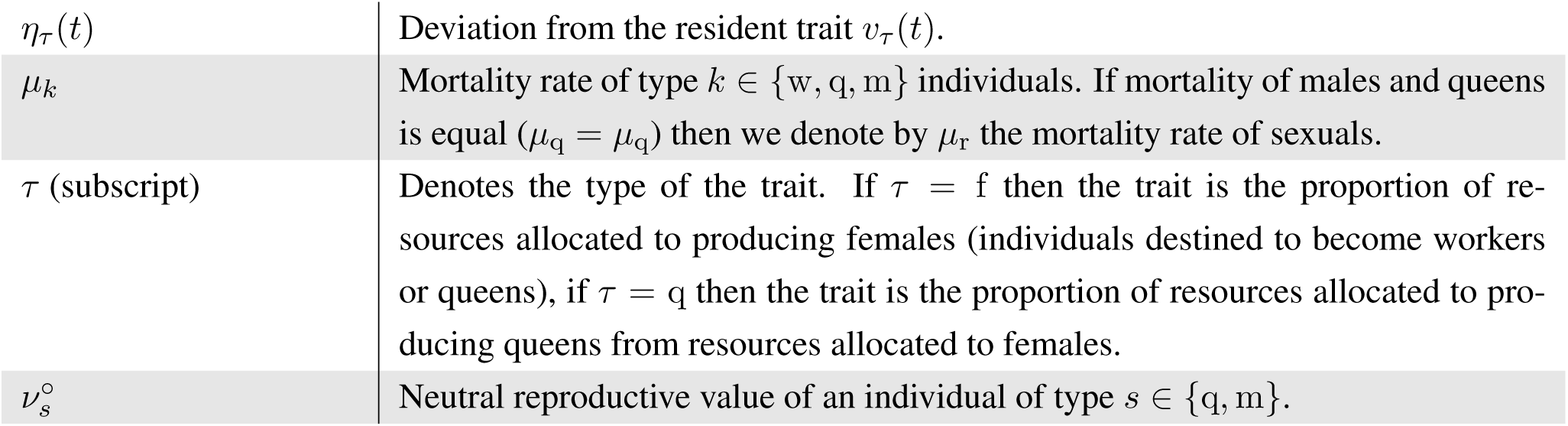

