## Supplementary Information for "Sex allocation conflict and sexual selection throughout the lifespan of eusocial colonies"

**10 Continuous stability of the candidate uninvadable allocation schedule** **91**

**11 Iterative scheme of the best response map** **92**

**12 Static resource allocation model with a linear relationship between colony productivity and colony size** **94**

**13 How our model connects to previous tightly related literature** **96**
13.1 Delayed dispersal . . . . . . . . . . . . . . . . . . . . . . . . . . . . . . . . . . . . . . . 96
13.2 Direct dispersal . . . . . . . . . . . . . . . . . . . . . . . . . . . . . . . . . . . . . . . . 97

**14 Summary of notation** **98**

### 1 Evolutionary analysis

#### 1.1 Mutant-resident system

We analyze the evolution of allocation schedules by performing an evolutionary invasion analysis (e.g., Caswell, 2001; Charlesworth, 1994; Eshel and Feldman, 1984; Ferrière and Gatto, 1995; Fisher, 1930; Lehmann et al., 2016; McNamara et al., 2001; Metz, 2011; Otto and Day, 2007) whereby we consider the fate (invasion or extinction) of a single mutant allele introduced into a population of resident individuals, where the mutant allele determines an allocation schedule that is different from the resident allocation schedule throughout the entire season. We assume that the traits are determined at separate single loci with two segregating alleles (resident and mutant).

Let $v_\tau(t)$ and $u_\tau(t)$ denote the resident and mutant resource allocation phenotypes for a trait of type $\tau \in \{\mathrm{f}, \mathrm{q}\}$, respectively. It will turn out to be useful to define the mutant phenotype $u_\tau(t)$ as a trait expressed by a (hypothetical) colony where all the genes in control of the trait are mutant alleles (i.e. individuals whose genes are in control of the trait are homozygous for the mutant allele).

Since the fate (invasion or extinction) of a mutant allele is determined when it is rare in the population, then only one of the colony "founding" individuals is carrying a single copy of the mutant allele (heterozygous diploid female or a hemizygous haploid male). By writing that a mutant male "founds" a colony, we mean that a mutant male has mated with a resident female that gives rise to a focal colony, where the mutant allele is present in the genes of the workers. Furthermore, in haplodiploid systems, where females are diploid and males are haploid, the phenotypes of colonies founded by mutant individuals of opposite sexes will be different. The distinction between the phenotypes expressed in these two types of colonies will also turn out to be useful when describing these phenotypes under the various assumptions of genetic control that different parties have over the traits.

Thus, we will denote by $u_{\mathrm{f}}^{\mathrm{q}}(t)$ (and respectively, by $u_{\mathrm{f}}^{\mathrm{m}}(t)$) the proportion of resources allocated at time $t$ to producing females in a colony founded by a focal mutant heterozygous female (hemizygous male) and by $u_{\mathrm{q}}^{\mathrm{q}}(t)$ (and respectively, by $u_{\mathrm{q}}^{\mathrm{m}}(t)$) the proportion of resources allocated to producing queens from resources allocated to females at time $t$ in a colony founded by a focal mutant heterozygous female (hemizygous male). Let $u_\tau^s(t)$ denote the resource allocation phenotype for a trait of type $\tau \in \{\mathrm{f}, \mathrm{q}\}$ of a colony founded by a heterozygous (hemizygous) individual of type $s \in \{\mathrm{q}, \mathrm{m}\}$ and it can be expressed as (assuming additive genetic effects)

$$u_\tau^s(t) = p_c^s u_\tau(t) + (1 - p_c^s) v_\tau(t), \tag{S1}$$

where $p_c^s > 0$ is the expected frequency of the mutant allele in party $c \in \{\mathrm{q}, \mathrm{w}\}$ in full control of the trait of type $\tau$ in a colony founded by a mutant individual of type $s$ ($s \in \{\mathrm{q}, \mathrm{m}\}$). Hereinafter, the subscript $c = \mathrm{q}$ denotes a scenario of full queen control and $c = \mathrm{w}$ denotes a scenario of full worker control, and $c = \mathrm{mx}$ denotes a scenario of mixed control.

Under queen control of the trait, the expected colony phenotype $u_\tau^s(t)$ is determined from the frequency of the mutant allele in the colony-founding queen. If the colony is founded by a heterozygous mutant female then the frequency of the mutant allele in the colony-founding queen is $p_{\mathrm{q}}^{\mathrm{q}} = 1/2$. Under queen control, mutant males who have mated with a colony-founding queen have no genetic influence on the resource allocation traits and thus $p_{\mathrm{q}}^{\mathrm{m}} = 0$. Hence, female mating frequency will also have no affect on the trait under queen control. Under worker control of the trait, the expected colony phenotype $u_{\tau}^{s}(t)$ is determined from the expected frequency of the mutant allele in workers. The expected frequency of the mutant allele in workers in a colony founded by a heterozygous mutant female is $p_{\mathrm{w}}^{\mathrm{q}} = 1/4$ and it is not affected by the mating frequency of the queen because a mutant female will only encounter resident males since the mutant allele is considered to be rare. The expected frequency of the mutant allele in workers in a colony founded by a mutant male is $p_{\mathrm{w}}^{\mathrm{m}} = 1/(2M)$, where $M$ is the number of times the female has mated (when the mutant allele is rare, only one of the males is carrying the mutant allele).

Let $\mathbf{v} = \{v_{\mathrm{f}}(t), v_{\mathrm{q}}(t)\}_{t\in[0,T]}$ denote the full allocation schedule of a colony founded by resident individuals, i.e. it describes how colony resources are allocated throughout the entire season $t \in [0, T]$. Similarly, let $\mathbf{u} = \{u_{\mathrm{f}}(t), u_{\mathrm{q}}(t)\}_{t\in[0,T]}$ denote the full allocation schedule of a colony founded by individuals who carry only mutant alleles for both of the evolving traits. Similarly, let $\mathbf{u}^{s} = \{u_{\mathrm{f}}^{s}(t), u_{\mathrm{q}}^{s}(t)\}_{t\in[0,T]}$ denote the full allocation schedule of a colony founded by a heterozygous (hemizygous) individual of type $s$ for each of the evolving traits, hence $\mathbf{u}^{s}$ depends on $\mathbf{u}$. This notation turns out to be useful for performing the invasion fitness analysis, but it does not imply that we are assuming pleiotropic effects.

Let $a_{k,\mathrm{u}}^{s}(t)$ be the proportion of resources allocated to producing type $k \in \{\mathrm{w}, \mathrm{q}, \mathrm{m}\}$ individuals in a colony founded by a heterozygous individual of type $s \in \{\mathrm{q}, \mathrm{m}\}$, where the subscript "u" in $a_{s,\mathrm{u}}^{s}(t)$ emphasizes that it is the mutant allocation schedule, which, according to eq. (3) is

$$a_{\mathrm{w,u}}^{s}(t) = u_{\mathrm{f}}^{s}(t)(1 - u_{\mathrm{q}}^{s}(t)), \qquad a_{\mathrm{q,u}}^{s}(t) = u_{\mathrm{f}}^{s}(t)u_{\mathrm{q}}^{s}(t), \qquad a_{\mathrm{m,u}}^{s}(t) = (1 - u_{\mathrm{f}}^{s}(t)). \tag{S2}$$

The rate of change in the number of type $k \in \{\mathrm{w}, \mathrm{q}, \mathrm{m}\}$ individuals alive at time $t$, that have been produced in a colony founded by a mutant individual of type $s$, is given by the equation

$$\frac{\mathrm{d}x_{k}^{s}(t)}{\mathrm{d}t} = ba_{k,\mathrm{u}}^{s}(t)x_{\mathrm{w}}^{s}(t) - \mu_{k}x_{k}^{s}(t), \ x_{k}^{s}(t_0) = y_{k0}, \tag{S3}$$

where $x_{k}^{s}(t)$ denotes the number of individuals of type $k$ alive at time $t$ that have been produced in a colony founded by a mutant individual of type $s$. The rate of change of females $x_{\mathrm{iq}}^{s}(t)$ alive at time $t$, who have been inseminated by the males produced in the focal colony (under a monandrous mating system) founded by a mutant individual of type $s$, is given by the equation

$$\frac{\mathrm{d}x_{\mathrm{iq}}^{s}(t)}{\mathrm{d}t} = \begin{cases} 0, \ \text{for } t < T, \ \text{with } x_{\mathrm{iq}}^{s}(T) = x_{\mathrm{m}}^{s}(T)\frac{y_{\mathrm{q}}(T)}{y_{\mathrm{m}}(T)} & \text{(delayed dispersal)}, \\ x_{\mathrm{m}}^{s}(t)\frac{ba_{\mathrm{q}}(t)y_{\mathrm{w}}(t)}{y_{\mathrm{m}}(t)} - \mu_{\mathrm{q}}x_{\mathrm{iq}}^{s}(t), \ x_{\mathrm{iq}}^{s}(0) = 0 & \text{(direct dispersal)}. \end{cases} \tag{S4}$$

Note that the number of females alive at time $t$, who have been inseminated by the males produced in the focal colony founded by a mutant individual of type $s$ in a mating system where females mate $M$ times is $Mx_{\mathrm{iq}}^{s}(t)$.

The rate of change of individuals of type $k \in \{\mathrm{w}, \mathrm{q}, \mathrm{m}\}$ produced in a resident colony and females inseminated by males from the resident colony are given by eq. (1) and eq. (2), respectively.

##### 1.2 Fitness functions of mutant individuals

We will express the invasion fitness of the mutant allele in terms of gene transmission frequencies and fitness functions of mutant individual of type $s$ ($s \in \{\mathrm{q}, \mathrm{m}\}$), who have founded colonies in the current season. The fitness is measured as the expected number of daughters and sons who become colony founders in the next generation. Note that, since females mate with $M$ males, a son can become a colony founder for multiple colonies. Hence, the number of sons who become colony founders in the next generation can be measured in the number of females inseminated by sons, who will become colony-founding queens in the next generation (Bulmer, 1994, p. 213). Hence, one fitness measurement cycle lasts from the beginning of the current season to the beginning of the following season and we keep track of the genes in mutant colony-founding individuals of type $s$.

Let $w_{s's}(\mathbf{u}^s, \mathbf{v})$ denote the expected number of mutant colony-founding individuals of type $s' \in \{\mathrm{q}, \mathrm{m}\}$ in the following season that descend from a current colony-founding mutant individual of type $s \in \{\mathrm{q}, \mathrm{m}\}$ in a resident population. The fitness function $w_{s's}(\mathbf{u}^s, \mathbf{v})$ is a function of the allocation schedule $\mathbf{u}^s$ of a colony founded by an individual of type $s$ (by way of eqs. S2–S4). Note that the fitness function $w_{s's}(\mathbf{u}^s, \mathbf{v})$ is ultimately a function of the mutant schedule $\mathbf{u}$, the frequency $p_c^s$ of the mutant allele in the average individual in control of a resource allocation trait, and of the resident allocation schedule $\mathbf{v}$ (by way of eq. S1). However, since the mutant allele is considered to be rare for the invasion analysis and the population size is large, then the fitness function $w_{s's}(\mathbf{u}^s, \mathbf{v})$ is independent of the number (or frequency) of mutants in the population.

For calculating the fitness functions, we only need to specify the number of individuals alive at the end of the season $t = T$. To that end it is useful to set

$$y_k(T) = y_k(\mathbf{v}) \quad \text{and} \quad x_k^s(T) = x_k(\mathbf{u}^s), \tag{S5}$$

which gives the number of individuals of type $k \in \{\mathrm{w}, \mathrm{q}, \mathrm{m}, \mathrm{iq}\}$ at the end of the season $t = T$ associated with a resident colony and a colony founded by a mutant individual of type $s$, respectively (by way of eqs. 1–2 and S3–S4). Note that type $k \in \{\mathrm{w}, \mathrm{q}, \mathrm{m}\}$ individuals are individuals produced in a focal colony and type $k = \mathrm{iq}$ individuals are females inseminated by sons produced in the focal colony. In eq. (S5) we have emphasized the functional dependence of the number of individuals at the end of the season on the allocation schedules, $\mathbf{u}^s$ and $\mathbf{v}$ (recall eqs. S2 and 3).

Next, we derive the fitness functions $w_{s's}(\mathbf{u}^s, \mathbf{v})$. A colony-founding female is expected to have $x_\mathrm{q}(\mathbf{u}^\mathrm{q})$ surviving daughters (juvenile queens) at the end of the breeding season and her sons are expected to have inseminated $Mx_\mathrm{q}(\mathbf{u}^\mathrm{q})$ surviving females at the end of the breeding season. The probability that a daughter or a female inseminated by a son will gain any one of the $n$ breeding spots is $n/ny_\mathrm{q}(\mathbf{v})$, since there are total number of $ny_\mathrm{q}(\mathbf{v})$ juvenile queens competing for these spots. Hence, the number of mutant colony-founding individuals of type $s'$ in the next generation that descend from a mutant colony-founding female can be written as

$$w_{\mathrm{qq}}(\mathbf{u}^{\mathrm{q}},\mathbf{v}) = \frac{x_{\mathrm{q}}(\mathbf{u}^{\mathrm{q}})}{y_{\mathrm{q}}(\mathbf{v})}, \qquad w_{\mathrm{mq}}(\mathbf{u}^{\mathrm{q}},\mathbf{v}) = \frac{Mx_{\mathrm{iq}}(\mathbf{u}^{\mathrm{q}})}{y_{\mathrm{q}}(\mathbf{v})}. \tag{S6}$$

Since females mate with $M$ males, each colony-founding male is expected to only father $1/M$ of the offspring (here we have not yet taken into account the transmission frequencies specific to different genetic systems). Hence, a colony-founding male is expected to father $x_{\mathrm{q}}(\mathbf{u}^{\mathrm{m}})/M$ surviving daughters at the end of the breeding season and his sons are expected to have inseminated $Mx_{\mathrm{iq}}(\mathbf{u}^{\mathrm{m}})$ surviving females at the end of the breeding season. Here we formulate the fitness functions for any genetic system and by "sons" we mean males produced by a queen that the male has mated with. Note that, in haplodiploids, males do not pass their genes to male offspring. The probability that the daughters and the females inseminated by sons will gain any one of the $n$ breeding spots is $n/ny_{\mathrm{q}}(\mathbf{v})$. Hence, the number of colony-founding individuals of type $s'$ in the next generation that descend from a colony-founding male can be written as

$$w_{\mathrm{qm}}(\mathbf{u}^{\mathrm{m}},\mathbf{v}) = \frac{1}{M}\frac{x_{\mathrm{q}}(\mathbf{u}^{\mathrm{m}})}{y_{\mathrm{q}}(\mathbf{v})}, \qquad w_{\mathrm{mm}}(\mathbf{u}^{\mathrm{m}},\mathbf{v}) = \frac{1}{M}\frac{Mx_{\mathrm{iq}}(\mathbf{u}^{\mathrm{m}})}{y_{\mathrm{q}}(\mathbf{v})}. \tag{S7}$$

The number of individuals at the end of the season $y_{\mathrm{q}}(\mathbf{v})$, $x_{\mathrm{q}}(\mathbf{u}^{s})$ and $x_{\mathrm{iq}}(\mathbf{u}^{s})$ ($s \in \{\mathrm{q},\mathrm{m}\}$) in eqs. (S6)–(S7) are determined from eq. (1) (with eq. 3) and eqs. (S3)–(S4) (with eqs. S1 and S2), respectively.

#### 1.3 The invasion fitness

We now have all the elements to obtain an expression for the invasion fitness, which allows to ascertain the fate of the mutant allele. Let us denote by $n_{\mathrm{q,u}}$ (and respectively, by $n_{\mathrm{m,u}}$) the number of mutant allele copies in females (males with whom the females have mated with), measured at time $t = T$ in the population. The change in the vector $\mathbf{n}_{\mathrm{u}} = (n_{\mathrm{q,u}}, n_{\mathrm{m,u}})^{\intercal}$ of number of gene copies from one generation to the next $\mathbf{n}'_{\mathrm{u}} = (n'_{\mathrm{q,u}}, n'_{\mathrm{m,u}})^{\intercal}$, when the mutant allele for a trait that is under genetic control of party $c \in \{\mathrm{q},\mathrm{w}\}$ is still rare in the population, is given by the matrix

$$\mathbf{A}_c(\mathbf{u}^{\mathrm{q}}(\mathbf{u}),\mathbf{u}^{\mathrm{m}}(\mathbf{u}),\mathbf{v}) = \mathbf{A}_c(\mathbf{u},\mathbf{v}) = \begin{bmatrix} \gamma_{\mathrm{qq}}w_{\mathrm{qq}}(\mathbf{u}^{\mathrm{q}},\mathbf{v}) & \gamma_{\mathrm{qm}}w_{\mathrm{qm}}(\mathbf{u}^{\mathrm{m}},\mathbf{v}) \\ \gamma_{\mathrm{mq}}w_{\mathrm{mq}}(\mathbf{u}^{\mathrm{q}},\mathbf{v}) & \gamma_{\mathrm{mm}}w_{\mathrm{mm}}(\mathbf{u}^{\mathrm{m}},\mathbf{v}) \end{bmatrix} \tag{S8}$$

where $\gamma_{s's}$ is the probability that a gene sampled in an individual of type $s' \in \{\mathrm{q},\mathrm{m}\}$ was contributed by an individual of type $s \in \{\mathrm{q},\mathrm{m}\}$, i.e. a transmission frequency of type $s$ to type $s'$ (for haplodiploids $\gamma_{\mathrm{qq}} = 1/2$, $\gamma_{\mathrm{qm}} = 1/2$, $\gamma_{\mathrm{mq}} = 1$, $\gamma_{\mathrm{mm}} = 0$). Hence, elements $a_{s's}$ of matrix $\mathbf{A}_c(\mathbf{u},\mathbf{v})$ give the expected number of mutant gene copies in a type $s' \in \{\mathrm{q},\mathrm{m}\}$ individual that descends from an individual of type $s \in \{\mathrm{q},\mathrm{m}\}$ carrying the mutant allele. Note that in eq. (S8), the dependence on the party $c \in \{\mathrm{q},\mathrm{w}\}$ who has the genetic control enters into the right-hand-side implicitly via the mutant schedules $\mathbf{u}^{\mathrm{q}}$ and $\mathbf{u}^{\mathrm{m}}$ (recall eq. S1).

The invasion fitness $W_c(\mathbf{u},\mathbf{v})$ of the mutant allele is then given by the leading eigenvalue of the matrix $\mathbf{A}_c(\mathbf{u},\mathbf{v})$ (eq. S8), where the subscript $c \in \{\mathrm{q},\mathrm{w}\}$ emphasizes the party in control of the focal trait. Hence, it satisfies

$$W_c(\mathbf{u},\mathbf{v})\mathbf{q}(\mathbf{u},\mathbf{v}) = \mathbf{A}_c(\mathbf{u},\mathbf{v})\mathbf{q}(\mathbf{u},\mathbf{v}), \tag{S9}$$

where $\mathbf{q}(\mathbf{u},\mathbf{v}) = (q_{\mathrm{q}}(\mathbf{u},\mathbf{v}), q_{\mathrm{m}}(\mathbf{u},\mathbf{v}))^{\intercal}$ is the normalized right leading eigenvector of $\mathbf{A}_c(\mathbf{u},\mathbf{v})$. Here, normalization means that $q_{\mathrm{q}}(\mathbf{u},\mathbf{v}) + q_{\mathrm{m}}(\mathbf{u},\mathbf{v}) = 1$. Pre-multiplying eq. (S9) by the vector $(1,1)$ yields

$$W_c(\mathbf{u},\mathbf{v}) = \left[\gamma_{\mathrm{qq}} w_{\mathrm{qq}}(\mathbf{u}^{\mathrm{q}},\mathbf{v}) + \gamma_{\mathrm{mq}} w_{\mathrm{mq}}(\mathbf{u}^{\mathrm{q}},\mathbf{v})\right] q_{\mathrm{q}}(\mathbf{u},\mathbf{v}) + \left[\gamma_{\mathrm{qm}} w_{\mathrm{qm}}(\mathbf{u}^{\mathrm{m}},\mathbf{v}) + \gamma_{\mathrm{mm}} w_{\mathrm{mm}}(\mathbf{u}^{\mathrm{m}},\mathbf{v})\right] q_{\mathrm{m}}(\mathbf{u},\mathbf{v}), \tag{S10}$$

since $q_{\mathrm{q}}(\mathbf{u},\mathbf{v}) = (1 - q_{\mathrm{m}}(\mathbf{u},\mathbf{v}))$ (see Lehmann et al., 2016, Appendices A-C for more details of how to express invasion fitness in terms of leading left and right eigenvectors of the transition matrix). Note that in eq. (S10), the dependence on the party $c \in \{\mathrm{q}, \mathrm{w}\}$ who has the genetic control, enters into the right-hand-side implicitly via the mutant schedules $\mathbf{u}^{\mathrm{q}}$ and $\mathbf{u}^{\mathrm{m}}$ (recall eq. S1).

The invasion fitness can be interpreted here as the geometric growth rate (generational growth rate) of the mutant allele. This is the asymptotic per capita number of mutant copies produced by the mutant lineage descending from the initial mutation, when overall still rare in the population (see Lehmann et al., 2016 for more details and connections to different fitness measures used in evolutionary biology).

Direct calculation of the normalized right eigenvectors yields

$$\begin{aligned} q_{\mathrm{q}}(\mathbf{u},\mathbf{v}) &= \frac{2\gamma_{\mathrm{qm}} w_{\mathrm{qm}}(\mathbf{u}^{\mathrm{q}},\mathbf{v}) - X(\mathbf{u},\mathbf{v}) + \sqrt{(X(\mathbf{u},\mathbf{v}))^2 + 4\gamma_{\mathrm{qm}}\gamma_{\mathrm{mq}} w_{\mathrm{qm}}(\mathbf{u}^{\mathrm{m}},\mathbf{v})\, w_{\mathrm{mq}}(\mathbf{u}^{\mathrm{q}},\mathbf{v})}}{2\left(\gamma_{\mathrm{qm}} w_{\mathrm{qm}}(\mathbf{u}^{\mathrm{q}},\mathbf{v}) - \gamma_{\mathrm{mq}} w_{\mathrm{mq}}(\mathbf{u}^{\mathrm{q}},\mathbf{v}) - X(\mathbf{u},\mathbf{v})\right)}, \\ q_{\mathrm{m}}(\mathbf{u},\mathbf{v}) &= \frac{2\gamma_{\mathrm{mq}} w_{\mathrm{mq}}(\mathbf{u}^{\mathrm{q}},\mathbf{v})}{2\gamma_{\mathrm{mq}} w_{\mathrm{mq}}(\mathbf{u}^{\mathrm{q}},\mathbf{v}) + X(\mathbf{u},\mathbf{v}) + \sqrt{(X(\mathbf{u},\mathbf{v}))^2 + 4\gamma_{\mathrm{qm}}\gamma_{\mathrm{mq}} w_{\mathrm{qm}}(\mathbf{u}^{\mathrm{m}},\mathbf{v})\, w_{\mathrm{mq}}(\mathbf{u}^{\mathrm{q}},\mathbf{v})}}, \end{aligned} \tag{S11}$$

where $X(\mathbf{u},\mathbf{v}) = \gamma_{\mathrm{qq}} w_{\mathrm{qq}}(\mathbf{u}^{\mathrm{q}},\mathbf{v}) - \gamma_{\mathrm{mm}} w_{\mathrm{mm}}(\mathbf{u}^{\mathrm{m}},\mathbf{v})$.

The quantity $q_s(\mathbf{u},\mathbf{v})$ can be interpreted as the asymptotic probability that a mutant allele is sampled in a class $s$ individual. It follows that the maximization of the invasion fitness (S10) depends on both the fitnesses of carriers of the mutant allele (the $w_{s's}(\mathbf{u}^s,\mathbf{v})$ functions) and how the mutant allele is distributed across classes (the $q_s(\mathbf{u},\mathbf{v})$ functions which also depend on the evolving traits themselves).

##### 1.4 Uninvadable allocation schedule

An uninvadable schedule $\mathbf{u}^* = \{u_{\mathrm{f}}^*(t), u_{\mathrm{q}}^*(t)\}_{t\in[0,T]}$ is a resident schedule that is resistant to invasion by any mutant $\mathbf{u} \in \mathbb{U} = \mathbb{U}_{\mathrm{f}} \times \mathbb{U}_{\mathrm{q}}$ schedule. Here, $\mathbb{U}$ is a set of all possible allocation schedules, while $\mathbb{U}_{\mathrm{f}}$ and $\mathbb{U}_{\mathrm{q}}$ are sets of full trajectories of the traits $\mathbf{u}_{\mathrm{f}} = \{u_{\mathrm{f}}(t)\}_{t\in[0,T]}$ and $\mathbf{u}_{\mathrm{q}} = \{u_{\mathrm{q}}(t)\}_{t\in[0,T]}$ under consideration. Notice that in order to simplify notations in the main text we used $\mathbf{v}^* \equiv \mathbf{u}^*$, $v_{\mathrm{f}}^*(t) \equiv u_{\mathrm{f}}^*(t)$, and $v_{\mathrm{q}}^*(t) \equiv u_{\mathrm{q}}^*(t)$ for the uninvadable schedule, but in this S.I. it is more convenient to use the letter $u$ for that, basically throughout the S.I. we always distinguish mutant and resident, both at the level of state variables ($x$ vs. $y$) and at the level of evolving traits ($u$ vs. $v$).

If party $c \in \{\mathrm{q}, \mathrm{w}\}$ is in full control of the two traits (i.e. single-party control), then the uninvadable schedule

$\mathbf{u}^*$ satisfies the condition

$$\mathbf{u}^* \in \arg\max_{\mathbf{u}\in\mathbb{U}} W_c(\mathbf{u}, \mathbf{u}^*), \tag{S12}$$

that is, a mutant that adopts the resident schedule $\mathbf{u}^*$ has the highest invasion fitness from all possible strategies in a population, for a population expressing schedule $\mathbf{u}^*$. Hence, an uninvadable schedule $\mathbf{u}^*$ is a candidate endpoint of the evolutionary process.

Under mixed control, where the queen is in control of the trait $u_{\mathrm{f}}$ and the workers are in control of the trait $u_{\mathrm{q}}$, the uninvadable schedule $\mathbf{u}^*$ satisfies condition

$$\mathbf{u}_{\mathrm{f}}^* \in \arg\max_{\mathbf{u}_{\mathrm{f}}\in\mathbb{U}_{\mathrm{f}}} W_{\mathrm{q}}\big((\mathbf{u}_{\mathrm{f}}, \mathbf{u}_{\mathrm{q}}^*), \mathbf{u}^*\big) \quad \text{and} \quad \mathbf{u}_{\mathrm{q}}^* \in \arg\max_{\mathbf{u}_{\mathrm{q}}\in\mathbb{U}_{\mathrm{q}}} W_{\mathrm{w}}((\mathbf{u}_{\mathrm{f}}^*, \mathbf{u}_{\mathrm{q}}), \mathbf{u}^*)\,. \tag{S13}$$

Hence, the uninvadable allocation schedules to individuals of type $k$ ($k \in \{\mathrm{w}, \mathrm{q}, \mathrm{m}\}$) can be written as follows

$$a_{\mathrm{w}}^*(t) = u_{\mathrm{f}}^*(t)(1 - u_{\mathrm{q}}^*(t)), \qquad a_{\mathrm{q}}^*(t) = u_{\mathrm{f}}^*(t)u_{\mathrm{q}}^*(t), \qquad a_{\mathrm{m}}^*(t) = (1 - u_{\mathrm{f}}^*(t)) \tag{S14}$$

and we denote by $x_k^*(t)$ the resulting number of individuals at time $t$.

### 2 First-order condition

In this section, we derive the first-order necessary condition for uninvadability (eq. 5 of the main text) and show that it has a similar structure that first-order conditions in static allocation models (e.g. eq. (1)–(2) in Reuter and Keller, 2001) and applies regardless of colony growth dynnamic. In the next section, we the show how to formulate the necessary first-order condition in terms of pointwise marginal change using optimal control theory and then solve explicitly for the (candidate) uninvadable schedule for our model.

#### 2.1 Eigenvalue perturbation

##### 2.1.1 Perturbations in terms of Gâteaux derivatives and relatedness assymetry

We consider a small variation $\epsilon\eta_\tau$ ($\eta_\tau = \{\eta_\tau(t)\}_{t=0}^{T}$) in the trait $u_\tau^*(t)$ $\tau \in \{\mathrm{f}, \mathrm{q}\}$ of the uninvadable schedule, such that the mutant trait can be written as

$$u_\tau(t) = u_\tau^*(t) + \epsilon_\tau\eta_\tau(t) \qquad \forall t \in [0, T], \tag{S15}$$

for any feasible deviation $\eta_\tau(t)$ (such that $0 \le u_\tau(t) \le 1$) from the resident schedule $u_\tau^*(t)$, where $\epsilon_\tau \ll 1$ is a small parameter measuring the intensity of the mutant deviation. Hence, we consider a change in the candidate uninvadable allocation trait $u_\tau^*(t)$ that remains very close to it for all $t \in [0, T]$. The direction of selection for trait $v_\tau(t) = u_\tau^*(t)$ is indicated by the sign of perturbation in invasion fitness

$$\left.\frac{\mathrm{d}W_c(\mathbf{u}, \mathbf{u}^*)}{\mathrm{d}\epsilon_\tau}\right|_{\epsilon_{\mathrm{f}}=0, \epsilon_{\mathrm{q}}=0} \qquad \text{for } \tau = \mathrm{f} \text{ and } \tau = \mathrm{q}, \tag{S16}$$

for any feasible deviation $\eta_\tau(t)$ from the uninvadable schedule $\mathbf{u}^*$. The derivative in eq. (S16) is a Gâteaux derivative (a type of functional or variational derivative) of invasion fitness (e.g., Weber and Arfken, 2003, p. 827–830, Troutman, 2012, p. 45–50, Luenberger, 1997, p. 171–178, Gelfand and Fomin, 1963, p. 54–63). In other words, it gives the infinitesimal change in invasion fitness resulting from a change in the whole mutant schedule into the direction of $\eta(t)$ (Gâteaux derivative can be thought of as a generalization of directional derivative from differential calculus). Gâteaux derivatives are useful to generalize evolutionary stability conditions (e.g., Eshel, 1983, eq. 3, Taylor, 1989, eq. 2.1) to function-valued traits.

Because the functional derivative, $\mathrm{d}W_c(\mathbf{u},\mathbf{u}^*)/\,\mathrm{d}\epsilon_\tau$ is an ordinary function in $\epsilon_\tau$, it follows from standard results of eigenvalue perturbation (Caswell, 2001, p. 209, eq. 9.10) that

$$\frac{\mathrm{d}W_c(\mathbf{u},\mathbf{u}^*)}{\mathrm{d}\epsilon_\tau} = \frac{\boldsymbol{\nu}^\circ \frac{\mathrm{d}\mathbf{A}_c(\mathbf{u},\mathbf{u}^*)}{\mathrm{d}\epsilon_\tau}(\mathbf{q}^\circ)^\intercal}{\boldsymbol{\nu}^\circ(\mathbf{q}^\circ)^\intercal}, \tag{S17}$$

where superscript $\intercal$ denotes transpose, $\boldsymbol{\nu}^\circ = (\nu^\circ_\mathrm{q}, \nu^\circ_\mathrm{m})$ is a vector of neutral reproductive values of colony-founding individuals of type $s \in \{\mathrm{q},\mathrm{m}\}$ and $\mathbf{q}^\circ = (q^\circ_\mathrm{q}, q^\circ_\mathrm{m})$ is a vector of the neutral frequencies of class $s \in \{\mathrm{q},\mathrm{m}\}$ individuals. Throughout, the superscript $\circ$ will denote a quantity that is evaluated in the absence of natural selection, i.e., by a process determined by the monomorphic resident population. Substituting eq. (S8) into (S17) and given that $\boldsymbol{\nu}^\circ(\mathbf{q}^\circ)^\intercal = 1$ (total class reproductive values of all individuals add up to one) yields

$$\begin{aligned}\frac{\mathrm{d}W_c(\mathbf{u},\mathbf{u}^*)}{\mathrm{d}\epsilon_\tau} &= \left(\nu^\circ_\mathrm{q}\gamma_\mathrm{qq}\frac{\mathrm{d}w_\mathrm{qq}(\mathbf{u}^\mathrm{q},\mathbf{u}^*)}{\mathrm{d}\epsilon_\tau} + \nu^\circ_\mathrm{m}\gamma_\mathrm{mq}\frac{\mathrm{d}w_\mathrm{mq}(\mathbf{u}^\mathrm{q},\mathbf{u}^*)}{\mathrm{d}\epsilon_\tau}\right) q^\circ_\mathrm{q} \\ &\quad + \left(\nu^\circ_\mathrm{q}\gamma_\mathrm{qm}\frac{\mathrm{d}w_\mathrm{qm}(\mathbf{u}^\mathrm{m},\mathbf{u}^*)}{\mathrm{d}\epsilon_\tau} + \nu^\circ_\mathrm{m}\gamma_\mathrm{mm}\frac{\mathrm{d}w_\mathrm{mm}(\mathbf{u}^\mathrm{m},\mathbf{u}^*)}{\mathrm{d}\epsilon_\tau}\right) q^\circ_\mathrm{m},\end{aligned} \tag{S18}$$

where all derivatives are evaluated at $\epsilon_\tau = 0$.

In the absence of natural selection, the number of gene copies from one generation to the next can be described by a matrix

$$\mathbf{A}_c(\mathbf{v},\mathbf{v}) = \mathbf{A}(\mathbf{v},\mathbf{v}) = \begin{bmatrix} \gamma_\mathrm{qq} & \gamma_\mathrm{qm}\frac{1}{M} \\ \gamma_\mathrm{mq}M & \gamma_\mathrm{mm}\end{bmatrix}, \tag{S19}$$

which does not depend on the mode of control and whose dominant eigenvalue is one (given that $\gamma_\mathrm{qq} = 1/2$, $\gamma_\mathrm{qm} = 1/2$, $\gamma_\mathrm{mq} = 1$, $\gamma_\mathrm{mm} = 0$). The reproductive values $\boldsymbol{\nu}^\circ$ and class frequencies $\mathbf{q}^\circ$ are, respectively, given by the left and right unit eigenvectors of $\mathbf{A}(\mathbf{v},\mathbf{v})$, and we normalize these vectors such that the total class reproductive values defined by

$$\alpha^\circ_s = \nu^\circ_s q^\circ_s \tag{S20}$$

(e.g., Rousset, 2004; Taylor, 1990; Taylor and Frank, 1996) of all individuals add up to one: $\alpha^\circ_\mathrm{q} + \alpha^\circ_\mathrm{m} = 1$. This normalization entails the use of the perturbation formula eq. (S18) (e.g., Caswell, 2001), with which we obtain

$$\begin{aligned} \alpha_{\mathrm{q}}^{\circ} &= \frac{1-\gamma_{\mathrm{mm}}}{2-\gamma_{\mathrm{qq}}-\gamma_{\mathrm{mm}}} \qquad \text{and} \quad \alpha_{\mathrm{m}}^{\circ} = \frac{1-\gamma_{\mathrm{qq}}}{2-\gamma_{\mathrm{qq}}-\gamma_{\mathrm{mm}}}, \\ q_{\mathrm{q}}^{\circ} &= \frac{1}{1+M} \qquad \text{and} \quad q_{\mathrm{m}}^{\circ} = \frac{M}{1+M}. \end{aligned} \tag{S21}$$

It follows from the class frequencies $q_s^{\circ}$, that under neutrality there are $M$ times as much colony-founding males than females, which is in accordance with the fact that females mate $M$ times.

Substituting the transmission frequencies $\gamma_{s's}$ for haplodiploids [$\gamma_{\mathrm{qq}} = 1/2$, $\gamma_{\mathrm{mq}} = 1$, $\gamma_{\mathrm{qm}} = 1/2$, $\gamma_{\mathrm{mm}} = 0$] then we have the class reproductive values for haplodiploids

$$\alpha_{\mathrm{q}}^{\circ} = \frac{2}{3} \quad \text{and} \quad \alpha_{\mathrm{m}}^{\circ} = \frac{1}{3}. \tag{S22}$$

In eq. (S18), the derivative $\mathrm{d}w_{s's}(\mathbf{u}^s, \mathbf{v}) / \mathrm{d}\epsilon_\tau$ is the total variation of individual fitness with respect to mutant values, which acts on $\mathbf{u}^s$ (by way of eq. S1). By substituting eq. (S15) into eq. (S1) (where we take $v_\tau = u_\tau^*$), we have for $\tau \in \{\mathrm{f}, \mathrm{q}\}$ that

$$u_\tau^s(t) = u_\tau^*(t) + \epsilon_\tau \eta_\tau(t) p_c^s \tag{S23}$$

and owing to eq. (S15) and the constant factor rule in differentiation, we can write

$$\left.\frac{\mathrm{d}w_{s's}(\mathbf{u}^s, \mathbf{u}^*)}{\mathrm{d}\epsilon_\tau}\right|_{\epsilon_\tau = 0} = \left.\frac{\mathrm{d}w_{s's}(\mathbf{u}, \mathbf{u}^*)}{\mathrm{d}\epsilon_\tau}\right|_{\epsilon_\tau = 0} \times p_c^s. \tag{S24}$$

Substituting eq. (S24) into eq. (S18), we have for control mode $c \in \{\mathrm{q}, \mathrm{w}\}$ that

$$\begin{aligned} \frac{\mathrm{d}W_c(\mathbf{u}, \mathbf{u}^*)}{\mathrm{d}\epsilon_\tau} = &\left(\nu_{\mathrm{q}}^{\circ}\gamma_{\mathrm{qq}}\frac{\mathrm{d}w_{\mathrm{qq}}(\mathbf{u}, \mathbf{u}^*)}{\mathrm{d}\epsilon_\tau} + \nu_{\mathrm{m}}^{\circ}\gamma_{\mathrm{mq}}\frac{\mathrm{d}w_{\mathrm{mq}}(\mathbf{u}, \mathbf{u}^*)}{\mathrm{d}\epsilon_\tau}\right) p_c^{\mathrm{q}} q_{\mathrm{q}}^{\circ} \\ &+ \left(\nu_{\mathrm{q}}^{\circ}\gamma_{\mathrm{qm}}\frac{\mathrm{d}w_{\mathrm{qm}}(\mathbf{u}, \mathbf{u}^*)}{\mathrm{d}\epsilon_\tau} + \nu_{\mathrm{m}}^{\circ}\gamma_{\mathrm{mm}}\frac{\mathrm{d}w_{\mathrm{mm}}(\mathbf{u}, \mathbf{u}^*)}{\mathrm{d}\epsilon_\tau}\right) p_c^{\mathrm{m}} q_{\mathrm{m}}^{\circ}, \end{aligned} \tag{S25}$$

where all derivatives are evaluated at $\epsilon_\tau = 0$ and thus all trait values (allocation schedules) are set to the resident schedule $\mathbf{v}$. Substituting eq. (S20) into eq. (S25) yields

$$\begin{aligned} \frac{\mathrm{d}W_c(\mathbf{u}, \mathbf{u}^*)}{\mathrm{d}\epsilon_\tau} = &\,\alpha_{\mathrm{q}}^{\circ}\left(\gamma_{\mathrm{qq}} p_c^{\mathrm{q}} \frac{\mathrm{d}w_{\mathrm{qq}}(\mathbf{u}, \mathbf{u}^*)}{\mathrm{d}\epsilon_\tau} + \frac{\nu_{\mathrm{m}}^{\circ}}{\nu_{\mathrm{q}}^{\circ}}\gamma_{\mathrm{qm}} p_c^{\mathrm{m}} \frac{\mathrm{d}w_{\mathrm{qm}}(\mathbf{u}, \mathbf{u}^*)}{\mathrm{d}\epsilon_\tau}\right) \\ &+ \alpha_{\mathrm{m}}^{\circ}\left(\frac{\nu_{\mathrm{q}}^{\circ}}{\nu_{\mathrm{m}}^{\circ}}\gamma_{\mathrm{mq}} p_c^{\mathrm{q}} \frac{\mathrm{d}w_{\mathrm{mq}}(\mathbf{u}, \mathbf{u}^*)}{\mathrm{d}\epsilon_\tau} + \gamma_{\mathrm{mm}} p_c^{\mathrm{m}} \frac{\mathrm{d}w_{\mathrm{mm}}(\mathbf{u}, \mathbf{u}^*)}{\mathrm{d}\epsilon_\tau}\right). \end{aligned} \tag{S26}$$

Substituting the fitness functions (S6)–(S7) into eq. (S26) yields

$$\frac{\mathrm{d}W_c(\mathbf{u},\mathbf{u}^*)}{\mathrm{d}\epsilon_\tau} = \alpha_{\mathrm{q}}^{\circ}\left(\gamma_{\mathrm{qq}}p_c^{\mathrm{q}}\frac{\mathrm{d}x_{\mathrm{q}}(\mathbf{u})}{\mathrm{d}\epsilon_\tau} + \frac{\nu_{\mathrm{m}}^{\circ}}{\nu_{\mathrm{q}}^{\circ}}\gamma_{\mathrm{qm}}p_c^{\mathrm{m}}\frac{1}{M}\frac{\mathrm{d}x_{\mathrm{q}}(\mathbf{u})}{\mathrm{d}\epsilon_\tau}\right)+\alpha_{\mathrm{m}}^{\circ}\left(\frac{\nu_{\mathrm{q}}^{\circ}}{\nu_{\mathrm{m}}^{\circ}}\gamma_{\mathrm{mq}}p_c^{\mathrm{q}}M\frac{\mathrm{d}x_{\mathrm{iq}}(\mathbf{u})}{\mathrm{d}\epsilon_\tau} + \gamma_{\mathrm{mm}}p_c^{\mathrm{m}}\frac{\mathrm{d}x_{\mathrm{iq}}(\mathbf{u})}{\mathrm{d}\epsilon_\tau}\right). \tag{S27}$$

By considering that eqs. (S20) and (S21) yield that $\nu_{\mathrm{m}}^{\circ}/\nu_{\mathrm{q}}^{\circ} = M$ and $\nu_{\mathrm{q}}^{\circ}/\nu_{\mathrm{m}}^{\circ} = 1/M$, then eq. (S27) as

$$\frac{\mathrm{d}W_c(\mathbf{u},\mathbf{u}^*)}{\mathrm{d}\epsilon_\tau} = \alpha_{\mathrm{q}}^{\circ}\frac{\mathrm{d}x_{\mathrm{q}}(\mathbf{u})}{\mathrm{d}\epsilon_\tau}\left(\gamma_{\mathrm{qq}}p_c^{\mathrm{q}} + \gamma_{\mathrm{qm}}p_c^{\mathrm{m}}\right) + \alpha_{\mathrm{m}}^{\circ}\frac{\mathrm{d}x_{\mathrm{iq}}(\mathbf{u})}{\mathrm{d}\epsilon_\tau}\left(\gamma_{\mathrm{mq}}p_c^{\mathrm{q}} + \gamma_{\mathrm{mm}}p_c^{\mathrm{m}}\right). \tag{S28}$$

Rearranging, we can write eq. (S28) as

$$\frac{\mathrm{d}W_c(\mathbf{u},\mathbf{u}^*)}{\mathrm{d}\epsilon_\tau} = K\left[R_c\frac{\mathrm{d}x_{\mathrm{q}}(\mathbf{u})}{\mathrm{d}\epsilon_\tau} + \frac{\mathrm{d}x_{\mathrm{iq}}(\mathbf{u})}{\mathrm{d}\epsilon_\tau}\right]. \tag{S29}$$

where $K = \alpha_{\mathrm{m}}^{\circ}(\gamma_{\mathrm{mq}}p_c^{\mathrm{q}} + \gamma_{\mathrm{mm}}p_c^{\mathrm{m}}) > 0$ is a positive constant and

$$R_c = \frac{\alpha_{\mathrm{q}}^{\circ}}{\alpha_{\mathrm{m}}^{\circ}} \times \left(\frac{\gamma_{\mathrm{qq}}p_c^{\mathrm{q}} + \gamma_{\mathrm{qm}}p_c^{\mathrm{m}}}{\gamma_{\mathrm{mq}}p_c^{\mathrm{q}} + \gamma_{\mathrm{mm}}p_c^{\mathrm{m}}}\right) \tag{S30}$$

is the so-called relatedness asymmetry (see Boomsma and Grafen, 1991, p. 386 and section 2.1.2 for the biological interpretation).

##### 2.1.2 Interpretation of relatedness asymmetry

The *relatedness asymmetry* $R_c$ gives the ratio of sex-specific potentials for the party $c$ in control to contribute genes into the distant future (Boomsma and Grafen, 1991, p. 386). In order to see this, we note that owing to eq. (S21), the first ratio in eq. (S30) is the ratio $\alpha_{\mathrm{q}}^{\circ}/\alpha_{\mathrm{m}}^{\circ}$ of class reproductive values. Furthermore, notice that $(\gamma_{\mathrm{qq}}p_c^{\mathrm{q}} + \gamma_{\mathrm{qm}}p_c^{\mathrm{m}}) = \Phi_{\mathrm{q},c}$ and $(\gamma_{\mathrm{mq}}p_c^{\mathrm{q}} + \gamma_{\mathrm{mm}}p_c^{\mathrm{m}}) = \Phi_{\mathrm{m},c}$ are the probabilities that a gene randomly sampled in a recipient female and male, respectively, is identical-by-descent to a gene randomly sampled from party $p$ in control of resource allocation; that is, the *coancestry* (or *consanguinity*) between a female (male) and the (average) individual whose genes are in control of the resource allocation trait. Hence, the relatedness asymmetry is

$$R_c = \frac{v_{\mathrm{q}}^{\circ}q_{\mathrm{q}}^{\circ}\Phi_{\mathrm{q},c}}{v_{\mathrm{m}}^{\circ}q_{\mathrm{m}}^{\circ}\Phi_{\mathrm{m},c}}. \tag{S31}$$

Here, $q_s^{\circ}\Phi_{s,c}$ is the asymptotic probability that a randomly sampled gene from a colony-founding individual finds itself in an individual of type $s$ and is a replica copy of a gene sampled from a party $c$. Then, since $\nu_s^{\circ}$ is the long-term contribution of genes in individual of type $s$ to the gene pool, we can interpret the relatedness asymmetry as giving the ratio of sex-specific potentials for party $p$ in control to contribute (in a neutral process) to the gene pool in the distant future.

Since the ratio of consanguinity is equivalent to the ratio of relatedness, we can write the second ratio in eq. (S30) as $r^\circ_{\mathrm{q},c}/r^\circ_{\mathrm{m},c}$, where $r^\circ_{s,c} = \Phi_{s,c}/\Phi_c$ is the relatedness between an individual of type $s$ and the average individual whose genes are in control of the resource allocation trait, and this depends on the coefficient of coancestry $\Phi_c$ of the average individual in control of the resource allocation trait with itself (i.e. the probability that two homologous genes, drawn randomly with replacement from party $c$, are identical by descent). With this eq. (S31) is also

$$R_c = \frac{\nu^\circ_{\mathrm{q}} q^\circ_{\mathrm{q}} r^\circ_{\mathrm{q},c}}{\nu^\circ_{\mathrm{m}} q^\circ_{\mathrm{m}} r^\circ_{\mathrm{m},c}} = \frac{\alpha^\circ_{\mathrm{q}} r^\circ_{\mathrm{q},c}}{\alpha^\circ_{\mathrm{m}} r^\circ_{\mathrm{m},c}}, \tag{S32}$$

where the second equality displays the classical form of the relatedness asymmetry (Boomsma and Grafen, 1991, p. 386). For haplodiploids eq. (S32) simplifies to

$$R_c = 2\frac{r^\circ_{\mathrm{q},c}}{r^\circ_{\mathrm{m},c}}. \tag{S33}$$

Since relatedness if given by the ratio of the coefficient of coancestry of party $c$ with an individual of type $s$ ($\Phi_{s,c}$, which is given by the transmission frequencies $\gamma_{s's}$ and the expected frequency $p^s_c$ of mutant allele residing in party $c$) to the coefficient of coancestry $\Phi_c$ of party $c$ with itself ($\Phi_{\mathrm{q}}$=$\Phi_{\mathrm{w}}$=1/2). Substituting the frequencies for haplodiploids [$\gamma_{\mathrm{qq}} = 1/2$, $\gamma_{\mathrm{mq}} = 1$, $\gamma_{\mathrm{qm}} = 1/2$, $\gamma_{\mathrm{mm}} = 0$, $p^{\mathrm{q}}_{\mathrm{q}} = 1/2$, $p^{\mathrm{m}}_{\mathrm{q}} = 0$, $p^{\mathrm{q}}_{\mathrm{w}} = 1/4$, and $p^{\mathrm{m}}_{\mathrm{w}} = 1/(2M)$] gives the relatedness coefficients for haplodiploids

$$\begin{aligned} &r^\circ_{\mathrm{q,q}} = \frac{1}{2} \quad \text{and} \quad r^\circ_{\mathrm{m,q}} = 1 \quad (\mathbf{u}_\tau, \mathbf{v}_\tau \text{ under queen control}),\\ &r^\circ_{\mathrm{q,w}} = \frac{2+M}{4M} \quad \text{and} \quad r^\circ_{\mathrm{m,w}} = \frac{1}{2} \quad (\mathbf{u}_\tau, \mathbf{v}_\tau \text{ under worker control}) \end{aligned} \tag{S34}$$

which are classical expressions (e.g., Frank, 1998, Fig. 10.4, p. 209). Substituting the relatedness coefficients into eq. (S33) yields the relatedness asymmetry for haplodiploids

$$R_{\mathrm{q}} = 1 \quad (\mathbf{u}_\tau, \mathbf{v}_\tau \text{ under queen control}) \qquad R_{\mathrm{w}} = \frac{2+M}{M} \quad (\mathbf{u}_\tau, \mathbf{v}_\tau \text{ under worker control}). \tag{S35}$$

#### 2.2 First-order condition for uninvadability

The necessary first-order condition for the candidate uninvdable schedule $\mathbf{u}^* = (\mathbf{u}^*_{\mathrm{f}}, \mathbf{u}^*_{\mathrm{q}})$ is given by

$$\left.\frac{\mathrm{d}W_c(\mathbf{u},\mathbf{u}^*)}{\mathrm{d}\epsilon_\tau}\right|_{\epsilon_{\mathrm{f}}=0,\epsilon_{\mathrm{q}}=0} \leq 0 \qquad \text{for } \tau = \mathrm{f} \text{ and } \tau = \mathrm{q}, \tag{S36}$$

for any feasible deviation $\eta(t)$. Substituting eq. (S29) into (S36) yields that we can express the necessary first-order condition for uninvadability under queen ($c = \mathrm{q}$) or worker ($c = \mathrm{w}$) control as

$$R_c \left.\frac{\mathrm{d}x_{\mathrm{q}}(\mathbf{u})}{\mathrm{d}\epsilon_\tau}\right|_{\epsilon_{\mathrm{f}}=0,\epsilon_{\mathrm{q}}=0} + \left.\frac{\mathrm{d}x_{\mathrm{iq}}(\mathbf{u})}{\mathrm{d}\epsilon_\tau}\right|_{\epsilon_{\mathrm{f}}=0,\epsilon_{\mathrm{q}}=0} \leq 0 \tag{S37}$$

and under mixed control as

$$
\begin{aligned}
&R_c \frac{\mathrm{d}x_{\mathrm{q}}(\mathbf{u})}{\mathrm{d}\epsilon_{\mathrm{f}}}\bigg|_{\epsilon_{\mathrm{f}}=0,\epsilon_{\mathrm{q}}=0} + \frac{\mathrm{d}x_{\mathrm{iq}}(\mathbf{u})}{\mathrm{d}\epsilon_{\mathrm{f}}}\bigg|_{\epsilon_{\mathrm{f}}=0,\epsilon_{\mathrm{q}}=0} \leq 0 \quad \text{and} \\
&R_c \frac{\mathrm{d}x_{\mathrm{q}}(\mathbf{u})}{\mathrm{d}\epsilon_{\mathrm{q}}}\bigg|_{\epsilon_{\mathrm{f}}=0,\epsilon_{\mathrm{q}}=0} + \frac{\mathrm{d}x_{\mathrm{iq}}(\mathbf{u})}{\mathrm{d}\epsilon_{\mathrm{q}}}\bigg|_{\epsilon_{\mathrm{f}}=0,\epsilon_{\mathrm{q}}=0} \leq 0.
\end{aligned}
\tag{S38}
$$

Hence, the first-order condition given by eqs. (S37) and (S38) can be expressed in terms of variational derivatives $\mathrm{d}x_k(\mathbf{u})/\,\mathrm{d}\epsilon_\tau$ and relatedness asymmetry $R_c$. The variational derivative $\mathrm{d}x_k(\mathbf{u})/\,\mathrm{d}\epsilon_\tau$ measures the change in the number of individuals of type $k \in \{\mathrm{q}, \mathrm{iq}\}$ associated with a focal colony where phenotype $\mathbf{u}$ is expressed. In the next section we give the interpretation for relatedness asymmetry. Note that the first-order condition given by eqs. (S37) and (S38) is a dynamic version of first-order condition in a comparable static allocation model (e.g. eq. (1)–(2) in Reuter and Keller, 2001). Note that we the first-order condition (given by eqs. S37 and S38) in the main text (recall 5) using a different notation (to simplify the readability for the general audience), where $y_{\mathrm{q}}(\mathbf{u}) \equiv x_{\mathrm{q}}(\mathbf{u})$ and $y_{\mathrm{iq}}(\mathbf{u}) \equiv x_{\mathrm{iq}}(\mathbf{u})$.

#### 2.3 Pointwise eigenvalue perturbation

It is useful to also consider pointwise perturbations in invasion fitness, which would allow to describe the direction of selection on trait $v_\tau(t) = u_\tau^*(t)$ for each $t$. That is, we consider for every $t \in [0, T]$

$$
\frac{\delta W_c(\mathbf{u}, \mathbf{u}^*)}{\delta \eta_\tau(t)} = \lim_{\eta_\tau(t) \to \delta_t(t')} \frac{\mathrm{d}W_c(\mathbf{u}, \mathbf{u}^*)}{\mathrm{d}\epsilon_\tau}\bigg|_{\epsilon_{\mathrm{f}}=0,\epsilon_{\mathrm{q}}=0}, \tag{S39}
$$

where the derivative on the left-hand-side is a pointwise functional derivative (the so-called Volterra derivative) of invasion fitness at time $t \in [0, T]$ (see e.g. Parr and Yang, 1989, p. 246-247 and eq. (3a) in Dieckmann et al., 2006) and $\delta_t(t') = \delta(t' - t)$ is the Dirac delta function, which is 0, except at $t' = t$, when it is 1 (here, $t'$ is just a dummy variable for time $t' \in [0, T]$) Note that using the $\delta$-notation (not to be confused with the Dirac delta function) to refer to the pointwise functional derivative is standard notation in the physical literature (see e.g. Giaquinta and Hildebrandt, 1996, p. 18).

It follows from eq. (S29) that we can express the pointwise perturbations as follows

$$
\frac{\delta W_c(\mathbf{u}, \mathbf{u}^*)}{\delta \eta_\tau(t)}\bigg|_{\epsilon_{\mathrm{f}}=0,\epsilon_{\mathrm{q}}=0} = \lim_{\eta_\tau(t) \to \delta_t(t')} K\left(R_c \frac{\mathrm{d}x_{\mathrm{q}}(\mathbf{u})}{\mathrm{d}\epsilon_\tau} + \frac{\mathrm{d}x_{\mathrm{iq}}(\mathbf{u})}{\mathrm{d}\epsilon_\tau}\right) = K\left(R_c \frac{\delta x_{\mathrm{q}}(\mathbf{u})}{\delta \eta_\tau(t)} + \frac{\delta x_{\mathrm{iq}}(\mathbf{u})}{\delta \eta_\tau(t)}\right). \tag{S40}
$$

#### 2.4 Pointwise first-order condition for a singular arc and the marginal substitution rate

We call the uninvadable allocation trait $u_\tau^*(t)$ a singular arcs, when it does not reside on the bounds of the feasible set (i.e., when $0 \leq u_\tau^*(t) = \hat{u}_\tau^*(t) \leq 1$) over a finite period of time. Now we will show that the pointwise first-order conditions for singular arcs can be expressed in terms of marginal substitution rates. We will show in section 3 (see eqs. S66 and S73) that at the singular arc

$$\left.\frac{\delta W_c(\mathbf{u},\mathbf{u}^*)}{\delta \eta_\tau(t)}\right|_{\epsilon_\mathrm{f}=0,\epsilon_\mathrm{q}=0} = 0. \tag{S41}$$

Hence, we can express the necessary first-order condition for the singular arc $\hat{u}_\tau^*(t)$ to be uninvadable under queen ($c = \mathrm{q}$) or worker ($c = \mathrm{w}$) as

$$R_c \frac{\delta x_\mathrm{q}(\mathbf{u})}{\delta \eta_\tau(t)} + \frac{\delta x_\mathrm{iq}(\mathbf{u})}{\delta \eta_\tau(t)} = 0 \tag{S42}$$

and under mixed control as

$$\begin{aligned} R_\mathrm{q} \frac{\delta x_\mathrm{q}(\mathbf{u})}{\delta \eta_\mathrm{f}(t)} + \frac{\delta x_\mathrm{iq}(\mathbf{u})}{\delta \eta_\mathrm{f}(t)} &= 0 \quad \text{and} \\ R_\mathrm{w} \frac{\delta x_\mathrm{q}(\mathbf{u})}{\delta \eta_\mathrm{q}(t)} + \frac{\delta x_\mathrm{iq}(\mathbf{u})}{\delta \eta_\mathrm{q}(t)} &= 0. \end{aligned} \tag{S43}$$

Rearranging eqs. (S42) and (S43) yields for queen ($c = \mathrm{q}$) and worker ($c = \mathrm{w}$) control

$$\left.-\frac{\delta x_\mathrm{iq}(\mathbf{u})/\delta \eta_\tau(t)}{\delta x_\mathrm{q}(\mathbf{u})/\delta \eta_\tau(t)}\right|_{\epsilon_\mathrm{f}=0,\epsilon_\mathrm{q}=0} = R_c \qquad \text{for } \tau \in \{\mathrm{f},\mathrm{q}\}, \tag{S44}$$

and for mixed control

$$\left.-\frac{\delta x_\mathrm{iq}(\mathbf{u})/\,\mathrm{d}\eta_\mathrm{f}(t)}{\delta x_\mathrm{q}(\mathbf{u})/\delta \eta_\mathrm{f}(t)}\right|_{\epsilon_\mathrm{f}=0,\epsilon_\mathrm{q}=0} = R_\mathrm{q} \quad \text{and} \quad \left.-\frac{\delta x_\mathrm{iq}(\mathbf{u})/\delta \eta_\mathrm{q}(t)}{\delta x_\mathrm{q}(\mathbf{u})/\delta \eta_\mathrm{q}(t)}\right|_{\epsilon_\mathrm{f}=0,\epsilon_\mathrm{q}=0} = R_\mathrm{w}, \tag{S45}$$

The left-hand side in eqs. (S44) and (S45) gives the ratio of the marginal change in the number of inseminated queens to the marginal change in the number of queens produced when the allocation schedule is varied. This ratio is expressed in terms of a variational derivatives $\delta x_k(\mathbf{u})/\delta \eta_\tau(t)$ measuring the change in the number of individuals of type $k \in \{\mathrm{q}, \mathrm{iq}\}$ associated with a focal colony where phenotype $\mathbf{u}$ is expressed. Hence, we have showed that when $u_\mathrm{f}^*(t) = \hat{u}_\mathrm{f}^*(t)$ is a singular arc, the marginal substitution rate of inseminated queens with produced queens is given be the relatedness asymmetry $R_c$.

###### 2.4.1 The critical sex ratio under delayed dispersal

In this section, we derive the condition for the (critical) sex ratio at the end of the season ($t = T$) under delayed dispersal. Equation (S4) for delayed dispersal together with eq. (S5) yields that

$$x_\mathrm{iq}(\mathbf{u}^s) = x_\mathrm{m}(\mathbf{u}^s)\frac{y_\mathrm{q}(\mathbf{v})}{y_\mathrm{m}(\mathbf{v})}. \tag{S46}$$

Hence, it follows from eq. (S23) that under delayed dispersal

$$\left.\frac{\delta x_{\mathrm{iq}}(\mathbf{u})}{\delta \eta_{\tau}(t)}\right|_{\epsilon_{\mathrm{f}}=0,\epsilon_{\mathrm{q}}=0} = \frac{x_{\mathrm{q}}^{*}(\mathbf{u}^{*})}{x_{\mathrm{m}}^{*}(\mathbf{u}^{*})}\left.\frac{\delta x_{\mathrm{m}}(\mathbf{u})}{\delta \eta_{\tau}(t)}\right|_{\epsilon_{\mathrm{f}}=0,\epsilon_{\mathrm{q}}=0}. \tag{S47}$$

We will show later in section 5 that under delayed dispersal $\hat{u}_{\mathrm{f}}^{*}(t)$ is a singular arc during $t \in [t_{c,1}^{*}, T]$, where $t_{c,1}^{*}$ is the time, when $u_{\mathrm{f}}^{*}(t)$ becomes a singular arc under the control mode $c \in \{\mathrm{q}, \mathrm{w}, \mathrm{mx}\}$. Substituting eq. (S47) into eqs. (S44)–(S45) yields for queen ($c = \mathrm{q}$) and worker ($c = \mathrm{w}$) control

$$\frac{x_{\mathrm{q}}^{*}(\mathbf{u}^{*})}{x_{\mathrm{m}}^{*}(\mathbf{u}^{*})} = -\left.\frac{\delta x_{\mathrm{q}}(\mathbf{u})/\delta \eta_{\mathrm{f}}(t)}{\delta x_{\mathrm{m}}(\mathbf{u})/\delta \eta_{\mathrm{f}}(t)}\right|_{\epsilon_{\mathrm{f}}=0,\epsilon_{\mathrm{q}}=0} \times R_{c}, \tag{S48}$$

and mixed control

$$\frac{x_{\mathrm{q}}^{*}(\mathbf{u}^{*})}{x_{\mathrm{m}}^{*}(\mathbf{u}^{*})} = -\left.\frac{\delta x_{\mathrm{q}}(\mathbf{u})/\delta \eta_{\mathrm{f}}(t)}{\delta x_{\mathrm{m}}(\mathbf{u})/\delta \eta_{\mathrm{f}}(t)}\right|_{\epsilon_{\mathrm{f}}=0,\epsilon_{\mathrm{q}}=0} \times R_{\mathrm{q}} \tag{S49}$$

where the right-hand side depends on the ratio of the marginal values of males relative to queens (i.e., the marginal rate of substitution of producing males instead of new queens). If males and juvenile queens are equally costly to produce and they have the same mortality rate ($\mu_{\mathrm{q}} = \mu_{\mathrm{m}} = \mu_{\mathrm{r}}$) and the same growth schedule, then the marginal product is negative one. Hence, the (critical) sex ratio at the end of the season ($t = T$) under delayed dispersal for equal mortality rates of males and queens is equal to the relatedness asymmetry (recall eq. S35), i.e.

$$\begin{aligned} \frac{x_{\mathrm{q}}^{*}(\mathbf{u}^{*})}{x_{\mathrm{m}}^{*}(\mathbf{u}^{*})} &= R_{\mathrm{q}} \qquad \text{(queen and mixed control)} \\ \frac{x_{\mathrm{q}}^{*}(\mathbf{u}^{*})}{x_{\mathrm{m}}^{*}(\mathbf{u}^{*})} &= R_{\mathrm{w}} \qquad \text{(worker control).} \end{aligned} \tag{S50}$$

### 3 First-order condition expressed in terms of optimal control problem

In this section, we use optimal control theory in order to solve the marginal value equation (S44) for different scenarios of our model by way of applying Pontryagin's weak maximum principle (e.g., Bryson and Ho, 1975; Sydsæter et al., 2008 for broad introductions and Day and Taylor, 2000; González-Forero et al., 2017; Iwasa and Roughgarden, 1984; Macevicz and Oster, 1976; Perrin, 1992 for previous application to evolutionary biology).

#### 3.1 Formulation of the optimal control problem

##### 3.1.1 The basic problem

We start by formulating the dynamic resource allocation problem as a classical optimal control problem. Finding the candidate uninvadable schedule entails establishing an optimal pair

$$(\mathbf{u}^{*}, \mathbf{x}^{*}) = \left(\{\mathbf{u}^{*}(t)\}_{t\in[0,T]}, \{\mathbf{x}^{*}(t)\}_{t\in[0,T]}\right), \tag{S51}$$

where $\mathbf{u}^*(t) = (u_\mathrm{f}^*(t), u_\mathrm{q}^*(t))$ and $\mathbf{x}^*(t) = (x_\mathrm{w}^*(t), x_\mathrm{q}^*(t), x_\mathrm{m}^*(t))$ are vectors of uninvadable (optimal) control and state variables, respectively. The optimal pair $(\mathbf{u}^*, \mathbf{x}^*)$ is a solution to eqs. (S12) and (S13) under single-party control and mixed control, respectively. That is, it maximizes the invasion fitness $W_c(\mathbf{u}, \mathbf{v})$ (as given by eq. S10).

The so-called control variables for the maximization problems are the (resource allocation) phenotypes expressed in colonies founded by individuals who are homozygous for the mutant allele (recall eq. S1)

$$u_\mathrm{f}(t) \text{ and } u_\mathrm{q}(t)\ \forall t \in [0, T] \tag{S52}$$

where

$$0 \leq u_\tau(t) \leq 1\ \forall t \in [0, T],\ \tau \in \{\mathrm{f}, \mathrm{q}\}\,. \tag{S53}$$

For delayed dispersal, the vector of the so-called state variables for the optimal control problems can be expressed as

$$\mathbf{x}(t) = \begin{cases} (x_\mathrm{w}^\mathrm{q}(t), x_\mathrm{q}^\mathrm{q}(t), x_\mathrm{m}^\mathrm{q}(t)) & (\mathbf{u}_\tau,\, \mathbf{v}_\tau \text{ under queen control}), \\ (x_\mathrm{w}^\mathrm{q}(t), x_\mathrm{q}^\mathrm{q}(t), x_\mathrm{m}^\mathrm{q}(t), x_\mathrm{w}^\mathrm{m}(t), x_\mathrm{q}^\mathrm{m}(t), x_\mathrm{m}^\mathrm{m}(t)) & (\mathbf{u}_\tau,\, \mathbf{v}_\tau \text{ under worker control}). \end{cases} \tag{S54}$$

For direct dispersal, the vector of state variables for the optimal control problems can be expressed as

$$\mathbf{x}(t) = \begin{cases} (x_\mathrm{w}^\mathrm{q}(t), x_\mathrm{q}^\mathrm{q}(t), x_\mathrm{m}^\mathrm{q}(t), x_\mathrm{iq}^\mathrm{q}(t)) & (\mathbf{u}_\tau,\, \mathbf{v}_\tau \text{ under queen control}), \\ (x_\mathrm{w}^\mathrm{q}(t), x_\mathrm{q}^\mathrm{q}(t), x_\mathrm{m}^\mathrm{q}(t), x_\mathrm{iq}^\mathrm{q}(t), x_\mathrm{w}^\mathrm{m}(t), x_\mathrm{q}^\mathrm{m}(t), x_\mathrm{m}^\mathrm{m}(t), x_\mathrm{iq}^\mathrm{m}(t)) & (\mathbf{u}_\tau,\, \mathbf{v}_\tau \text{ under worker control}). \end{cases} \tag{S55}$$

In addition, the vector of dynamical variables involved in the invasion fitness (eqs. S62–S62) for the optimal control problems can be expressed as

$$\mathbf{y}(t) = (y_\mathrm{w}(t), y_\mathrm{q}(t), y_\mathrm{m}(t)). \tag{S56}$$

The rate of change in state variables appearing in eqs. (S54)–(S55) is described by the differential equations

$$\dot{x}_k^s(t) = g_{k,\mathrm{u}}^s(\mathbf{x}(t), \mathbf{u}(t)) = g_{k,\mathrm{u}}^s(t), \text{ with } x_k^s(0) = y_{k,0} \text{ for } k \in \{\mathrm{q}, \mathrm{m}, \mathrm{w}, \mathrm{iq}\} \text{ and } s \in \{\mathrm{q}, \mathrm{m}\}\,, \tag{S57}$$

where upper "$\cdot$" denotes the time derivative, $(y_{\mathrm{w},0}, y_{\mathrm{q},0}, y_{\mathrm{m},0}, y_{\mathrm{iq},0}) = (1, 0, 0, 0)$ (fixed) and $x_k^s(T) = x_k(\mathbf{u}^s)$ is free and the differential equations can be expressed as

$$\begin{aligned} g_{k,\mathrm{u}}^s(t) &= b a_{k,\mathrm{u}}^s(t) x_k^s(t) - \mu_k x_k^s(t) \text{ for } k \in \{\mathrm{q}, \mathrm{m}, \mathrm{w}\} \text{ and } s \in \{\mathrm{q}, \mathrm{m}\}\,, \\ g_{\mathrm{iq},\mathrm{u}}^s(t) &= x_\mathrm{m}^s \frac{b a_\mathrm{q}(t) y_\mathrm{w}(t)}{y_\mathrm{m}(t)} - \mu_\mathrm{q} x_\mathrm{iq}^s(t) \text{ for } s \in \{\mathrm{q}, \mathrm{m}\}\,, \end{aligned} \tag{S58}$$

where the mutant allocation schedules $a_{k,\mathrm{u}}^s(t)$ and resident allocation schedules $a_k(t)$, are given by eqs. (S2) and (3) of the main text, respectively.

The rate of change in dynamic variables appearing in eq. (S56) is described by the differential equations

$$\dot{y}_k = g_k(\mathbf{y}(t), \mathbf{v}(t), t) = g_k(t), \text{ with } y_k(0) = y_{k,0} \text{ for } k \in \{\mathrm{q}, \mathrm{m}, \mathrm{w}\}\,, \tag{S59}$$

where $(y_{\mathrm{w},0}, y_{\mathrm{q},0}, y_{\mathrm{m},0}) = (1, 0, 0)$ (fixed) and $y_k(T) = y_k(\mathbf{v})$ is free and the differential equations can be expressed as

$$g_k(t) = ba_k(\mathbf{v}(t))y_{\mathrm{w}}(t) - \mu_k y_k(t) \text{ for } k \in \{\mathrm{q}, \mathrm{m}, \mathrm{w}\} \text{ and } s \in \{\mathrm{q}, \mathrm{m}\}\,, \tag{S60}$$

where the resident allocation schedules, $a_k(\mathbf{v}(t))$, are given by and eq. (3) of the main text.

##### 3.1.2 Explicit expression for invasion fitness function

Henceforth, we write the invasion fitness of a mutant allele as $W_c(\mathbf{u}, \mathbf{v}) \equiv W_{c,d}(\mathbf{u}, \mathbf{v})$, where the additional subscript $d \in \{\mathrm{del}, \mathrm{dir}\}$ emphasizes the scenario of dispersal of sexuals, delayed and direct dispersal, respectively. Substituting the transmission frequencies for haplodiploids ($\gamma_{\mathrm{qq}} = 1/2$, $\gamma_{\mathrm{mq}} = 1$, $\gamma_{\mathrm{qm}} = 1/2$, $\gamma_{\mathrm{mm}} = 0$) into eq. (S10) and using eq. (S11) we can simplify the expression for the invasion fitness (eq. S10) under delayed dispersal to

$$W_{c,\mathrm{del}}(\mathbf{u}, \mathbf{v}) = \begin{cases} \dfrac{1}{4}\dfrac{x_{\mathrm{q}}(\mathbf{u}^{\mathrm{q}})}{y_{\mathrm{q}}(\mathbf{v})} + \sqrt{\dfrac{1}{16}\left[\dfrac{x_{\mathrm{q}}(\mathbf{u}^{\mathrm{q}})}{y_{\mathrm{q}}(\mathbf{v})}\right]^2 + \dfrac{1}{2}\dfrac{x_{\mathrm{m}}(\mathbf{u}^{\mathrm{q}})}{y_{\mathrm{m}}(\mathbf{v})}} & (\mathbf{u}_{\mathcal{T}},\, \mathbf{v}_{\mathcal{T}} \text{ under queen control}), \\ \dfrac{1}{4}\dfrac{x_{\mathrm{q}}(\mathbf{u}^{\mathrm{q}})}{y_{\mathrm{q}}(\mathbf{v})} + \sqrt{\dfrac{1}{16}\left[\dfrac{x_{\mathrm{q}}(\mathbf{u}^{\mathrm{q}})}{y_{\mathrm{q}}(\mathbf{v})}\right]^2 + \dfrac{1}{2}\dfrac{x_{\mathrm{q}}(\mathbf{u}^{\mathrm{m}})}{y_{\mathrm{q}}(\mathbf{v})}\dfrac{x_{\mathrm{m}}(\mathbf{u}^{\mathrm{q}})}{y_{\mathrm{m}}(\mathbf{v})}} & (\mathbf{u}_{\mathcal{T}},\, \mathbf{v}_{\mathcal{T}} \text{ under worker control}) \end{cases} \tag{S61}$$

and under direct dispersal to

$$W_{c,\mathrm{dir}}(\mathbf{u}, \mathbf{v}) = \begin{cases} \dfrac{1}{4}\dfrac{x_{\mathrm{q}}(\mathbf{u}^{\mathrm{q}})}{y_{\mathrm{q}}(\mathbf{v})} + \sqrt{\dfrac{1}{16}\left[\dfrac{x_{\mathrm{q}}(\mathbf{u}^{\mathrm{q}})}{y_{\mathrm{q}}(\mathbf{v})}\right]^2 + \dfrac{1}{2}\dfrac{x_{\mathrm{iq}}(\mathbf{u}^{\mathrm{q}})}{y_{\mathrm{q}}(\mathbf{v})}} & (\mathbf{u}_{\mathcal{T}},\, \mathbf{v}_{\mathcal{T}} \text{ under queen control}), \\ \dfrac{1}{4}\dfrac{x_{\mathrm{q}}(\mathbf{u}^{\mathrm{q}})}{y_{\mathrm{q}}(\mathbf{v})} + \sqrt{\dfrac{1}{16}\left[\dfrac{x_{\mathrm{q}}(\mathbf{u}^{\mathrm{q}})}{y_{\mathrm{q}}(\mathbf{v})}\right]^2 + \dfrac{1}{2}\dfrac{x_{\mathrm{q}}(\mathbf{u}^{\mathrm{m}})}{y_{\mathrm{q}}(\mathbf{v})}\dfrac{x_{\mathrm{iq}}(\mathbf{u}^{\mathrm{q}})}{y_{\mathrm{q}}(\mathbf{v})}} & (\mathbf{u}_{\mathcal{T}},\, \mathbf{v}_{\mathcal{T}} \text{ under worker control}). \end{cases} \tag{S62}$$

Note that for mixed control we have the invasion fitness function under queen control $W_{\mathrm{q},d}(\mathbf{u}, \mathbf{v})$ to determine $u_{\mathrm{f}}^*(t)$ and the invasion fitness function under worker control $W_{\mathrm{w},d}(\mathbf{u}, \mathbf{v})$ to determine $u_{\mathrm{q}}^*$. These simplified expressions of invasion fitness will turn out useful for solving numerically the optimal control problem (see section 11) and also conceptually, because it makes it explicit how the invasion fitness depends on the state $\mathbf{x}(t)$ and dynamic variables $\mathbf{y}(t)$.

##### 3.2 Pontryagin's weak maximum principle and the Hamiltonian

The necessary first-order condition for uninvadability (given by eq. S36) can be expressed in terms of pointwise functional derivatives (given by eq. S39, see e.g. Parr and Yang, 1989, p. 246); that is

$$\left.\frac{\mathrm{d}W_{c,d}(\mathbf{u},\mathbf{v})}{\mathrm{d}\epsilon_\tau}\right|_{\epsilon_\mathrm{f}=0,\epsilon_\mathrm{q}=0} = \int_0^T \left.\frac{\delta W_{c,d}(\mathbf{u},\mathbf{u}^*)}{\delta\eta_\tau(t)}\right|_{\epsilon_\mathrm{f}=0,\epsilon_\mathrm{q}=0} \eta_\tau(t)dt. \tag{S63}$$

This expression can be thought of as a functional analogue of the formula for the total derivative of a function $W(\eta_1(t),\eta_2(t),...)$: $\mathrm{d}W/\,\mathrm{d}t = \sum_i(\partial W/\partial\eta_i)(\partial\eta_i/\partial t)$ (see e.g. Parr and Yang, 1989, p. 246).

Hence, the first-order condition for uninvadability (eq. S36) can be expressed in terms of point-wise marginal change, which can be expressed under single-party control as

$$\int_0^T \left.\frac{\delta W_{c,d}(\mathbf{u},\mathbf{u}^*)}{\delta\eta_\tau(t)}\right|_{\epsilon_\mathrm{f}=0,\epsilon_\mathrm{q}=0} \eta_\tau(t)dt \le 0 \qquad \text{for } \tau=\mathrm{f} \text{ and } \tau=\mathrm{q}, \tag{S64}$$

and for mixed party control as

$$\int_0^T \left.\frac{\delta W_{c,d}(\mathbf{u},\mathbf{u}^*)}{\delta\eta_\tau(t)}\right|_{\epsilon_\mathrm{f}=0,\epsilon_\mathrm{q}=0} \eta_\tau(t)dt \le 0 \quad \text{and} \quad \int_0^T \left.\frac{\delta W_{c,d}(\mathbf{u},\mathbf{u}^*)}{\delta\eta_\tau(t)}\right|_{\epsilon_\mathrm{f}=0,\epsilon_\mathrm{q}=0} \eta_\tau(t)dt \le 0. \tag{S65}$$

for any feasible $\eta_\tau(t)$.

Pontryagin's weak maximum principle yields that the pointwise marginal change (as in eq. S63) can be restated for both single-party (see e.g. Speyer and Jacobson, 2010, p. 61) and mixed control (see e.g. Mazalov, 2014, p. 372, Theorem 10.8) as follows

$$\left.\frac{\delta W_{c,d}(\mathbf{u},\mathbf{u}^*)}{\delta\eta_\tau(t)}\right|_{\epsilon_\mathrm{f}=0,\epsilon_\mathrm{q}=0} = \left.\frac{\partial H_{c,d}(\mathbf{u}(t),\mathbf{x}^*(t),\boldsymbol{\lambda}(t))}{\partial u_\tau(t)}\right|_{\mathbf{u}=\mathbf{v}=\mathbf{u}^*} \qquad \forall t\in[0,T]. \tag{S66}$$

Here, $H_{c,d}(\mathbf{u}(t),\mathbf{x}^*(t),\boldsymbol{\lambda}(t))$ is the Hamiltonian function, which allows to transform a dynamic optimization problem into a sequence of static optimization problems by providing a marginal condition that must be satisfied over the whole time schedule (see section 3.3 for the full interpretation of the Hamiltonian), and which for our problem can be expressed as

$$\begin{aligned} H_{c,d}(\mathbf{u}(t),\mathbf{x}^*(t),\boldsymbol{\lambda}(t)) = &\sum_{k\in\{\mathrm{w,q,m}\}} \lambda_k^\mathrm{q}(t)g_{k,\mathrm{u}}^\mathrm{q}(t) + \\ &\delta_\mathrm{cw}\sum_{k\in\{\mathrm{w,q,m}\}} \lambda_k^\mathrm{m}(t)g_{k,\mathrm{u}}^\mathrm{m}(t) + \delta_{d\mathrm{dir}}\sum_{s\in\{\mathrm{q,m}\}} \lambda_\mathrm{iq}^s(t)g_{\mathrm{iq,u}}^s(t), \end{aligned} \tag{S67}$$

where index $c\in\{\mathrm{q,w,mx}\}$ emphasizes the mode of control and $d\in\{\mathrm{del,dir}\}$ emphasizes the time of dispersal of sexuals. In eq. (S67), $\delta_{ij}$ is the Kronecker delta function, i.e.

$$\delta_{ij} = \begin{cases} 1 & \text{for } i = j, \\ 0 & \text{for } i \neq j \end{cases} \tag{S68}$$

$\lambda_k^s(t)$ is a costate variable associated with the state variable $x_k^s(t)$ and $\boldsymbol{\lambda}(t)$ is a vector of costate variables and for delayed dispersal it can be expressed as

$$\boldsymbol{\lambda}(t) = \begin{cases} (\lambda_{\mathrm{w}}^{\mathrm{q}}(t), \lambda_{\mathrm{q}}^{\mathrm{q}}(t), \lambda_{\mathrm{m}}^{\mathrm{q}}(t)) & (\mathbf{u}_\tau, \mathbf{v}_\tau \text{ under queen control}), \\ (\lambda_{\mathrm{w}}^{\mathrm{q}}(t), \lambda_{\mathrm{q}}^{\mathrm{q}}(t), \lambda_{\mathrm{m}}^{\mathrm{q}}(t), \lambda_{\mathrm{w}}^{\mathrm{m}}(t), \lambda_{\mathrm{q}}^{\mathrm{m}}(t), \lambda_{\mathrm{m}}^{\mathrm{m}}(t)) & (\mathbf{u}_\tau, \mathbf{v}_\tau \text{ under worker control}), \end{cases} \tag{S69}$$

and for direct dispersal it can be expressed as

$$\boldsymbol{\lambda}(t) = \begin{cases} (\lambda_{\mathrm{w}}^{\mathrm{q}}(t), \lambda_{\mathrm{q}}^{\mathrm{q}}(t), \lambda_{\mathrm{m}}^{\mathrm{q}}(t), \lambda_{\mathrm{iq}}^{\mathrm{q}}(t)) & (\mathbf{u}_\tau, \mathbf{v}_\tau \text{ under queen control}), \\ (\lambda_{\mathrm{w}}^{\mathrm{q}}(t), \lambda_{\mathrm{q}}^{\mathrm{q}}(t), \lambda_{\mathrm{m}}^{\mathrm{q}}(t), \lambda_{\mathrm{iq}}^{\mathrm{q}}(t), \lambda_{\mathrm{w}}^{\mathrm{m}}(t), \lambda_{\mathrm{q}}^{\mathrm{m}}(t), \lambda_{\mathrm{m}}^{\mathrm{m}}(t), \lambda_{\mathrm{iq}}^{\mathrm{m}}(t)) & (\mathbf{u}_\tau, \mathbf{v}_\tau \text{ under worker control}). \end{cases} \tag{S70}$$

The differential equations for the costate variables appearing eqs. (S69)–(S70) are given by the derivatives of the Hamiltonian with respect to the corresponding state variables, i.e.

$$\dot{\lambda}_k^s(t) = -\left.\frac{\partial H_{c,d}(\mathbf{u}^*(t), \mathbf{x}(t), \boldsymbol{\lambda}(t))}{\partial x_k^s(t)}\right|_{\mathbf{x}=\mathbf{y}=\mathbf{x}^*} \tag{S71}$$

Since $\mathbf{x}(T)$ is free, the transversality conditions for the co-state variables are given by

$$\lambda_k^s(T) = \left.\frac{\partial W_{c,d}(\mathbf{u}, \mathbf{v})}{\partial x_k^s(T)}\right|_{\mathbf{x}=\mathbf{y}=\mathbf{x}^*} \tag{S72}$$

(e.g., Bryson and Ho, 1975; Sydsæter et al., 2008).

##### 3.3 Interpretation of the Hamiltonian and costate variable

The quantity $H_{c,d}(\mathbf{u}(t), \mathbf{x}^*(t), \boldsymbol{\lambda}(t))\,\mathrm{d}t = H_{c,d}(t)\,\mathrm{d}t$ can be interpreted as the total contribution to the invasion fitness $W_{c,d}(\mathbf{u}, \mathbf{v})$ by an increase in the production of individuals of different types for a certain (constant) allocation schedule $u_\tau(t) = \bar{u}_\tau$ during the interval $[t, t + \mathrm{d}t]$ (e.g. Dorfman, 1969, Sethi and Thompson, 2006, p. 34). As a consequence, the control variables $u_\tau(t)$ for a given interval should be chosen such that to maximize $H_{c,d}(t)$. This implies that the dynamic optimization problem of maximizing the invasion fitness $W_{c,d}(\mathbf{u}, \mathbf{v})$ can be transformed into a sequence of static problems of maximizing the corresponding Hamiltonian $H_{c,d}(t)$ at instants $t \in [0, T]$. Hence, the Hamiltonian can be interpreted as a rate at which the invasion fitness (which is defined at final time $T$) increases at time $t$ and $\partial H_{c,d}(t)/\partial u_\tau(t)$ represents a variation in invasion fitness due to a unit impulse (Dirac function) in $u_\tau(t)$ at time $t$, while satisfying the state equations (Bryson and Ho, 1975, p. 49). More precisely, $\partial H_{c,d}(t)/\partial u_\tau(t)$ measures the net effect on invasion fitness that the marginal change in the trait value $u_\tau(t)$ has through immediate change in the trait value $u_\tau(t)$ at time $t$ and through the cascading effects that this change has on the invasion fitness by changing the state variables (the numbers of individuals of different types) from time $t$ onward until time $T$.

A costate variable $\lambda_k^s(t)$ can be interpreted as the effect on invasion fitness for a marginal change in the corresponding state variable $x_k^s(t)$ when colony resources are allocated to production of type $k$ individuals at time $t$ in a colony founded by a mutant individual of type $s$. Therefore, informally, a costate variable $\lambda_k^s(t)$ gives the value (measured as the increase in invasion fitness) of each unit resource invested at time $t$ into the production of type $k$ individuals at time $t$ in a colony founded by a mutant individual of type $s$. Hence, the costate variables are of extreme importance since only the individuals that yield the highest investment value in invasion fitness should be produced at any given time.

##### 3.4 Derivatives of the Hamiltonian

It follows from eqs. (S64), (S65), (S66) and (S53) (see e.g. Kamien and Schwartz, 2012, p. 185-186 for full explanation) that for all $t \in [0, T]$

$$\begin{aligned}
&\text{if} \quad \left.\frac{\partial H_{c,d}(\mathbf{u}(t), \mathbf{x}^*(t), \boldsymbol{\lambda}(t))}{\partial u_\tau(t)}\right|_{\mathbf{u}=\mathbf{v}=\mathbf{u}^*} < 0 \quad \text{then} \quad u_\tau^*(t) = 0, \\
&\text{if} \quad \left.\frac{\partial H_{c,d}(\mathbf{u}(t), \mathbf{x}^*(t), \boldsymbol{\lambda}(t))}{\partial u_\tau(t)}\right|_{\mathbf{u}=\mathbf{v}=\mathbf{u}^*} = 0 \quad \text{then} \quad 0 \leq u_\tau^*(t) = \hat{u}_\tau^*(t) \leq 1, \\
&\text{if} \quad \left.\frac{\partial H_{c,d}(\mathbf{u}(t), \mathbf{x}^*(t), \boldsymbol{\lambda}(t))}{\partial u_\tau(t)}\right|_{\mathbf{u}=\mathbf{v}=\mathbf{u}^*} > 0 \quad \text{then} \quad u_\tau^*(t) = 1,
\end{aligned} \tag{S73}$$

where $\hat{u}_\tau^*(t)$ denotes that the control $u_\tau^*(t)$ is a singular arc (Sethi and Thompson, 2006, p. 407). An allocation trait is a singular arc ($u_\tau^*(t) = \hat{u}_\tau^*(t)$) when the Hamiltonian is linear (or more strictly, affine) in the control and the derivative $\partial H_{c,d}(\mathbf{u}(t), \mathbf{x}^*(t), \boldsymbol{\lambda}(t))/\partial u_\tau(t)|_{\mathbf{u}=\mathbf{u}^*} = 0$. Hence, the first-order condition (given by eq. S64 or eq. S65) is satisfied, but the control variable $u_\tau(t)$ does not directly appear in the first-order condition. More generally, an optimal control is a singular arc, if the value of variational Hamiltonian is unchanged to the second order from a weak first-order variation of the control at each point of the arc (Robbins, 1967).

In order to ascertain the uninvadable allocation schedule $\mathbf{u}^*$ from eq. (S73), we need to determine the derivatives of the Hamiltonian with respect to $u_\tau(t)$. Substituting eq. (S58) into eq. (S67) and taking the derivative with respect to $u_\tau$ produces

$$\begin{aligned}
\left.\frac{\partial H_{c,d}(\mathbf{u}(t), \mathbf{x}^*(t), \boldsymbol{\lambda}(t))}{\partial u_\tau(t)}\right|_{\mathbf{u}=\mathbf{v}=\mathbf{u}^*} =& b x_{\mathrm{w}}^*(t) \left[ \lambda_{\mathrm{w}}^{\mathrm{q}}(t) \frac{\partial a_{\mathrm{w,u}}^{\mathrm{q}}(t)}{\partial u_\tau(t)} + \lambda_{\mathrm{q}}^{\mathrm{q}}(t) \frac{\partial a_{\mathrm{q,u}}^{\mathrm{q}}(t)}{\partial u_\tau(t)} + \lambda_{\mathrm{m}}^{\mathrm{q}}(t) \frac{\partial a_{\mathrm{m,u}}^{\mathrm{q}}(t)}{\partial u_\tau(t)} \right. \\
& \left. + \delta_{\mathrm{cw}} \left( \lambda_{\mathrm{w}}^{\mathrm{m}}(t) \frac{\partial a_{\mathrm{w,u}}^{\mathrm{m}}(t)}{\partial u_\tau(t)} + \lambda_{\mathrm{m}}^{\mathrm{m}}(t) \frac{\partial a_{\mathrm{q,u}}^{\mathrm{m}}(t)}{\partial u_\tau(t)} + \lambda_{\mathrm{m}}^{\mathrm{m}}(t) \frac{\partial a_{\mathrm{m,u}}^{\mathrm{m}}(t)}{\partial u_\tau(t)} \right) \right],
\end{aligned} \tag{S74}$$

with partial derivatives

$$
\begin{aligned}
\frac{\partial a^s_{\mathrm{w,u}}(t)}{\partial u_{\mathrm{f}}(t)} &= (1-u^*_{\mathrm{q}}(t))p^s_c\,, &\quad \frac{\partial a^s_{\mathrm{q,u}}(t)}{\partial u_{\mathrm{f}}(t)} &= u^*_{\mathrm{q}}(t)p^s_c\,, &\quad \frac{\partial a^s_{\mathrm{m,u}}(t)}{\partial u_{\mathrm{f}}(t)} &= -p^s_c\,,\\
\frac{\partial a^s_{\mathrm{w,u}}(t)}{\partial u_{\mathrm{q}}(t)} &= -u^*_{\mathrm{f}}(t)p^s_c\,, & \frac{\partial a^s_{\mathrm{q,u}}(t)}{\partial u_{\mathrm{q}}(t)} &= u^*_{\mathrm{f}}(t)p^s_c, \text{ and} & \frac{\partial a^s_{\mathrm{m,u}}(t)}{\partial u_{\mathrm{q}}(t)} &= 0.
\end{aligned}
\tag{S75}
$$

Hence, the derivatives of the Hamiltonian with respect to controls $u_{\mathrm{f}}$ and $u_{\mathrm{q}}$ can be written as

$$
\begin{aligned}
\left.\frac{\partial H_{c,d}(\mathbf{u}(t),\mathbf{x}^*(t),\boldsymbol{\lambda}(t))}{\partial u_{\mathrm{f}}(t)}\right|_{\mathbf{u}=\mathbf{v}=\mathbf{u}^*} &= b x^*_{\mathrm{w}}(t)\left(u^*_{\mathrm{q}}(t)\sigma^c_1(t)-\sigma^c_2(t)\right)\\
\left.\frac{\partial H_{c,d}(\mathbf{u}(t),\mathbf{x}^*(t),\boldsymbol{\lambda}(t))}{\partial u_{\mathrm{q}}(t)}\right|_{\mathbf{u}=\mathbf{v}=\mathbf{u}^*} &= b x^*_{\mathrm{w}}(t)u^*_{\mathrm{f}}(t)\sigma^c_1(t).
\end{aligned}
\tag{S76}
$$

Expressions $\sigma^c_1(t)$ and $\sigma^c_2(t)$ and $\sigma^c_1(t)-\sigma^c_2(t)$ in eq. (S76) are functions of the costate variables $\lambda^s_k(t)$ and the expected frequencies $p^s_c$ of the mutant allele in the party $c$

$$
\begin{aligned}
\sigma^c_1(t) &= p^{\mathrm{q}}_c\left(\lambda^{\mathrm{q}}_{\mathrm{q}}(t)-\lambda^{\mathrm{q}}_{\mathrm{w}}(t)\right)+\delta_{c\mathrm{w}}p^{\mathrm{m}}_c\left(\lambda^{\mathrm{m}}_{\mathrm{q}}(t)-\lambda^{\mathrm{m}}_{\mathrm{w}}(t)\right),\\
\sigma^c_2(t) &= p^{\mathrm{q}}_c\left(\lambda^{\mathrm{q}}_{\mathrm{m}}(t)-\lambda^{\mathrm{q}}_{\mathrm{w}}(t)\right)+\delta_{c\mathrm{w}}p^{\mathrm{m}}_c\left(\lambda^{\mathrm{m}}_{\mathrm{m}}(t)-\lambda^{\mathrm{m}}_{\mathrm{w}}(t)\right),\\
\sigma^c_1(t)-\sigma^c_2(t) &= p^{\mathrm{q}}_c\left(\lambda^{\mathrm{q}}_{\mathrm{q}}(t)-\lambda^{\mathrm{q}}_{\mathrm{m}}(t)\right)+\delta_{c\mathrm{w}}p^{\mathrm{m}}_c\left(\lambda^{\mathrm{m}}_{\mathrm{q}}(t)-\lambda^{\mathrm{m}}_{\mathrm{m}}(t)\right),
\end{aligned}
\tag{S77}
$$

where $p^{\mathrm{q}}_{\mathrm{q}} = 1/2$, $p^{\mathrm{m}}_{\mathrm{q}} = 0$, $p^{\mathrm{q}}_{\mathrm{w}} = 1/4$, and $p^{\mathrm{m}}_{\mathrm{w}} = 1/(2M)$. It follows from eq. (S76) and (S73) that $u^*_{\mathrm{f}}(t)$ is determined from the sign of expression $u^*_{\mathrm{q}}(t)\sigma^c_1(t)-\sigma^c_2(t)$ and $u^*_{\mathrm{q}}(t)$ is determined from the sign of expression $u^*_{\mathrm{f}}(t)\sigma^c_1(t)$, since $b>0$ and $x^*_{\mathrm{w}}(t)>0$ for biological reasons. Since, the signs of functions $\sigma^c_1(t)$ and $\sigma^c_2(t)$ will be instrumental in determining the signs of these expressions, we call them the switching functions, which are analogous to the switching functions in linear optimal control problems (e.g., Bryson and Ho, 1975, p. 111).

Substituting the expected frequencies $p^s_c$ of the mutant allele in the party $c$ and eq. (S91) into eq. (S77) yields that for queen control

$$
\begin{aligned}
\sigma^{\mathrm{q}}_1(t) &= \frac{1}{2}\left(\lambda^{\mathrm{q}}_{\mathrm{q}}(t)-\lambda^{\mathrm{q}}_{\mathrm{w}}(t)\right),\\
\sigma^{\mathrm{q}}_2(t) &= \frac{1}{2}\left(\lambda^{\mathrm{q}}_{\mathrm{m}}(t)-\lambda^{\mathrm{q}}_{\mathrm{w}}(t)\right),\\
\sigma^{\mathrm{q}}_1(t)-\sigma^{\mathrm{q}}_2(t) &= \frac{1}{2}\left(\lambda^{\mathrm{q}}_{\mathrm{q}}(t)-\lambda^{\mathrm{q}}_{\mathrm{m}}(t)\right)
\end{aligned}
\tag{S78}
$$

and for worker control

$$
\begin{aligned}
\sigma^{\mathrm{w}}_1(t) &= \frac{2+M}{4M}\lambda^{\mathrm{q}}_{\mathrm{q}}(t)-\frac{1}{4}\lambda^{\mathrm{q}}_{\mathrm{w}}(t)-\frac{1}{2M}\lambda^{\mathrm{m}}_{\mathrm{w}}(t),\\
\sigma^{\mathrm{w}}_2(t) &= \frac{1}{4}\lambda^{\mathrm{q}}_{\mathrm{q}}(t)-\frac{1}{4}\lambda^{\mathrm{q}}_{\mathrm{w}}(t)-\frac{1}{2M}\lambda^{\mathrm{m}}_{\mathrm{w}}(t),\\
\sigma^{\mathrm{w}}_1(t)-\sigma^{\mathrm{w}}_2(t) &= \frac{1}{4}\left(\frac{2+M}{M}\lambda^{\mathrm{q}}_{\mathrm{q}}(t)-\lambda^{\mathrm{q}}_{\mathrm{m}}(t)\right).
\end{aligned}
\tag{S79}
$$

Under delayed dispersal if the mortality rates of males and queens are equal (i.e. $\mu_{\mathrm{q}} = \mu_{\mathrm{m}} = \mu_{\mathrm{r}}$), it follows from the costate equations (S82) and from the transversality conditions (S86) that the switching functions for worker control further simplify to

$$\begin{aligned} \sigma_1^{\mathrm{w}}(t) &= \frac{2+M}{4M}\left(\lambda_{\mathrm{q}}^{\mathrm{q}}(t) - \lambda_{\mathrm{w}}^{\mathrm{q}}(t)\right), \\ \sigma_2^{\mathrm{w}}(t) &= \frac{1}{4}\left(\lambda_{\mathrm{m}}^{\mathrm{q}}(t) - \frac{2+M}{M}\lambda_{\mathrm{w}}^{\mathrm{q}}(t)\right), \\ \sigma_1^{\mathrm{w}}(t) - \sigma_2^{\mathrm{w}}(t) &= \frac{1}{4}\left(\frac{2+M}{M}\lambda_{\mathrm{q}}^{\mathrm{q}}(t) - \lambda_{\mathrm{m}}^{\mathrm{q}}(t)\right). \end{aligned} \tag{S80}$$

### 4 Global qualitative properties of the uninvadable allocation schedule

Here we describe the scheme of deriving the uninvadable allocation schedule $\mathbf{u}^*$ under different assumptions of the model. First we present the conditions that the candidate allocation schedule has to satisfy to be consistent with the first-order condition for uninvadability, which gives rise to different phases of colony growth. Then we describe the scheme of determining the uninvadable allocation schedule that consists of these possible phases.

#### 4.1 Conditions for candidate uninvadable allocation schedules

We now have all the elements to characterize the first-order conditions given by eqs. (S64) and (S65). We have from eqs. (S73) and (S76) that the conditions for the candidate optimal controls can be expressed as

$$u_{\mathrm{q}}^*(t) = \begin{cases} 0 & \text{if } u_{\mathrm{f}}^*(t) > 0 \text{ and } \sigma_1^c(t) < 0 \\ 1 & \text{if } u_{\mathrm{f}}^*(t) > 0 \text{ and } \sigma_1^c(t) > 0 \\ 0 < \hat{u}_{\mathrm{q}}^*(t) < 1 & \text{if } u_{\mathrm{f}}^*(t) > 0 \text{ and } \sigma_1^c(t) = 0 \\ 0 \le \tilde{u}_{\mathrm{q}}^*(t) \le 1 & \text{if } u_{\mathrm{f}}^*(t) = 0 \end{cases}$$

$$u_{\mathrm{f}}^*(t) = \begin{cases} 0 & \text{if } \left(u_{\mathrm{q}}^*(t) = 0 \text{ and } \sigma_2^c(t) > 0\right) \text{ or } \left(u_{\mathrm{q}}^*(t) = 1 \text{ and } \sigma_1^c(t) < \sigma_2^c(t)\right) \\ & \text{or } \left(u_{\mathrm{q}}^*(t) = \hat{u}_{\mathrm{q}}^*(t) \text{ and } \hat{u}_{\mathrm{q}}^*(t)\sigma_1^c(t) < \sigma_2^c(t)\right) \\ & \text{or } \left(u_{\mathrm{q}}^*(t) = \tilde{u}_{\mathrm{q}}^*(t) \text{ and } \tilde{u}_{\mathrm{q}}^*(t)\sigma_1^c(t) \le \sigma_1^c(t) < \sigma_2^c(t) \text{ and } \sigma_2^c(t) > 0\right) \\ 1 & \text{if } \left(u_{\mathrm{q}}^*(t) = 0 \text{ and } \sigma_2^c(t) < 0\right) \text{ or } \left(u_{\mathrm{q}}^*(t) = 1 \text{ and } \sigma_1^c(t) > \sigma_2^c(t)\right) \\ & \text{or } \left(u_{\mathrm{q}}^*(t) = \hat{u}_{\mathrm{q}}^*(t) \text{ and } \sigma_1^c(t) > \hat{u}_{\mathrm{q}}^*(t)\sigma_1^c(t) > \sigma_2^c(t)\right) \\ 0 < \hat{u}_{\mathrm{f}}^*(t) < 1 & \text{if } \left(u_{\mathrm{q}}^*(t) = 0 \text{ and } \sigma_2^c(t) = 0\right) \text{ or } \left(u_{\mathrm{q}}^*(t) = 1 \text{ and } \sigma_1^c(t) = \sigma_2^c(t)\right) \\ & \text{or } \left(\sigma_1^c(t) = \sigma_2^c(t) = 0\right) \text{ or } \left(u_{\mathrm{q}}^*(t) = \hat{u}_{\mathrm{q}}^*(t) \text{ and } \hat{u}_{\mathrm{q}}^*(t)\sigma_1^c(t) = \sigma_2^c(t)\right), \end{cases} \tag{S81}$$

where $\hat{u}_\tau^*(t)$ $\tau \in \{\mathrm{f}, \mathrm{q}\}$ denotes that the uninvadable control variable $u_\tau^*(t)$ is a singular arc and $\tilde{u}_{\mathrm{q}}^*(t)$ denotes that $u_{\mathrm{q}}^*(t)$ can not be determined and hence can take any value in the range $[0, 1]$ (reflecting the fact that during the phase when only males are produced the trait that affects how resources are allocated between different types of females does not have any effect on the allocation to individuals of different types). It follows from the conditions for the candidate optimal controls, given by eq. (S81), that there are seven possible phases of colony growth (see table S1). We will call phase W (exclusive worker production) the ergonomic phase of colony growth, while phases F (production of only queens), M (production of only males) and FM (production of queens and males simultaneously) are called the reproductive phases, since all the colony resources are directed towards producing sexuals. During phases WF, WM, FM, WFM there is simultaneous production of individuals of different types and in the context of our optimal control problem, it means that one or both of the control variables are singular arcs. In the following sections, we determine from the candidate optimal controls given by eq. (S81), the uninvadable allocation schedules for different assumptions of the time of dispersal of sexuals and different scenarios of genetic control.

| Regime | Individuals produced | $(u_{\mathrm{f}}^*(t), u_{\mathrm{q}}^*(t))$ | $\mathrm{sgn}(\sigma_1^c(t), \sigma_2^c(t), \sigma_1^c(t) - \sigma_2^c(t))$ |
|---|---|---|---|
| W | Workers | $(\bar{u}_{\mathrm{f}}^* = 1, \bar{u}_{\mathrm{q}}^* = 0)$ | $(-, -, \cdot)$ |
| F | Females (new queens) | $(\bar{u}_{\mathrm{f}}^* = 1, \bar{u}_{\mathrm{q}}^* = 1)$ | $(+, \cdot, +)$ |
| M | Males | $(\bar{u}_{\mathrm{f}}^* = 0, \tilde{u}_{\mathrm{q}}^*(t))$ | $(\cdot, +, -)$ |
| WF | Workers and females | $(\bar{u}_{\mathrm{f}}^* = 1, \hat{u}_{\mathrm{q}}^*(t))$ | $(0, -, +)$ |
| WM | Workers and males | $(\hat{u}_{\mathrm{f}}^*, \bar{u}_{\mathrm{q}}^* = 0)$ | $(-, 0, \cdot)$ |
| FM | Females and males | $(\hat{u}_{\mathrm{f}}^*(t), \bar{u}_{\mathrm{q}}^* = 1)$ | $(+, +, 0)$ |
| WFM | Workers, females and males | $(\hat{u}_{\mathrm{f}}^*(t), \hat{u}_{\mathrm{q}}^*(t))$ | $(0, 0, 0)$ |

Table S1: Candidate optimal controls and conditions for the signs of switching functions for all possible regimes of colony growth. Note that "$\cdot$" means any sign, $\bar{u}_\tau^*$ denotes that $u_\tau^*(t)$ is constant, $\hat{u}_\tau^*(t)$ denotes that $u_\tau^*(t)$ is a singular arc (if the singular arc is constant we write it as $\hat{\bar{u}}_\tau^*$) and $\tilde{u}_{\mathrm{q}}^*(t)$ denotes that $u_{\mathrm{q}}^*(t)$ is undetermined because no females are produced, since it does not appear in the Hamiltonian when $u_{\mathrm{f}}^*(t) = \bar{u}_{\mathrm{f}}^* = 0$.

#### 4.2 Short description of the derivations of the analytical results

We describe the scheme of obtaining the uninvadable allocation schedule $\mathbf{u}^*$ that consists of possible phases outlined in the table S1. The uninvadable allocation schedule $\mathbf{u}^*$ can explicitly be determined from the table S1, given that we know the switching functions $\sigma_1^c(t)$ and $\sigma_2^c(t)$ throughout the period $t \in [0, T]$. According to eq. (S77) the switching functions depend on the costate variables $\lambda_k^s(t)$ and the expected frequencies $p_c^s$ of the mutant allele in party $c$. Hence, establishing the uninvadable allocation schedule $\mathbf{u}^*$ reduces to determining the costate variables throughout the period $t \in [0, T]$. The dynamical behaviour of the costate variables is given backwards in time by the differential equations in eq. (S71) and the terminal conditions (transversality conditions) in eq. (S72). During candidate phases W, F, and M of colony growth, the allocation variables $u_{\mathrm{f}}^*(t) = \bar{u}_{\mathrm{f}}^*$ and $u_{\mathrm{q}}^*(t) = \bar{u}_{\mathrm{q}}^*$ are constant. Hence, it will turn out to be useful to derive the equations for costate variables from eq. (S71), assuming that $u_{\mathrm{f}}^*(t) = \bar{u}_{\mathrm{f}}^*$ and $u_{\mathrm{q}}^*(t) = \bar{u}_{\mathrm{q}}^*$ are constant (see eqs. S90–S91). The costate variables at time $t = T$ are given by the transversality conditions (S83), (S86), and (S87), which deduce from eq. (S72). In addition, we derive eqs. (S88) and (S89) that describe the dynamical behaviour of the state and dynamic variables, respectively, which are solutions to eqs. (S57), (S58), (S57), and (S58) assuming that $u_{\mathrm{f}}^*(t) = \bar{u}_{\mathrm{f}}^*$ and $u_{\mathrm{q}}^*(t) = \bar{u}_{\mathrm{q}}^*$ are constant. These expressions together with table S1 give the necessary elements to obtain the uninvadable allocation schedule $\mathbf{u}^*$.

We then proceed by determining the switching functions $\sigma_1^c(t)$ and $\sigma_2^c(t)$ backwards in time. At time $t = T$, the switching functions can be directly determined by substituting the transversality conditions (S83), (S86), and (S87) into eq. (S77). Considering that the costate variables are continuous, then it follows that we can determine $\mathbf{u}^*(t)$ for some time interval $t \in [t_{c,i}^*, T]$ before the end of the season, where $t = t_{c,i}^*$ is the time for which one or more of the switching functions (S77) change their signs. Here, the subscript "$c$" emphasizes the scenario of genetic control and "$i$" denotes the number of switches between different phases that make up the uninvadable allocation schedule $\mathbf{u}^*$. Hereby, we have established the uninvadable allocation strategy $\mathbf{u}^*(t)$ for the last phase $t \in [t_{c,i}^*, T]$. We proceed by using information about $\mathbf{u}^*(t)$ during the last phase and determining the switching functions $\sigma_1^c(t)$ and $\sigma_2^c(t)$ for the penultimate phase $t \in [t_{c,i-1}^*, t_{c,i}^*]$. We iterate this process, until the switching functions will not change their signs any more. This scheme allows us to determined all the $i+1$ phases of the uninvadable allocation schedule $\mathbf{u}^*$ and all the switching times ($t_{c,1}^*$,...,$t_{c,i}^*$) between the different phases.

Since the properties of the uninvadable resource allocation strategies under delayed dispersal and direct dispersal turn out to be substantially different, then we will present the derivations for these two cases separately. For delayed dispersal, we were able to derive the uninvadable resource allocation schedule analytically only assuming equal mortality of sexuals $\mu_{\mathrm{q}} = \mu_{\mathrm{m}} = \mu_{\mathrm{r}}$. For direct dispersal, we were able to derive the full uninvadable resource allocation schedule analytically under single party control only for when $R_c\mu_{\mathrm{q}} \geq \mu_{\mathrm{m}}$ and under mixed control only for when $\mu_{\mathrm{q}} = \mu_{\mathrm{m}}$.

Our analytical results are in accordance with the numerical results derived with GPOPS that employs the direct approach of finding the uninvadable allocation schedule instead of the indirect approach given by Pontryagin's maximum principle. Note that the derivation of some of our analytical results (for direct dispersal) entails using the intuition from our numerical results about the general properties of the optimal allocation schedule and then verifying that this solution satisfies the first-order conditions for uninvadability (eqs. S64–S65).

##### 4.3 State and costate variables

We outlined in section 4.2 that in order to determine the univadable allocation schedule $\mathbf{u}^*$ from the first-order condition, we need to derive the costate equations, transversality conditions and the equations that describe the dynamics of the costate and the state variables for a period of time $[t_0, t_1]$ during which the resource allocation is constant. We now present these expressions.

###### 4.3.1 Costate equations and transversality conditions

Substitution of eqs. (S58) and (S67) into eq. (S71) yields the following differential equations for the costate variables

$$\begin{aligned}
\dot{\lambda}^s_{\mathrm{w}}(t) &= -\left[\lambda^s_{\mathrm{w}}(t)(ba^*_{\mathrm{w}}(t)-\mu_{\mathrm{w}}(t)) + \lambda^s_{\mathrm{q}}(t)ba^*_{\mathrm{q}}(t) + \lambda^s_{\mathrm{m}}(t)ba^*_{\mathrm{m}}(t)\right], \\
\dot{\lambda}^s_{\mathrm{q}}(t) &= \mu_{\mathrm{q}}\lambda^s_{\mathrm{q}}(t), \\
\dot{\lambda}^s_{\mathrm{m}}(t) &= \begin{cases} \mu_{\mathrm{m}}\lambda^s_{\mathrm{m}}(t) & \text{(under delayed dispersal)}, \\ \mu_{\mathrm{m}}\lambda^s_{\mathrm{m}}(t) - \frac{ba^*_{\mathrm{q}}(t)x^*_{\mathrm{w}}(t)}{x^*_{\mathrm{m}}(t)}\lambda^s_{\mathrm{iq}}(t) & \text{(under direct dispersal)}, \end{cases} \\
\dot{\lambda}^s_{\mathrm{iq}}(t) &= \mu_{\mathrm{q}}\lambda^s_{\mathrm{iq}}(t),
\end{aligned} \tag{S82}$$

where $a^*_{\mathrm{w}}(t) = u^*_{\mathrm{f}}(t)(1-u^*_{\mathrm{q}}(t))$, $a^*_{\mathrm{q}}(t) = u^*_{\mathrm{f}}(t)u^*_{\mathrm{q}}(t)$, and $a^*_{\mathrm{m}}(t) = (1-u^*_{\mathrm{f}}(t))$. The terminal conditions for these differential equations (S82) are given by the transversality conditions (S72).

Because the number of workers does not appear in the expression of invasion fitness, we have, regardless of the mode of control of traits, that

$$\lambda^s_{\mathrm{w}}(T) = 0 \text{ for } s \in \{\mathrm{q},\mathrm{m}\}. \tag{S83}$$

Otherwise, we have from the perturbation formula for eigenvalues (eq. S18 and S72) that

$$\begin{aligned}
\lambda^s_k(T) &= \nu^\circ_k \gamma_{ks} \left.\frac{\partial w_{ks}(\mathbf{u}^s,\mathbf{v})}{\partial x^s_k(T)}\right|_{\mathbf{x}=\mathbf{y}=\mathbf{x}^*} q^\circ_s \text{ for } k \in \{\mathrm{q},\mathrm{m}\} \text{ and } s \in \{\mathrm{q},\mathrm{m}\}, \\
\lambda^s_{\mathrm{iq}}(T) &= \nu^\circ_{\mathrm{m}} \gamma_{\mathrm{m}s} \left.\frac{\partial w_{\mathrm{m}s}(\mathbf{u}^s,\mathbf{v})}{\partial x^s_{\mathrm{iq}}(T)}\right|_{\mathbf{x}=\mathbf{y}=\mathbf{x}^*} q^\circ_s \text{ for } s \in \{\mathrm{q},\mathrm{m}\}.
\end{aligned} \tag{S84}$$

Furthermore $x^s_k$ affects only the component $w_{ks}(\mathbf{u}^s,\mathbf{v})$ of invasion fitness for $k \in \{\mathrm{q},\mathrm{m}\}$ and $x^s_{\mathrm{iq}}$ only affects the component $w_{\mathrm{m}s}(\mathbf{u}^s,\mathbf{v})$. Hence, owing to eq. (S20) and the fitness functions (eqs. S6–S7), we have

$$\begin{aligned}
&\lambda^s_k(T) = \frac{\alpha^\circ_k \gamma_{ks}}{x^*_k(\mathbf{u}^*)} && \text{(under delayed dispersal)}, \\
&\lambda^s_{\mathrm{q}}(T) = \frac{\alpha^\circ_{\mathrm{q}} \gamma_{\mathrm{q}s}}{x^*_{\mathrm{q}}(\mathbf{u}^*)},\ \lambda^s_{\mathrm{m}}(T) = 0 \text{ and } \lambda^s_{\mathrm{iq}}(T) = \frac{\alpha^\circ_{\mathrm{m}} \gamma_{\mathrm{m}s}}{x^*_{\mathrm{q}}(\mathbf{u}^*)} && \text{(under direct dispersal)}.
\end{aligned} \tag{S85}$$

For haplodiploids, $\gamma_{\mathrm{qq}} = 1/2$, $\gamma_{\mathrm{mq}} = 1$, $\gamma_{\mathrm{qm}} = 1/2$ and $\gamma_{\mathrm{mm}} = 0$ and consequently eq. (S21) yields that $\alpha_{\mathrm{q}} = 2/3$ and $\alpha_{\mathrm{m}} = 1/3$. Substituting these parameters into eq. (S85) yields the following transversality conditions for sexuals under delayed dispersal

$$
\begin{aligned}
&\lambda_{\mathrm{q}}^{\mathrm{q}}(T)=\frac{1}{3x_{\mathrm{q}}^{*}(\mathbf{u}^{*})}\,,\lambda_{\mathrm{m}}^{\mathrm{q}}(T)=\frac{1}{3x_{\mathrm{m}}^{*}(\mathbf{u}^{*})}\,,\\
&\lambda_{\mathrm{q}}^{\mathrm{m}}(T)=0\,,\lambda_{\mathrm{m}}^{\mathrm{m}}(T)=0 \qquad\qquad (\mathbf{u}_{\mathcal{T}},\,\mathbf{v}_{\mathcal{T}}\text{ under queen control});\\
&\lambda_{\mathrm{q}}^{\mathrm{q}}(T)=\frac{1}{3x_{\mathrm{q}}^{*}(\mathbf{u}^{*})}\,,\lambda_{\mathrm{m}}^{\mathrm{q}}(T)=\frac{1}{3x_{\mathrm{m}}^{*}(\mathbf{u}^{*})}\,,\\
&\lambda_{\mathrm{q}}^{\mathrm{m}}(T)=\frac{1}{3x_{\mathrm{q}}^{*}(\mathbf{u}^{*})}\,,\lambda_{\mathrm{m}}^{\mathrm{m}}(T)=0 \qquad\qquad (\mathbf{u}_{\mathcal{T}},\,\mathbf{v}_{\mathcal{T}}\text{ under worker control})
\end{aligned}
\tag{S86}
$$

and under direct dispersal the transversality conditions are given by

$$
\begin{aligned}
&\lambda_{\mathrm{q}}^{\mathrm{q}}(T)=\frac{1}{3x_{\mathrm{q}}^{*}(\mathbf{u}^{*})}\,,\lambda_{\mathrm{m}}^{\mathrm{q}}(T)=0\,,\lambda_{\mathrm{q}}^{\mathrm{m}}(T)=0\,,\lambda_{\mathrm{m}}^{\mathrm{m}}(T)=0\,,\\
&\lambda_{\mathrm{iq}}^{\mathrm{q}}(T)=\frac{1}{3x_{\mathrm{q}}^{*}(\mathbf{u}^{*})}\,,\lambda_{\mathrm{iq}}^{\mathrm{m}}(T)=0 \qquad\qquad (\mathbf{u}_{\mathcal{T}},\,\mathbf{v}_{\mathcal{T}}\text{ under queen control});\\
&\lambda_{\mathrm{q}}^{\mathrm{q}}(T)=\frac{1}{3x_{\mathrm{q}}^{*}(\mathbf{u}^{*})}\,,\lambda_{\mathrm{m}}^{\mathrm{q}}(T)=0\,,\lambda_{\mathrm{q}}^{\mathrm{m}}(T)=\frac{1}{3x_{\mathrm{q}}^{*}(\mathbf{u}^{*})}\,,\lambda_{\mathrm{m}}^{\mathrm{m}}(T)=0\,,\\
&\lambda_{\mathrm{iq}}^{\mathrm{q}}(T)=\frac{1}{3x_{\mathrm{q}}^{*}(\mathbf{u}^{*})}\,,\lambda_{\mathrm{iq}}^{\mathrm{m}}(T)=0 \qquad\qquad (\mathbf{u}_{\mathcal{T}},\,\mathbf{v}_{\mathcal{T}}\text{ under worker control}).
\end{aligned}
\tag{S87}
$$

###### 4.3.2 Solutions to the state and costate equations

Let the upper bar denote that the variable is constant. Hence, let $\bar{a}_{k,u}^{s}$ denote constant proportional allocation to individuals of type $k$ in a colony founded by a mutant individual of type $s$ during a given time interval $[t_0, t_1]$. The solutions to the differential equations (S57)–(S58) that give the expressions for the state variables at time $t \in [t_0, t_1]$ are

$$
\begin{aligned}
x_{\mathrm{w}}^{s}(t)&=x_{\mathrm{w}}^{s}(t_0)e^{(t-t_0)\left(b\bar{a}_{\mathrm{w,u}}^{s}-\mu_{\mathrm{w}}\right)},\\
x_{\mathrm{q}}^{s}(t)&=\frac{e^{-\mu_{\mathrm{q}}(t-t_0)}\left[b\bar{a}_{\mathrm{q,u}}^{s}x_{\mathrm{w}}^{s}(t_0)\left(e^{(t-t_0)\left(b\bar{a}_{\mathrm{w,u}}^{s}+\mu_{\mathrm{q}}-\mu_{\mathrm{w}}\right)}-1\right)+x_{\mathrm{q}}^{s}(t_0)\left(b\bar{a}_{\mathrm{w,u}}^{s}+\mu_{\mathrm{q}}-\mu_{\mathrm{w}}\right)\right]}{b\bar{a}_{\mathrm{w,u}}^{s}+\mu_{\mathrm{q}}-\mu_{\mathrm{w}}},\\
x_{\mathrm{m}}^{s}(t)&=\frac{e^{-\mu_{\mathrm{m}}(t-t_0)}\left[b\bar{a}_{\mathrm{m,u}}^{s}x_{\mathrm{w}}^{s}(t_0)\left(e^{(t-t_0)\left(b\bar{a}_{\mathrm{w,u}}^{s}+\mu_{\mathrm{m}}-\mu_{\mathrm{w}}\right)}-1\right)+x_{\mathrm{m}}^{s}(t_0)\left(b\bar{a}_{\mathrm{w,u}}^{s}+\mu_{\mathrm{m}}-\mu_{\mathrm{w}}\right)\right]}{b\bar{a}_{\mathrm{w,u}}^{s}+\mu_{\mathrm{m}}-\mu_{\mathrm{w}}},\\
x_{\mathrm{iq}}^{s}(t)&=x_{\mathrm{iq}}^{s}(t_0)+\int_{t_0}^{t}x_{\mathrm{m}}^{s}(\tau)\frac{ba_{\mathrm{q}}^{s}(\tau)y_{\mathrm{w}}(\tau)}{y_{\mathrm{m}}(\tau)}e^{-\mu_{\mathrm{q}}(t-\tau)}d\tau,
\end{aligned}
\tag{S88}
$$

Let $\bar{a}_k$ denote the constant proportional allocation to individuals of type $k$ in a resident colony during a given time interval $[t_0, t_1]$. The solutions to differential equations (S59)–(S60) that give the expressions for the dynamic variables at time $t \in [t_0, t_1]$ are

$$
\begin{aligned}
y_{\mathrm{w}}(t) &= y_{\mathrm{w}}(t_0) e^{(t-t_0)(b\bar{a}_{\mathrm{w}}-\mu_{\mathrm{w}})}, \\
y_{\mathrm{q}}(t) &= \frac{e^{-\mu_{\mathrm{q}}(t-t_0)}\left[b\bar{a}_{\mathrm{q}} y_{\mathrm{w}}(t_0)\left(e^{(t-t_0)(b\bar{a}_{\mathrm{w}}+\mu_{\mathrm{q}}-\mu_{\mathrm{w}})}-1\right)+y_{\mathrm{q}}(t_0)\left(b\bar{a}_{\mathrm{w}}+\mu_{\mathrm{q}}-\mu_{\mathrm{w}}\right)\right]}{b\bar{a}_{\mathrm{w}}+\mu_{\mathrm{q}}-\mu_{\mathrm{w}}}, \\
y_{\mathrm{m}}(t) &= \frac{e^{-\mu_{\mathrm{m}}(t-t_0)}\left[b\bar{a}_{\mathrm{m}} y_{\mathrm{w}}(t_0)\left(e^{(t-t_0)(b\bar{a}_{\mathrm{w}}+\mu_{\mathrm{m}}-\mu_{\mathrm{w}})}-1\right)+y_{\mathrm{m}}(t_0)\left(b\bar{a}_{\mathrm{w}}+\mu_{\mathrm{m}}-\mu_{\mathrm{w}}\right)\right]}{b\bar{a}_{\mathrm{w}}+\mu_{\mathrm{m}}-\mu_{\mathrm{w}}}.
\end{aligned}
\tag{S89}
$$

Let $\bar{a}_k^*$ denote a constant uninvadable proportional allocation to individuals of type $k$ during a given time interval $[t_0, t]$. The solutions to differential equations eq. (S82) that give the expressions for the costate variables at time $t \in [t_0, t_1]$ are

$$
\begin{aligned}
\lambda_{\mathrm{w}}^s(t) &= \lambda_{\mathrm{w}}^s(t_1) e^{(b\bar{a}_{\mathrm{w}}^*-\mu_{\mathrm{w}})(t_1-t)} + \sum_{k\in\{\mathrm{q},\mathrm{m}\}} \lambda_k^s(t_1) \frac{b\bar{a}_k^*}{b\bar{a}_{\mathrm{w}}^*-\mu_{\mathrm{w}}+\mu_k}\left(e^{(b\bar{a}_{\mathrm{w}}^*-\mu_{\mathrm{w}})(t_1-t)} - e^{-\mu_k(t_1-t)}\right), \\
\lambda_{\mathrm{q}}^s(t) &= \lambda_{\mathrm{q}}^s(t_1) e^{-\mu_{\mathrm{q}}(t_1-t)}, \\
\lambda_{\mathrm{m}}^s(t) &= \begin{cases} \lambda_{\mathrm{m}}^s(t_1) e^{-\mu_{\mathrm{m}}(t_1-t)} & \text{(delayed dispersal)}, \\ \lambda_{\mathrm{m}}^s(t_1) e^{-\mu_{\mathrm{m}}(t_1-t)} + \lambda_{\mathrm{iq}}^s(t_1) e^{\mu_{\mathrm{m}} t-\mu_{\mathrm{q}} t_1} \displaystyle\int_t^{t_1} e^{(\mu_{\mathrm{q}}-\mu_{\mathrm{m}})\tau} \frac{b a_{\mathrm{q}}^*(\tau) x_{\mathrm{w}}^*(\tau)}{x_{\mathrm{m}}^*(\tau)} d\tau & \text{(direct dispersal)}, \end{cases} \\
\lambda_{\mathrm{iq}}^s(t) &= \lambda_{\mathrm{iq}}^s(t_1) e^{-\mu_{\mathrm{q}}(t_1-t)} \quad .
\end{aligned}
\tag{S90}
$$

Note that the solutions given by eq. (S90) to costate equations hold for $\lambda_{\mathrm{q}}^s(t)$, $\lambda_{\mathrm{q}}^s(t)$, $\lambda_{\mathrm{q}}^s(t)$ even if the uninvadable allocation schedule $\mathbf{u}^*(t)$ is not constant during $[t_0, t]$. As opposed to state variables, the dynamics of costate variables is described backwards in time, where $\lambda_{\mathrm{m}}^s(t_1)$ is the terminal condition.The transversality conditions (S86)-(S87) together with eq. (S90) imply that

$$
\begin{aligned}
\lambda_{\mathrm{m}}^{\mathrm{m}}(t) = 0,\ \lambda_{\mathrm{q}}^{\mathrm{q}}(t) = \lambda_{\mathrm{q}}^{\mathrm{m}}(t), &\quad \forall\, t \in [0,T], \quad \text{(delayed dispersal)}, \\
\lambda_{\mathrm{m}}^{\mathrm{m}}(t) = \lambda_{\mathrm{iq}}^{\mathrm{m}}(t) = 0,\ \lambda_{\mathrm{q}}^{\mathrm{q}}(t) = \lambda_{\mathrm{q}}^{\mathrm{m}}(t) = \lambda_{\mathrm{iq}}^{\mathrm{q}}(t), &\quad \forall\, t \in [0,T], \quad \text{(direct dispersal)}.
\end{aligned}
\tag{S91}
$$

### 5 The candidate uninvadable allocation schedule under delayed dispersal

#### 5.1 Equal male and female mortality ($\mu_{\mathrm{q}} = \mu_{\mathrm{m}}$)

Here we show analytically that the candidate uninvadable allocation schedule under delayed dispersal (assuming equal mortality rates of sexuals, i.e. $\mu_{\mathrm{q}} = \mu_{\mathrm{m}} = \mu_{\mathrm{r}}$) consists of only two growth regimes: (i) the ergonomic phase (regime W) and (ii) the reproductive phase, where males and new queens are produced simultaneously (regime FM). The uninvadable allocation schedule can be described by phases W and FM of table S1:

$$
\begin{aligned}
&\textbf{Phase W:} \quad (u_{\mathrm{f}}^*(t), u_{\mathrm{q}}^*(t)) = (\bar{u}_{\mathrm{f}}^* = 1, \bar{u}_{\mathrm{q}}^* = 0) \quad \text{for } t \in \left[0, t_{c,1}^*\right]; \\
&\textbf{Phase FM:} \quad (u_{\mathrm{f}}^*(t), u_{\mathrm{q}}^*(t)) = (\hat{\bar{u}}_{\mathrm{f}}^*, \bar{u}_{\mathrm{q}}^* = 1) \quad \text{for } t \in \left[t_{c,1}^*, T\right].
\end{aligned}
\tag{S92}
$$

The uninvadable allocation schedule has this form regardless of the assumptions about the genetic control of the resource allocation traits. However, the switching time $t^*_{c,1}$ from the ergonomic phase to the reproductive phase depends on the scenario $c$ of the genetic control of the resource allocation traits. We will proceed according to the scheme for deriving the uninvadable allocation schedule $\mathbf{u}^*$ outlined in section 4.2.

##### 5.1.1 Regime FM: $\sigma^c_1(t) > 0$, $\sigma^c_2(t) > 0$, and $\sigma^c_1(t) - \sigma^c_2(t) = 0$

We know from biological considerations that $x^*_{\mathrm{q}}(\mathbf{u}^*) > 0$ and $x^*_{\mathrm{m}}(\mathbf{u}^*) > 0$. Taking this into account and substituting the transversality conditions (S83) and (S86) into the switching functions (S78) (for queen control) and (S80) (for worker control), implies that $\sigma^c_1(T) > 0$ and $\sigma^c_2(T) > 0$. According to table S1, we need to also determine the sign of $\sigma^c_1(T) - \sigma^c_2(T)$. Assuming equal mortality rates of males and queens ($\mu_{\mathrm{q}} = \mu_{\mathrm{m}} = \mu_{\mathrm{r}}$) and substituting eq. (S86) into eqs. (S78) (for queen control) and (S80) (for worker control) yields

$$\begin{aligned} \sigma^{\mathrm{q}}_1(T) - \sigma^{\mathrm{q}}_2(T) &= \frac{1}{6}\left(\frac{1}{x^*_{\mathrm{q}}(\mathbf{u}^*)} - \frac{1}{x^*_{\mathrm{m}}(\mathbf{u}^*)}\right) && (\mathbf{u}_{\mathrm{f}}, \mathbf{v}_{\mathrm{f}} \text{ under queen control}), \\ \sigma^{\mathrm{w}}_1(T) - \sigma^{\mathrm{w}}_2(T) &= \frac{1}{12}\left(\frac{2+M}{M}\frac{1}{x^*_{\mathrm{q}}(\mathbf{u}^*)} - \frac{1}{x^*_{\mathrm{m}}(\mathbf{u}^*)}\right) && (\mathbf{u}_{\mathrm{f}}, \mathbf{v}_{\mathrm{f}} \text{ under worker control}). \end{aligned} \tag{S93}$$

We know from eq. (S35) that the first-order condition implies that $x^*_{\mathrm{q}}(\mathbf{u}^*)/x^*_{\mathrm{m}}(\mathbf{u}^*) = R_c$, where $R_c = 1$ for queen control and $R_c = (2+M)/M$ for worker control (recall S35), which, on substitution, implies that

$$\sigma^c_1(T) - \sigma^c_2(T) = 0 \tag{S94}$$

for both queen and worker control of the focal trait of type $\tau$.

Hence, we have shown that $\sigma^c_1(T) > 0$, $\sigma^c_2(T) > 0$, $\sigma^c_1(T) - \sigma^c_2(T) = 0$. This implies via table S1 that $u^*_{\mathrm{q}}(t) = \bar{u}^*_{\mathrm{q}} = 1$ during the final growth regime. It also follows from eq. (S81) and $\sigma^c_1(T) - \sigma^c_2(T) = 0$, that $u^*_{\mathrm{f}}(t)$ might be a singular arc during the final growth regime. More precisely, for $u^*_{\mathrm{f}}(t) = \hat{u}^*_{\mathrm{f}}(t)$ to be a singular arc,

$$\left.\frac{\partial H_{c,d}(\mathbf{u}(t), \mathbf{x}^*(t), \boldsymbol{\lambda}(t))}{\partial u_{\mathrm{f}}(t)}\right|_{\mathbf{u}=\mathbf{v}=\mathbf{u}^*} = b x^*_{\mathrm{w}}(t)\left(u^*_{\mathrm{q}}(t)\sigma^c_1(t) - \sigma^c_2(t)\right) = 0 \tag{S95}$$

must hold for a finite interval of time (e.g., Bryson and Ho, 1975, p. 248). It follows, that also higher order time derivatives have to vanish along the singular arc and this condition can be used to determine $u^*_{\mathrm{f}}(t) = \hat{u}^*_{\mathrm{f}}(t)$, i.e. the constraints for the singular arc are given by the sequence

$$\left(\frac{\mathrm{d}}{\mathrm{d}t}\right)^i \left.\frac{\partial H_{c,d}(\mathbf{u}(t), \mathbf{x}^*(t), \boldsymbol{\lambda}(t))}{\partial u_{\mathrm{f}}(t)}\right|_{\mathbf{u}=\mathbf{v}=\mathbf{u}^*} = \left(\frac{\mathrm{d}}{\mathrm{d}t}\right)^i b x^*_{\mathrm{w}}(t)\left(u^*_{\mathrm{q}}(t)\sigma^c_1(t) - \sigma^c_2(t)\right) = 0 \qquad i = 0, 1, 2, .... \tag{S96}$$

Note that $b > 0$ for biological reasons and $u^*_{\mathrm{q}}(t) = \bar{u}^*_{\mathrm{q}} = 1$ during the last growth regime. It follows from eq. (S88) (by taking $\bar{u}^*_{\mathrm{q}} = 1$, $t_0 = t^*_{c,1}$, and $t_1 = T$) that $x^*_{\mathrm{w}}(t) = x^*_{\mathrm{w}}(t^*_{c,1})\exp(-\mu_{\mathrm{w}}(t - t^*_{c,1}))$ during the last growth regime. Since, $x^*_{\rm w}(t^*_{c,1}) > 0$, then eq. (S96) simplifies to

$$\left(\frac{\rm d}{{\rm d}t}\right)^i (\sigma_1^c(t) - \sigma_2^c(t)) = 0 \qquad i = 0, 1, 2, \ldots . \tag{S97}$$

Substituting the switching functions eq. (S78) (for queen control) and (S80) (for worker control) into eq. (S97) implies for $i = 1$ that on a singular arc

$$\begin{aligned} \dot\lambda^{\rm q}_{\rm q}(t) &= \dot\lambda^{\rm q}_{\rm m}(t) \quad (\mathbf{u}_\tau,\, \mathbf{v}_\tau \text{ under queen control}),\\ \frac{1}{4}\dot\lambda^{\rm q}_{\rm q}(t) + \frac{1}{2M}\dot\lambda^{\rm m}_{\rm q}(t) &= \frac{1}{4}\dot\lambda^{\rm q}_{\rm m}(t) + \frac{1}{2M}\dot\lambda^{\rm m}_{\rm m}(t) \quad (\mathbf{u}_\tau,\, \mathbf{v}_\tau \text{ under worker control}).\end{aligned} \tag{S98}$$

must hold. Simplifying eq. (S98) by using eq. (S91); namely, $\dot\lambda^{\rm q}_{\rm q}(t) = \dot\lambda^{\rm m}_{\rm q}(t)$ and $\dot\lambda^{\rm m}_{\rm m}(t) = 0$, and further using the expression for relatedness asymmetry $R_c$ given by eq. (S35), we obtain

$$\frac{\dot\lambda^{\rm q}_{\rm m}(t)}{\dot\lambda^{\rm q}_{\rm q}(t)} = R_c, \tag{S99}$$

where $R_c$ is the relatedness asymmetry associated with the party $c \in \{{\rm q}, {\rm w}\}$ in control of the trait of type f. Substituting the differential equations for costate variables (S82) into eq. (S99) gives

$$\frac{\lambda^{\rm q}_{\rm m}(t)}{\lambda^{\rm q}_{\rm q}(t)} = \frac{\mu_{\rm q}}{\mu_{\rm m}} R_c. \tag{S100}$$

Note that the control variable $u_{\rm f}(t)$ itself does not appear in eq. (S100). Furthermore, it follows from the simple form of the costate equations (S82) that the $i$-th order time derivative of the coefficient $(\sigma_1^c(t) - \sigma_2^c(t))$ produces

$$\frac{\lambda^{\rm q}_{\rm m}(t)}{\lambda^{\rm q}_{\rm q}(t)} = \left(\frac{\mu_{\rm q}}{\mu_{\rm m}}\right)^i R_c. \tag{S101}$$

Since the control variable does $u^*_{\rm f}(t)$ not appear in any order time derivative of the coefficient $(\sigma_1^c(t) - \sigma_2^c(t))$, we can use the constraints (S101) that the singular arc has to satisfy to indirectly obtain the expression for the singular arc $\hat u^*_{\rm f}(t)$. Substituting the costate equations (S90) (and setting $t_1 = T$) into eq. (S101), using the transversality conditions given by eq. (S86), and assuming that the mortality rates of queens and males are equal ($\mu_{\rm q} = \mu_{\rm m}$) yields

$$\frac{x^*_{\rm q}(\mathbf{u}^*)}{x^*_{\rm m}(\mathbf{u}^*)} = R_c. \tag{S102}$$

Here, we have recovered the critical sex ratio given by eq. (S50), which essentially implies that if males and queens have equal mortality rates, there are no additional dynamic constraints besides the critical sex ratio at the end of the season. Since, there are no additional dynamic constraints for the control variable, we assume (for simplicity) that it is constant, i.e. $\hat u^*_{\rm f}(t) = \bar u^*_{\rm f}$. This allows us to substitute the equations for state variables for constant allocation strategy ($\hat u^*_{\rm f}$,$\bar u^*_{\rm q}$=1), given by eq. (S88) with eq. (3) into eq. (S102) for the time interval

$t \in [t^*_{c,1}, T]$ and assuming that $\mu_\mathrm{q} = \mu_\mathrm{m} = \mu_\mathrm{r}$ and $x^*_\mathrm{q}(t^*_{c,1}) = x^*_\mathrm{m}(t^*_{c,1}) = 0$ (at the start of the reproductive phase, there are no males or juvenile queens), we obtain

$$\begin{aligned}\frac{x^*_\mathrm{m}(\mathbf{u}^*)}{x^*_\mathrm{q}(\mathbf{u}^*)} &= \frac{b(1-\hat{u}^*_\mathrm{f} x^*_\mathrm{w}(t^*_{c,1})e^{-\mu_\mathrm{r}\left(T-t^*_{c,1}\right)}\left[e^{\left(T-t^*_{c,1}\right)(\mu_\mathrm{r}-\mu_\mathrm{w})}-1\right](\mu_\mathrm{r}-\mu_\mathrm{w})}{b\hat{u}^*_\mathrm{f} x^*_\mathrm{w}(t^*_{c,1})e^{-\mu_\mathrm{r}\left(T-t^*_{c,1}\right)}\left[\left(e^{\left(T-t^*_{c,1}\right)(\mu_\mathrm{r}-\mu_\mathrm{w})}-1\right)\right](\mu_\mathrm{r}-\mu_\mathrm{w})}, \\ &= \frac{(1-\hat{u}^*_\mathrm{f})}{\hat{u}^*_\mathrm{f}} = R_c.\end{aligned} \tag{S103}$$

Solving for $\hat{u}^*_\mathrm{f}$ yields

$$\hat{u}^*_\mathrm{f} = \begin{cases} \dfrac{R_\mathrm{q}}{1+R_\mathrm{q}} & (\mathbf{u}_\mathrm{f},\, \mathbf{v}_\mathrm{f} \text{ under queen control}), \\ \dfrac{R_\mathrm{w}}{1+R_\mathrm{w}} & (\mathbf{u}_\mathrm{f},\, \mathbf{v}_\mathrm{f} \text{ under worker control}) \end{cases} \tag{S104}$$

and for haplodiploids it simplifies to

$$\hat{u}^*_\mathrm{f} = \begin{cases} \dfrac{1}{2} & (\mathbf{u}_\mathrm{f},\, \mathbf{v}_\mathrm{f} \text{ under queen control})), \\ \dfrac{2+M}{2(1+M)} & (\mathbf{u}_\mathrm{f},\, \mathbf{v}_\mathrm{f} \text{ under worker control}). \end{cases} \tag{S105}$$

Hence, we have determined the uninvadable allocation schedule $\mathbf{u}^*(t)$ for the final growth regime FM ($t \in [t^*_{c,1}, T]$), assuming equal mortality rates of males and queens.

##### 5.1.2 Regime W: $\sigma^c_1(t) < 0$, $\sigma^c_2(t) < 0$, and $\sigma^c_1(t) - \sigma^c_2(t) = 0$

In order to determine the preceding phase, we need to determine which switching function expression $\sigma^c_1(t) < 0$, $\sigma^c_2(t) < 0$, and $\sigma^c_1(t) - \sigma^c_2(t) = 0$ changes their sign.

Lets first examine expression $\sigma^c_1(t) - \sigma^c_2(t) = 0$. Substituting the costate variables given by eq. (S90) (for the time interval $t \in [0, T]$) into the expression $\sigma^c_1(t) - \sigma^c_2(t)$ given by eq. (S78) (for queen control) and eq. (S80) (for worker control) yields for any $t \in [0, T]$

$$\begin{aligned}\sigma^\mathrm{q}_1(t) - \sigma^\mathrm{q}_2(t) &= e^{-\mu_\mathrm{r}(T-t)}\frac{1}{2}\left(\lambda^\mathrm{q}_\mathrm{q}(T) - \lambda^\mathrm{q}_\mathrm{m}(T)\right) && (\mathbf{u}_\mathrm{f},\, \mathbf{v}_\mathrm{f} \text{ under queen control}), \\ \sigma^\mathrm{w}_1(t) - \sigma^\mathrm{w}_2(t) &= e^{-\mu_\mathrm{r}(T-t)}\frac{1}{4}\left(\frac{2+M}{M}\lambda^\mathrm{q}_\mathrm{q}(T) - \lambda^\mathrm{q}_\mathrm{m}(T)\right) && (\mathbf{u}_\mathrm{f},\, \mathbf{v}_\mathrm{f} \text{ under worker control}).\end{aligned} \tag{S106}$$

Notice that from the expression $\sigma^c_1(t) - \sigma^c_2(t)$ given by eqs. (S78) (for queen control) and (S80) (for worker control) that eq. (S106) can also be expressed in terms of

$$\sigma^c_1(t) - \sigma^c_2(t) = e^{-\mu_\mathrm{r}(T-t)}\left[\sigma^c_1(T) - \sigma^c_2(T)\right]. \tag{S107}$$

It follows from eq. (S94) that $\sigma^c_1(t) - \sigma^c_2(t) = 0$ throughout the entire time interval $t \in [0, T]$ for both queen and worker control.

Next, we will show that there is a switch from regime FM to regime W. If there exists at least one root of $t$ in equation $\sigma_1^c(t) = 0$, given that the costate variables in $\sigma_1^c(t)$ are obtained through integrating eq. (S82) over the final growth regime $[t^*_{c,1}, T]$, assuming that $\mathbf{u}^*(t) = (\bar{u}^*_{\mathrm{f}} = \hat{u}^*_{\mathrm{f}}, \bar{u}^*_{\mathrm{q}} = 1)$, then there is a switch in $u_{\mathrm{q}}(t)$ and $u_{\mathrm{f}}(t)$ (since $\sigma_2^c(t)$ also changes its sign because $\sigma_1^c(t) = \sigma_2^c(t)\ \forall t \in [0, T]$). The switching time $t^*_{c,1} < T$ from phase FM to phase W is given by the largest root of $\sigma_1^c(t) = 0$ (assuming that $\mathbf{u}^*(t) = (\bar{u}^*_{\mathrm{f}} = \hat{u}^*_{\mathrm{f}}, \bar{u}^*_{\mathrm{q}} = 1)$ in the last phase) and its existence is shown in sections 5.1.3 for single-party control and 5.1.4 for mixed control, respectively. We can infer from our numerical solutions that $u_{\mathrm{q}}$ is not a singular arc. Hence, $\sigma_1^c(t) = \sigma_2^c(t) = 0$ only at time $t = t^*_{c,1}$ and $\sigma_1^c(t) = \sigma_2^c(t) < 0$ for $t < t^*_{c,1}$. Hence, we have determined that phase W (with allocation schedule $(\bar{u}^*_{\mathrm{f}} = 1, \bar{u}^*_{\mathrm{q}} = 0)$) is the penultimate phase.

In order to determine if there are additional switches during time $0 < t < t^*_{c,1}$, one can further look for roots of the switching function $\sigma_1^c(t) = 0$ that satisfy $0 < t < t^*_{c,1}$. It follows from $\sigma_1^c(t) = 0$ and eq. (S78) for queen control and eq. (S80) for worker control that

$$\begin{aligned}
\sigma_1^{\mathrm{q}}(t) &= \frac{1}{2}\left(\lambda_{\mathrm{q}}^{\mathrm{q}}(t) - \lambda_{\mathrm{w}}^{\mathrm{q}}(t)\right) = 0 \qquad &(\mathbf{u}_{\mathrm{q}}, \mathbf{v}_{\mathrm{q}} \text{ under queen control}),\\
\sigma_1^{\mathrm{w}}(t) &= \frac{2+M}{4M}\left(\lambda_{\mathrm{q}}^{\mathrm{q}}(t) - \lambda_{\mathrm{w}}^{\mathrm{q}}(t)\right) = 0 \qquad &(\mathbf{u}_{\mathrm{q}}, \mathbf{v}_{\mathrm{q}} \text{ under worker control}),
\end{aligned} \tag{S108}$$

where costate variables at time $t$ are evaluated by using eq. (S82) $[t^*_{c,1}, T]$, assuming that $\mathbf{u}^*(t) = (\bar{u}^*_{\mathrm{f}} = \hat{u}^*_{\mathrm{f}}, \bar{u}^*_{\mathrm{q}} = 1)$ during $[t^*_{c,1}, T]$ and $\mathbf{u}^*(t) = (\bar{u}^*_{\mathrm{f}} = 1\bar{u}^*_{\mathrm{q}} = 0)$ during $[t, T]$. Our numerical solutions indicate that there are no additional growth regimes (see Figs. 1–2). This can be also shown analytically, by showing that there are no additional switches in the switching functions for $t < t^*_{c,1}$. However, for conciseness, we do not provide the proofs here.

In conclusion, we have determined that the optimal allocation schedule consists of two growth regimes: starting with the ergonomic regime W followed by reproductive regime FM, where the switch from regime W to regime FM happens at time $t^*_{c,1}$. We also verified that our analytical results are in line with our numerical solutions (see Fig. 1, where $\mu_{\mathrm{q}} = \mu_{\mathrm{m}} = \mu_{\mathrm{r}}$).

##### 5.1.3 Switching time for single-party control

We have determined the candidate optimal controls for the ergonomic $t \in [0, t^*_{c,1}]$ and the reproductive $t \in [t^*_{c,1}, T]$ phase (eq. S92 and S104), and we are now going to determine the switching time $t^*_{c,1}$ for single-party control (i.e. $c \in \{\mathrm{q}, \mathrm{w}\}$) that marks the time when the growth schedule switches from one regime to another.

We showed in the previous section that the control variable $u^*_{\mathrm{q}}(t)$ switches its value from 1 to 0, when $\sigma_1^c(t) = 0$. Hence, solving the equation $\sigma_1^c(t) = 0$ for $t$ gives the switching time $t^*_{c,1}$. For queen control of $\mathbf{u}_{\mathrm{q}}$ and $\mathbf{v}_{\mathrm{q}}$, eq. (S78) and $\sigma_1^{\mathrm{q}}(t^*_{\mathrm{q},1}) = 0$ this yields

$$\sigma_1^{\mathrm{q}}(t^*_{\mathrm{q},1}) = \frac{1}{2}\left(\lambda_{\mathrm{q}}^{\mathrm{q}}(t^*_{\mathrm{q},1}) - \lambda_{\mathrm{w}}^{\mathrm{q}}(t^*_{\mathrm{q},1})\right) = 0, \tag{S109}$$

and using the solutions for costate equations (S90), transversality conditions (S83) and (S86), and assuming

$\mu_{\rm q} = \mu_{\rm m} = \mu_{\rm r}$ it leads to the following transcendental equation for finding $t^*_{\rm q,1}$

$$\lambda^{\rm q}_{\rm q}(T)e^{-\mu(T-t^*_{\rm q,1})} - \frac{b\left(a^*_{\rm q}(t)\lambda^{\rm q}_{\rm q}(T) + a^*_{\rm m}(t)\lambda^{\rm q}_{\rm m}(T)\right)\left(e^{-\mu_{\rm w}\left(T-t^*_{\rm q,1}\right)} - e^{-\mu_{\rm r}\left(T-t^*_{\rm q,1}\right)}\right)}{\mu_{\rm r} - \mu_{\rm w}} = 0. \quad \text{(S110)}$$

Similarly, for worker control of $\mathbf{u}_{\rm q}$ and $\mathbf{v}_{\rm q}$, eq. (S80) and $\sigma^{\rm w}_1(t^*_{\rm w,1}) = 0$ yields

$$\sigma^{\rm w}_1(t^*_{\rm w,1}) = \frac{2+M}{4M}\left(\lambda^{\rm q}_{\rm q}(t^*_{\rm w,1}) - \lambda^{\rm q}_{\rm w}(t^*_{\rm w,1})\right) = 0 \quad \text{(S111)}$$

and using the solutions for costate equations (S90), transversality conditions (S83) and (S86), and assuming $\mu_{\rm q} = \mu_{\rm m} = \mu_{\rm r}$, it leads to the following transcendental equation for finding $t^*_{\rm w,1}$

$$\lambda^{\rm q}_{\rm q}(T)e^{-\mu_{\rm r}(T-t^*_{\rm w,1})} + \frac{2\lambda^{\rm m}_{\rm q}(T)}{M}\left(e^{-\mu_{\rm r}(T-t^*_{\rm w,1})} + e^{-\mu_{\rm w}\left(T-t^*_{\rm w,1}\right)} - e^{-\mu_{\rm r}\left(T-t^*_{\rm w,1}\right)}\right)$$
$$- \frac{a^*_{\rm q}(t)\lambda^{\rm q}_{\rm q}(T) + a^*_{\rm m}(t)\lambda^{\rm q}_{\rm m}(T)}{\mu_{\rm r} - \mu_{\rm w}}\left(e^{-\mu_{\rm w}\left(T-t^*_{\rm w,1}\right)} - e^{-\mu_{\rm r}\left(T-t^*_{\rm w,1}\right)}\right) = 0. \quad \text{(S112)}$$

By solving eq. (S110) for $t^*_{\rm q,1}$ and (S112) for $t^*_{\rm w,1}$ and taking $u^*_{\rm q}(t) = 1$, we obtain

$$t^*_{c,1} = T - \frac{\ln\left(1 + \theta_c\frac{\mu_{\rm r}-\mu_{\rm w}}{r}\right)}{\mu_{\rm r} - \mu_{\rm w}}, \quad \text{(S113)}$$

where

$$\frac{1}{\theta_c} = \begin{cases} u^*_{\rm f}(t) + (1-u^*_{\rm f}(t))\frac{\lambda^{\rm q}_{\rm m}(T)}{\lambda^{\rm q}_{\rm q}(T)} & \text{for } c = \text{q}, \\ u^*_{\rm f}(t) + (1-u^*_{\rm f}(t))\frac{\lambda^{\rm q}_{\rm m}(T)}{\lambda^{\rm q}_{\rm q}(T)+(2/M)\lambda^{\rm m}_{\rm q}(T)} & \text{for } c = \text{w}. \end{cases} \quad \text{(S114)}$$

Using eq. (S86) we obtain

$$\frac{1}{\theta_c} = u^*_{\rm f}(t) + (1-u^*_{\rm f}(t))\frac{1}{R_c}\frac{x^*_{\rm q}(\mathbf{u}^*)}{x^*_{\rm m}(\mathbf{u}^*)}. \quad \text{(S115)}$$

After substituting the control variable $u^*_{\rm f}(t) = \hat{\hat{u}}^*_{\rm f}$ from equation (S104) for the respective case of control and using eq. (S102), we finally have

$$\theta_c = \frac{1+R_c}{1+R_c} = 1. \quad \text{(S116)}$$

We have obtained that the switching time for single-party control is

$$t^*_{c,1} = T - \frac{\ln\left(1 + \frac{\mu_{\rm r}-\mu_{\rm w}}{b}\right)}{\mu_{\rm r} - \mu_{\rm w}} \text{ for } c \in \{\text{q}, \text{w}\}. \quad \text{(S117)}$$

In the limit where the mortality of sexuals becomes equal to the mortality of workers ($\mu_{\rm r} \to \mu_{\rm w}$) the switching time simplifies to

$$t^*_{c,1} = T - \frac{1}{b} \text{ for } c \in \{\text{q}, \text{w}\}. \quad \text{(S118)}$$

Note that in our model $(1/b)$ can be loosely interpreted as the time it takes for one worker to help produce one offspring, i.e. a generation time. Hence, when the mortality rate of sexuals is roughly equal to the mortality rate of workers, then the switching time from the ergonomic to the reproductive phase $t^*_{c,1}$ under single-party control ($c = \{\mathrm{q}, \mathrm{w}\}$) approaches to one generation time $(1/b)$ before the end of the season, i.e. only the last generation of brood is reproductive.

##### 5.1.4 Switching time for mixed control

It follows from eqs. (S65) and (S76) that under mixed control the trait $u^*_{\mathrm{q}}(t)$ is determined from the sign of $u^*_{\mathrm{f}}(t)\sigma^{\mathrm{w}}_1(t)$ and the trait $u^*_{\mathrm{f}}(t)$ is determined from from the sign of $(u^*_{\mathrm{q}}(t)\sigma^{\mathrm{q}}_1(t) - \sigma^{\mathrm{q}}_2(t))$. Hence, the switching time $t^*_{\mathrm{mx},1}$ (the switch in the trait $u^*_{\mathrm{q}}(t)$) from ergonomic phase to reproductive phase under mixed control can be found by solving $\sigma^{\mathrm{w}}_1(t) = 0$ for $t = t^*_{\mathrm{mx},1}$, which by way of eq. (S80) yields

$$\sigma^{\mathrm{w}}_1(t^*_{\mathrm{mx},1}) = \frac{2+M}{4M}\left(\lambda^{\mathrm{q}}_{\mathrm{q}}(t^*_{\mathrm{mx},1}) - \lambda^{\mathrm{q}}_{\mathrm{w}}(t^*_{\mathrm{mx},1})\right) = 0 \tag{S119}$$

and using the solutions for costate equations (S90), transversality conditions (S83) and (S86), and assuming $\mu_{\mathrm{q}} = \mu_{\mathrm{m}} = \mu_{\mathrm{r}}$, it leads to the following transcendental equation for finding $t^*_{\mathrm{mx},1}$

$$\begin{aligned}\lambda^{\mathrm{q}}_{\mathrm{q}}(T)e^{-\mu_{\mathrm{r}}(T-t^*_{\mathrm{mx},1})} + \frac{2\lambda^{\mathrm{m}}_{\mathrm{q}}(T)}{M}\left(e^{-\mu_{\mathrm{r}}(T-t^*_{\mathrm{mx},1})} + e^{-\mu_{\mathrm{w}}\left(T-t^*_{\mathrm{mx},1}\right)} - e^{-\mu_{\mathrm{r}}\left(T-t^*_{\mathrm{mx},1}\right)}\right) \\ - \frac{a^*_{\mathrm{q}}\lambda^{\mathrm{q}}_{\mathrm{q}}(T) + a^*_{\mathrm{m}}\lambda^{\mathrm{q}}_{\mathrm{m}}(T)}{\mu_{\mathrm{r}} - \mu_{\mathrm{w}}}\left(e^{-\mu_{\mathrm{w}}\left(T-t^*_{\mathrm{mx},1}\right)} - e^{-\mu_{\mathrm{r}}\left(T-t^*_{\mathrm{mx},1}\right)}\right) = 0.\end{aligned} \tag{S120}$$

Solving eq. (S120) for $t^*_{\mathrm{mx},1}$ yields

$$t^*_{\mathrm{mx},1} = T - \frac{\ln\left(1 + \theta_{\mathrm{mx}}\frac{\mu_{\mathrm{r}}-\mu_{\mathrm{w}}}{b}\right)}{\mu_{\mathrm{r}} - \mu_{\mathrm{w}}}, \tag{S121}$$

where

$$\frac{1}{\theta_{\mathrm{mx}}} = u^*_{\mathrm{f}}(t) + (1 - u^*_{\mathrm{f}}(t))\frac{\lambda^{\mathrm{q}}_{\mathrm{m}}(T)}{\lambda^{\mathrm{q}}_{\mathrm{q}}(T) + (2/M)\lambda^{\mathrm{m}}_{\mathrm{q}}(T)}.$$

Since according to eq. (S76) for mixed control, the trait $u^*_{\mathrm{f}}(t)$ is determined from equation $\sigma^{\mathrm{q}}_1(t) - \sigma^{\mathrm{q}}_2(t) = 0$. We have shown earlier (see eqs. S98–S104 for queen control) that this leads to an equation for $u^*_{\mathrm{f}}(t)$ given by eq. (S104) (for queen control). Substituting $u^*_{\mathrm{f}}(t) = \hat{u}^*_{\mathrm{f}}$ from eq. (S104) (for queen control) and with transversality conditions (S86) and eq. (S102) (for queen control) and using eq. (8) of the main text, we obtain

$$\theta_{\mathrm{mx}} = \frac{2+M}{1+M}. \tag{S122}$$

It follows from eq. (S121) that $t^*_{\mathrm{mx},1} < t^*_{\mathrm{q},1} = t^*_{\mathrm{w},1}$ and as $M \to \infty$, $t^*_{\mathrm{mx},1} \to t^*_{\mathrm{q},1} = t^*_{\mathrm{w},1}$ (see eq. S117). We should also mention that equations (S117) and (S121) hold if $\mu_{\mathrm{w}} < b/\theta_c + \mu_{\mathrm{r}}$. This not biologically restrictive, since the reproduction rate has to be significantly higher than worker mortality otherwise the population will go extinct. Finally, in the limit where the mortality of sexuals approaches the mortality of workers ($\mu_{\mathrm{r}} \to \mu_{\mathrm{w}}$) the switching time simplifies to

$$t^*_{\mathrm{mx},1} = T - \frac{\theta_{\mathrm{mx}}}{b} \text{ for } c \in \{\mathrm{q}, \mathrm{w}\}\,. \tag{S123}$$

Hence, when the mortality rate of sexuals is roughly equal to the mortality rate of workers, then under mixed control the switch happens $\theta_{\mathrm{mx}}$ generations earlier. For example, when females mate only once ($M = 1$) the switch to reproductive phase happens one and a half generations before the end of the season.

#### 5.2 Unequal male and female mortality ($\mu_{\mathrm{q}} \neq \mu_{\mathrm{m}}$)

The above analytical results hold for $\mu_{\mathrm{q}} = \mu_{\mathrm{m}}$. Our numerical solutions indicate that if the mortality rates of queens and males are not equal ($\mu_{\mathrm{q}} \neq \mu_{\mathrm{m}}$), then the sex that has the lower mortality rate is produced first. Furthermore, the numerical solutions confirm that males and queens are produced such that by the end of the season the ratio of queens to males is given by the relatedness asymmetry, i.e. eq. (S50) holds, regardless of the mortality rates of males and queens. In Figs. S1–S2, we have depicted our numerical results for the uninvadable proportional allocation $a^*_k(t)$ to individuals of different types and the corresponding number of individuals $x^*_k$, respectively, assuming that queen mortality is lower than male mortality.

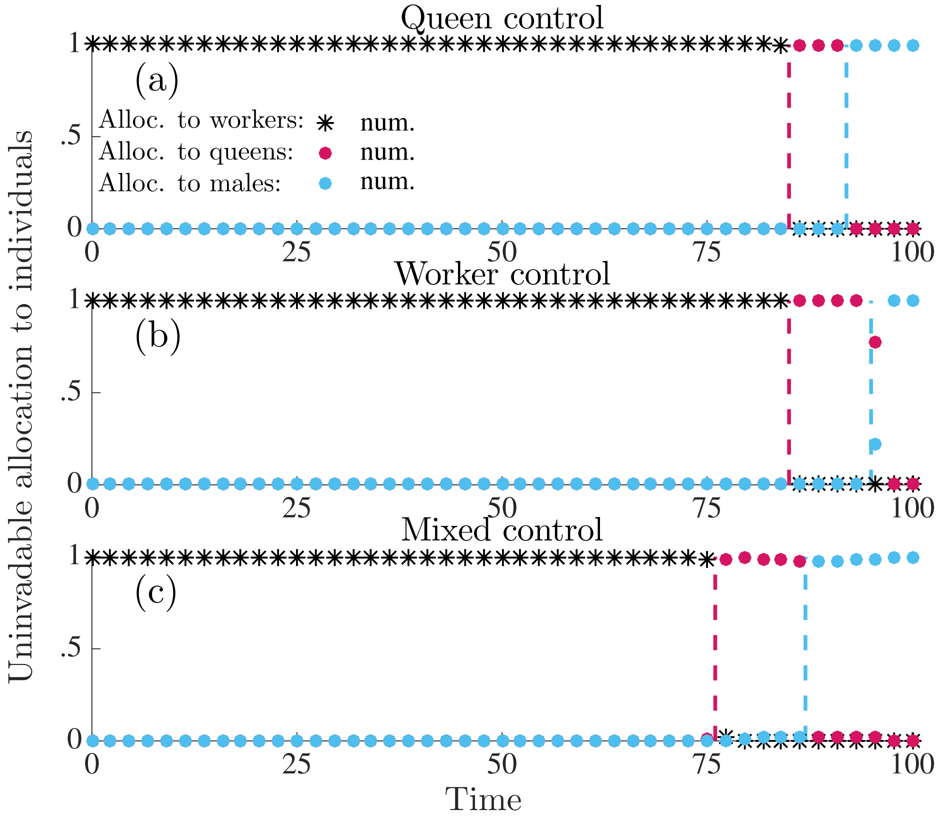

Figure S1: Uninvadable proportional allocation (under delayed dispersal) to workers $a^*_{\mathrm{w}}(t) = u^*_{\mathrm{f}}(t)(1 - u^*_{\mathrm{q}}(t))$ (black asterisks), queens $a^*_{\mathrm{q}}(t) = u^*_{\mathrm{f}}(t)u^*_{\mathrm{q}}(t)$ (red circles), and males $a^*_{\mathrm{m}}(t) = (1 - u^*_{\mathrm{f}}(t))$ (blue circles). Results here are only numerically derived. Panel (a): queen control. Panel (b): worker control. Panel (c): mixed control. Parameter values: $M = 1$ (queen monandry), $b = 0.07$, $\mu_{\mathrm{w}} = 0.015$, $\mu_{\mathrm{q}} = 0.001$, $\mu_{\mathrm{m}} = 0.02$, $T = 100$.

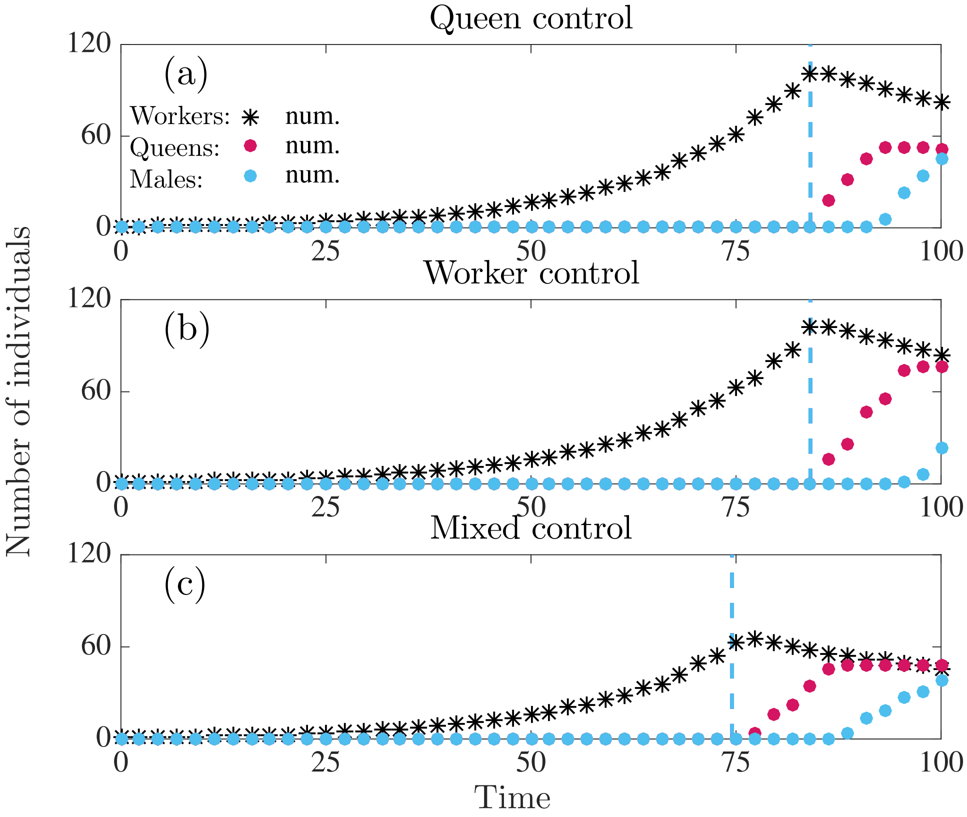

Figure S2: Number of individuals produced in a colony following the uninvadable resource allocation schedule $\mathbf{u}^*$ under delayed dispersal. Number of workers (black asterisks), number of juvenile queens (red circles), number of males (blue circles). Results here are only numerically derived. Panel (a): full queen control. Panel (b): full worker control. Panel (c): mixed control. Parameter values: $M = 1$ (queen monandry), $b = 0.07$, $\mu_{\mathrm{w}} = 0.015$, $\mu_{\mathrm{q}} = 0.001$, $\mu_{\mathrm{m}} = 0.02$, $T = 100$.

### 6 The candidate uninvadable allocation schedule under direct dispersal

#### 6.1 The cases $R_{\mathrm{c}}\mu_{\mathrm{q}} \geq \mu_{\mathrm{m}}$ (single-party control) and $R_{\mathrm{q}}\mu_{\mathrm{q}} \geq \mu_{\mathrm{m}}$ (mixed control)

Here, we determine analytically the candidate uninvadable allocation schedule under direct dispersal assuming $R_{\mathrm{c}}\mu_{\mathrm{q}} \geq \mu_{\mathrm{m}}$ (single-party control) and $R_{\mathrm{q}}\mu_{\mathrm{q}} \geq \mu_{\mathrm{m}}$ (mixed control). We find that the optimal allocation schedule consists of the following three growth regimes: (i) ergonomic phase (production of workers), (ii) reproductive phase where only males are produced, (iii) reproductive phase where only new queens are produced. Under these conditions, the uninvadable allocation schedule has the following properties

$$\begin{aligned}
&\textbf{Regime W:} \quad (u_{\mathrm{f}}^*(t), u_{\mathrm{q}}^*(t)) = (\bar{u}_{\mathrm{f}}^* = 1, \bar{u}_{\mathrm{q}}^* = 0) \quad \text{for } t \in \left[0, t_{c,1}^*\right], \\
&\textbf{Regime M:} \quad (u_{\mathrm{f}}^*(t), u_{\mathrm{q}}^*(t)) = (\bar{u}_{\mathrm{f}}^* = 0, \tilde{u}_{\mathrm{q}}^*(t)) \quad \text{for } t \in \left[t_{c,1}^*, t_{c,2}^*\right], \\
&\textbf{Regime F:} \quad (u_{\mathrm{f}}^*(t), u_{\mathrm{q}}^*(t)) = (\bar{u}_{\mathrm{f}}^* = 1, \bar{u}_{\mathrm{q}}^* = 1) \quad \text{for } t \in \left[t_{c,2}^*, T\right],
\end{aligned} \tag{S124}$$

where $t_{c,1}^*$ and $t_{c,2}^*$ in denote the switching times from ergonomic to reproductive phase and from male production to queen production, respectively, and they depend on the mode of control $c \in \{\mathrm{q}, \mathrm{w}, \mathrm{mx}\}$. We now derive this schedule by working backwards in time.

##### 6.1.1 Regime F: $\sigma_1^c(t) > 0$ and $\sigma_1^c(t) - \sigma_2^c(t) > 0$

The transversality conditions (S83) and (S87) yield that $\lambda_{\mathrm{w}}^{\mathrm{q}}(T) = \lambda_{\mathrm{w}}^{\mathrm{m}}(T) = 0$, $\lambda_{\mathrm{q}}^{\mathrm{q}}(T) = \lambda_{\mathrm{q}}^{\mathrm{m}}(T) > 0$, and $\lambda_{\mathrm{m}}^{\mathrm{q}}(T) = \lambda_{\mathrm{m}}^{\mathrm{m}}(T) = 0$. Hence, it follows from eqs. (S78) and (S79) that $\sigma_1^c(T) > 0$, $\sigma_2^c(T) = 0$, and $(\sigma_1^c(T) - \sigma_2^c(T)) > 0$. Therefore, from table S1 it follows that $(\bar{u}_{\mathrm{f}}^* = 1, \bar{u}_{\mathrm{q}}^* = 1)$ during $t \in [t_{c,2}^*, T]$, where $t_{c,2}^*$ marks the beginning of the last growth regime.

##### 6.1.2 Regime M: $\sigma_1^c(t) > 0$ and $\sigma_1^c(t) - \sigma_2^c(t) > 0$

If at least one of the switching functions (S77) changes its sign at time $t_{c,2}^*$, then one of the three alternative conditions must hold: (i) $\sigma_1^c(t_{c,2}^*) = 0$, which implies a change in the control variable $u_{\mathrm{q}}^*(t)$, (ii) $(\sigma_1^c(t_{c,2}^*) - \sigma_2^c(t_{c,2}^*)) = 0$, which means that the control variable $u_{\mathrm{f}}^*(t)$ changes, or (iii) $(\sigma_1^c(t_{c,2}^*) - \sigma_2^c(t_{c,2}^*)) = \sigma_1^c(t_{c,2}^*) = 0$, which means that both control variables change. The switching time $t_{c,2}^*$ of an uninvadable allocation schedule (S124) is given by the largest root $t = t_{c,2}^*$ that satisfies one of the conditions given by these scenarios.

Next, we will solve eqs. $\sigma_1^c(t_{c,2}^*) = 0$ and $(\sigma_1^c(t_{c,2}^*) - \sigma_2^c(t_{c,2}^*)) = 0$ for $t_{c,2}^*$, taking into account that that $(\bar{u}_{\mathrm{f}}^* = 1, \bar{u}_{\mathrm{q}}^* = 1)$ during $t \in [t_{c,2}^*, T]$ and eq. (S91) that implies that $\lambda_{\mathrm{w}}^{\mathrm{q}}(t) = \lambda_{\mathrm{w}}^{\mathrm{m}}(t)$. After which we will compare the the roots $t_{c,2}^*$ for these two equations in order to determine which of the three alternative above-mentioned scenarios holds.

Firstly, we determine the root $t_{c,2}^*$ of eq. $\sigma_1^c(t_{c,2}^*) = 0$. Substituting the transversality conditions (S83) and (S87) into eq. (S90), where we take $t_1 = T$ and assume that $(\bar{u}_{\mathrm{f}}^* = 1, \bar{u}_{\mathrm{q}}^* = 1)$ we obtain for $t \in [t_{c,2}^*, T]$

$$\lambda_{\mathrm{w}}^s(t) = \lambda_{\mathrm{q}}^s(t) \frac{b}{\mu_{\mathrm{q}} - \mu_{\mathrm{w}}} \left(e^{(-\mu_{\mathrm{w}})(T-t)} - e^{-\mu_{\mathrm{q}}(T-t)}\right). \tag{S125}$$

Substituting eq. (S91) into eq. (S125) yields that $\lambda_{\mathrm{w}}^{\mathrm{q}}(t) = \lambda_{\mathrm{w}}^{\mathrm{m}}(t)$. Taking this into account and substituting eqs. (S78) and (S79) into $\sigma_1^c(t_{c,2}^*) = 0$ yields

$$\begin{aligned}
\sigma_1^{\mathrm{q}}(t_{c,2}^*) &= \frac{1}{2}\left(\lambda_{\mathrm{q}}^{\mathrm{q}}(t_{c,2}^*) - \lambda_{\mathrm{w}}^{\mathrm{q}}(t_{c,2}^*)\right) = 0 \qquad (\mathbf{u}_{\mathrm{q}}, \mathbf{v}_{\mathrm{q}} \text{ under queen control}), \\
\sigma_1^{\mathrm{w}}(t_{c,2}^*) &= \frac{2+M}{4M}\left(\lambda_{\mathrm{q}}^{\mathrm{q}}(t_{c,2}^*) - \lambda_{\mathrm{w}}^{\mathrm{q}}(t_{c,2}^*)\right) = 0 \qquad (\mathbf{u}_{\mathrm{q}}, \mathbf{v}_{\mathrm{q}} \text{ under worker control}).
\end{aligned} \tag{S126}$$

Substituting the solutions to the costate equations (S90) (assuming that $t_1 = T$ and $t = t^*_{c,2}$), the transversality conditions (S83), (S87) into eq. (S126) assuming that $(\bar{u}^*_\mathrm{f} = 1, \bar{u}^*_\mathrm{q} = 1)$ during $t \in [t^*_{c,2}, T]$ and solving for $t^*_{c,2}$ yields

$$t^*_{c,2} = T - \frac{\ln\left(1 + \frac{\mu_\mathrm{r} - \mu_\mathrm{w}}{b}\right)}{\mu_\mathrm{r} - \mu_\mathrm{w}}. \tag{S127}$$

Note that the derivation of eq. (S127) from eq. (S126) is not shown here, since it is very similar to derivation of eq. (S117) from eqs. (S109) and (S111).

Secondly, we need to determine the root $t^*_{c,2}$ of eq. $(\sigma^c_1(t^*_{c,2}) - \sigma^c_2(t^*_{c,2})) = 0$. Taking into account that $\lambda^\mathrm{q}_\mathrm{w}(t) = \lambda^\mathrm{m}_\mathrm{w}(t)$ for $t \in [t^*_{c,2}, T]$ (recall the implications of eqs. S125 and S91) and substituting eqs. (S78) and (S79) (assuming that $t = t^*_{c,2}$) into $(\sigma^c_1(t^*_{c,2}) - \sigma^c_2(t^*_{c,2})) = 0$ yields for $t = t^*_{c,2}$ that

$$\begin{aligned} &\lambda^\mathrm{q}_\mathrm{q}(t) = \lambda^\mathrm{q}_\mathrm{m}(t) && (\mathbf{u}_\mathrm{f}, \mathbf{v}_\mathrm{f} \text{ under queen control}), \\ &\frac{1}{4}\lambda^\mathrm{q}_\mathrm{q}(t) + \frac{1}{2M}\lambda^\mathrm{m}_\mathrm{q}(t) = \frac{1}{4}\lambda^\mathrm{q}_\mathrm{m}(t) + \frac{1}{2M}\lambda^\mathrm{m}_\mathrm{m}(t) && (\mathbf{u}_\mathrm{f}, \mathbf{v}_\mathrm{f} \text{ under worker control}). \end{aligned} \tag{S128}$$

We will show in sections 6.1.4 and 6.1.5 that substituting the solutions to the costate equations (S90) (assuming that $t_1 = T$ and $t = t^*_{c,2}$) and the transversality conditions (S83), (S87) into eq. (S128) yields a switching time $t^*_{c,2}$ given by eq. (S139) for single-party control (assuming $R_c\mu_\mathrm{q} \geq \mu_\mathrm{m}$) and eqs. (S147) and (S146) for mixed control (assuming $\mu_\mathrm{q} = \mu_\mathrm{m}$).

Finally, comparing the root $t^*_{c,2}$ of eq. $\sigma^c_1(t^*_{c,2}) = 0$ given by eq. (S127) with roots of eq. $(\sigma^c_1(t^*_{c,2}) - \sigma^c_2(t^*_{c,2})) = 0$, given by eq. (S139) (for single-party control) and eqs. (S147) and (S146) (for mixed control), yields that for biologically realistic parameter values, the root $t^*_{c,2}$ of eq. $\sigma^c_1(t^*_{c,2}) = 0$ is smaller than the roots of eq. $(\sigma^c_1(t^*_{c,2}) - \sigma^c_2(t^*_{c,2})) = 0$. Hence, we have verified that the switch $t^*_{c,2}$ is given by the root of $(\sigma^c_1(t^*_{c,2}) - \sigma^c_2(t^*_{c,2})) = 0$ and it follows that the control variable $u^*_\mathrm{f}(t)$ changes its sign at this time.

We have established that $(\sigma^c_1(t) - \sigma^c_2(t)) = 0$ at time $t = t^*_{c,2}$. Next we have to determine if $(\sigma^c_1(t) - \sigma^c_2(t)) = 0$ only at time $t = t^*_{c,2}$ or if $(\sigma^c_1(t) - \sigma^c_2(t)) = 0$ during a finite period of time that ends at time $t = t^*_{c,2}$. It follows from the definition of the singular arc that if $(\sigma^c_1(t) - \sigma^c_2(t)) = 0$ holds for a finite interval of time then $u_\mathrm{f}(t) = \hat{u}^*_\mathrm{f}(t)$ is a singular arc during that time (e.g., Bryson and Ho, 1975, p. 246–249). Furthermore, if $(\sigma^c_1(t) - \sigma^c_2(t)) = 0$ during a finite period of time then it also follows that $(\dot{\sigma}_1(t) - \dot{\sigma}_2(t)) = 0$ holds during that time. Furthermore, it follows from the time derivative of eq. (S128) that a condition for a singular arc to exist is given by

$$\begin{aligned} &\dot{\lambda}^\mathrm{q}_\mathrm{q}(t) = \dot{\lambda}^\mathrm{q}_\mathrm{m}(t) && (\mathbf{u}_\mathrm{f}, \mathbf{v}_\mathrm{f} \text{ under queen control}), \\ &\frac{1}{4}\dot{\lambda}^\mathrm{q}_\mathrm{q}(t) + \frac{1}{2M}\dot{\lambda}^\mathrm{m}_\mathrm{q}(t) = \frac{1}{4}\dot{\lambda}^\mathrm{q}_\mathrm{m}(t) + \frac{1}{2M}\dot{\lambda}^\mathrm{m}_\mathrm{m}(t) && (\mathbf{u}_\mathrm{f}, \mathbf{v}_\mathrm{f} \text{ under worker control}). \end{aligned} \tag{S129}$$

Simplifying eq. (S129) by considering that $\dot{\lambda}^\mathrm{q}_\mathrm{q}(t) = \dot{\lambda}^\mathrm{m}_\mathrm{q}(t)$ and $\dot{\lambda}^\mathrm{m}_\mathrm{m}(t) = 0$ (by way of eq. S91) and using eq. (S35) we obtain

$$\frac{\dot{\lambda}^\mathrm{q}_\mathrm{m}(t)}{\dot{\lambda}^\mathrm{q}_\mathrm{q}(t)} = R_c, \tag{S130}$$

where $c$ denotes the party in control of $\mathbf{u}_\mathrm{f}$ and $\mathbf{v}_\mathrm{f}$. Substituting the costate equations (S82) into eq. (S130) yields

$$\frac{\mu_\mathrm{m}\lambda_\mathrm{m}^\mathrm{q}(t) - \frac{ba_\mathrm{q}^*(t)x_\mathrm{w}^*(t)\lambda_\mathrm{iq}^\mathrm{q}(t)}{x_\mathrm{m}^*(t)}}{\mu_\mathrm{q}\lambda_\mathrm{q}^\mathrm{q}(t)} = R_c. \tag{S131}$$

Taking this account together with eq. (S91) and $a_\mathrm{q}^*(t) = u_\mathrm{f}^*(t)u_\mathrm{q}^*(t)$ implies that

$$\frac{ba_\mathrm{q}^*(t)x_\mathrm{w}^*(t)}{x_\mathrm{m}^*(t)} = \frac{bu_\mathrm{f}^*(t)u_\mathrm{q}^*(t)x_\mathrm{w}^*(t)}{x_\mathrm{m}^*(t)} = \mu_\mathrm{m} - R_c\mu_\mathrm{q}. \tag{S132}$$

Given that $u_\mathrm{q}^* = \bar{u}_\mathrm{q}^* = 1$ for $t < t_{c,2}^*$ (since $\sigma_1^c(t) > 0$ for $t < t_{c,2}^*$) then eq. (S131) implies that $u_\mathrm{f}^*(t) = \hat{u}_\mathrm{f}^*(t)$ can only be positive if $(\mu_\mathrm{m} - R_c\mu_\mathrm{q}) > 0$. Note that here $c$ denotes the party in control of $\mathbf{u}_\mathrm{f}$ and $\mathbf{v}_\mathrm{f}$. Hence, $u_\mathrm{f}^*$ can not be a singular arc before $t_{c,2}^*$ if juvenile male mortality is lower or equal than that of $R_c$ times juvenile queen mortality (i.e. $\mu_\mathrm{m} \leq R_c\mu_\mathrm{q}$).

Hence we have determined that in the penultimate phase, which ends at time, $t_{c,2}^*$ that $\sigma_1^c(t) > 0$ and $\sigma_1^c(t) - \sigma_2^c(t) < 0$ if $R_c\mu_\mathrm{q} \geq \mu_\mathrm{m}$ (under single-party control) or $R_\mathrm{q}\mu_\mathrm{q} \geq \mu_\mathrm{m}$ (under mixed control). This means that if $R_c\mu_\mathrm{q} \geq \mu_\mathrm{m}$ (under single-party control) or $R_\mathrm{q}\mu_\mathrm{q} \geq \mu_\mathrm{m}$ (under mixed control) then regime M (exclusive production of juvenile males) precedes the final regime F, where control variables are given by $(\bar{u}_\mathrm{f}^* = 0, \bar{u}_\mathrm{q}^* = 1)$. Later in this section we will revisit the case when $R_c\mu_\mathrm{q} < \mu_\mathrm{m}$ (under single-party control) and $R_\mathrm{q}\mu_\mathrm{q} < \mu_\mathrm{m}$ (under mixed control).

##### 6.1.3 Regime W: $\sigma_1^c < 0$ and $\sigma_2^c < 0$

We use the intuition from our numerical solutions (see Fig. 2) that there exists only one additional switching time $t_{c,1}^*$, when $\sigma_1^c(t)$ and $\sigma_2^c(t)$ become negative, which represent the first growth regime of the uninvadable allocation schedule. Hence, regime W (worker production) is the first growth regime of the uninvadable allocation schedule, where $\mathbf{u}^*(t) = (\bar{u}_\mathrm{f}^* = 1, \bar{u}_\mathrm{q}^* = 0)$. We can determine the switching time $t_{c,1}^*$ from the condition $\sigma_1^c(t) = 0$, where the costate variables are obtained by integrating them over the last two growth regimes. It turns out that we can explicitly calculate the switching times $t_{c,1}^*$ and $t_{c,2}^*$ only if $R_c\mu_\mathrm{q} \geq \mu_\mathrm{m}$ under single-party control and only if $\mu_\mathrm{q} = \mu_\mathrm{q} = \mu_\mathrm{r}$ under mixed control.

##### 6.1.4 Switching times under single-party control $R_c\mu_\mathrm{q} \geq \mu_\mathrm{m}$

If $\mu_\mathrm{m} \leq R_c\mu_\mathrm{q}$ then the condition for the switching time $t_{c,2}^*$ which marks the transition from production of exclusively males to the production of exclusively sexual females is given by (resulting from $\sigma_1^c(t_{c,2}^*) - \sigma_2^c(t_{c,2}^*) = 0$)

$$\begin{aligned}
&\lambda_\mathrm{q}^\mathrm{q}(t_{c,2}^*) = \lambda_\mathrm{m}^\mathrm{q}(t_{c,2}^*) && (\mathbf{u}_\mathrm{f}, \mathbf{v}_\mathrm{f} \text{ under queen control}),\\
&\frac{1}{4}\lambda_\mathrm{q}^\mathrm{q}(t_{c,2}^*) + \frac{1}{2M}\lambda_\mathrm{q}^\mathrm{m}(t_{c,2}^*) = \frac{1}{4}\lambda_\mathrm{m}^\mathrm{q}(t_{c,2}^*) + \frac{1}{2M}\lambda_\mathrm{m}^\mathrm{m}(t_{c,2}^*) && (\mathbf{u}_\mathrm{f}, \mathbf{v}_\mathrm{f} \text{ under worker control}).
\end{aligned} \tag{S133}$$

Using eq. (S90) for phase $t \in \left[t^*_{c,2}, T\right]$, where $\bar{u}^*_{\mathrm{f}} = 1$ and $\bar{u}^*_{\mathrm{q}} = 1$ and simplifying, we get

$$\begin{aligned}
\lambda^s_{\mathrm{q}}(t^*_{c,2}) &= \lambda^s_{\mathrm{q}}(T) e^{-\mu_{\mathrm{q}}(T-t^*_{c,2})},\\
\lambda^{\mathrm{q}}_{\mathrm{m}}(t^*_{c,2}) &= \lambda^{\mathrm{q}}_{\mathrm{iq}}(T) \frac{e^{-\mu_{\mathrm{q}}(T-t^*_{c,2})}(\mu_{\mathrm{m}}-\mu_{\mathrm{w}})\left(e^{(\mu_{\mathrm{q}}-\mu_{\mathrm{w}})(T-t^*_{c,2})}-1\right)}{(\mu_{\mathrm{q}}-\mu_{\mathrm{w}})\left(1-e^{-(\mu_{\mathrm{m}}-\mu_{\mathrm{w}})(t^*_{c,2}-t^*_{c,1})}\right)},\\
\lambda^{\mathrm{m}}_{\mathrm{m}}(t^*_{c,2}) &= 0.
\end{aligned} \tag{S134}$$

Substituting of eq. (S134) into (S133) and using eq. (S87) implies

$$\begin{aligned}
&(\mu_{\mathrm{m}}-\mu_{\mathrm{w}})e^{(\mu_{\mathrm{q}}-\mu_{\mathrm{w}})(T-t^*_{c,2})} + (\mu_{\mathrm{q}}-\mu_{\mathrm{w}})e^{-(\mu_{\mathrm{m}}-\mu_{\mathrm{w}})(t^*_{c,2}-t^*_{c,1})} =\\
&(\mu_{\mathrm{q}}-\mu_{\mathrm{w}}) + (\mu_{\mathrm{m}}-\mu_{\mathrm{w}}) \qquad (\mathbf{u}_{\mathrm{f}}, \mathbf{v}_{\mathrm{f}} \text{ under queen control}),\\
&\frac{M}{2+M}(\mu_{\mathrm{m}}-\mu_{\mathrm{w}})e^{(\mu_{\mathrm{q}}-\mu_{\mathrm{w}})(T-t^*_{c,2})} + (\mu_{\mathrm{q}}-\mu_{\mathrm{w}})e^{-(\mu_{\mathrm{m}}-\mu_{\mathrm{w}})(t^*_{c,2}-t^*_{c,1})} =\\
&(\mu_{\mathrm{q}}-\mu_{\mathrm{w}}) + \frac{M}{2+M}(\mu_{\mathrm{m}}-\mu_{\mathrm{w}}) \qquad (\mathbf{u}_{\mathrm{f}}, \mathbf{v}_{\mathrm{f}} \text{ under worker control}).
\end{aligned} \tag{S135}$$

The switching time $t^*_{c,1}$ which marks the transition from production of exclusively workers to the production of exclusively males can be found by solving the eq. $\sigma^c_2(t^*_{c,1}) = 0$ for $t^*_{c,1}$ (see table S1)

$$\begin{aligned}
&\lambda^{\mathrm{q}}_{\mathrm{w}}(t^*_{c,1}) = \lambda^{\mathrm{q}}_{\mathrm{m}}(t^*_{c,1}) \qquad (\mathbf{u}_{\mathrm{f}}, \mathbf{v}_{\mathrm{f}} \text{ under queen control}),\\
&\frac{1}{4}\lambda^{\mathrm{q}}_{\mathrm{w}}(t^*_{c,1}) + \frac{1}{2M}\lambda^{\mathrm{m}}_{\mathrm{w}}(t^*_{c,1}) = \frac{1}{4}\lambda^{\mathrm{q}}_{\mathrm{m}}(t^*_{c,1}) + \frac{1}{2M}\lambda^{\mathrm{m}}_{\mathrm{m}}(t^*_{c,1}) \qquad (\mathbf{u}_{\mathrm{f}}, \mathbf{v}_{\mathrm{f}} \text{ under worker control}).
\end{aligned} \tag{S136}$$

Using eq. (S90) for phase $t \in \left[t^*_{c,1}, t^*_{c,2}\right]$, for which $\bar{u}^*_{\mathrm{f}} = 0$ and simplifying, we get

$$\begin{aligned}
\lambda^s_{\mathrm{w}}(t^*_{c,1}) &= \lambda^s_{\mathrm{w}}(t^*_{c,2})e^{-\mu_{\mathrm{w}}(t^*_{c,2}-t^*_{c,1})} + \frac{b\lambda^s_{\mathrm{m}}(t^*_{c,2})}{(\mu_{\mathrm{m}}-\mu_{\mathrm{w}})}(e^{-\mu_{\mathrm{w}}(t^*_{c,2}-t^*_{c,1})} - e^{-\mu_{\mathrm{m}}(t^*_{c,2}-t^*_{c,1})}),\\
\lambda^s_{\mathrm{m}}(t^*_{c,1}) &= \lambda^s_{\mathrm{m}}(t^*_{c,2})e^{-\mu_{\mathrm{m}}(t^*_{c,2}-t^*_{c,1})}.
\end{aligned} \tag{S137}$$

Substituting of eq. (S137) into (S136) and solving for $t^*_{c,1} = t^*_{\mathrm{q},1}$ (under queen control) and $t^*_{c,1} = t^*_{\mathrm{w},1}$ (under worker control) implies

$$\begin{aligned}
t^*_{\mathrm{q},1} &= t^*_{\mathrm{q},2} - \frac{1}{(\mu_{\mathrm{m}}-\mu_{\mathrm{w}})}\ln\left(\frac{2b+(\mu_{\mathrm{m}}-\mu_{\mathrm{w}})}{2b}\right) \qquad \text{(queen control)},\\
t^*_{\mathrm{w},1} &= t^*_{\mathrm{w},2} - \frac{1}{(\mu_{\mathrm{m}}-\mu_{\mathrm{w}})}\ln\left(\frac{2b+\frac{M}{1+M}(\mu_{\mathrm{m}}-\mu_{\mathrm{w}})}{2b}\right) \qquad \text{(worker control)}.
\end{aligned} \tag{S138}$$

Substituting eq. (S138) into eq. (S135) and solving for $t^*_{c,2}$ we obtain

$$\begin{aligned} t^*_{\mathrm{q},2} &= T - \frac{1}{(\mu_\mathrm{q} - \mu_\mathrm{w})} \ln\left(\frac{2b + \mu_\mathrm{m} + \mu_\mathrm{q} - 2\mu_\mathrm{w}}{2b + \mu_\mathrm{m} - \mu_\mathrm{w}}\right) && \text{(queen control)}, \\ t^*_{\mathrm{w},2} &= T - \frac{1}{(\mu_\mathrm{q} - \mu_\mathrm{w})} \ln\left(\frac{2b + \frac{1}{1+M}(M\mu_\mathrm{m} + (2+M)\mu_\mathrm{q} - 2(1+M)\mu_\mathrm{w})}{2b + \frac{M}{1+M}(\mu_\mathrm{m} - \mu_\mathrm{w})}\right) && \text{(worker control)}. \end{aligned} \tag{S139}$$

##### 6.1.5 Switching time under mixed control when $\mu_\mathrm{q} = \mu_\mathrm{m} = \mu_\mathrm{r}$

For mixed control, we will derive the switching times $t^*_{\mathrm{mx},1}$ and $t^*_{\mathrm{mx},2}$ assuming that juvenile queen and male mortality is equal, i.e. $\mu_\mathrm{m} = \mu_\mathrm{q} = \mu_\mathrm{r}$, since this represents the only case where we were able to derive analytical expressions. Under mixed control, the workers control the trait $\mathbf{u}_\mathrm{q}$ and the queen controls the trait $\mathbf{u}_\mathrm{f}$. Hence, it follows from eqs. (S76) and (S124) that the switching time $t^*_{\mathrm{mx},1}$ is determined from equation $\sigma^\mathrm{w}_1(t^*_{\mathrm{mx},1}) = 0$ and $t^*_{\mathrm{mx},2}$ is determined from equation $\sigma^\mathrm{q}_1(t^*_{\mathrm{mx},2}) - \sigma^\mathrm{q}_2(t^*_{\mathrm{mx},2}) = 0$.

It follows from eq. (S77) that condition $\sigma^\mathrm{w}_1(t^*_{\mathrm{mx},1}) = 0$ yields

$$\frac{1}{4}\lambda^\mathrm{q}_\mathrm{w}(t^*_{\mathrm{mx},1}) + \frac{1}{2M}\lambda^\mathrm{m}_\mathrm{w}(t^*_{\mathrm{mx},1}) = \frac{1}{4}\lambda^\mathrm{q}_\mathrm{q}(t^*_{\mathrm{mx},1}) + \frac{1}{2M}\lambda^\mathrm{m}_\mathrm{q}(t^*_{\mathrm{mx},1}). \tag{S140}$$

Using eq. (S90) for phase $t \in \left[t^*_{\mathrm{mx},2}, T\right]$, where $\bar{u}^*_\mathrm{f} = 1$ and $\bar{u}^*_\mathrm{q} = 1$ and simplifying, we get

$$\begin{aligned} \lambda^s_\mathrm{w}(t^*_{\mathrm{mx},2}) &= \frac{b\lambda^s_\mathrm{q}(T)}{(\mu_\mathrm{q} - \mu_\mathrm{w})}(e^{-\mu_\mathrm{w}(T - t^*_{\mathrm{mx},2})} - e^{-\mu_\mathrm{q}(T - t^*_{\mathrm{mx},2})}), \\ \lambda^s_\mathrm{q}(t^*_{\mathrm{mx},2}) &= \lambda^s_\mathrm{q}(T)e^{-\mu_\mathrm{q}(T - t^*_{\mathrm{mx},2})}. \end{aligned} \tag{S141}$$

Using eq. (S90) for phase $t \in \left[t^*_{\mathrm{mx},1}, t^*_{\mathrm{mx},2}\right]$, where $\bar{u}^*_\mathrm{f} = 0$ and simplifying, we get

$$\begin{aligned} \lambda^s_\mathrm{w}(t^*_{\mathrm{mx},1}) &= \lambda^s_\mathrm{w}(t^*_{\mathrm{mx},2})e^{-\mu_\mathrm{w}(t^*_{\mathrm{mx},2} - t^*_{\mathrm{mx},1})} + \frac{b\lambda^s_\mathrm{m}(t^*_{\mathrm{mx},2})}{(\mu_\mathrm{m} - \mu_\mathrm{w})}(e^{-\mu_\mathrm{w}(t^*_{\mathrm{mx},2} - t^*_{\mathrm{mx},1})} - e^{-\mu_\mathrm{m}(t^*_{\mathrm{mx},2} - t^*_{\mathrm{mx},1})}), \\ \lambda^s_\mathrm{q}(t^*_{\mathrm{mx},1}) &= \lambda^s_\mathrm{q}(t^*_{\mathrm{mx},2})e^{-\mu_\mathrm{q}(t^*_{\mathrm{mx},2} - t^*_{\mathrm{mx},1})}. \end{aligned} \tag{S142}$$

Substituting the the costate variables from eq. (S142) together with (S141) into eq. (S140) and solving for $t^*_{\mathrm{mx},1}$ yields

$$t^*_{\mathrm{mx},1} = \frac{\mu_\mathrm{w}(T + t^*_{\mathrm{mx},2}) - \ln\left(-\frac{2(M+1)b\left(e^{\mu_\mathrm{q}T + \mu_\mathrm{w}t^*_{\mathrm{mx},2}} - e^{\mu_\mathrm{w}T + t^*_{\mathrm{mx},2}}\right)}{(M+2)(\mu_\mathrm{w} - \mu_\mathrm{q})}\right)}{\mu_\mathrm{w} - \mu_\mathrm{q}}. \tag{S143}$$

It follows from eq. (S77) that condition $\sigma^\mathrm{q}_1(t^*_{\mathrm{mx},2}) - \sigma^\mathrm{q}_2(t^*_{\mathrm{mx},2}) = 0$ yields

$$\lambda^\mathrm{q}_\mathrm{q}(t^*_{\mathrm{mx},2}) = \lambda^\mathrm{q}_\mathrm{m}(t^*_{\mathrm{mx},2}). \tag{S144}$$

Substituting the costate variables from eq. (S134) and the switching time $t^*_{\mathrm{mx},1}$ given by eq. (S143) into eq. (S144) and solving for $t^*_{\mathrm{mx},2}$ (assuming that $\mu_{\mathrm{q}} = \mu_{\mathrm{q}} = \mu_{\mathrm{r}}$) yields

$$t^*_{\mathrm{mx},2} = T - \frac{\ln\left(2 - \frac{b}{b+\frac{1}{2}\theta_{\mathrm{mx}}(\mu_{\mathrm{r}}-\mu_{\mathrm{w}})}\right)}{\mu_{\mathrm{r}} - \mu_{\mathrm{w}}}, \tag{S145}$$

where

$$\theta_{\mathrm{mx}} = \frac{2+M}{1+M}. \tag{S146}$$

Substituting $t^*_{\mathrm{mx},2}$ given by eq. (S145) back into eq. (S143) and simplifying yields

$$t^*_{\mathrm{mx},1} = T - \frac{\ln\left(1 + \theta_{\mathrm{mx}}\frac{\mu_{\mathrm{r}}-\mu_{\mathrm{w}}}{b}\right)}{\mu_{\mathrm{r}} - \mu_{\mathrm{w}}}. \tag{S147}$$

Hence, we have retrieved the same switching time from the ergonomic to the reproductive phase for mixed control under direct dispersal and delayed dispersal (given by eq. S121).

#### 6.2 Equal mortality rates of males and queens ($\mu_{\mathrm{q}} = \mu_{\mathrm{m}}$)

In this section, we present the results for the candidate uninvadable allocation schedule under direct dispersal assuming that the mortality rates of queens and males are equal ($\mu_{\mathrm{q}} = \mu_{\mathrm{m}} = \mu_{\mathrm{r}}$). It turns out these results can be directly obtained section 6.1 by equating the mortality rates of queens and males are equal ($\mu_{\mathrm{q}} = \mu_{\mathrm{m}} = \mu_{\mathrm{r}}$). This is because the results in section 6.1 were derived assuming that $R_c\mu_{\mathrm{q}} \geq \mu_{\mathrm{m}}$ (under single-party control) and $R_{\mathrm{q}}\mu_{\mathrm{q}} \geq \mu_{\mathrm{m}}$ (under mixed control), where relatedness asymmetry $R_c \geq 1$ (recall eq. (S35)).

The optimal allocation schedule consists of the following three growth regimes: (i) ergonomic phase (production of workers), (ii) reproductive phase where only males are produced, (iii) reproductive phase where only new queens are produced. The uninvadable allocation schedule has the following properties

$$\begin{aligned}
&\textbf{Regime W:} \quad (u^*_{\mathrm{f}}(t), u^*_{\mathrm{q}}(t)) = (\bar{u}^*_{\mathrm{f}} = 1, \bar{u}^*_{\mathrm{q}} = 0) \quad \text{for } t \in \left[0, t^*_{c,1}\right], \\
&\textbf{Regime M:} \quad (u^*_{\mathrm{f}}(t), u^*_{\mathrm{q}}(t)) = (\bar{u}^*_{\mathrm{f}} = 0, \tilde{u}^*_{\mathrm{q}}(t)) \quad \text{for } t \in \left[t^*_{c,1}, t^*_{c,2}\right], \\
&\textbf{Regime F:} \quad (u^*_{\mathrm{f}}(t), u^*_{\mathrm{q}}(t)) = (\bar{u}^*_{\mathrm{f}} = 1, \bar{u}^*_{\mathrm{q}} = 1) \quad \text{for } t \in \left[t^*_{c,2}, T\right],
\end{aligned} \tag{S148}$$

where $t^*_{c,1}$ and $t^*_{c,2}$ in denote the switching times from ergonomic to reproductive phase and from male production to queen production, respectively, and they depend on the mode of control $c \in \{\mathrm{q}, \mathrm{w}, \mathrm{mx}\}$.

##### Single-party control

If the mortality of juvenile queens and males is equal (i.e. $\mu_{\mathrm{q}} = \mu_{\mathrm{m}} = \mu_{\mathrm{r}}$) then the switching time $t^*_{c,1}$ simplifies to

$$t^*_{c,1} = T - \frac{\ln\left(1 + \frac{\mu_{\mathrm{r}}-\mu_{\mathrm{w}}}{b}\right)}{\mu_{\mathrm{r}} - \mu_{\mathrm{w}}} \text{ for } c \in \{\mathrm{q}, \mathrm{w}\}, \tag{S149}$$

which is equal to the switching time obtained for single-party control under delayed dispersal (eq. S117). For equal juvenile queen and male mortality (i.e. $\mu_{\mathrm{q}} = \mu_{\mathrm{m}} = \mu_{\mathrm{r}}$) the switching time $t^*_{c,2}$ simplifies to

$$\begin{aligned} t^*_{\mathrm{q},2} &= T - \frac{1}{(\mu_{\mathrm{r}} - \mu_{\mathrm{w}})} \ln\left(\frac{b + \mu_{\mathrm{r}} - \mu_{\mathrm{w}}}{b + \frac{1}{2}(\mu_{\mathrm{r}} - \mu_{\mathrm{w}})}\right) && \text{(queen control)}, \\ t^*_{\mathrm{w},2} &= T - \frac{1}{(\mu_{\mathrm{r}} - \mu_{\mathrm{w}})} \ln\left(\frac{b + \mu_{\mathrm{r}} - \mu_{\mathrm{w}}}{b + \frac{M}{2(1+M)}(\mu_{\mathrm{r}} - \mu_{\mathrm{w}})}\right) && \text{(worker control)}. \end{aligned} \tag{S150}$$

In the limit where the mortality of sexuals becomes equal to the mortality of workers ($\mu_{\mathrm{r}} \to \mu_{\mathrm{w}}$) the switching times $t^*_{c,1}$ and $t^*_{c,2}$ simplify to

$$\begin{aligned} t^*_{c,1} &= T - \frac{1}{b} && \text{(queen and worker control, } c = \{\mathrm{q}, \mathrm{w}\}), \\ t^*_{c,2} &= \begin{cases} T - \dfrac{1}{2b}, & \text{(queen control, } c = \mathrm{q}), \\ T - \dfrac{1}{2b}\dfrac{2+M}{(1+M)}, & \text{(worker control, } c = \mathrm{w}). \end{cases} \end{aligned} \tag{S151}$$

Hence, when the mortality rate of sexuals is roughly equal to the mortality rate of workers, then the switching time from the ergonomic to the reproductive phase $t^*_{c,1}$ under single-party control ($c = \{\mathrm{q}, \mathrm{w}\}$) approaches to one generation time $(1/b)$ before the end of the season, i.e. only the last generation of brood is reproductive.

**Mixed control**

We showed previously that if the mortality rates of queens and males are equal (i.e. $\mu_{\mathrm{q}} = \mu_{\mathrm{m}} = \mu_{\mathrm{r}}$) then the switching time $t^*_{\mathrm{mx},1}$ from the ergonomic to the reproductive phase can be expressed as

$$t^*_{\mathrm{mx},1} = T - \frac{\ln\left(1 + \theta_{\mathrm{mx}} \frac{\mu_{\mathrm{r}} - \mu_{\mathrm{w}}}{b}\right)}{\mu_{\mathrm{r}} - \mu_{\mathrm{w}}}, \tag{S152}$$

where

$$\theta_{\mathrm{mx}} = \frac{2+M}{1+M}. \tag{S153}$$

and the he switching time $t^*_{\mathrm{mx},2}$ from the male production to the queen production can be expressed as

$$t^*_{\mathrm{mx},2} = T - \frac{\ln\left(2 - \frac{b}{b + \frac{1}{2}\theta_{\mathrm{mx}}(\mu_{\mathrm{r}} - \mu_{\mathrm{w}})}\right)}{\mu_{\mathrm{r}} - \mu_{\mathrm{w}}}. \tag{S154}$$

In the limit where the mortality of sexuals becomes equal to the mortality of workers ($\mu_{\mathrm{r}} \to \mu_{\mathrm{w}}$) the switching times $t^*_{\text{mx},1}$ and $t^*_{\text{mx},2}$ simplify to

$$\begin{aligned} t^*_{\text{mx},1} &= T - \frac{\theta_{\text{mx}}}{b}, \\ t^*_{\text{mx},2} &= T - \frac{\theta_{\text{mx}}}{2b}. \end{aligned} \tag{S155}$$

Hence, when the mortality rate of sexuals is roughly equal to the mortality rate of workers, under mixed control the switch happens $\theta_{\text{mx}}$ generations earlier. For example, when females mate only once ($M = 1$) the switch to reproductive phase happens one and a half generations before the end of the season.

##### 6.3 The cases $R_c\mu_{\text{q}} < \mu_{\text{m}}$ (single-party control) and $R_{\text{q}}\mu_{\text{q}} < \mu_{\text{m}}$ (mixed control)

It follows from eq. (S132) that if $R_c\mu_{\text{q}} < \mu_{\text{m}}$ (single-party control) or $R_{\text{q}}\mu_{\text{q}} < \mu_{\text{m}}$ (mixed control), then $u^*_{\text{f}}(t)$ can possibly be a singular arc during some period before $t^*_{c,2}$, where $R_c$ is the relatedness asymmetry associated with party $c$ in control of the trait of type f. Lets denote this singular arc by $\hat{u}^*_{\text{f}}(t) = \hat{u}_{\text{f,Bulmer}}(t)$, since it was originally derived under full queen control by Bulmer (1983). Furthermore, if the singular arc $\hat{u}_{\text{f,Bulmer}}(t)$ exists, it has to satisfy

$$\left(\frac{\mathrm{d}}{\mathrm{d}t}\right)^i \left.\frac{\partial H_{c,d}(\mathbf{u}(t), \mathbf{x}^*(t), \boldsymbol{\lambda}(t))}{\partial u_{\text{f}}}\right|_{\mathbf{u}=\mathbf{v}=\mathbf{u}^*} = \left(\frac{\mathrm{d}}{\mathrm{d}t}\right)^i \left[b x^*_{\text{w}}(t) \left(u^*_{\text{q}}(t)\sigma^c_1(t) - \sigma^c_2(t)\right)\right] = 0 \qquad i = 0, 1, 2, ... \tag{S156}$$

(e.g., Bryson and Ho, 1975, p. 248). And since we have shown previously that $\bar{u}^*_{\text{q}} = 1$ during the penultimate phase and hence $x^*_{\text{w}}(t) > 0$ it follows that $\hat{u}_{\text{f,Bulmer}}(t)$ has to satisfy

$$\left(\frac{\mathrm{d}}{\mathrm{d}t}\right)^i (\sigma^c_1(t) - \sigma^c_2(t)) = 0 \qquad i = 0, 1, 2, ... \,. \tag{S157}$$

We have already shown that $(\dot{\sigma}^c_1(t) - \dot{\sigma}^c_2(t)) = 0$ leads to to eq. (S132). Furthermore, $(\ddot{\sigma}_1(t) - \ddot{\sigma}_2(t)) = 0$ together with eqs. (S78), (S79), (S91) and (S35) implies that

$$\frac{\ddot{\lambda}^{\text{q}}_{\text{m}}(t)}{\ddot{\lambda}^{\text{q}}_{\text{q}}(t)} = R_c, \tag{S158}$$

Considering that $a^*_{\text{q}}(t) = u^*_{\text{f}}(t)u^*_{\text{q}}(t) = \hat{u}_{\text{f,Bulmer}}(t)\bar{u}^*_{\text{q}}$ and $\bar{u}^*_{\text{q}} = 1$ during the penultimate phase and substituting the costate equations (S82) into (S158) yields

$$\frac{\mu_{\text{m}}\dot{\lambda}^{\text{q}}_{\text{m}} - \frac{\mathrm{d}}{\mathrm{d}t}\left(\frac{b\hat{u}_{\text{f,Bulmer}}(t)x^*_{\text{w}}(t)}{x^*_{\text{m}}(t)}\right)\lambda^{\text{q}}_{\text{iq}} - \left(\frac{b\hat{u}_{\text{f,Bulmer}}(t)(t)x^*_{\text{w}}(t)}{x^*_{\text{m}}(t)}\right)\dot{\lambda}^{\text{q}}_{\text{iq}}}{\mu_{\text{q}}\dot{\lambda}^{\text{q}}_{\text{q}}} = R_c, \tag{S159}$$

Substituting eq. (S132) and considering eqs. (S90) and (S91) yields

$$\frac{\mathrm{d}}{\mathrm{d}t}\left(\frac{b\hat{u}_{\mathrm{f,Bulmer}}(t)x_{\mathrm{w}}^{*}(t)}{x_{\mathrm{m}}^{*}(t)}\right)=0, \tag{S160}$$

It follows from eq. (S160) that $\hat{u}_{\mathrm{f,Bulmer}}(t_{c,1}^{*})=0$, since $x_{\mathrm{m}}^{*}(t_{c,1}^{*})=0$. Using the quotient rule of taking derivatives yields

$$\frac{\mathrm{d}\hat{u}_{\mathrm{f,Bulmer}}(t)}{\mathrm{d}t}x_{\mathrm{w}}^{*}(t)x_{\mathrm{m}}^{*}(t)+\hat{u}_{\mathrm{f,Bulmer}}(t)\dot{x}_{\mathrm{w}}^{*}(t)x_{\mathrm{m}}^{*}(t)-\hat{u}_{\mathrm{f,Bulmer}}(t)x_{\mathrm{w}}^{*}(t)\dot{x}_{\mathrm{m}}^{*}(t)=0. \tag{S161}$$

Substituting eqs. (S57) and (S58) into (S161) implies the following differential equation

$$\frac{\mathrm{d}\hat{u}_{\mathrm{f,Bulmer}}(t)}{\mathrm{d}t}=(1-\hat{u}_{\mathrm{f,Bulmer}}(t))(\mu_{\mathrm{m}}-R_{c}\mu_{\mathrm{q}})-\hat{u}_{\mathrm{f,Bulmer}}(t)(\mu_{\mathrm{m}}-\mu_{\mathrm{w}}). \tag{S162}$$

Solving the differential equation for $\hat{u}_{\mathrm{f,Bulmer}}(t)$ with initial condition $\hat{u}_{\mathrm{f,Bulmer}}(t_{c,1}^{*})=0$ gives

$$\hat{u}_{\mathrm{f,Bulmer}}(t)=\frac{(\mu_{\mathrm{m}}-R_{c}\mu_{\mathrm{q}})\left(e^{(t-t_{c,1}^{*})(\mu_{\mathrm{w}}+R\mu_{\mathrm{q}}-2\mu_{\mathrm{m}})}-1\right)}{\mu_{\mathrm{w}}+R_{c}\mu_{\mathrm{q}}-2\mu_{\mathrm{m}}}. \tag{S163}$$

Thus far we have derived the singular arc from the first and second time derivative of the coefficient $(\sigma_{1}^{c}(t)-\sigma_{2}^{c}(t))=0$. However, it follows from eq. (S131) that the control variable $\hat{u}_{\mathrm{f,Bulmer}}$ first appears in the odd member ($i$ is odd) in the sequence given by eq. (S157) (i.e. the degree of singularity of the singular arc is odd). It has been proven that if the degree of singularity of the singular arc is odd then it is necessarily non-optimal (Robbins, 1967). This means that if the control variable first appears in the time derivative of the coefficient $(\sigma_{1}^{c}(t)-\sigma_{2}^{c}(t))$ to an odd order, then this singular arc is non-optimal.

Hence, we will only rely on numerical solutions in order to approximate the uninvadable allocation schedule $u_{\mathrm{f}}^{*}$ if $R_{c}\mu_{\mathrm{q}}<\mu_{\mathrm{m}}$. Our numerical solutions indicate that under single-party control $u_{\mathrm{f}}^{*}(t)$ is close to 0 during the penultimate phase $t\in[t_{c,1}^{*},t_{c,2}^{*}]$ if $R_{c}\mu_{\mathrm{q}}<\mu_{\mathrm{m}}$. In Fig. S3 we demonstrate for single-party control that even if the mortality of queens is 20 times lower than that of males, approximately only males are produced in the penultimate phase $t\in[t_{c,1}^{*},t_{c,2}^{*}]$. Hence, we find that under single-party control, for a large set of biologically realistic parameter values, approximately only males are produced in the penultimate phase.

We also observe from Fig. S3 that under mixed control $u_{\mathrm{f}}^{*}(t)=\hat{u}_{\mathrm{f}}^{*}(t)>0$ during the penultimate phase $t\in[t_{c,1}^{*},t_{c,2}^{*}]$ if $R_{c}\mu_{\mathrm{q}}<\mu_{\mathrm{m}}$. Hence under mixed control, we predict that males and queens are produced simultaneously under mixed control during the penultimate phase $t\in[t_{c,1}^{*},t_{c,2}^{*}]$ if $R_{c}\mu_{\mathrm{q}}<\mu_{\mathrm{m}}$.

We find that even if the mortality rate of queens is significantly lower than that of males, the overall sex allocation ratio $S_{c,\mathrm{dir}}$ under single-party control is only slightly more female-biased than the uninvadable sex allocation ratio predicted from the standard static models of sex allocation theory (Boomsma and Grafen, 1991; Reuter and Keller, 2001; Trivers and Hare, 1976). We find that the overall sex allocation ratio $S_{\mathrm{mx,dir}}$ under mixed control is close to the overall sex allocation ratio $S_{\mathrm{q,dir}}$ under full queen control (see Figs. S3–S4).

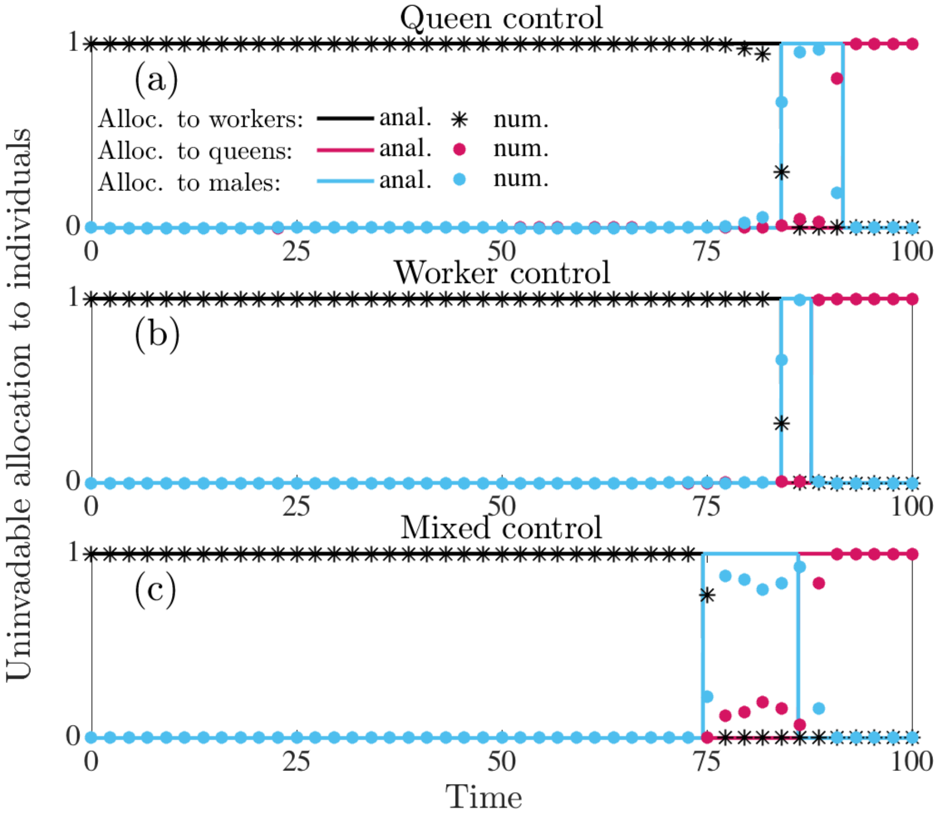

Figure S3: Uninvadable proportional allocation (under direct dispersal) to workers $a_{\mathrm{w}}^*(t) = u_{\mathrm{f}}^*(t)(1 - u_{\mathrm{q}}^*(t))$ (black), queens $a_{\mathrm{q}}^*(t) = u_{\mathrm{f}}^*(t)u_{\mathrm{q}}^*(t)$ (red), and males $a_{\mathrm{m}}^*(t) = (1 - u_{\mathrm{f}}^*(t))$ (blue). Panel (a): queen control. Panel (b): worker control. Panel (c): mixed control. Parameter values: $M = 1$ (queen monandry), $b = 0.07$, $\mu_{\mathrm{w}} = 0.015$, $\mu_{\mathrm{q}} = 0.001$, $\mu_{\mathrm{m}} = 0.02$, $T = 100$. Results here are only numerically derived and the correspondingly colored lines are analytically predicted results assuming that $\mu_{\mathrm{q}} = \mu_{\mathrm{m}} = 0.001$. Notice that these analytical predictions approximate the numerically derived predictions quite well under single party control.

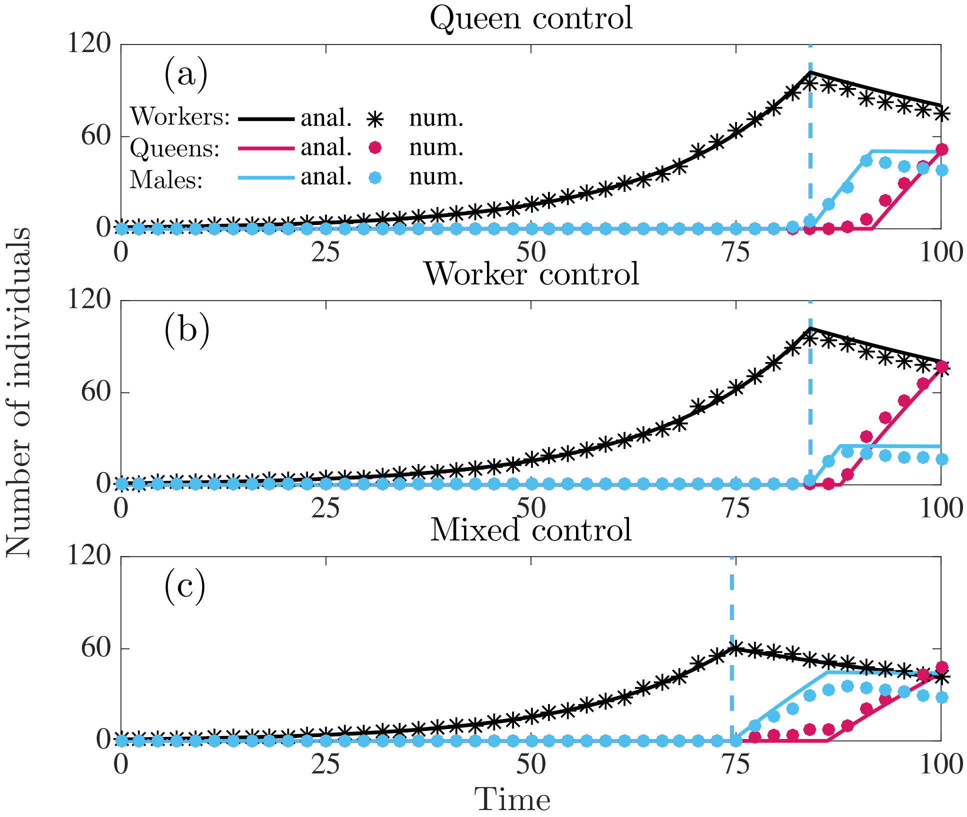

Figure S4: Number of individuals produced in a colony following the uninvadable resource allocation schedule $\mathbf{u}^*$ under direct dispersal. Panel (a): queen control. Panel (b): worker control. Panel (c): mixed control. Parameter values: $M = 1$ (queen monandry), $b = 0.07$, $\mu_\mathrm{w} = 0.015$, $\mu_\mathrm{q} = 0.001$, $\mu_\mathrm{m} = 0.02$, $T = 100$. The (numerical) overall sex allocation ratio $S_\mathrm{q} \approx 0.51$, $S_\mathrm{w} \approx 0.78$, $S_\mathrm{mx} \approx 0.53$. The correspondingly colored lines are analytically predicted results assuming that $\mu_\mathrm{q} = \mu_\mathrm{m} = 0.001$. Notice that these analytical predictions approximate the numerically derived predictions quite well under single party control.

#### 7 Macroscopic quantities describing resource allocation in colonies

##### 7.1 Colony size at maturity

It follows from eq. (S58) assuming that the allocation schedule to individuals corresponds to the uninvadable allocation schedule $\mathbf{u}^*$, given by eq. (S92) (for delayed dispersal) and eq. (S124) (for delayed dispersal) that during the ergonomic phase the number of workers grows exponentially at rate $b - \mu_\mathrm{w}$. Furthermore, the number of workers $x_\mathrm{w}^*(t_{c,1}^*)$ at the switching time $t_{c,1}^*$ from the ergonomic phase to the reproductive phase determines the colony size at maturity, which is given by

$$x_\mathrm{w}^*(t_{c,1}^*) = y_{\mathrm{w},0} e^{(b-\mu_\mathrm{w})t_{c,1}^*} = e^{(b-\mu_\mathrm{w})t_{c,1}^*}, \tag{S164}$$

and owing to mortality of workers it is also the maximal colony size.

#### 7.2 Colony productivity

The switching time $t^*_{c,1}$ also determines the colony productivity, which we define as the total number of males and females produced that have survived until the end of the season

$$B(t^*_{c,1}) = x^*_{\mathrm{q}}(\mathbf{u}^*) + x^*_{\mathrm{m}}(\mathbf{u}^*). \tag{S165}$$

Substituting eq. (S58) for state variables into eq. (S165) assuming that the allocation schedule to individuals corresponds to the uninvadable allocation schedule $\mathbf{u}^*$ under delayed dispersal (given by eq. S92) and the mortality rate of queens and males is equal ($\mu_{\mathrm{q}} = \mu_{\mathrm{m}} = \mu_{\mathrm{r}}$)

$$\begin{aligned}
B(t^*_{c,1}) &= \int_0^T b a^*_{\mathrm{q}}(t) x^*_{\mathrm{w}}(t) e^{-\mu_{\mathrm{q}}(T-t)}\,\mathrm{d}t + \int_0^T b a^*_{\mathrm{m}}(t) x^*_{\mathrm{w}}(t) e^{-\mu_{\mathrm{q}}(T-t)}\,\mathrm{d}t \\
&= \int_{t^*_{c,1}}^T b\hat{\hat{u}}^*_{\mathrm{f}} x^*_{\mathrm{w}}(t) e^{-\mu_{\mathrm{q}}(T-t)}\,\mathrm{d}t + \int_{t^*_{c,1}}^T b(1-\hat{\hat{u}}^*_{\mathrm{f}}) x^*_{\mathrm{w}}(t) e^{-\mu_{\mathrm{q}}(T-t)}\,\mathrm{d}t \\
&= \int_{t^*_{c,1}}^T b x^*_{\mathrm{w}}(t) e^{-\mu_{\mathrm{q}}(T-t)}\,\mathrm{d}t \\
&= \frac{b e^{b t^*_{c,1}} \left( e^{-\mu_{\mathrm{w}} T} - e^{-\mu_{\mathrm{r}} T + (\mu_{\mathrm{r}} - \mu_{\mathrm{w}}) t^*_{c,1}} \right)}{\mu_{\mathrm{r}} - \mu_{\mathrm{w}}}.
\end{aligned} \tag{S166}$$

Substituting eq. (S58) for state variables into eq. (S165) assuming that the allocation schedule to individuals corresponds to the uninvadable allocation schedule $\mathbf{u}^*$ under direct dispersal (given by eq. S124) and the mortality rate of queens and males is equal ($\mu_{\mathrm{q}} = \mu_{\mathrm{m}} = \mu_{\mathrm{r}}$)

$$\begin{aligned}
B(t^*_{c,1}) &= \int_0^T b a^*_{\mathrm{q}}(t) x^*_{\mathrm{w}}(t) e^{-\mu_{\mathrm{q}}(T-t)}\,\mathrm{d}t + \int_0^T b a^*_{\mathrm{m}}(t) x^*_{\mathrm{w}}(t) e^{-\mu_{\mathrm{q}}(T-t)}\,\mathrm{d}t \\
&= \int_{t^*_{c,2}}^T b x^*_{\mathrm{w}}(t) e^{-\mu_{\mathrm{q}}(T-t)}\,\mathrm{d}t + \int_{t^*_{c,1}}^{t^*_{c,2}} b x^*_{\mathrm{w}}(t) e^{-\mu_{\mathrm{q}}(T-t)}\,\mathrm{d}t \\
&= \int_{t^*_{c,1}}^T b x^*_{\mathrm{w}}(t) e^{-\mu_{\mathrm{q}}(T-t)}\,\mathrm{d}t \\
&= \frac{b e^{b t^*_{c,1}} \left( e^{-\mu_{\mathrm{w}} T} - e^{-\mu_{\mathrm{r}} T + (\mu_{\mathrm{r}} - \mu_{\mathrm{w}}) t^*_{c,1}} \right)}{\mu_{\mathrm{r}} - \mu_{\mathrm{w}}}.
\end{aligned} \tag{S167}$$

Hence, it follows from eqs. (S166) and (S167) that colony productivity (for delayed and direct dispersal) can be expressed as

$$B(t^*_{c,1}) = \frac{b e^{b t^*_{c,1}} \left( e^{-\mu_{\mathrm{w}} T} - e^{-\mu_{\mathrm{r}} T + (\mu_{\mathrm{r}} - \mu_{\mathrm{w}}) t^*_{c,1}} \right)}{\mu_{\mathrm{r}} - \mu_{\mathrm{w}}}. \tag{S168}$$

We can determine the switching time $t^*_{c,1}$ that maximizes colony productivity from

$$\frac{\mathrm{d}B(t^*_{c,1})}{\mathrm{d}t^*_{c,1}} = 0. \tag{S169}$$

Substituting eq. (S168) into eq. (S169) implies

$$\frac{\mathrm{d}B(t^*_{c,1})}{\mathrm{d}t^*_{c,1}} = be^{bt^*_{c,1}} \left( \left( \frac{b}{\mu_\mathrm{w} - \mu_\mathrm{r}} - 1 \right) e^{-\mu_\mathrm{r}T + (\mu_\mathrm{r} - \mu_\mathrm{w})t^*_{c,1}} + \frac{be^{-\mu_\mathrm{w}T}}{\mu_\mathrm{r} - \mu_\mathrm{w}} \right) = 0. \tag{S170}$$

Solving eq. (S170) for $t^*_{c,1}$ yields

$$t^*_{c,1} = T - \frac{\ln\left(1 + \frac{\mu_\mathrm{r} - \mu_\mathrm{w}}{b}\right)}{\mu_\mathrm{r} - \mu_\mathrm{w}}. \tag{S171}$$

The switching time given by eq. (S171) that maximizes the colony productivity is equal to the switching time $t^*_{c,1}$ under single-party control ($c \in \{\mathrm{q}, \mathrm{w}\}$) for both delayed (given by eq. S117) and direct dispersal (given by eq. S149) assuming that the mortality rates of queens and males are equal ($\mu_\mathrm{q} = \mu_\mathrm{m} = \mu_\mathrm{r}$).

#### 7.3 Overall sex allocation ratio

We define the overall sex allocation ratio as the proportion of the colony resources allocated to queens from the resources allocated to sexuals over the entire season (irrespective of whether they survive to reproduce), and it is thus given by

$$S_c = \frac{\int_0^T ba^*_\mathrm{q}(t)x^*_\mathrm{w}(t)\,\mathrm{d}t}{\int_0^T ba^*_\mathrm{q}(t)x^*_\mathrm{w}(t)\,\mathrm{d}t + \int_0^T ba^*_\mathrm{m}(t)x^*_\mathrm{w}(t)\,\mathrm{d}t}. \tag{S172}$$

**Sex allocation ratio under delayed dispersal**

Substituting the uninvadable allocation schedule $\mathbf{u}^*$ for delayed dispersal given by eq. (S92) with the solutions to the state equations given by (S89) into eq. (S172) yields

$$\begin{aligned} S_c =& \frac{\int_{t^*_{c,1}}^T b\hat{u}^*_\mathrm{f}x^*_\mathrm{w}(t)\,\mathrm{d}t}{\int_{t^*_{c,1}}^T b\hat{u}^*_\mathrm{f}x^*_\mathrm{w}(t)\,\mathrm{d}t + \int_{t^*_{c,1}}^T b(1-\hat{u}^*_\mathrm{f})x^*_\mathrm{w}(t)\,\mathrm{d}t} = \frac{x^*_\mathrm{w}(t^*_{c,1})\int_{t^*_{c,1}}^T \hat{u}^*_\mathrm{f}e^{-\mu_\mathrm{w}(t-t^*_{c,1})}\,\mathrm{d}t}{x^*_\mathrm{w}(t^*_{c,1})\int_{t^*_{c,1}}^T e^{-\mu_\mathrm{w}(t-t^*_{c,1})}\,\mathrm{d}t} = \\ =& \frac{\int_{t^*_{c,1}}^T \hat{u}^*_\mathrm{f}e^{-\mu_\mathrm{w}(t-t^*_{c,1})}\,\mathrm{d}t}{\int_{t^*_{c,1}}^T e^{-\mu_\mathrm{w}(t-t^*_{c,1})}\,\mathrm{d}t}. \end{aligned} \tag{S173}$$

If males and queens are equally costly to produce, then $u_{\mathrm{f}}^*(t)$ is constant in the reproductive phase and is given by eq. (S104). Hence, eq. (S173) simplifies to

$$\begin{aligned} S_c &= \hat{\hat{u}}_{\mathrm{f}}^* = \frac{R_c}{1+R_c} \text{ for } c \in \{\mathrm{q},\mathrm{w}\} && \text{(single-party control)} \\ S_{\mathrm{mx}} &= \hat{\hat{u}}_{\mathrm{f}}^* = \frac{R_{\mathrm{q}}}{1+R_{\mathrm{q}}} && \text{(mixed control),} \end{aligned} \tag{S174}$$

where $R_c$ is the relatedness asymmetry given by eq. (S35) and for haplodiploids (S174) simplifies to

$$\begin{aligned} S_{\mathrm{q}} &= \hat{\hat{u}}_{\mathrm{f}}^* = \frac{1}{2} && \text{(queen control)} \\ S_{\mathrm{w}} &= \hat{\hat{u}}_{\mathrm{f}}^* = \frac{2+M}{2(1+M)} && \text{(worker control)} \\ S_{\mathrm{mx}} &= \hat{\hat{u}}_{\mathrm{f}}^* = \frac{1}{2} && \text{(mixed control).} \end{aligned} \tag{S175}$$

**Sex allocation ratio under direct dispersal**

Recall that the overall sex allocation ratio (the proportion of the colony resources allocated to queens from the resources allocated to sexuals over the entire season) is

$$S_c = \frac{\int_0^T b a_{\mathrm{q}}^*(t) x_{\mathrm{w}}^*(t)\,\mathrm{d}t}{\int_0^T b a_{\mathrm{q}}^*(t) x_{\mathrm{w}}^*(t)\,\mathrm{d}t + \int_0^T b a_{\mathrm{m}}^*(t) x_{\mathrm{w}}^*(t)\,\mathrm{d}t}. \tag{S176}$$

Substituting the uninvadable resource allocation schedule $\mathbf{u}^*$ for $\mu_{\mathrm{m}} \leq \mu_{\mathrm{q}}$ under direct dispersal given by eq. (S124) with the solutions to state equations given by (S89) into eq. (S176) yields

$$\begin{aligned} S_c =& \frac{\int_{t_{c,2}^*}^T b x_{\mathrm{w}}^*(t)\,\mathrm{d}t}{\int_{t_{c,2}^*}^T b x_{\mathrm{w}}^*(t)\,\mathrm{d}t + \int_{t_{c,1}^*}^{t_{c,2}^*} b x_{\mathrm{w}}^*(t)\,\mathrm{d}t} = \frac{\int_{t_{c,2}^*}^T b x_{\mathrm{w}}^*(t)\,\mathrm{d}t}{\int_{t_{c,1}^*}^T b x_{\mathrm{w}}^*(t)\,\mathrm{d}t} = \frac{x_{\mathrm{w}}^*(t_{c,2}^*) \int_{t_{c,2}^*}^T e^{-\mu_{\mathrm{w}}(t-t_{c,2}^*)}\,\mathrm{d}t}{x_{\mathrm{w}}^*(t_{c,1}^*) \int_{t_{c,1}^*}^T e^{-\mu_{\mathrm{w}}(t-t_{c,1}^*)}\,\mathrm{d}t} \\ =& \frac{x_{\mathrm{w}}^*(t_{c,2}^*) e^{\mu_{\mathrm{w}} t_{c,2}^*} \int_{t_{c,2}^*}^T e^{-\mu_{\mathrm{w}} t}\,\mathrm{d}t}{x_{\mathrm{w}}^*(t_{c,1}^*) e^{\mu_{\mathrm{w}} t_{c,1}^*} \int_{t_{c,1}^*}^T e^{-\mu_{\mathrm{w}} t}\,\mathrm{d}t} = \frac{x_{\mathrm{w}}^*(t_{c,1}^*) e^{-\mu_{\mathrm{w}}(t_{c,2}^*-t_{c,1}^*)} e^{\mu_{\mathrm{w}} t_{c,2}^*} (-1/\mu_{\mathrm{w}})[e^{-\mu_{\mathrm{w}} T} - e^{-\mu_{\mathrm{w}} t_{c,2}^*}]}{x_{\mathrm{w}}^*(t_{c,1}^*) e^{\mu_{\mathrm{w}} t_{c,1}^*} (-1/\mu_{\mathrm{w}})[e^{-\mu_{\mathrm{w}} T} - e^{-\mu_{\mathrm{w}} t_{c,1}^*}]} \\ =& \frac{e^{-\mu_{\mathrm{w}} t_{c,2}^*} - e^{-\mu_{\mathrm{w}} T}}{e^{-\mu_{\mathrm{w}} t_{c,1}^*} - e^{-\mu_{\mathrm{w}} T}} \end{aligned} \tag{S177}$$

Hence, if $R_c \mu_{\mathrm{m}} > \mu_{\mathrm{q}}$ under single-party control and $\mu_{\mathrm{m}} > \mu_{\mathrm{q}}$ under mixed control, then the overall sex allocation ratio under direct dispersal is

$$S_c = \frac{e^{-\mu_{\mathrm{w}} t_{c,2}^*} - e^{-\mu_{\mathrm{w}} T}}{e^{-\mu_{\mathrm{w}} t_{c,1}^*} - e^{-\mu_{\mathrm{w}} T}}. \tag{S178}$$

### 8 Marginal return of changing the allocation trait for the ergonomic and reproductive phase under mixed control

The aim of this section is to show that under mixed control the queen determines the overall sex allocation ratio and workers determine the switching time $t^*_{\mathrm{mx},1}$ from the ergonomic to the reproductive phase (assuming equal mortality of males and queens, i.e $\mu_\mathrm{q} = \mu_\mathrm{m} = \mu_\mathrm{r}$). We do this by analyzing the marginal return $\partial H_{c,d}(\mathbf{u}(t), \mathbf{x}^*(t), \boldsymbol{\lambda}(t))/\partial u_\tau(t) = \partial H_{c,d}(t)/\partial u_\tau(t)$ of changing the allocation trait during the ergonomic ($\forall t \in [0, t^*_{\mathrm{mx},1}]$) and the reproductive phase ($\forall t \in [t^*_{\mathrm{mx},1}, T]$) under mixed control.

It follows from the first-order condition for uninvadability under mixed control (recall eq. S65 and eq. S73) for both delayed and direct dispersal ($d \in \{\mathrm{del}, \mathrm{dir}\}$) that

$$\begin{aligned}
&\left.\frac{\partial H_{\mathrm{q},d}(t)}{\partial u_\mathrm{f}(t)}\right|_{\mathbf{u}=\mathbf{v}=\mathbf{u}^*} \begin{cases} <0, & \Longrightarrow \quad u^*_\mathrm{f}(t) = 0 \\ =0, & \Longrightarrow \quad 0 \geq u^*_\mathrm{f}(t) = \hat{u}^*_\mathrm{f}(t) \geq 1 \\ >0, & \Longrightarrow \quad u^*_\mathrm{f}(t) = 1 \end{cases} \quad , \ \forall t \in [0,T], \\
&\left.\frac{\partial H_{\mathrm{w},d}(t)}{\partial u_\mathrm{q}(t)}\right|_{\mathbf{u}=\mathbf{v}=\mathbf{u}^*} \begin{cases} <0, & \Longrightarrow \quad u^*_\mathrm{q}(t) = 0 \\ =0, & \Longrightarrow \quad 0 \geq u^*_\mathrm{q}(t) = \hat{u}^*_\mathrm{q}(t) \geq 1 \\ >0, & \Longrightarrow \quad u^*_\mathrm{q}(t) = 1 \end{cases} \quad , \ \forall t \in [0,T].
\end{aligned} \tag{S179}$$

Hence, under mixed control the sign of $\partial H_{\mathrm{q},d}(t)/\partial u_\mathrm{f}(t)$, which is under queen control, determines $u^*_\mathrm{f}(t)$, while the sign of $\partial H_{\mathrm{w},d}(t)/\partial u_\mathrm{q}(t)$, which is under worker control, determines $u^*_\mathrm{q}(t)$.

Let sgn$(\cdot)$ denote a sign function, i.e.

$$\mathrm{sgn}(x) = \begin{cases} -1, & \text{if } x < 0, \\ 0, & \text{if } x = 0, \\ 1, & \text{if } x > 0. \end{cases} \tag{S180}$$

The signs of $\partial H_{c,d}(t)/\partial u_\tau(t)$ (assuming equal mortality of males and queens, i.e $\mu_\mathrm{q} = \mu_\mathrm{m} = \mu_\mathrm{r}$) can be inferred by way of eq. (S179) from the uninvadable allocation schedule $\mathbf{u}^*$ given by eq. (S92) for delayed dispersal and eq. (S148) for direct dispersal. Further, using eq. (S76), the fact that $b > 0$ and $x^*_\mathrm{w}(t) > 0$, and the switching functions eq. (S77) (where $p^\mathrm{q}_\mathrm{q} = 1/2$, $p^\mathrm{m}_\mathrm{q} = 0$, $p^\mathrm{q}_\mathrm{w} = 1/4$, and $p^\mathrm{m}_\mathrm{w} = 1/(2M)$), then we have for both delayed and direct dispersal ($d \in \{\mathrm{del}, \mathrm{dir}\}$) that during the ergonomic phase $\forall t \in [0, t^*_{c,1}]$:

$$\begin{aligned}
&\mathrm{sgn}\left(\left.\frac{\partial H_{\mathrm{q},d}(t)}{\partial u_\mathrm{f}(t)}\right|_{\mathbf{u}=\mathbf{v}=\mathbf{u}^*}\right) = \lambda^\mathrm{q}_\mathrm{w}(t) - \lambda^\mathrm{q}_\mathrm{m}(t) > 0 && \Longrightarrow u^*_\mathrm{f}(t) = 1 \\
&\mathrm{sgn}\left(\left.\frac{\partial H_{\mathrm{w},d}(t)}{\partial u_\mathrm{q}(t)}\right|_{\mathbf{u}=\mathbf{v}=\mathbf{u}^*}\right) = \left(\frac{1}{4}\lambda^\mathrm{q}_\mathrm{q}(t) + \frac{1}{2M}\lambda^\mathrm{m}_\mathrm{q}(t)\right) - \left(\frac{1}{4}\lambda^\mathrm{q}_\mathrm{w}(t) + \frac{1}{2M}\lambda^\mathrm{m}_\mathrm{w}(t)\right) < 0 && \Longrightarrow u^*_\mathrm{q}(t) = 0.
\end{aligned} \tag{S181}$$

Eq. (S181) shows that during the ergonomic phase the marginal return of workers is higher than that of queens and males. More precisely, the sign of $\partial H_{\mathrm{q},d}(t)/\partial u_{\mathrm{f}}(t)$ implies that during the ergonomic phase only females are produced ($u_{\mathrm{f}}^*(t)=1$), since the marginal return of producing workers $\lambda_{\mathrm{w}}^{\mathrm{q}}(t)$ is higher than that of males $\lambda_{\mathrm{m}}^{\mathrm{q}}(t)$ in colonies founded by queens carrying the mutant allele. In other words, during the ergonomic phase workers are more valuable than males to the genes residing in queens. Similarly, it follows from the the sign of $\partial H_{\mathrm{w},d}(t)/\partial u_{\mathrm{q}}(t)$ that all females produced during the ergonomic phase become workers ($u_{\mathrm{q}}^*(t)=0$), because the marginal return of queens $\lambda_{\mathrm{q}}^{s}(t)$ is lower that that of workers $\lambda_{\mathrm{w}}^{s}(t)$, where the marginal returns have been weighed by the expected frequency of mutant alleles in workers in colonies founded by type $s$ mutant individuals. In other words, workers are more valuable than queens to the genes residing in the workers. Hence, during the ergonomic phase, there is a latent trade-off between producing workers versus males from the perspective of the genes in the queens and a latent trade-off between producing workers versus queens from the perspective of the genes in the workers. Only workers are produced during the ergonomic phase, since workers have a higher marginal return for both parties.

During the reproductive phase $\forall t \in [t_{\mathrm{mx},1}^*, T]$, the signs of the marginal returns $\partial H_{c,\mathrm{del}}(t)/\partial u_{\tau}(t)$ are different for delayed and direct dispersal. Under delayed dispersal, we have

$$\begin{aligned}
&\operatorname{sgn}\left(\left.\frac{\partial H_{\mathrm{q,del}}(t)}{\partial u_{\mathrm{f}}(t)}\right|_{\mathbf{u}=\mathbf{v}=\mathbf{u}^*}\right) = \lambda_{\mathrm{q}}^{\mathrm{q}}(t) - \lambda_{\mathrm{m}}^{\mathrm{q}}(t) = 0 && \Longrightarrow\ u_{\mathrm{f}}^*(t) = \hat{u}_{\mathrm{f}}^* \\
&\operatorname{sgn}\left(\left.\frac{\partial H_{\mathrm{w,del}}(t)}{\partial u_{\mathrm{q}}(t)}\right|_{\mathbf{u}=\mathbf{v}=\mathbf{u}^*}\right) = \left(\frac{1}{4}\lambda_{\mathrm{q}}^{\mathrm{q}}(t) + \frac{1}{2M}\lambda_{\mathrm{q}}^{\mathrm{m}}(t)\right) - \left(\frac{1}{4}\lambda_{\mathrm{w}}^{\mathrm{q}}(t) + \frac{1}{2M}\lambda_{\mathrm{w}}^{\mathrm{m}}(t)\right) > 0 && \Longrightarrow\ u_{\mathrm{q}}^*(t) = 1.
\end{aligned} \tag{S182}$$

Under direct dispersal, we have during the time of male production $\forall t \in [t_{\mathrm{mx},1}^*, t_{\mathrm{mx},2}^*]$ that

$$\begin{aligned}
&\operatorname{sgn}\left(\left.\frac{\partial H_{\mathrm{q,dir}}(t)}{\partial u_{\mathrm{f}}(t)}\right|_{\mathbf{u}=\mathbf{v}=\mathbf{u}^*}\right) = \lambda_{\mathrm{q}}^{\mathrm{q}}(t) - \lambda_{\mathrm{m}}^{\mathrm{q}}(t) < 0 && \Longrightarrow\ u_{\mathrm{f}}^*(t) = 0 \\
&\operatorname{sgn}\left(\left.\frac{\partial H_{\mathrm{w,dir}}(t)}{\partial u_{\mathrm{q}}(t)}\right|_{\mathbf{u}=\mathbf{v}=\mathbf{u}^*}\right) = 0 && \Longrightarrow\ u_{\mathrm{q}}^*(t) = \tilde{u}_{\mathrm{q}}^*(t),
\end{aligned} \tag{S183}$$

while during the time of queen production $\forall t \in [t_{\mathrm{mx},2}^*, T]$:

$$\begin{aligned}
&\operatorname{sgn}\left(\left.\frac{\partial H_{\mathrm{q,dir}}(t)}{\partial u_{\mathrm{f}}(t)}\right|_{\mathbf{u}=\mathbf{v}=\mathbf{u}^*}\right) = \lambda_{\mathrm{q}}^{\mathrm{q}}(t) - \lambda_{\mathrm{m}}^{\mathrm{q}}(t) > 0 && \Longrightarrow\ u_{\mathrm{f}}^*(t) = 1 \\
&\operatorname{sgn}\left(\left.\frac{\partial H_{\mathrm{w,dir}}(t)}{\partial u_{\mathrm{q}}(t)}\right|_{\mathbf{u}=\mathbf{v}=\mathbf{u}^*}\right) = \left(\frac{1}{4}\lambda_{\mathrm{q}}^{\mathrm{q}}(t) + \frac{1}{2M}\lambda_{\mathrm{q}}^{\mathrm{m}}(t)\right) - \left(\frac{1}{4}\lambda_{\mathrm{w}}^{\mathrm{q}}(t) + \frac{1}{2M}\lambda_{\mathrm{w}}^{\mathrm{m}}(t)\right) > 0 && \Longrightarrow\ u_{\mathrm{q}}^*(t) = 1.
\end{aligned} \tag{S184}$$

The sign of $\partial H_{\mathrm{q},d}(t)/\partial u_{\mathrm{f}}(t)$ during the reproductive phase implies that females and males are produced simultaneously ($u_{\mathrm{f}}^*(t)=\hat{u}_{\mathrm{f}}^*$) under delayed dispersal (eq. S182) since the marginal return of queens $\lambda_{\mathrm{q}}^{\mathrm{q}}(t)$ and males $\lambda_{\mathrm{m}}^{\mathrm{q}}(t)$ is equal in colonies founded by queens carrying the mutant allele. However, under direct dispersal (eqs. S183 and S184) males are produced first, since the marginal return of males $\lambda_{\mathrm{m}}^{\mathrm{q}}(t)$ is initially higher and then becomes lower than that of queens $\lambda_{\mathrm{q}}^{\mathrm{q}}(t)$ in colonies founded by queens carrying the mutant allele. The sign of $\partial H_{\mathrm{w},d}(t)/\partial u_{\mathrm{q}}(t)$ during the reproductive phase (eqs. S182, S183 and S184) implies that if any females are produced ($u_{\mathrm{f}}^* \neq 0$ like in eq. S183) then all females become queens during the reproductive phase ($u_{\mathrm{q}}^*(t) = 1$), because the marginal return of queens $\lambda_{\mathrm{q}}^s(t)$ is higher that that of workers $\lambda_{\mathrm{w}}^s(t)$, where the marginal returns have been weighed by the expected frequency of mutant alleles in workers in colonies founded by type $s$ mutant individuals. In other words, queens are more valuable than workers to the genes residing in the workers. Under direct dispersal, during the production of males (eq. S183) the sign of $\partial H_{\mathrm{w,dir}}(t)/\partial u_{\mathrm{q}}(t) = 0$ because $u_{\mathrm{f}}^* = 0$. Hence, there is no directional selection on $u_{\mathrm{q}}(t)$ for this time period, since it has no effect on invasion fitness (i.e. during the time only males are produced, the proportion at which workers rear female eggs into queens does not affect invasion fitness). Hence, during the reproductive phase, there is a latent trade-off between producing queens versus males from the perspective of the genes in the queens and a latent trade-off between producing workers versus queens from the perspective of the genes in the workers.

In summary, the signs of marginal returns $\partial H_{c,d}(t)/\partial u_\tau(t)$ (eqs. S182, S183 and S184) defined over the entire season elucidate the main difference between the ergonomic and reproductive phase in terms of trade-offs between producing different individuals experienced by the two parties. From the perspective of the genes in the workers there is always a trade-off between producing queens versus workers, such that the balance is tipped in favour of workers during the ergonomic phase and in favour of queens during the reproductive phase. From the perspective of the genes in the queens, the trade-offs for the ergonomic and the reproductive phase are different. During the ergonomic phase, there is a trade-off between producing workers versus males and during the reproductive phase there is a trade-off between producing queens versus males. Intuitively, this "decoupling" of the trade-offs from the perspective of the queens happens because queens and workers are produced in separate phases. Hence, during the ergonomic phase the trade-off is only between producing workers and males, since workers rear all female eggs into workers and during the reproductive phase the trade-off is only between producing queens and males, since workers rear all female eggs into queens. We can now turn to explaining how the trade-offs experienced by the two parties during different phases of colony growth have shaped the evolutionary outcome of the sex allocation conflict.

The queen determines the overall sex allocation ratio $S_{\mathrm{mx}}$ under mixed control. To see this, first note that the primary sex allocation ratio $u_{\mathrm{f}}^*(t)$ during the reproductive determines the overall sex allocation ratio $S_{\mathrm{mx}}$, since all females become queens (i.e. $u_{\mathrm{q}}^*(t) = 1$) at the reproductive phase and sexuals are only produced during the reproductive phase (recall eqs. S92 and S148). It follows from eqs. (S182), (S183), (S184) that $u_{\mathrm{f}}^*(t)$ is determined by the difference between the marginal return of producing queens and males in a colony founded by queens carrying the mutant allele, i.e. the genes in the queens determines the overall sex allocation ratio $S_{\mathrm{mx}}$.

### 9 Marginal return of producing a queen versus a male

The overall sex allocation ratio $S_c$ is determined by the allocation trait $u^*_{\mathrm{f}}(t)$ during the reproductive phase (see sections 7.3 and 8). This means that allocation to queens versus males is determined by the sign of the marginal return $\partial H_{c,d}(t)/\partial u_{\mathrm{f}}(t)$ during the reproductive phase, which is given by eq. (S76) assuming that $u^*_{\mathrm{q}}(t) = 1$, which yields

$$\left.\frac{\partial H_{c,d}(\mathbf{u}(t), \mathbf{x}^*(t), \boldsymbol{\lambda}(t))}{\partial u_{\mathrm{f}}(t)}\right|_{\mathbf{u}=\mathbf{v}=\mathbf{u}^*} = b x^*_{\mathrm{w}}(t) \left(\sigma^c_1(t) - \sigma^c_2(t)\right). \tag{S185}$$

Substituting $\sigma^c_1(t) - \sigma^c_2(t)$ from eq. (S78) for queen control and eqs. (S79) and (S80) for worker control and using the expression for relatedness asymmetry $R_c$ (eq. S35), we obtain

$$\begin{aligned}
\left.\frac{\partial H_{\mathrm{q,dir}}(t)}{\partial u_{\mathrm{f}}(t)}\right|_{\mathbf{u}=\mathbf{v}=\mathbf{u}^*} &= \frac{b x^*_{\mathrm{w}}(t)}{2} \left(R_{\mathrm{q}} \lambda^{\mathrm{q}}_{\mathrm{q}}(t) - \lambda^{\mathrm{q}}_{\mathrm{m}}(t)\right) \qquad (\mathbf{u}_{\mathrm{f}}, \mathbf{v}_{\mathrm{f}} \text{ under queen control}),\\
\left.\frac{\partial H_{\mathrm{w,dir}}(t)}{\partial u_{\mathrm{f}}(t)}\right|_{\mathbf{u}=\mathbf{v}=\mathbf{u}^*} &= \frac{b x^*_{\mathrm{w}}(t)}{4} \left(R_{\mathrm{w}} \lambda^{\mathrm{q}}_{\mathrm{q}}(t) - \lambda^{\mathrm{q}}_{\mathrm{m}}(t)\right) \quad (\mathbf{u}_{\mathrm{f}}, \mathbf{v}_{\mathrm{f}} \text{ under worker control}).
\end{aligned} \tag{S186}$$

It follows from eq. (S90) that the marginal return of producing a male depends on the scenario of dispersal of reproductive individuals.

#### 9.1 Delayed dispersal

Assuming that $\mu_{\rm q} = \mu_{\rm m} = \mu_{\rm r}$, it follows from eq. (S90) (by taking $t_1 = T$) for delayed dispersal

$$\begin{aligned} \lambda_{\rm q}^{\rm q}(t) &= \lambda_{\rm q}^{\rm q}(T) e^{-\mu_{\rm r}(T-t)} \quad \forall t \in [t_{c,1}^*, T], \\ \lambda_{\rm m}^{\rm q}(t) &= \lambda_{\rm m}^{\rm q}(T) e^{-\mu_{\rm r}(T-t)} \quad \forall t \in [t_{c,1}^*, T]. \end{aligned} \tag{S187}$$

Substituting the transversality conditions (S86) and using eq. (S50), we obtain

$$\frac{\lambda_{\rm m}^{\rm q}(t)}{\lambda_{\rm q}^{\rm q}(t)} = R_c \quad \forall t \in [t_{c,1}^*, T]. \tag{S188}$$

Hence the ratio of the marginal return of a male to the marginal return of a queen is equal to the relatedness asymmetry $R_c$ at any time during the reproductive phase.

For consistency, we can substitute eq. (S188) into eq. (S186), which yields that $\partial H_{c,d}(t)/\partial u_{\rm f}(t) = 0$ throughout the reproductive phase ($\forall t \in [t_{c,1}^*, T]$).

#### 9.2 Direct dispersal

It follows from eq. (S90), where $t_1 = T$ and $\lambda_{\rm m}^{\rm q}(T) = 0$ (from eq. S87) for direct dispersal and re-arranging

$$\begin{aligned} \lambda_{\rm q}^{\rm q}(t) &= \lambda_{\rm q}^{\rm q}(T) e^{-\mu_{\rm q}(T-t)}, \\ \lambda_{\rm m}^{\rm q}(t) &= \lambda_{\rm iq}^{\rm q}(T) \frac{M F_c(t)}{M}, \end{aligned} \tag{S189}$$

where

$$F_c(t) = \int_t^T e^{-\mu_{\rm m}(\tau - t)} \frac{b a_{\rm q}^*(\tau) x_{\rm w}^*(\tau)}{x_{\rm m}^*(\tau)} e^{-\mu_{\rm q}(T-\tau)} d\tau. \tag{S190}$$

is the expected number of queens surviving until $T$ that are inseminated by a male born at $t$ (in a population, where the queens mates only once, i.e. $M = 1$). Note that, in a population where females mate $M$ times, the expected number of surviving queens inseminated by a male born at $t$ is equal to $MF_c(t)$. However, since the males are only expected to father $1/M$ of the offspring of the queen, that he would have fathered if the the queen would be singly mated, then the effect of multiple matings $M$ by the queen cancels out. Lets denote by $l(t)$ the probability that a queen produced at time $t$ survives until the end of the season, which is given by

$$l(t) = e^{-\mu_{\rm q}(\tau - t)}. \tag{S191}$$

Under direct dispersal, $\partial H_{c,\rm dir}(t)/\partial u_{\rm f}(t) < 0$ during $t \in [t_{c,1}^*, t_{c,2}^*]$ while only males are produced and $\partial H_{c,\rm dir}(t)/\partial u_{\rm f}(t) > 0$ during $t \in [t_{c,2}^*, T]$ while only queens are produced. Hence, at time $t_{c,2}^*$ the marginal value $\partial H_{c,\rm dir}(t_{c,2}^*)/\partial u_{\rm f}(t_{c,2}^*) = 0$. Assuming that the mortality of queens and males is equal ($\mu_{\rm q} = \mu_{\rm m} = \mu_{\rm r}$), it follows then from eqs. (S186) and (S191) (S189)

$$\frac{F_c(t_{c,2}^*)}{l_\mathrm{q}(t_{c,2}^*)} = \begin{cases} R_\mathrm{q} = 1 & \text{(queen control \& mixed control)}, \\ R_\mathrm{w} = \dfrac{2+M}{M} & \text{(worker control)}. \end{cases} \tag{S192}$$

It follows from eq. (S192) that the switch from male production to queen production happens when producing a male instead of a surviving queen yields $R_c$ ($R_\mathrm{q} = 1$ and $R_\mathrm{w} = (2+M)/M$) surviving inseminated queens.

#### 9.3 Verifying the consistency of eq. (S192)

Here, we show that the $F_c(t_{c,2}^*)/l_\mathrm{q}(t_{c,2}^*)$ indeed satisfies eq. (S192) given the the explicit solutions for $\mathbf{u}^*$, $\mathbf{x}^*$, $t_{c,1}^*$, and $t_{c,2}^*$ that we have already established. For this we need to evaluate $F(t)$ at time $t_{c,2}^*$ (assuming that the mortality of queens and males is equal, i.e. $\mu_\mathrm{q} = \mu_\mathrm{m} = \mu_\mathrm{r}$) and the expression of $x_\mathrm{w}^*(t)$ and $x_\mathrm{m}^*(t)$ for the last phase $t \in [t_{c,2}^*, T]$.

In order to establish the initial conditions $x_\mathrm{m}^*(t_{c,1}^*)$ and $x_\mathrm{w}^*(t_{c,1}^*)$ for the last phase, we use eq. (S88) for the penultimate phase ($t \in [t_{c,1}^*, t_{c,2}^*]$) of the uninvadable state ($\bar{a}_\mathrm{q} = 0$, $\bar{a}_\mathrm{m} = 1$, $\bar{a}_\mathrm{w} = 0$, $t_0 = t_{c,1}^*$, $t_1 = t_{c,2}^*$, $x_\mathrm{m}^*(t_{c,1}^*) = 0$) and assume that $\mu_\mathrm{q} = \mu_\mathrm{m} = \mu_\mathrm{r}$. This allows us to express $x_\mathrm{m}^*(t_{c,1}^*)$ and $x_\mathrm{w}^*(t_{c,1}^*)$ as

$$\begin{aligned} x_\mathrm{w}^*(t_{c,2}^*) &= x_\mathrm{w}^*(t_{c,1}^*) e^{-\mu_\mathrm{w}(t_{c,2}^* - t_{c,1}^*)}, \\ x_\mathrm{m}^*(t_{c,2}^*) &= \frac{b x_\mathrm{w}^*(t_{c,1}^*) e^{-\mu_\mathrm{r}(t_{c,2}^* - t_{c,1}^*)} \left( e^{(\mu_\mathrm{r} - \mu_\mathrm{w})(t_{c,2}^* - t_{c,1}^*)} - 1 \right)}{\mu_\mathrm{r} - \mu_\mathrm{w}}. \end{aligned} \tag{S193}$$

Next using eq. (S88) for the last phase $\tau = t \in [t_{c,2}^*, T]$ ($\bar{a}_\mathrm{q} = 1$, $\bar{a}_\mathrm{m} = 0$, $\bar{a}_\mathrm{w} = 0$, $t_0 = t_{c,2}^*$, $t_1 = T$), and taking eq. (S193) as an initial condition we obtain

$$\begin{aligned} x_\mathrm{w}^*(\tau) &= x_\mathrm{w}^*(t_{c,1}^*) e^{-\mu_\mathrm{w}(\tau - t_{c,1}^*)}, \\ x_\mathrm{m}^*(\tau) &= \frac{b x_\mathrm{w}^*(t_{c,1}^*) e^{-\mu_\mathrm{r}(\tau - t_{c,1}^*)} \left( e^{(\mu_\mathrm{r} - \mu_\mathrm{w})(t_{c,2}^* - t_{c,1}^*)} - 1 \right)}{\mu_\mathrm{r} - \mu_\mathrm{w}}. \end{aligned} \tag{S194}$$

Now we can evaluate $F(t)$ at time $t_{c,2}^*$ by substituting eq. (S194) into eq. (S190) (assuming $\mu_\mathrm{q} = \mu_\mathrm{m} = \mu_\mathrm{r}$) and taking into account that $a_\mathrm{q}^*(t) = 1$ during the last phase $t \in [t_{c,2}^*, T]$ yields

$$\begin{aligned} F_c(t_{c,2}^*) &= e^{-\mu_\mathrm{r}(T - t_{c,2}^*)} \int_{t_{c,2}^*}^{T} \frac{b x_\mathrm{w}^*(\tau)}{x_\mathrm{m}^*(\tau)} d\tau = e^{-\mu_\mathrm{r}(T - t_{c,2}^*)} \int_{t_{c,2}^*}^{T} \frac{(\mu_\mathrm{r} - \mu_\mathrm{w}) e^{(\mu_\mathrm{r} - \mu_\mathrm{w})(\tau - t_{c,1}^*)}}{e^{(\mu_\mathrm{r} - \mu_\mathrm{w})(t_{c,2}^* - t_{c,1}^*)} - 1} d\tau \\ &= e^{-\mu_\mathrm{r}(T - t_{c,2}^*)} \frac{e^{(\mu_\mathrm{r} - \mu_\mathrm{w}) t_{c,2}^*} - e^{(\mu_\mathrm{r} - \mu_\mathrm{w}) T}}{e^{(\mu_\mathrm{r} - \mu_\mathrm{w}) t_{c,1}^*} - e^{(\mu_\mathrm{r} - \mu_\mathrm{w}) t_{c,2}^*}}. \end{aligned} \tag{S195}$$

Substituting the expressions for the switching times under direct dispersal (eqs. S149, S150, S152, and S154), and simplifying, we get

$$F_c(t^*_{c,2})l_{\mathrm{q}}(t^*_{c,2}) = \begin{cases} e^{-\mu_{\mathrm{r}}(T-t^*_{c,2})}R_{\mathrm{q}} = l(t^*_{c,2})R_{\mathrm{q}} & \text{(queen control \& mixed control)}, \\ e^{-\mu_{\mathrm{r}}(T-t^*_{c,2})}R_{\mathrm{w}} = l(t^*_{c,2})R_{\mathrm{w}} & \text{(worker control)}. \end{cases} \tag{S196}$$

##### 9.4 The overall sex allocation ratio and the ratio of expected surviving inseminated queens to surviving queens

It is significant that the necessary condition for $\mathbf{u}^*$ to be uninvadable under direct dispersal, is given by the condition that the ratio $F_c(t)/l(t)$ at time $t^*_{c,2}$ would be equal to the relatedness asymmetry and not on the condition on the overall sex allocation ratio $S_c$. In this section we would like to make a connection between these two terms.

First, lets define the overall sex allocation ratio $Z_c = S_c/(1-S_c)$ (recall S172 for the formal definition of $S_c$) as the ratio of the proportion of colony resources allocated queens versus males throughout the entire season (which would make it easier to compare overall allocation to queens versus males to the ratio $F_c(t^*_{c,2})/l(t^*_{c,2})$). Hence, we can write $Z_c$ as

$$Z_c = \frac{\int_0^T b a^*_{\mathrm{q}}(t) x^*_{\mathrm{w}}(t)\,\mathrm{d}t}{\int_0^T b a^*_{\mathrm{m}}(t) x^*_{\mathrm{w}}(t)\,\mathrm{d}t}. \tag{S197}$$

Substituting the uninvadable resource allocation schedule $\mathbf{u}^*$ for $\mu_{\mathrm{m}} = \mu_{\mathrm{q}} = \mu_{\mathrm{r}}$ under direct dispersal given by eq. (S148) with the solutions to state equations given by (S89) into eq. (S197) yields

$$\begin{aligned} Z_c =& \frac{\int_{t^*_{c,2}}^T b x^*_{\mathrm{w}}(t)\,\mathrm{d}t}{\int_{t^*_{c,1}}^{t^*_{c,2}} b x^*_{\mathrm{w}}(t)\,\mathrm{d}t} = \frac{x^*_{\mathrm{w}}(t^*_{c,2})\int_{t^*_{c,2}}^T e^{-\mu_{\mathrm{w}}(t-t^*_{c,2})}\,\mathrm{d}t}{x^*_{\mathrm{w}}(t^*_{c,1})\int_{t^*_{c,1}}^{t^*_{c,2}} e^{-\mu_{\mathrm{w}}(t-t^*_{c,1})}\,\mathrm{d}t} \\ =& \frac{x^*_{\mathrm{w}}(t^*_{c,2})e^{\mu_{\mathrm{w}}t^*_{c,2}}\int_{t^*_{c,2}}^T e^{-\mu_{\mathrm{w}}t}\,\mathrm{d}t}{x^*_{\mathrm{w}}(t^*_{c,1})e^{\mu_{\mathrm{w}}t^*_{c,1}}\int_{t^*_{c,1}}^{t^*_{c,2}} e^{-\mu_{\mathrm{w}}t}\,\mathrm{d}t} = \frac{x^*_{\mathrm{w}}(t^*_{c,1})e^{-\mu_{\mathrm{w}}(t^*_{c,2}-t^*_{c,1})}e^{\mu_{\mathrm{w}}t^*_{c,2}}(-1/\mu_{\mathrm{w}})[e^{-\mu_{\mathrm{w}}T} - e^{-\mu_{\mathrm{w}}t^*_{c,2}}]}{x^*_{\mathrm{w}}(t^*_{c,1})e^{\mu_{\mathrm{w}}t^*_{c,1}}(-1/\mu_{\mathrm{w}})[e^{-\mu_{\mathrm{w}}t^*_{c,2}} - e^{-\mu_{\mathrm{w}}t^*_{c,1}}]} \\ =& \frac{e^{-\mu_{\mathrm{w}}t^*_{c,2}} - e^{-\mu_{\mathrm{w}}T}}{e^{-\mu_{\mathrm{w}}t^*_{c,1}} - e^{-\mu_{\mathrm{w}}t^*_{c,2}}}. \end{aligned} \tag{S198}$$

The overall allocation to queens versus males under direct dispersal is therefore given by

$$Z_c = \frac{e^{-\mu_{\mathrm{w}}t^*_{c,2}} - e^{-\mu_{\mathrm{w}}T}}{e^{-\mu_{\mathrm{w}}t^*_{c,1}} - e^{-\mu_{\mathrm{w}}t^*_{c,2}}}. \tag{S199}$$

It follows from eq. (S195) that the ratio $F_c(t^*_{c,2})/l(t^*_{c,2})$ can be expressed as

$$\frac{F_c(t^*_{c,2})}{l(t^*_{c,2})} = \frac{e^{(\mu_{\mathrm{r}}-\mu_{\mathrm{w}})t^*_{c,2}} - e^{(\mu_{\mathrm{r}}-\mu_{\mathrm{w}})T}}{e^{(\mu_{\mathrm{r}}-\mu_{\mathrm{w}})t^*_{c,1}} - e^{(\mu_{\mathrm{r}}-\mu_{\mathrm{w}})t^*_{c,2}}}. \tag{S200}$$

Hence, one can see from eqs. (S199) and (S200) that

$$Z_c < \frac{F_c(t^*_{c,2})}{l(t^*_{c,2})} \quad \forall c \in \{\mathrm{q}, \mathrm{w}, \mathrm{mx}\} \tag{S201}$$

whenever $\mu_\mathrm{r} > 0$ and the difference $F_c(t^*_{c,2})/l(t^*_{c,2}) - Z_c$ is larger for higher values of $\mu_\mathrm{r}$. Hence, the overall sex allocation ratio is more male-biased under direct dispersal than expected from the classical results (e.g. Reuter and Keller, 2001) for higher values of mortality of reproductive individuals.

In order to get a better intuition why the overall sex allocation ratio is more male-biased for higher values of mortality $\mu_\mathrm{r}$ of reproductive individuals, lets examine how $\mu_\mathrm{r}$ influences $F_c(t)/l(t)$ during the reproductive phase which we can express as (using eq. S190 and S191)

$$\frac{F_c(t)}{l(t)} = \int_t^T \frac{ba^*_\mathrm{q}(\tau)x^*_\mathrm{w}(\tau)}{x^*_\mathrm{m}(\tau)} d\tau. \tag{S202}$$

Here, $ba^*_\mathrm{q}(t)x^*_\mathrm{w}(\tau)/x^*_\mathrm{m}(\tau)$ gives the mating success of a male (the number of queens available to mate per male at time $\tau$). Here, $ba^*_\mathrm{q}(t)x^*_\mathrm{w}(\tau)$ is independent of the mortality of sexuals $\mu_\mathrm{r}$, since ($a^*_\mathrm{q}(\tau) = 0$, during male production and $a^*_\mathrm{q}(\tau) = 1$ during queen production) and number of workers $x^*_\mathrm{w}(\tau)$ does not depend on the mortality of sexuals. In contrast, the number of males $x^*_\mathrm{m}(\tau)$ alive at time $\tau$ during the last phase, when only females are produced is smaller for higher values of $\mu_\mathrm{r}$. Hence, it follows that $F_c(t)/l(t)$ is higher for higher mortality rate of sexuals (for a given $t$ in the reproductive phase), because higher mortality increases the mating success of a male alive at a given time $t$. Since, the ratio $F_c(t)/l(t)$ gives the expected number of surviving queens inseminated by a male produced instead of a queen, then the surviving probability of a focal male together with the surviving probability of the queen(s) he inseminates cancels out with the surviving probability of a queen that would have been otherwise produced (since we assumed that the mortality of queens and males is equal). Because of this, the ratio $F_c(t)/l(t)$ increases with the increase in the mortality of sexuals via the mating success of a focal male $1/x^*_\mathrm{m}(\tau)$.

### 10 Continuous stability of the candidate uninvadable allocation schedule

In this section we address the issue of (continuous) stability of the candidate uninvadable allocation schedule $\mathbf{u}^*$ given by eq. (S92) for delayed dispersal and eq. (S148) for direct dispersal. We only discuss the continuous stability of the candidate uninvadable allocation schedule $\mathbf{u}^*$ for equal mortality rate of queens and males ($\mu_\mathrm{q} = \mu_\mathrm{m} = \mu_\mathrm{r}$) because we have fully derived the analytical results only under this assumption. Continuous stability is given by two separate properties of the of the candidate uninvadable allocation schedule $\mathbf{u}^*$: (i) the local uninvadablity and (ii) convergence stability (e.g. see Christiansen, 1991; Eshel, 1983; Taylor, 1989 and for functioned-valued traits see Dieckmann et al., 2006). The candidate uninvadable allocation schedule $\mathbf{u}^*$ is locally uninvadable if a monomorphic population following the strategy $\mathbf{u}^*$ can resist invasion by any mutant whose strategy is close to the the strategy $\mathbf{u}^*$. The candidate uninvadable allocation schedule $\mathbf{u}^*$ is convergence stable if a population will converge to this schedule $\mathbf{u}^*$ through recurrent substitutions, meaning that a mutant whose schedule is closer to $\mathbf{u}^*$ will invade a monomorphic population that follows a schedule further away from $\mathbf{u}^*$.

Firstly, we would like to point out that the continuous stability of the candidate uninvadable allocation schedule $\mathbf{u}^*$ (assuming that $\mu_{\rm q} = \mu_{\rm m} = \mu_{\rm r}$) is not directly given from the first-order condition only for $t \in\in \left[t^*_{c,1}, T\right]$ under delayed dispersal. Indeed, if $\partial H_{c,d}(\mathbf{u}(t), \mathbf{x}^*(t), \boldsymbol{\lambda}(t))/\partial u_\tau(t)|_{\mathbf{u}=\mathbf{u}^*} > 0$ then a mutant allele with $u_\tau(t) > v_\tau(t)$ can always spread (recall eq. S73). Similarly, if $\partial H_{c,d}(\mathbf{u}(t), \mathbf{x}^*(t), \boldsymbol{\lambda}(t))/\partial u_\tau(t)|_{\mathbf{u}=\mathbf{u}^*} < 0$ then a mutant allele with $u_\tau(t) < v_\tau(t)$ can always spread. However, for a finite period of time, for which $\partial H_{c,d}(\mathbf{u}(t), \mathbf{x}^*(t), \boldsymbol{\lambda}(t))/\partial u_\tau(t)|_{\mathbf{u}=\mathbf{u}^*} = 0$ holds and the focal trait is a singular arc $u^*_\tau(t)$, then there is no directional selection. Hence, the two properties of continuous stability of $\mathbf{u}^*$ has to be only addressed for $t \in \left[t^*_{c,1}, T\right]$ under delayed dispersal, where $u^*_{\rm f}(t)$ is a singular arc.

The conditions for local uninvadability and convergence stability are given in terms of second-order functional derivatives of the invasion fitness (S10) (Dieckmann et al., 2006). Since the formal continuous stability analysis for function-valued traits is out of the scope of this paper, we will not address the questions of continuous stability of $\mathbf{u}^*$ for $t \in \left[t^*_{c,1}, T\right]$ under delayed dispersal any further in this paper. However, since we recover the uninvadable allocation schedule numerically through iteration, which is analogous to the evolutionary dynamics of the population, then we might regard this as giving support that the proper evolutionary dynamics would also converge to this schedule. Under this heuristic approach, the iterative scheme of the best response map also implies continuous stability of the candidate uninvadable schedule found by it (Houston and McNamara, 1999).

### 11 Iterative scheme of the best response map

Here, we describe the computational technique for finding the locally uninvadable strategies for our optimal control problems. This method is known as the iterative scheme of the best response map (see p. 187 in Houston and McNamara, 1999).

A mutant schedule $\left(\beta^{\rm f}_{c,d}(\mathbf{v}), \beta^{\rm q}_{c,d}(\mathbf{v})\right)$ that yields the highest invasion fitness in a population, where resident schedule is $\mathbf{v}$, i.e.

$$\beta^{\rm f}_{c,d}(\mathbf{v}) = \arg\max_{\mathbf{u}_{\rm f}\in\mathbb{U}_{\rm f}} W_{c,d}\left[(\mathbf{u}_{\rm f}, \mathbf{v}_{\rm q}), \mathbf{v}\right] \quad \text{and} \quad \beta^{\rm q}_{c,d}(\mathbf{v}) = \arg\max_{\mathbf{u}_{\rm q}\in\mathbb{U}_{\rm q}} W_{c,d}\left[(\mathbf{v}_{\rm f}, \mathbf{u}_{\rm q}), \mathbf{v}\right], \tag{S203}$$

is said be the best response to the resident schedule $\mathbf{v}$. Here, $\beta^\tau_{c,d} : \mathbb{U} \to \mathbb{U}_\tau$, where $\tau \in {\rm f}, {\rm q}$ is the best-response correspondence which maps a resident schedule $\mathbf{v} \in \mathbb{U}$ to a (unique) trajectory $\beta^\tau_{c,d}(\mathbf{v}) \in \mathbb{U}_\tau$ for a trait type $\tau$, such that no other trajectory for a focal trait gives a higher invasion fitness to a mutant in a population, where resident individuals follow the schedule $\mathbf{v} \in \mathbb{U}$. Here, $\mathbb{U} = \mathbb{U}_{\rm f} \times \mathbb{U}_{\rm q}$ is a set of all possible allocation strategies, $\mathbb{U}_{\rm f}$ and $\mathbb{U}_{\rm q}$ are sets of all possible trajectories for the traits $\mathbf{u}_{\rm f}$ ($\mathbf{v}_{\rm f}$) and $\mathbf{u}_{\rm q}$ ($\mathbf{v}_{\rm q}$), respectively. In the notation of the best-response correspondence $\beta^\tau_{c,d}$, the subscripts $c \in \{{\rm q}, {\rm w}\}$ and $d \in \{{\rm del}, {\rm dir}\}$ emphasize the party in control and the time of dispersal of sexuals, respectively, and superscript $\tau \in \{{\rm f}, {\rm f}\}$ emphasizes the trait type.

Hence, under single-party control, where party $c \in \{{\rm q}, {\rm w}\}$ is in full control, the best response schedule

$\beta_{c,d}(\mathbf{v}) = \left(\beta^{\mathrm{f}}_{c,d}(\mathbf{v}), \beta^{\mathrm{q}}_{c,d}(\mathbf{v})\right)$ can be written as

$$\beta_{c,d}(\mathbf{v}) = \arg\max_{\mathbf{u}\in\mathbb{U}} W_{c,d}\left(\mathbf{u},\mathbf{v}\right) \tag{S204}$$

and under mixed control the best response schedule $\beta_{\mathrm{mx},d}(\mathbf{v}) = \left(\beta^{\mathrm{f}}_{\mathrm{mx},d}(\mathbf{v}), \beta^{\mathrm{q}}_{\mathrm{mx},d}(\mathbf{v})\right)$ can be written as

$$\beta^{\mathrm{f}}_{\mathrm{mx},d}(\mathbf{v}) = \arg\max_{\mathbf{u}_\mathrm{f}\in\mathbb{U}_\mathrm{f}} W_{\mathrm{q},d}\left[(\mathbf{u}_\mathrm{f},\mathbf{v}_\mathrm{q}),\mathbf{v}\right] \quad \text{and} \quad \beta^{\mathrm{q}}_{\mathrm{mx},d}(\mathbf{v}) = \arg\max_{\mathbf{u}_\mathrm{q}\in\mathbb{U}_\mathrm{q}} W_{\mathrm{w},d}\left[(\mathbf{v}_\mathrm{f},\mathbf{u}_\mathrm{q}),\mathbf{v}\right], \tag{S205}$$

where $\beta_c : \mathbb{U} \to \mathbb{U}$ under control mode $c \in \{\mathrm{q},\mathrm{w},\mathrm{mx}\}$ is the best-response correspondence which maps to a (resident) schedule $\mathbf{v} \in \mathbb{U}$ a schedule $\beta_c(\mathbf{v}) \in \mathbb{U}$, such that no other schedule gives a higher invasion fitness to a mutant in a population, where resident individuals follow the schedule $\mathbf{v} \in \mathbb{U}$.

It follows from the definition of the uninvadable schedule given by eq. (S12) for single-party control and by eq. (S13) for mixed control that the uninvadable schedule is a best response to itself, i.e.

$$\mathbf{u}^* = \beta_{c,d}(\mathbf{u}^*). \tag{S206}$$

Note that, here we have assumed that the best response is always unique.

In order to approximate the uninvadable schedule numerically, we start out from some initial resource allocation schedule for the resident population $\mathbf{u}^0$ and using GPOPS (Patterson and Rao, 2014) we find the mutant schedule that has the highest fitness $\mathbf{u}^1 = \beta_{c,d}(\mathbf{u}^0)$. The software GPOPS uses a direct approach to find the best response $\beta_{c,d}(\mathbf{v})$ for a given environment $\mathbf{v}$ in contrast to the indirect approach of Pontryagin's maximum principle (see section 3.2), which gives a necessary condition for optimality. We then update the resident schedule for the next iteration

$$\mathbf{u}^i = \delta\beta_{c,d}(\mathbf{u}^{i-1}) + (1-\delta)\mathbf{u}^{i-1} \tag{S207}$$

and repeat the process. Here, $0 > \delta > 1$ is called the replacement factor. We can interpret this new resident schedule as a polymorphism - each individual adopting a schedule $\beta_{c,d}(\mathbf{u}^{i-1})$ with probability $\alpha$ and schedule $\mathbf{u}^{i-1}$ with probability $(1-\delta)$ (Houston and McNamara, 1999). To improve convergence after iterating from some while we can decrease $\delta$ with further iterations (Houston and McNamara, 1999; Krawczyk and Uryasev, 2000).

This iterative scheme forms a sequence of strategies $(\mathbf{u}^0; \mathbf{u}^1; \mathbf{u}^2; ...)$ where each schedule is derived from the best response to the previous schedule according to equation (S207). If the difference between the best response and resident schedule approaches zero as the number of iterations increases, i.e.

$$\left|\mathbf{u}^i - \mathbf{u}^{i-1}\right| \to 0 \text{ as } i \to \infty, \tag{S208}$$

then we have arrived at the uninvadable schedule (Nash equilibrium).

For single-party control we use GPOPS to find the best response $\beta_{c,d}(\mathbf{v})$ that maximizes the objective $W_{c,d}(\mathbf{u},\mathbf{v})$ given by eq. (S10) of the party $c$ in control. For mixed control, the best response $\beta^{\mathrm{f}}_{c,d}(\mathbf{v})$ maximizes the the objective of the queen $W_{\mathrm{q},d}(\mathbf{u},\mathbf{v})$ and $\beta^{\mathrm{q}}_{c,d}(\mathbf{v})$ maximizes the objective of the workers $W_{\mathrm{w},d}(\mathbf{u},\mathbf{v})$.

### 12 Static resource allocation model with a linear relationship between colony productivity and colony size

In this section we re-derive the main results of Reuter and Keller (2001) with a slight modification in the assumption about how the overall colony productivity scales with colony size. Here we assume that the overall colony productivity scales linearly with colony size and show that the main predictions of their model about how sex allocation conflict affect allocation strategy in the colony are not affected by the assumption about how overall colony productivity scales with colony size. In order to make a direct comparison easier for the reader, we slightly modified the notation in Reuter and Keller (2001) to make it easier to compare our model with theirs, however the modelling approach in this section is exactly identical to Reuter and Keller (2001). Here we present the main features of the model assuming monogamy, for the full model and further details, see the original paper by Reuter and Keller (2001).

Let $0 \leq u_{\mathrm{f}} \leq 1$ ($u_{\mathrm{f}} = f$ in their notation) be the proportion of colony resources allocated into producing females in a focal colony and let $0 \leq u_{\mathrm{q}} \leq 1$ ($u_{\mathrm{q}} = 1 - w$ in their notation) be the proportion of resources allocated into queens from the resources allocated into females. Let the the corresponding population average traits be $0 \leq v_{\mathrm{f}} \leq 1$ and $0 \leq v_{\mathrm{q}} \leq 1$ ($F$ and $1-W$ in their notation), respectively. Here the allocation strategies $u_{\mathrm{f}}$, $u_{\mathrm{q}}$ ($v_{\mathrm{f}}$, $v_{\mathrm{q}}$) give the allocation of all colony resources over the entire season.

Since this model is a static resource allocation model, colony size is given by the proportional investment $u_{\mathrm{f}}(1-u_{\mathrm{q}})$ into workers. Reuter and Keller (2001) assume that per-worker productivity declines with the number of workers in the colony, such that overall colony productivity $b(u_{\mathrm{f}}, u_{\mathrm{q}})$ (interpreted in their paper as the total biomass of all individuals produced) in the focal colony follows a diminishing return function

$$b(u_{\mathrm{f}}, u_{\mathrm{q}}) = 1 - (1 - u_{\mathrm{f}}(1 - u_{\mathrm{q}}))^2. \tag{S209}$$

Next, they formulate an expression for the fitness function ($V_X$ in their notation)

$$W_c = b(u_{\mathrm{f}}, u_{\mathrm{q}}) \left[ r^{\circ}_{\mathrm{q},c} \alpha^{\circ}_{\mathrm{q}} \frac{u_{\mathrm{f}} u_{\mathrm{q}}}{v_{\mathrm{f}} v_{\mathrm{q}}} + r^{\circ}_{\mathrm{m},c} \alpha^{\circ}_{\mathrm{q}} \frac{(1 - u_{\mathrm{f}})}{(1 - v_{\mathrm{f}})} \right], \tag{S210}$$

where the subscript $c \in \{\mathrm{q}, \mathrm{w}\}$ emphasizes the party in control. See eq. 2 in Reuter and Keller, 2001 for further details about the interpretation of this fitness function. Essentially, $W_c$ in (S210) is comparable to the invasion fitness $W_c$ in (S9). However, we note that this way of defining an invasion fitness function is heuristic and implicitly makes the assumption that only a first-order analysis will be carried out (e.g. otherwise relatedness coefficients cannot be hold at neutrality), and so the eq. (S210) can not be used to check second order conditions (further it should be normalized so that in a resident population fitness is one). Candidate uninvadable allocation strategy $u^*_{\mathrm{f}}$, $u^*_{\mathrm{q}}$ (called evolutionary stable strategy and noted as $f^*$, $w^*$ in Reuter and Keller, 2001) under single-party $c$ control is given by

$$\frac{\mathrm{d}W_c}{\mathrm{d}u_\mathrm{f}} = 0 \quad \text{and} \quad \frac{\mathrm{d}W_c}{\mathrm{d}u_\mathrm{q}} = 0 \tag{S211}$$

and under mixed control is given by

$$\frac{\mathrm{d}W_\mathrm{q}}{\mathrm{d}u_\mathrm{f}} = 0 \quad \text{and} \quad \frac{\mathrm{d}W_\mathrm{w}}{\mathrm{d}u_\mathrm{q}} = 0. \tag{S212}$$

The results of Reuter and Keller (2001) assuming monogamy are outlined in table S2. The main results of Reuter and Keller (2001) can be summarized as follows: less resources are allocated into worker production under single-party control than under mixed control, the uninvadable sex allocation ratio is equal to the relatedness asymmetry under single-party control ($R_\mathrm{q} = 1$ for queen control and $R_\mathrm{w} = 3$ under worker control, assuming monogamy). Under mixed control the uninvadable sex allocation ratio has a value intermediate $\approx 1.26$ between the relatedness asymmetries for queen and worker control.

| Control mode | Queens ($u_\mathrm{f}^* u_\mathrm{q}^*$) | Males ($1 - u_\mathrm{f}^*$) | Workers ($u_\mathrm{f}^*(1 - u_\mathrm{q}^*)$) | $S_c$ |
|---|---|---|---|---|
| Queen control | $\approx 0.289$ | $\approx 0.289$ | $\approx 0.423$ | 0.5 |
| Worker control | $\approx 0.433$ | $\approx 0.144$ | $\approx 0.423$ | 0.75 |
| Mixed control | $\approx 0.353$ | $\approx 0.281$ | $\approx 0.365$ | $\approx 0.56$ |

Table S2: Uninvadable allocation into queen, males, and workers and the overall sex allocation ratio $S_c$ (proportional allocation to queens from resources allocated to sexuals) predicted by Reuter and Keller (2001).

Lets now modify the overall colony productivity $b(u_\mathrm{f}, u_\mathrm{q})$, such that it increases linearly with the amount of resources invested into workers $u_\mathrm{f}(1 - u_\mathrm{q})$.

$$b_{\mathrm{lin}}(u_\mathrm{f}, u_\mathrm{q}) = u_\mathrm{f}(1 - u_\mathrm{q}). \tag{S213}$$

Substituting $b_{\mathrm{lin}}(u_\mathrm{f}, u_\mathrm{q})$ into the fitness function (S210) instead of $b(u_\mathrm{f}, u_\mathrm{q})$ and solving for the uninvadable allocation strategy according to eqs. (S211)–(S212), we obtain the uninvadable allocation strategy $(u_\mathrm{f}^*, u_\mathrm{q}^*)$ assuming that colony productivity scales linearly with colony size. The results are outlined in table S3. The main results of the static model, where colony productivity scaled linearly with colony size can be summarized as follows: less resources are allocated into worker production under single-party control than under mixed control, the uninvadable sex allocation ratio is equal to the relatedness asymmetry under single-party control ($R_\mathrm{q} = 1$ for queen control and $R_\mathrm{w} = 3$ under worker control, assuming monogamy). Under mixed control the uninvadable sex allocation ratio has a value intermediate $\approx 1.28$ between the relatedness asymmetries for queen and worker control.

Hence, we can conclude, that the main results predicted by a static allocation model of Reuter and Keller (2001) about how sex allocation conflict affects allocation strategies in the colony are not qualitatively affected by the assumption about how colony productivity scales with colony size.

| Control mode | Queens ($u_{\mathrm{f}}^{*}u_{\mathrm{q}}^{*}$) | Males ($1-u_{\mathrm{f}}^{*}$) | Workers ($u_{\mathrm{f}}^{*}(1-u_{\mathrm{q}}^{*})$) | $S_c$ |
|---|---|---|---|---|
| Queen control | 0.25 | 0.25 | 0.5 | 0.5 |
| Worker control | 0.375 | 0.125 | 0.5 | 0.75 |
| Mixed control | $\approx 0.321$ | $\approx 0.25$ | $\approx 0.429$ | $\approx 0.56$ |

Table S3: Uninvadable allocation into queen, males, and workers and the overall sex allocation ratio $S_c$ (proportional allocation to queens from resources allocated to sexuals) predicted by a static model similar to Reuter and Keller (2001), assuming that colony productivity scales linearly with colony size.

#### 14 Summary of notation

| Symbol | Meaning |
|---|---|
| $a_k(t)$ | Proportion of resources allocated to producing type $k \in \{\mathrm{w}, \mathrm{q}, \mathrm{m}\}$ individuals at time $t$ in a colony founded by resident individuals. |
| $a^s_{k,\mathrm{u}}(t)$ | Proportion of resources allocated to producing type $k \in \{\mathrm{w}, \mathrm{q}, \mathrm{m}\}$ individuals at time $t$ in a colony founded by a mutant individual of type $s \in \{\mathrm{q}, \mathrm{m}\}$. |
| $\mathbf{A}_c(\mathbf{u}, \mathbf{v})$ | Matrix whose elements $a_{s's}$ give the expected number of mutant gene copies in a type $s' \in \{\mathrm{q}, \mathrm{m}\}$ individual that descend from an individual of type $s \in \{\mathrm{q}, \mathrm{m}\}$ ($\mathbf{A}_c(\mathbf{u}, \mathbf{v}) = \mathbf{A}_c(\mathbf{u}^{\mathrm{q}}(\mathbf{u}), \mathbf{u}^{\mathrm{m}}(\mathbf{u}), \mathbf{v})$ and so matrix elements depend on $u^s_\tau(t)$). |
| $b$ | Individual productivity rate of a worker. |
| $B(t^*_{c,1})$ | Colony productivity under the univadable allocation schedule $\mathbf{u}^*$, number of sexuals produced in the focal colony, that survive until the end of the season. |
| $C(M) = C$ | Potential for conflict. |
| $H_{c,d}(\mathbf{u}(t), \mathbf{x}(t), \boldsymbol{\lambda}(t))$ | Hamiltonian function if the evolving trait is under the genetic control of a party $c \in \{\mathrm{w}, \mathrm{q}\}$ for a scenario $d \in \{\mathrm{del}, \mathrm{dir}\}$ of dispersal of sexuals. |
| $M$ | Queen mating frequency. |
| $n$ | Number of colonies or breeding sites (large and constant). |
| $p^s_c$ | Expected frequency of the mutant allele in party $c$ in a colony founded by a mutant individual of type $s \in \{\mathrm{q}, \mathrm{m}\}$. |
| $q_s(\mathbf{u}, \mathbf{v})$ | Asymptotic probability that a mutant allele is sampled in an individual of type $s \in \{\mathrm{q}, \mathrm{m}\}$. |
| $R_c$ | Relatedness asymmetry, i.e, the ratio of sex-specific (females to males) contributions of genes of a party $c \in \{\mathrm{w}, \mathrm{q}\}$ (in a neutral process) to the gene pool in distant future. |
| $r^\circ_{s,c}$ | Average relatedness between an individual of party $c \in \{\mathrm{w}, \mathrm{q}\}$ and a juvenile individual of type $s \in \{\mathrm{q}, \mathrm{m}\}$. |
| $t$ | Time of the season defined over a period $[0, T]$. |
| $t^*_{c,1}$ | Switching time from the ergonomic phase to the reproductive phase under the scenario $c \in \{\mathrm{q}, \mathrm{w}, \mathrm{mx}\}$ of genetic control of resource allocation traits. |
| $t^*_{c,2}$ | Switching time from the production of only males to the production of only queens (for direct dispersal) under the scenario $c \in \{\mathrm{q}, \mathrm{w}, \mathrm{mx}\}$ of genetic control of resource allocation traits. |
| $u_\tau(t)$ | Resource allocation trait of type $\tau \in \{\mathrm{f}, \mathrm{q}\}$ of a colony where all the genes in control of the trait carry only mutant alleles. |
| $u^s_\tau(t)$ | Resource allocation trait of type $\tau \in \{\mathrm{f}, \mathrm{q}\}$ of a colony founded by a mutant individual of type $s \in \{\mathrm{q}, \mathrm{m}\}$. |
| $u^*_\tau(t)$ | Uninvadable resource allocation trait of type $\tau \in \{\mathrm{f}, \mathrm{q}\}$. |

| | |
|---|---|
| $\mathbf{u}$ | Full resource allocation schedule of a colony founded by an individual of type $s$ who carries a mutant allele for each of the evolving traits, i.e. $\mathbf{u} = \{u_{\mathrm{f}}(t), u_{\mathrm{q}}(t)\}_{t\in[0,T]}$. Note that $\mathbf{u}(t) = (u_{\mathrm{f}}(t), u_{\mathrm{q}}(t))$. |
| $\mathbf{u}^s$ | A hypothetical resource allocation schedule of a colony founded by an individual of type $s \in \{\mathrm{q}, \mathrm{m}\}$ who is homozygous for the mutant alleles for both of the evolving traits, i.e. $\mathbf{u}^s = \{u^s_{\mathrm{f}}(t), u^s_{\mathrm{q}}(t)\}_{t\in[0,T]}$. Note that $\mathbf{u}^s(t) = (u^s_{\mathrm{f}}(t), u^s_{\mathrm{q}}(t))$. |
| $\mathbf{u}^*$ | Full uninvadable allocation schedule defined over the entire season $t \in [0, T]$, $\mathbf{u}^* = \{u^*_{\mathrm{f}}(t), u^*_{\mathrm{q}}(t)\}_{t\in[0,T]}$. Note that $\mathbf{u}^*(t) = (u^*_{\mathrm{f}}(t), u^*_{\mathrm{q}}(t))$. Note that in the main text we use a different notation, whereby $\mathbf{v}^* \equiv \mathbf{u}^*$. |
| $v_\tau(t)$ | Resource allocation trait of type $\tau \in \{\mathrm{f}, \mathrm{q}\}$ of a colony founded by resident individuals. |
| $\mathbf{v}$ | Full resource allocation schedule that describes how resources are allocated throughout the season $t \in [0, T]$ in a colony founded by resident individuals, i.e. $\mathbf{v} = \{v_{\mathrm{f}}(t), v_{\mathrm{q}}(t)\}_{t\in[0,T]}$. Note that $\mathbf{v}(t) = (v_{\mathrm{f}}(t), v_{\mathrm{q}}(t))$. |
| $w_{s's}(\mathbf{u}^s, \mathbf{v})$ | Expected number of juveniles of type $s' \in \{\mathrm{q}, \mathrm{m}\}$ that descend from a mutant colony-founding individual of type $s \in \{\mathrm{q}, \mathrm{m}\}$ carrying the mutant allele. |
| $W_c(\mathbf{u}, \mathbf{v})$ or $W_{c,d}(\mathbf{u}, \mathbf{v})$ | Invasion fitness of the mutant allele for a trait that is under the genetic control of a party $c \in \{\mathrm{w}, \mathrm{q}\}$. The additional subscript in the latter notation further emphasizes the scenario $d \in \{\mathrm{del}, \mathrm{dir}\}$ of dispersal of sexuals. |
| $y_k(t)$ | Number of type $k \in \{\mathrm{w}, \mathrm{q}, \mathrm{m}\}$ individuals alive at time $t$, who have been produced in a colony founded by resident individuals. Note that in the main text we use a different notation, whereby $x_k \equiv y_k$. |
| $x^s_k(t)$ | Number of type $k \in \{\mathrm{w}, \mathrm{q}, \mathrm{m}\}$ individuals alive at time $t$, who have been produced in a colony founded by a mutant individual of type $s \in \{\mathrm{q}, \mathrm{m}\}$. |
| $x^s_{\mathrm{iq}}(t)$ | Number of females inseminated by males produced in a colony founded by a mutant individual of type $s \in \{\mathrm{q}, \mathrm{m}\}$. |
| $x^*_k(t)$ | Number of type $k \in \{\mathrm{w}, \mathrm{q}, \mathrm{m}\}$ individuals alive at time $t$, who have been produced in the colony following the uninvadable allocation schedule $\mathbf{u}^*$. |
| $x^*_{\mathrm{w}}(t^*_{c,1})$ | Colony size at maturity under the univadable allocation schedule $\mathbf{u}^*$, i.e. colony size at $t^*_{c,1}$ when colony switches from the ergonomic phase to the reproductive phase. |
| $\alpha^\circ_s$ | Neutral class reproductive value of an individual of type $s \in \{\mathrm{q}, \mathrm{m}\}$. |
| $\delta$ | Replacement factor in the iterative scheme of best response map. |
| $\gamma_{s's}$ | Probability that a gene sampled in an individual of type $s' \in \{\mathrm{q}, \mathrm{m}\}$ was contributed by an individual of type $s \in \{\mathrm{q}, \mathrm{m}\}$; i.e. transmission frequency of type $s$ to type $s'$. |
| $\lambda^s_k(t)$ | Costate variable associated with the state variable $x^s_{k,\mathrm{u}}(t)$. |
| $\epsilon_\tau$ | The intensity of deviation from the resident trait $v_\tau(t)$. |

| | |
|---|---|
| $\eta_\tau(t)$ | Deviation from the resident trait $v_\tau(t)$. |
| $\mu_k$ | Mortality rate of type $k \in \{\mathrm{w}, \mathrm{q}, \mathrm{m}\}$ individuals. If mortality of males and queens is equal ($\mu_\mathrm{q} = \mu_\mathrm{q}$) then we denote by $\mu_\mathrm{r}$ the mortality rate of sexuals. |
| $\tau$ (subscript) | Denotes the type of the trait. If $\tau = \mathrm{f}$ then the trait is the proportion of resources allocated to producing females (individuals destined to become workers or queens), if $\tau = \mathrm{q}$ then the trait is the proportion of resources allocated to producing queens from resources allocated to females. |
| $\nu_s^\circ$ | Neutral reproductive value of an individual of type $s \in \{\mathrm{q}, \mathrm{m}\}$. |
